# Gamma protocadherins in vascular endothelial cells inhibit Klf2/4 to promote atherosclerosis

**DOI:** 10.1101/2024.01.16.575958

**Authors:** Divyesh Joshi, Brian G. Coon, Raja Chakraborty, Hanqiang Deng, Pablo Fernandez-Tussy, Emily Meredith, James G. Traylor, Anthony Wayne Orr, Carlos Fernandez-Hernando, Martin A. Schwartz

## Abstract

Atherosclerotic cardiovascular disease (ASCVD) is the leading cause of mortality worldwide^1^. Laminar shear stress (LSS) from blood flow in straight regions of arteries protects against ASCVD by upregulating the Klf2/4 anti-inflammatory program in endothelial cells (ECs)^2–8^. Conversely, disturbed shear stress (DSS) at curves or branches predisposes these regions to plaque formation^9,10^. We previously reported a whole genome CRISPR knockout screen^11^ that identified novel inducers of Klf2/4. Here we report suppressors of Klf2/4 and characterize one candidate, protocadherin gamma A9 (Pcdhga9), a member of the clustered protocadherin gene family^12^. Pcdhg deletion increases Klf2/4 levels in vitro and in vivo and suppresses inflammatory activation of ECs. Pcdhg suppresses Klf2/4 by inhibiting the Notch pathway via physical interaction of cleaved Notch1 intracellular domain (NICD Val1744) with nuclear Pcdhg C-terminal constant domain (CCD). Pcdhg inhibition by EC knockout (KO) or blocking antibody protects from atherosclerosis. Pcdhg is elevated in the arteries of human atherosclerosis. This study identifies a novel fundamental mechanism of EC resilience and therapeutic target for treating inflammatory vascular disease.

## Introduction

ASCVD, characterized by fatty plaques within arterial walls, results from converging metabolic, inflammatory, and biomechanical factors including hypertension, hyperlipidemia, smoking, and age^3,6^. Atherosclerotic plaques form preferentially at curved or branched regions of arteries experiencing low, multi-directional shear stress from blood flow, termed disturbed shear stress (DSS)^3,6^. DSS is also the most reliable predictor of plaque erosion^2,8^ and plaque vulnerability to rupture^13^. Conversely, straight regions with high unidirectional laminar shear (LSS) suppress plaque formation mainly by upregulating the Kruppel-like transcription factors 2 and 4 (Klf2/4) in ECs. These two genes are generally co-regulated in ECs and have partially redundant gene targets and functions. Klf2/4 govern ∼70% of LSS induced anti-inflammatory and anti-thrombotic protective genes^4,5,7,9,10,14,15^. Extensive studies in mice using both EC specific knockout (ECKO) and transgenic overexpression identify Klf2/4 as potent mediators of resilience against a multiplicity of vascular conditions including atherosclerosis, pulmonary hypertension, and Covid19-mediated vascular dysfunction^15,16^. Reduced endothelial Klf2/4 expression is similarly associated with worsened cardiovascular outcomes in humans. Restoring high Klf2/4 expression may therefore protect against a range of inflammatory CVDs^17^. However, fundamental mechanisms regulating Klf2/4 expression remain unanswered.

Towards this goal, we carried out a genome wide CRISPR knockout screen using a GFP reporter driven by the human Klf2 promoter that identified ∼300 genes required for shear stress induction of Klf2^11^. Systematic analysis of the genes whose CRISPR KO reduced Klf2 (activators) identified a novel contribution from mitochondrial metabolism that synergizes with the established mechanism Mekk2/3-Mek5-Erk5 kinase cascade^11,18^. Unexpectedly, this screen uncovered an additional ∼160 genes whose KO increased Klf2 (suppressors). These genes are of great interest both to gain mechanistic insight into Klf2 regulation and as candidate therapeutic targets in vascular inflammatory diseases.

We selected Protocadherin gamma A9 (Pcdhga9) for further study. This gene is a member of the 22 gene Pcdhg subfamily, among the clustered protocadherin (cPcdh) family of homophilic adhesion receptors. Clustered protocadherins mediate a wide range of functions in the central nervous system including adhesion, signaling, and cell sorting^12,19^. Pcdhg has an essential role in neurons and its mis-expression is associated with neuronal disorders^20,21^, however, little is known about Pcdhg in other cell types. Here, we describe a role for the Pcdhg cluster in vasculature inflammation and ASCVD. Pcdhg functions by inhibiting the Notch pathway which is critical for transcriptional induction of Klf2/4 by LSS.

## Results

### Genome wide CRISPR screen identifies suppressors of hemodynamic Klf2 induction

To identify the mechanisms underlying mechano-regulation of Klf2, a genome wide CRISPR screen was performed using a *Klf2*:GFP reporter expressed in Mouse Aortic ECs (MAECs) and stimulated with LSS for 24h (Figures 1a, b, S1a). In addition to the ∼300 genes whose CRISPR KO decreased Klf2^11^, this screen also identified ∼160 Klf2 genes whose KO increased *Klf2*:GFP reporter levels (Figure S1b). Network analysis showed enrichment of pathways/processes that affect vascular and endothelial functions (Figure 1c), consistent with a central role for Klf2/4 in ECs.

**Figure 1.**
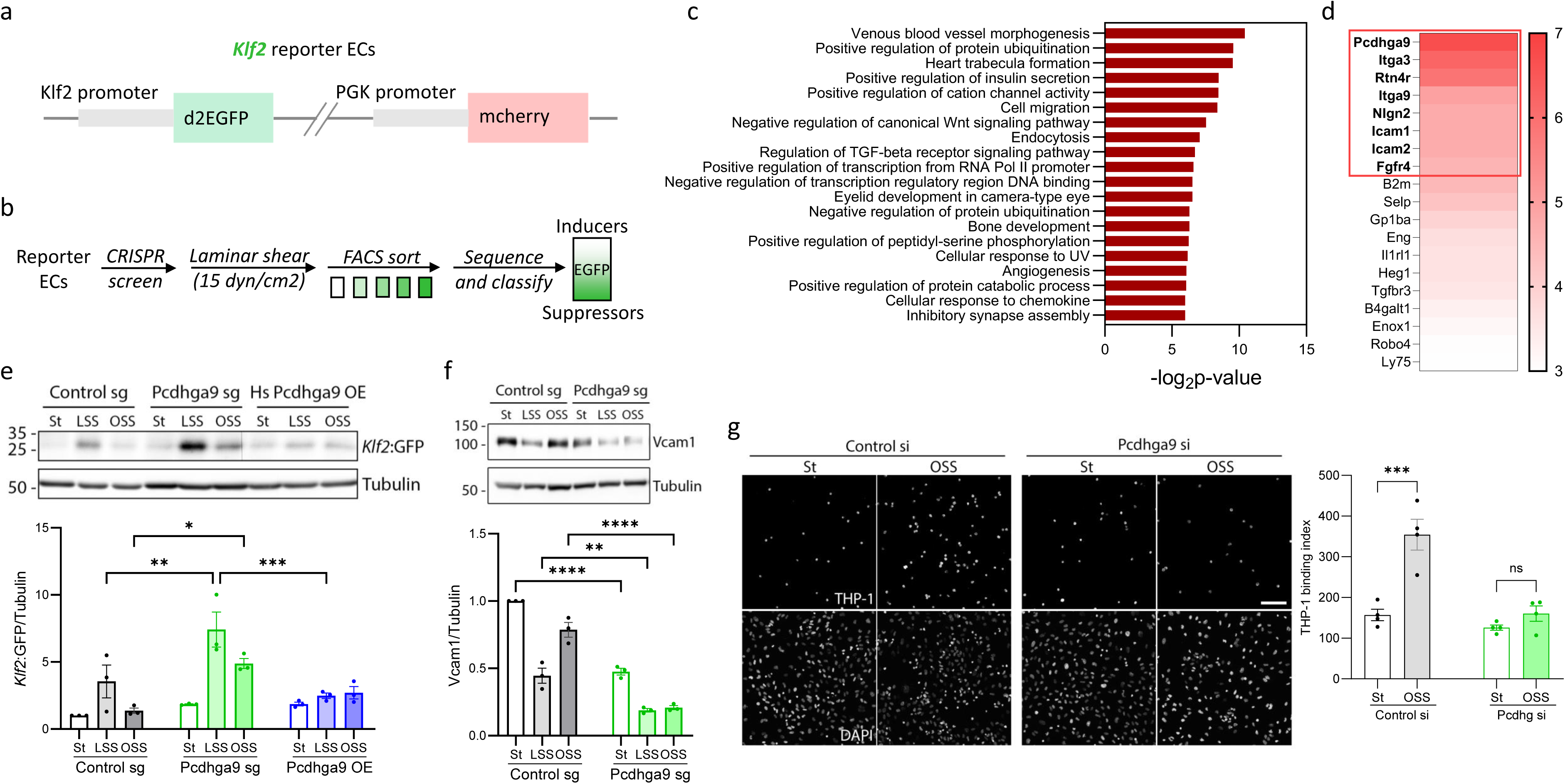
Genome wide CRISPR screen identifies Klf2 suppressors. (a) Schematic of the Klf2 reporter stably expressed in Mouse Aortic Endothelial Cells (MAEC). Destabilized EGFP (d2EGFP) is driven by the Klf2 promoter while mCherry is driven by a constitutive PGK promoter as an internal control. (b) Schematic of the genome wide CRISPR screen to identify modifiers (activators and suppressors) of Klf2 induction under laminar shear stress (LSS; 15 dyn/cm^2^ LSS for 16 h). (c) Functional categorization of the candidate Klf2 suppressors (cumulative z score >4, 160 hits) using Ingenuity Pathway Analysis (IPA, Qiagen). (d) candidate suppressors intersecting with ‘cell-surface exposed proteins on outer plasma membrane’ GO term to identify candidates amenable to function neutralization by blocking antibodies (red box: validated hits). (e) *Klf2*:GFP reporter MAECs were transduced with Cas9 plus Control sg, Pcdhga9 sg or with a human Pcdhga9 (Hs Pcdhga9) vector. Cells were exposed to static (St), LSS, or disturbed oscillatory shear stress (OSS) for 16 h and immunoblotted for *Klf2*:GFP (N=3). Graph: quantitation of *Klf2*:GFP normalized to Tubulin loading control. (f) Immunoblot for the inflammatory marker VCAM1 in Control or Pcdhga9 depleted MAECs exposed to St, LSS or OSS (N=3). Graph: quantitation of VCAM1 normalized to Tubulin loading control. (g) THP-1 monocyte binding to Control si or Pcdhga9 si Human Umbilical Vein Endothelial Cells (HUVECs) exposed to St or OSS for 16 h (N=4). Graph: quantitation of bound THP1 monocytes per field (THP1 binding index). Statistical analysis used one-way ANOVA. Scale bar: (g) 100 μm.

Reasoning that transmembrane proteins would be readily accessible for therapeutic approaches, we focused on the cell surface candidates (Figure 1d). Of these, the top 8 candidates were functionally verified using independent CRISPR sgRNAs by assaying for *Klf2*:GFP induction upon LSS and oscillatory shear (OSS), commonly used to model in vivo DSS (Figure 1d red box). The top candidate, Pcdhga9, was chosen for further study. We next examined induction of the *Klf2*:GFP in the MAEC reporter line under both LSS and OSS. CRISPR KO of Pcdhga9 increased *Klf2*:GFP in cells under LSS and did so even more strongly under OSS, where expression is normally low (Figure 1e). Conversely, OE of human Pcdhga9 suppressed reporter expression. Knockdown of Pcdhga9 using siRNA similarly increased Klf2:GFP which was reversed by OE of human protein (Figure S1c-e). Examining endogenous Klf2 in primary human umbilical vein endothelial cells (HUVECs) replicated this behavior, demonstrating conservation across species and EC types (Figure S1f). In ECs under pro-inflammatory OSS, Pcdhga9 depletion also suppressed the induction of the leukocyte adhesion receptor VCAM1 (Figure 1f) and the adhesion of THP1 monocytes (Figure 1g). Pcdhga9 thus restrains LSS-dependent induction of anti-inflammatory Klf2 and potentiates OSS-dependent pro-inflammatory activation.

### Protocadherin gamma (Pcdhg) gene cluster suppresses Klf2/4

Pcdhga9 is a member of the clustered protocadherin family, which belongs to the cadherin superfamily of cell-cell adhesion receptors. The clustered protocadherin genes, classified into alpha (a or α), beta (b or β), and gamma (g or γ) clusters, are organized in tandem in the genome. The mouse Pcdhg gene cluster comprises 22 genes (Figure 2a), several of which are expressed in ECs (this study). Exon 1 is unique, while exons 2-4 are shared by all 22 members allowing for targeting of the entire Pcdhg cluster by siRNA mediated silencing (Figure 2a, region enclosed between dashed lines). Silencing the Pcdhg gene cluster in HUVECs (Figures 2b, S2a) strongly induced endogenous *Klf2 and 4* mRNA and suppressed pro-inflammatory *E-selectin* (*Sele*) even under OSS, (Figures 2c, d, S2b). Depletion of the entire Pcdhg gene cluster suppressed adhesion of THP1 monocytes to ECs under pro-inflammatory OSS to the same extent as Pcdhga9 depletion (Figure 2e). Targeting Pcdhga9 or the entire cluster thus give indistinguishable results.

**Figure 2.**
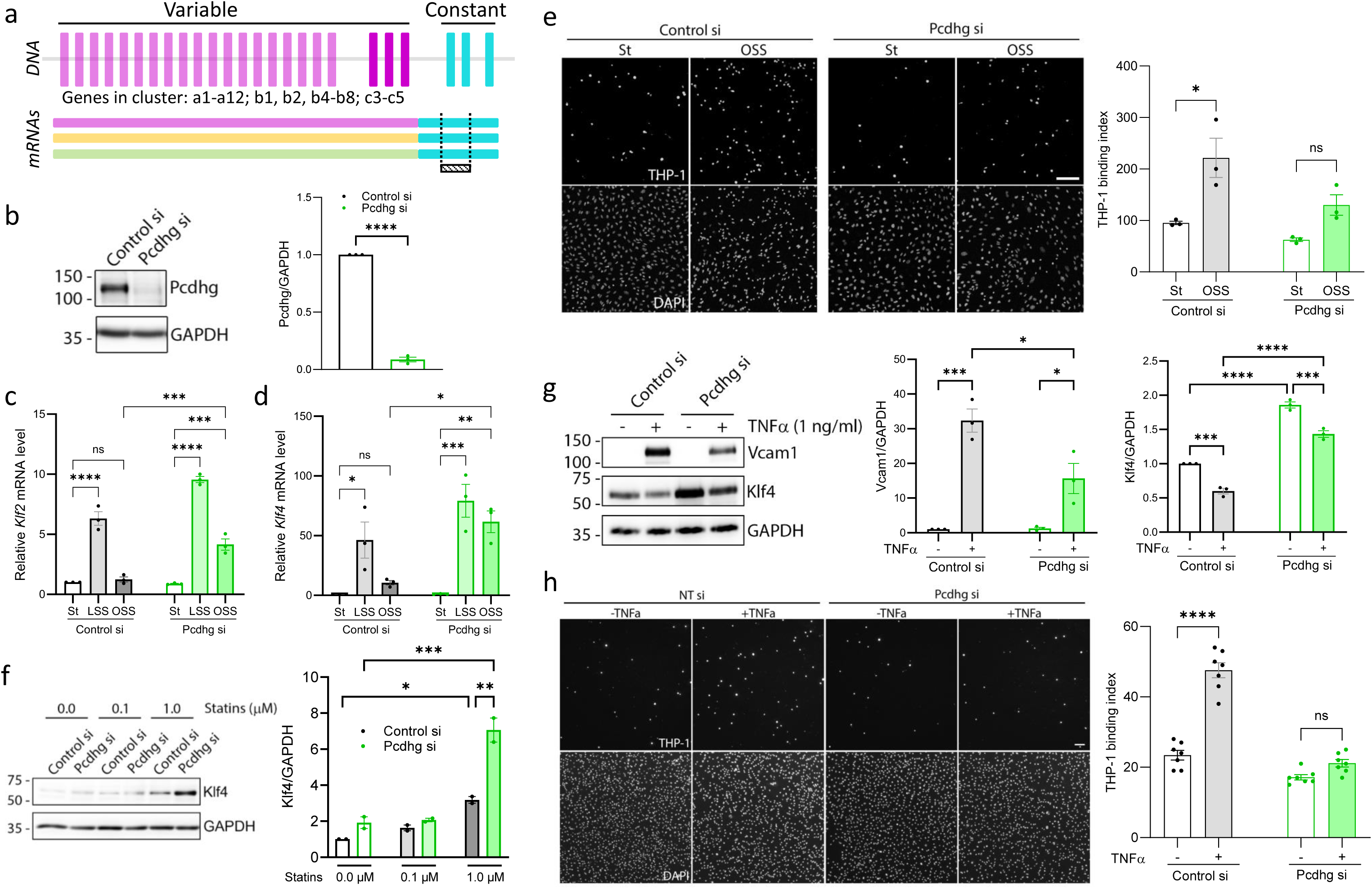
Protocadherin gamma (Pcdhg) gene cluster suppresses hemodynamic Klf2/4. (a) Top: schematic of Pcdhg gene cluster organization consisting of 22 genes with their unique/variable first exon (and individual promoter, not shown) and three common/constant 3’ exons that form the common cytoplasmic domain (CCD). Bottom: strategy for silencing (knocking down) the entire Pcdhg gene cluster with a siRNA targeting the common region (shaded bar and dashed lines). (b) Validation of siRNA mediated knockdown of the entire Pcdhg gene cluster (22 genes) in HUVECs by immunoblot with an antibody to the common domain (N=3). Graph: quantitation of Pcdhg levels normalized to GAPDH loading control. (c, d) qRT-PCR for *Klf2* (c) and *Klf4* (d) in control and Pcdhg-depleted HUVECs exposed to St, LSS and OSS for 16 h. (e) THP-1 monocyte binding to Control si or Pcdhg si HUVECs exposed to St or OSS for 16 h (N=3). Graph: quantitation of THP1 monocytes per field per condition (THP1 binding index). (f) Immunoblot for Klf4 in Control si or Pcdhg si HUVECs, treated with lovastatin (0.0, 0.1 or 1.0 μM) for 16 h. Graph: quantitation of Klf4 normalized to GAPDH loading control. (g) Immunoblot for VCAM1 and Klf4 in Control si or Pcdhg si HUVECs, untreated or treated with TNFα (1 ng/ml) for 16 h (N=3). Graphs: quantitation of VCAM1 and Klf4 normalized to GAPDH loading control with or without TNFα. (h) THP-1 monocyte binding to Control si or Pcdhg si HUVECs untreated or treated with TNFα (1ng/ml) for 16h (N=6). Graph: quantitation of THP1 monocytes per field per condition. Statistical analysis was carried out using one-way ANOVA. Scale bar: 100 μm, (h) 250 μm.

In addition to LSS, Klf2/4 are also induced by statins^22^, the cholesterol lowering drugs used as first line therapy for patients at risk of ASCVD; an effect that is believed to contribute to their therapeutic benefits. To test for interactions between Pcdhg and statins, control or Pcdhg-depleted HUVECs were treated with lovastatin for 16 h. Pcdhg depletion greatly amplified statin induction of Klf4 (Figure 2f) showing clear synergy. On the other hand, the pro-inflammatory cytokine TNFα suppresses Klf2/4 and induces pro-inflammatory genes including, the leucocyte adhesion receptor, VCAM1. Pcdhg depletion strongly suppressed TNFα-induced VCAM1 expression and subsequent adhesion of THP1 monocytes to ECs, and alleviated Klf4 suppression by TNFα (Figure 2g, h). Pcdhg depletion significantly reduced induction of VCAM1 even at the highest doses of TNFα tested (Figure S2c). Pcdhg loss thus suppressed EC inflammatory activation in multiple contexts.

### Pcdhg endothelial knockout protects against atherosclerosis

EC Klf2/4 are essential for vascular resilience against inflammatory CVDs, including atherosclerosis. Importantly, elevating Klf4 in ECs protects against ASCVD, demonstrating its sufficiency for atheroprotection^23^. To test if effects of Pcdhg seen in culture are conserved in vivo, *Pcdhg^fcon^*^3^ (Pcdhg constant exon 3 floxed and tagged with gfp) mice^24,25^ were crossed with constitutive *Cdh5Cre* mice to delete the entire Pcdhg cluster in ECs (Pcdhg EC knockout or ECKO) (Figure 3a). Homozygous Pcdhg ECKO pups were obtained at the expected Mendelian frequency and were phenotypically normal, showing that Pcdhg expression in ECs, unlike in neurons, is not essential for development^26,27^ (Figures 3b, S3a-c). These mice were thus further analyzed.

**Figure 3.**
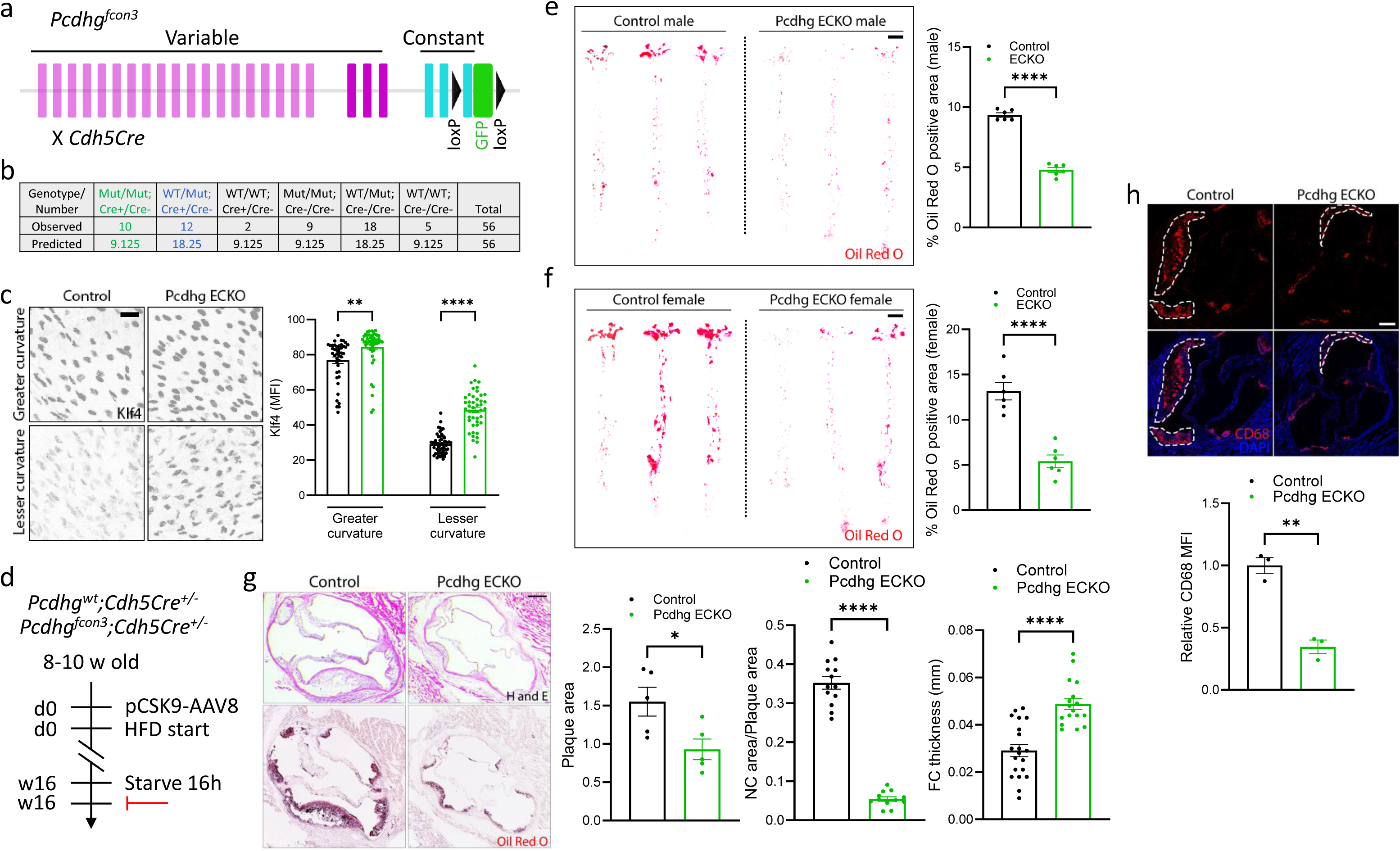
Pcdhg endothelial knockout protects against atherosclerosis. (a) Schematic showing Pcdhg floxed mouse (*Pcdhg^fcon^*^3^ mouse line^24,25^) and generation of Pcdhg endothelial knockout (ECKO) by crossing *Pcdhg^fcon^*^3^ with *Cdh5Cre*. The third constant exon is tagged with GFP and floxed. Cre-mediated recombination excises the third constant exon creating a premature stop codon leading to non-sense mediated decay. (b) Analysis of progeny genotype showing Pcdhg ECKO at the expected Mendelian ratio. (c) Aortic arch from Control or Pcdhg ECKO adult mice immunostained for Klf4 and examined *en face* (N=3). Geater curvature denotes the regions of laminar shear while lesser curvature denotes the regions of low, disturbed shear. Graph: quantitation of the mean fluorescence intensity (MFI) per cell. (d) Schematic of atherosclerosis study. 8-10 week-old Control or Pcdhg ECKO mice were injected with pCSK9-Adeno Associated Virus 8 (AAV8) and maintained on High Fat Diet (HFD) for 16 weeks, starved overnight, and analyzed for blood lipids and vessel histology. (e, f) Whole aortas were stained with Oil Red O to mark lipid-rich regions and imaged *en face* (N=6 each sex). Graph: quantitation of percent Oil Red O positive area for males (e) and females (f). (g) Sections from aortic roots were stained with Hematoxylin and Eosin (H&E) (N=5) or Oil Red O. Graphs: quantitation of atherosclerotic plaque area, necrotic core (NC) area and fibrous cap (FC) thickness. (h) Atherosclerotic plaques (dotted area) in aortic root sections were stained for the macrophage/monocyte marker CD68 (N=3). Graph: quantitation of CD68 MFI. Statistical analysis was carried out using Student’s t-test. Scale bar: (c) 50 μm, (e, f) 500 μm, (g, h) 200 μm.

The aortic arch contains the athero-resistant greater curvature that is under LSS and the athero-susceptible lesser curvature that is under DSS, offering a well-established model. Examination *en face* revealed modestly increased Klf4 expression in Pcdhg ECKO ECs in the athero-resistant greater curvature and a markedly larger increase in the athero-susceptible lesser curvature, compared to controls (Figure 3c). We next induced hyperlipidemia by injection of AAV8-PCSK9 and feeding a high fat diet (HFD) for 16 weeks (Figure 3d)^28,29^. Pcdhg ECKO mice showed reduced atherosclerotic plaques as observed by Oil Red O staining in aortas from both males and females (N=6, each sex) (Figure 3e, f). Examination of the aortic root using hematoxylin, and eosin (H&E) and Oil Red O-stains showed reduced atherosclerotic plaques with drastically smaller necrotic cores (NC), thicker fibrous caps (FC) (Figure 3g), and greatly reduced macrophage/monocyte content in Pcdhg ECKO (Figure 3h). No differences were observed in blood lipids (triglycerides, cholesterol, HDL-C) (Figure S3d-f) or body weights (Figure S3g). Pcdhg expression in ECs thus exerts a pro-atherogenic role.

### Nuclear Pcdhg intracellular domain suppresses Klf2/4

To investigate the molecular mechanism by which Pcdhg suppresses Klf2/4, we performed a structure-function analysis using the alternative splicing of Pcdhg cluster members to define domain boundaries (Figure 4a). Each Pcdhg protein is organized into an extracellular domain (ECD) containing 6 cadherin domains involved in homophilic or heterophilic cis/trans interactions, a transmembrane (TM) domain governing its membrane localization, and an intracellular domain (ICD) involved in intracellular trafficking, surface delivery, localization, and signaling. The ICD is sub-divided into a variable C-terminal domain (VCD) and a constant C-terminal domain (CCD). The conserved CCD is encoded by the three 3’ exons shared by all 22 members (Figure 4a, b).

**Figure 4.**
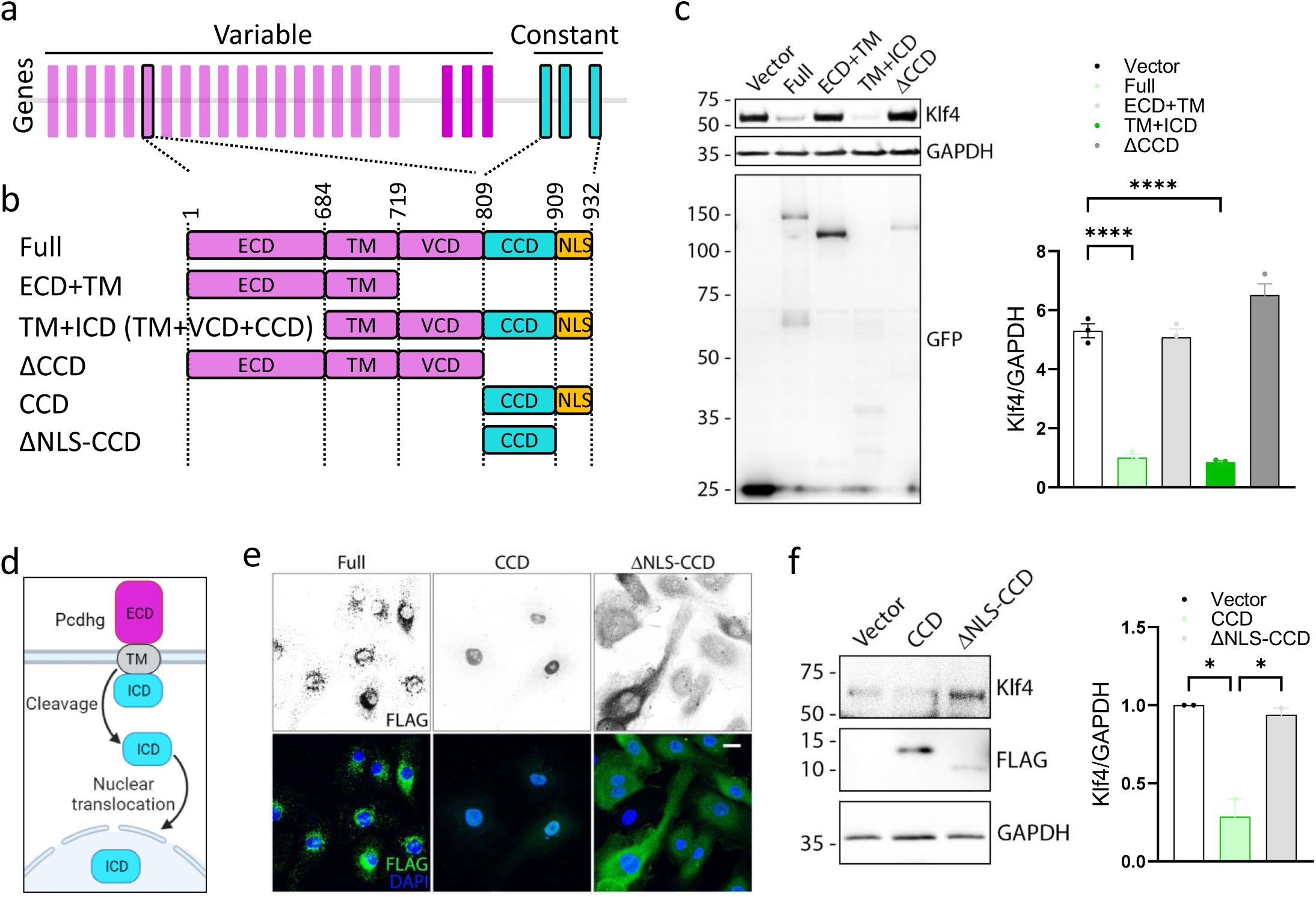
Signaling via nuclear ICD is necessary and sufficient for suppression of Klf2/4. (a, b) Gene organization of Pcdhg cluster with the three 3’ exons (blue), coding for the Carboxy terminal constant domain (CCD) shared by all 22 members. (b) Mutant constructs. (c) HUVECs transfected with indicated constructs and treated with LSS for 16h. Immunoblotting confirms expression of mutants and effects on endogenous Klf4 (N=3). Graph: quantitation of Klf4 normalized to GAPDH loading control. The C-terminal GFP-tagged mutants were expressed at comparable levels (GFP immunoblot). (d) Schematic showing cleavage and nuclear translocation of the Pcdhg ICD. (e) HUVECs expressing additional mutants (C-terminal FLAG-tagged) were immunostained for FLAG and counterstained with DAPI to mark nuclei (N=3). (e) HUVECs expressing the mutants were exposed to LSS and immunoblotted for Klf4 (N=3). Graph: quantitation of Klf4 normalized to GAPDH loading control. The C-terminal FLAG-tagged mutants were expressed at comparable levels (FLAG immunoblot). Full: Full length; ECD: Extracellular domain; TM: Transmembrane domain; VCD: Variable C-terminal domain; CCD: Constant C-terminal domain; ΔCCD: Deleted CCD; ICD: Intracellular domain (VCD+CCD); NLS: Nuclear Localization Signal; ΔNLS: Deleted NLS. Statistical analysis used Student’s t-test. Scale bar: (e) 20 μm. *p < 0.05, **p < 0.01, ***p < 0.001, ****p < 0.0001.

Overexpression of Pcdhga9 suppressed Klf2/4 (Figures 1e, S1e). We thus used this assay to identify functional domains. Full length (Full), ECD+TM, TM+ICD and CCD-lacking (ΔCCD) mutants (Figure 4b) were expressed in HUVECs, which were treated with LSS and OSS and assayed for Klf4 protein. The mutants were expressed at comparable levels and localization was analyzed by GFP fluorescence (Figures 4c, S5a, b). While the full length localized to both cell-cell junctions and intracellular vesicles, ECD+TM and ΔCCD were more junctional as expected (owing to reduced internalization in the absence of the CCD), while TM+ICD was punctate and distributed evenly (Figure S5b). Klf4 Western blotting revealed that the CCD is essential for Klf2/4 suppression, whereas the ECD is dispensable (Figures 4c, S5a). Further experiments thus focused on the CCD.

Unlike classical cadherins, Pcdhg members are processed by cleavage, which releases the ICD, which translocates to the nucleus ^30^ via its nuclear localization signal (NLS) (Figure 4b, d). To test the role of cleavage and nuclear translocation, Flag-tagged version of CCD and an NLS-deleted CCD mutant (ΔNLS-CCD) were expressed in HUVECs, stimulated with LSS, and Klf4 levels assayed (Figure 4b). The mutants were expressed at comparable levels and localized as expected, with the full-length present in the secretory system and plasma membrane, the CCD present mainly in the nucleus and the ΔNLS-CCD present in the cytoplasm (Figure 4e). The CCD suppressed Klf4 whereas the ΔNLS mutant was ineffective (Figure 4f). Interestingly, the CCD is very highly conserved across species, supporting its critical role (Figure S5c). Pcdhg thus signals via cleavage and nuclear translocation of its common cytoplasmic regions.

### Pcdhg regulates Klf2/4 via the Notch pathway

For an unbiased assessment of the genes and processes regulated by Pcdhg in ECs, we performed bulk RNAseq analysis of cells under OSS (where Pcdhg had the greatest effect). In addition to control si and Pcdhg si HUVECs, Pcdhg+Klf2/4 triple si was included to identify Klf2/4-independent effects (Figure 5a, b). Upstream regulatory pathway analysis of these Pcdhg-dependent and Klf2/4-independent DEGs identified Notch as the most over-represented pathway (Figure 5c), confirmed by strong upregulation of known Notch target genes in ECs (Figure 5d). Notch1-4 family transmembrane receptors are critical for EC functions including determination of arterial identity and vascular stability, with Notch1 being the main isoform. Binding of Notch ligands such as DLL4 to receptors triggers proteolysis by γ-secretase, releasing the Notch intracellular domain (NICD), which translocates to the nucleus and binds the transcription factor RBPJ to induce transcription of target genes^31^. Interestingly, LSS both activates Notch and induces Klf2/4, both of which stabilize and protect vessels against inflammation and atherosclerosis^32–35^, though effects of Notch and Klf2/4 have not been linked. We found that Pcdhg depletion led to increased levels of the cleaved, activated Notch1 intracellular domain [NICD (Val1744)] (Figure 5e). Notably, the promoters for both human and mouse Klf2 and Klf4 contain previously unappreciated canonical Notch-RBPJ binding sites (Figure S6a, b). To test the role of Notch in Klf2/4 induction by LSS, RBPJ was blocked (small molecule inhibitor RIN1 or RBPJi) or depleted (RBPJ si), with or without Pcdhg depletion. ECs under LSS were then examined. The increase in Klf4 levels upon Pcdhg si was completely prevented by inhibition of the Notch pathway (Figure 5f). We conclude that Pcdhg regulates Klf4 via Notch activation.

**Figure 5.**
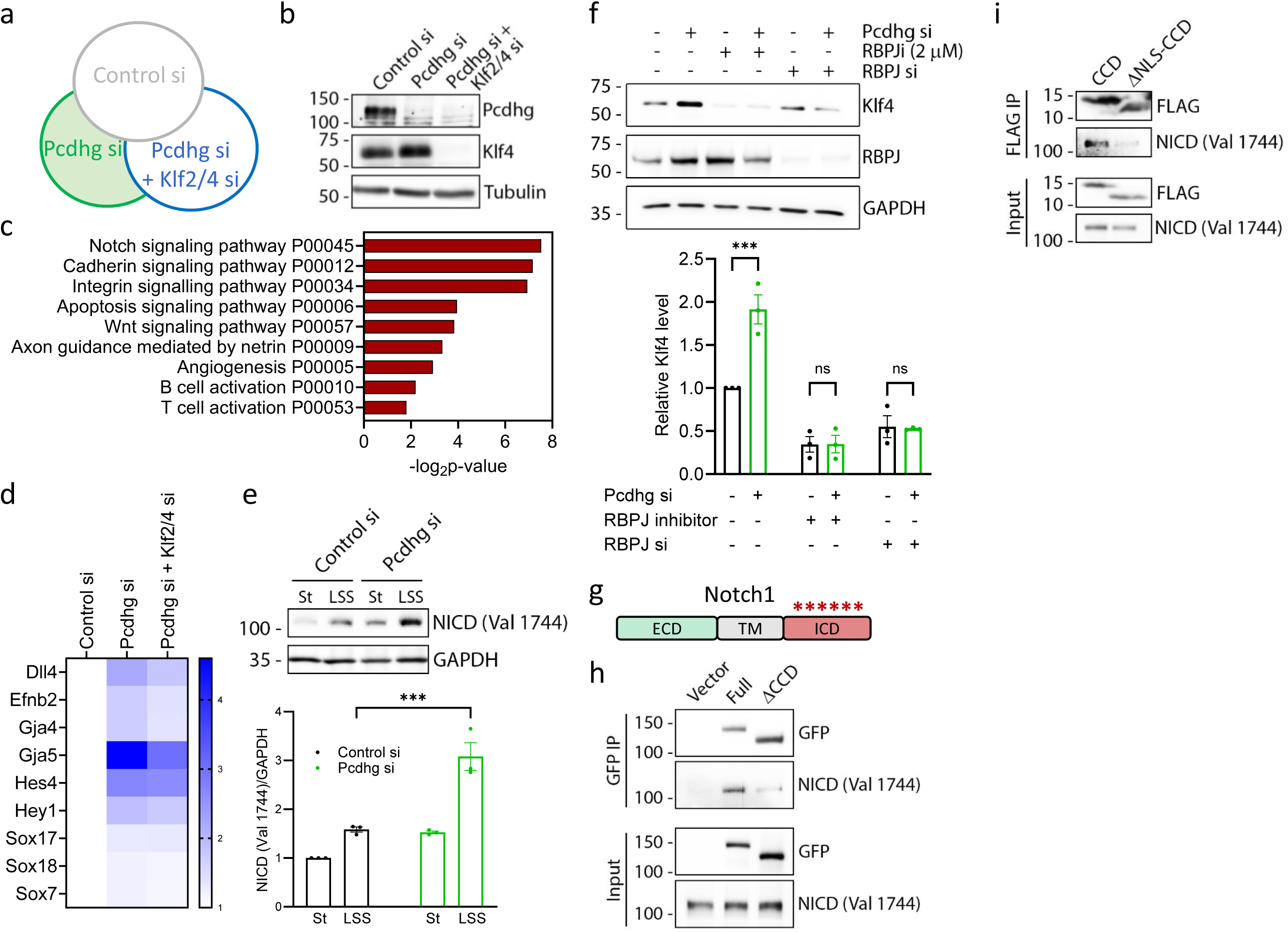
A physical and functional Pcdhg-Notch axis regulates hemodynamic Klf2/4. (a) Schematic of bulk RNAseq analysis of <u>Control si</u>, <u>Pcdhg si</u> and <u>Pcdhg+Klf2/4 triple si</u> HUVECs to identify Pcdhg-dependent, Klf2/4 independent DEGs (green shaded area). (b) Immunoblot validation of Pcdhg and Klf4 depletion. (c) Upstream regulatory pathway/process analysis from the DEGs using Enrichr. (d) Heat map of upregulated Notch target genes in Pcdhg si and Pcdhg si+Klf2/4 si compared to Control si from the RNAseq analysis. (e) Control si or Pcdhg si HUVECs were exposed to St or LSS for 2h and immunoblotted for the cleaved Notch1 Intracellular Domain (NICD Val1744) (N=3). Graph: quantitation of NICD Val1744 normalized to GAPDH loading control. (f) The NICD-dependent transcription was blocked with RIN1, a pharmacological inhibitor of Notch-RBPJ interaction (RBPJi) or RBPJ siRNA (RBPJ si). HUVECs were exposed to LSS for 16h and immunoblotted for Klf2 and Klf4 (N=3). Graph: quantitation of Klf2 and Klf4 normalized to GAPDH loading control. (g) Schematic showing Notch1. ECD: Extracellular domain; TM: Transmembrane domain; ICD: Intracellular Domain. The asterisks (*) show the NICD peptides identified in the Mass Spectrometry analysis of the Ips of full but not the ΔCCD Pcdhg mutant. (h, i) Co-IP of Pcdhg and NICD Val1744. Inputs used as loading controls for IPs. (h) HUVECs expressing full or ΔCCD Pcdhg (C-term GFP-tagged) were immunoprecipitated (IP) with GFP nanobody beads and immunoblotted for NICD Val1744 (N=3). Vector alone used as negative control for the IP. (i) HUVECs expressing CCD or ΔNLS-CCD mutant (C-term FLAG-tagged) were immunoprecipitated (IP) with FLAG antibody beads and immunoblotted for NICD Val1744 (N=3). Statistical analysis was carried out using Student’s t-test or one-way ANOVA. *p < 0.05, **p < 0.01, ***p < 0.001, ****p < 0.0001.

Structure-function analysis identified CCD of Pcdhg as the functional domain that inhibits Klf2/4 induction. To identify interacting proteins, we did a proteomic analysis of immunoprecipitates (IP) from full length Pcdhga9 and the ΔCCD mutant, looking for binding partners that required the CCD. Notch1 peptides were detected in the IPs of full length Pcdhga9, but not the ΔCCD mutant or vector alone. Sequence analysis showed that these Notch1 peptides came from the NICD region (Figures 5g asterisks, S6c), consistent with functional effects. IP and immunoblot analysis showed that in addition to full length Pcdhg, the CCD fragment but not the ΔNLS-CCD physically associated with the NICD (Figure 5 h-i). Taken together, these data support a model in which the PICD physically associates with NICD to suppress Notch signaling and its downstream target Klf2/4.

### Pcdhga9 blocking antibody in experimental atherosclerosis

Pcdhg was chosen as a target in part because its cell surface localization makes it amenable to inhibition by antibodies or other cell-impermeant reagents. Towards this goal, we purified Pcdhga9 ECD protein, which was used both for generating monoclonal antibodies (mAbs) and for developing a cell adhesion assay (Figures 6a, S7a). When cells were plated in 96 wells coated with Pcdhga9 ECD, cells adhered over time with minimal adhesion to uncoated wells (Figure 6b), demonstrating specificity.

**Figure 6.**
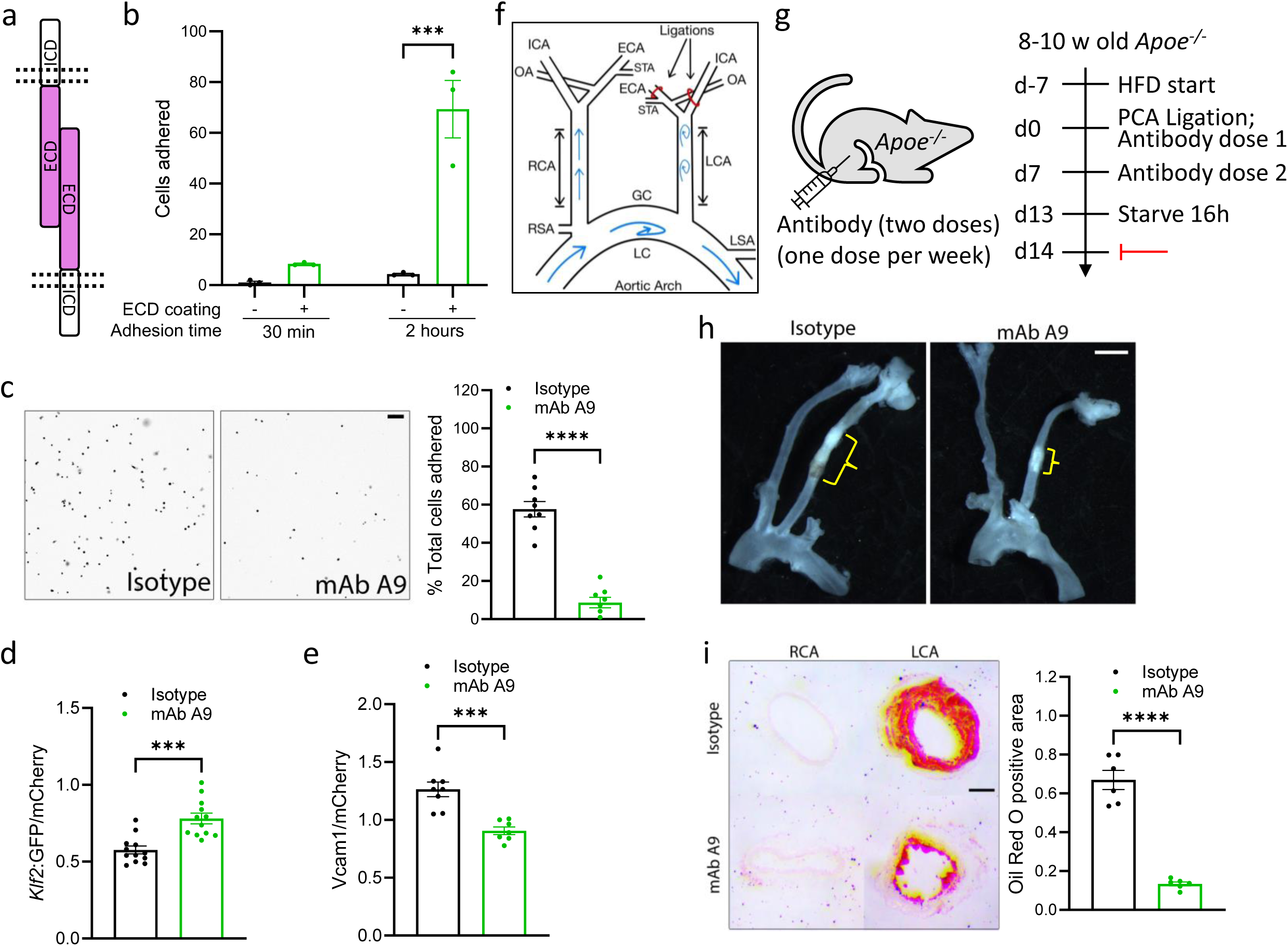
Pcdhga9 blocking antibody in experimental atherosclerosis. (a) Schematic of homophilic adhesion assay and function blocking monoclonal antibody (mAb) generation from Pcdhga9 ECD. (b) Pcdhga9 ECD was immobilized in 96 wells and percent MAECs that adhered measured (N=3). (c) Adhesion of MAECs to ECD in the presence of Isotype control or function-blocking mAb A9 (N=4). Graph: quantitation of percent total cells adhered to ECD. (d, e) *Klf2*:GFP reporter MAECs with Isotype control or mAb A9 tested for *Klf2*:GFP expression after 16h LSS (N=12) or immunostained for VCAM1 after OSS for 16 h OSS (N=7). Graphs: quantitation of *Klf2*:GFP and VCAM1 levels normalized to mCherry internal control. (f) Schematic of Partial Carotid Artery (PCA) Ligation model of accelerated atherosclerosis. LCA: Left Carotid Artery; RCA: Right Carotid Artery. (g) Experimental design. Antibodies were administration via intraperitoneal (IP) injection into in Apoe^-/-^ mice. (h) Whole mount brightfield image showing branching of LCA and RCA from aorta, with atherosclerotic plaque (yellow brackets) visible in the ligated LCA. (i) LCA and RCA sections were stained with Oil Red O to identify for lipid-rich atherosclerotic plaque (N=6). Graph: quantitation of Oil Red O positive area fraction. Statistical analysis was carried out using Student’s t-test. Scale bar: (c, i) 100 μm, (h) 1 mm. *p < 0.05, **p < 0.01, ***p < 0.001, ****p < 0.0001.

Rat hybridomas that showed the strongest positive signals in ELISA assays against Pcdhga9 ECD (Figure S7a,b) were purified and tested in the homophilic adhesion assay^36^. Antibodies A9, B1 and B4 strongly blocked adhesion (Figures 6c, S7c). Antibody specificity was verified by Western blotting using purified ECD-GST or ECD alone (Figure S7d). Next, these function blocking mAbs were tested for their effect on Klf2/4 upon LSS, and VCAM1 upon OSS. Blocking antibodies increased flow-induction of the *Klf2:*GFP reporter and decreased VCAM1, normalized to the internal mCherry control (Figures 6d, e, S7e, f). mAb A9 showed the strongest effect on adhesion to ECD, Klf2 induction upon LSS, VCAM1 induction upon OSS and the highest signal to noise ratio in immunofluorescence assay (Figure S7g). Hence, mAb A9 was purified in bulk and studied further.

A9 specificity was verified by Western blotting of cell lysates from control vs Pcdhg or Pcdhga9 knockdown MAECs (Figure S8a), with purified ECD as a positive control. We therefore examined its efficacy in a model of experimental atherosclerosis in mice. Because A9 is a rat monoclonal, we avoided immune recognition and clearance of the rat IgG by studying accelerated atherosclerosis model using partial carotid artery (PCA) ligation in *Apoe^-/-^* mice where lesions develop within 1 week (Figure 6f)^29,37,38^. Mice were maintained on HFD for a total of 3 weeks (1 week prior to ligation and 2 weeks post ligation). A9 half-life, determined by IP injecting 2 μg of isotype control or mAb A9 per mouse and measuring blood plasma levels, was ∼8 days (Figure S8b). Hence, *Apoe^-/-^* mice on HFD were subject to surgery and A9 vs control IgG was injected once per week for two weeks as described in Methods (Figure 6g). The operated Left Carotid Artery (LCA) was then compared to the unligated Right Carotid Artery (RCA) as an internal control. Treatment with mAb A9 strongly reduced plaque in this acute model of ASCVD as seen by whole mount carotid preparations (Figure 6h, brackets) and verified by staining carotid sections with Oil Red O to detect lipid-rich plaques (Figure 6i). The reduction in lumen diameter was also limited by A9 (Figure S8c). No difference was observed in blood lipids (triglycerides and cholesterol) (Figure S8d, e). Taken together, these results identify Pcdhg as a novel therapeutic target in atherosclerosis.

### Pcdhg expression in atherosclerosis

Lastly, we analyzed levels of Pcdhg in human arteries. Staining for Pcdhg using an antibody against the CCD showed ∼3× higher expression of Pcdhg in endothelium (marked by Erg) from CVD donors compared to healthy, age-matched donors (Figure 7a). Atherosclerotic mouse arteries also showed elevated Pcdhg staining (Figure S9a). Coronary arteries from 3 asymptomatic elderly donors stained for Pcdhg showed ∼4× higher signal in the regions of plaque compared to regions of the same artery without evident plaque (Figure 7b). Antibody specificity was verified by Western blots using control and Pcdhg si HUVECs (Figure 2b) and staining of control vs. Pcdhg ECKO retinas (Figure S9b). Pcdhg is thus upregulated in endothelium from ASCVD.

**Figure 7.**
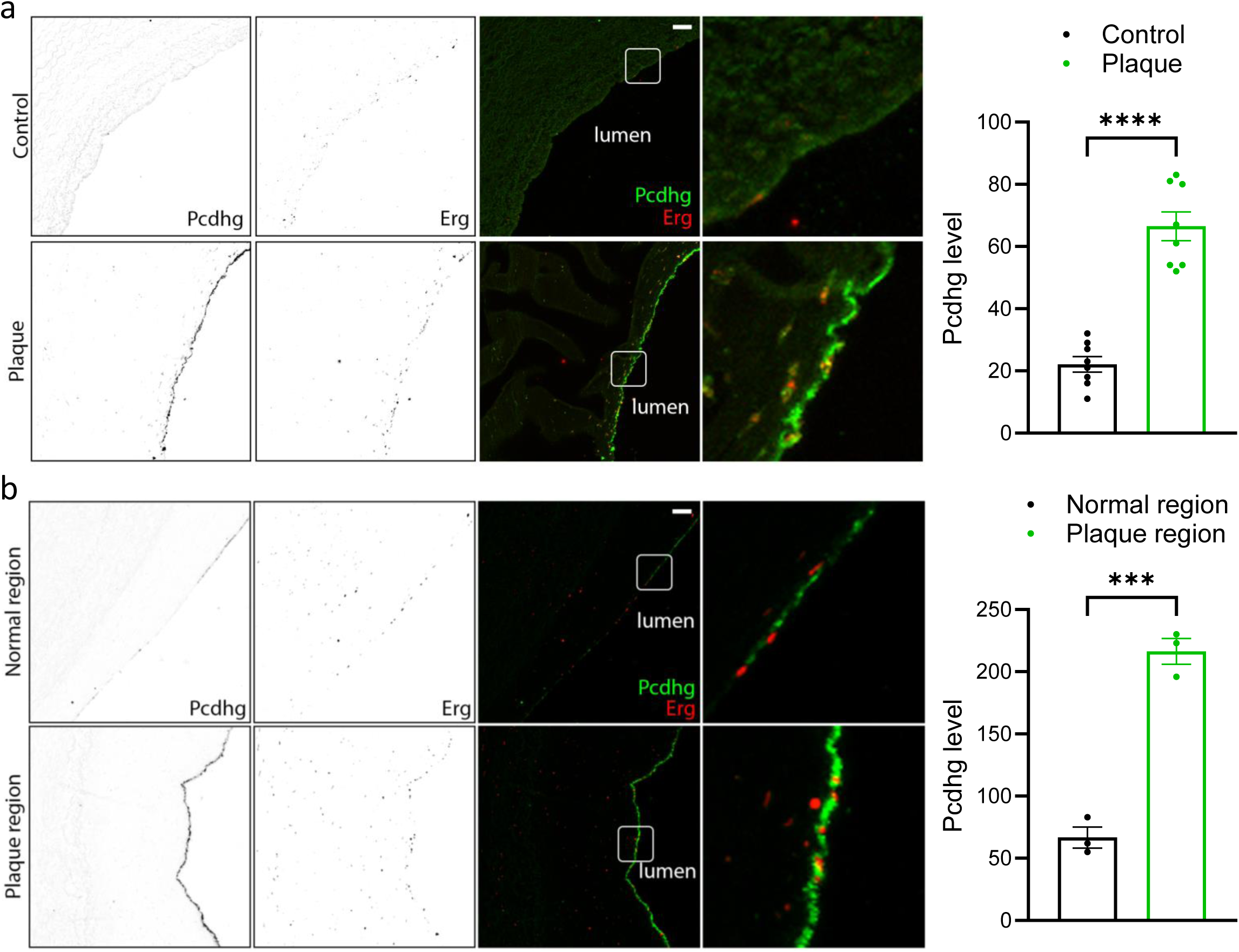
Pcdhg in human atherosclerosis. (a) Artery sections from human CVD patients and healthy controls stained for Pcdhg and for Erg to mark ECs (N=8). Graph: quantitation of Pcdhg level in ECs. (b) Human coronary artery sections from elderly donors stained for Pcdhg levels, comparing Plaque region to segments of the same artery without evident plaque (Normal region). Sections were stained for Erg to mark ECs (N=3). Graph: quantitation of Pcdhg staining intensity in ECs. Statistical analysis used Student’s t-test. Scale bar: (a, b) 50 μm. *p < 0.05, **p < 0.01, ***p < 0.001, ns (not significant) > 0.05.

## Discussion

ECs play a pivotal role in vascular physiology and pathology via functions range from regulating vessel diameter to immune responses to nutrient transport. Cell-cell adhesions are a locus of EC signaling and function, including solute transport, leukocyte trafficking, growth control and shear stress signaling. This study is based on a genome wide CRISPR knockout screen that identified ∼160 novel genes that suppress Klf2. We selected the strongest cell surface localized Klf2 suppressor, Pcdhga9, a member of the Pcdhg family of adhesion receptors, which has been extensively studied in neuronal systems^12,20,21,24,26,27,39,40^. However, only two studies report roles for another Pcdhg family member (Pcdhgc3) in the vasculature, with effects on EC tube formation, migration, and permeability in vitro, suggestive of a role in angiogenesis^41,42^. We found that Pcdhg is a pro-inflammatory gene that suppresses Klf2/4 through a pathway that involves its cleavage and release of the intracellular domain, which translocates to the nucleus and suppresses Notch1-dependent transcription. EC deletion of Pcdhg in mice increases Klf2/4 levels and protects against ASCVD without apparent effects on development or viability. Antibodies that block homophilic Pcdhg adhesion also increase Klf2/4 and limit experimental atherosclerosis. Within the vasculature, Pcdhg is expressed primarily in ECs, with marked increases in atherosclerotic regions and low expression in other cell types.

Analysis of Pcdhg-dependent genes identified the Notch pathway as a major downstream target. Notch1 limits angiogenesis in development, and promotes arterial identity and limits atherosclerosis in adults, thus was further investigated. LSS induction of both Klf2/4 transcription and Notch activation are well established, both of which promote vascular stability^32–35^, but an interconnection has not to our knowledge been reported. Mechanistic data in this study propose Notch as a direct inducer of Klf2/4 via consensus sites in their promoters/enhancers, which is suppressed by Pcdhg, which is upregulated in inflammatory settings.

While this study focused on atherosclerosis, vascular inflammation driven by EC activation is a major contributing factor in many additional diseases including pulmonary arterial hypertension, venous thrombosis, and auto-immune disease to name a few^12,19^. The CANTOS clinical trial identified benefits of immune suppression, but these were balanced by increased deaths from infections^43^. We suggest that targeting endothelial inflammatory activation rather than the immune system offers a path to limiting vascular inflammation without compromising host pathogen defense. Further work to fully elucidate mechanisms of Pcdhg expression, activation and contributions to inflammatory disease are thus important directions for future work. Additionally, the remaining 159 hits whose CRISPR KO increase Klf2 levels offer a rich source of new therapeutic targets.

## Materials and methods

### Primary cells, cell lines, and cell culture reagents

The PyMT-immortalized Mouse Aortic ECs that express the *Klf2* promoter reporter (*Klf2*:GFP MAEC) was described previously^11^. MAECs were maintained in complete EC Medium (Cell Biologics, M1166)^44^. HUVECs obtained from Yale Vascular Biology and Therapeutics Core were pooled from 3 donors. They were screened for the absence of pathogens, maintained in EGM2 Endothelial Cell Growth Medium (Lonza, CC-3162) and used at passages 2-5. All cells were routinely screened for the absence of mycoplasma. SiRNA transfection was performed with Lipofectamine RNAiMAX (ThermoFisher, 13778150) in Opti-MEM medium (ThermoFisher, 31985070) using ON-TARGET plus SMARTpool siRNAs from Horizon Discovery (Dharmacon). For lentiviral transduction, HEK293T cells were transfected with lentiviral vectors along with pVSV-G (Addgene, #138479) and psPAX2 (Addgene, #12260) packaging plasmids using lipofectamine 2000 (ThermoFisher, 11668019) following the manufacturer’s instructions. Supernatants were collected 48-96 h after transfection and filtered through a 0.45 μm low-protein binding filter. Primary HUVECs were infected with lentivirus for 24h, then the medium replaced with EMG2 medium. For monocyte adhesion assays, THP-1 labeled with CellTracker Deep Red (ThermoFisher, C34565) were resuspended in HBSS supplemented with 1 mM Ca2+, 0.5 mM Mg2+, and 0.5% BSA, added to the slides with HUVECs, incubated for 20 min at 37°C, washed 3× in HBSS and fixed with 3.7% formaldehyde. Cells were counterstained with DAPI before fluorescence imaging. Drugs used were Lovastatin (Sigma, 438185), TNFα (PeproTech, 300-01A) and RIN1 (RBPJ Inhibitor-1) (Selleckchem, SS3376). All antibodies were validated in knockdown/knockout depletion by immunofluorescence and/or immunoblotting. The details of antibodies, plasmid vectors, sgRNA sequences and primers are present in supporting information.

### CRISPR library screen and Klf2 suppressor (gain-of-function) phenotype determination

CRISPR library screening, library preparation and next-generation sequencing was described previously^11^. Briefly, immortalized *Klf2*:GFP reporter MAECs were infected with genome wide CRISPR library of ∼160,000 single guide (sg) CRISPR RNAs, treated with 15 dyn/cm^2^ LSS for 16 hours, FACS sorted based on *Klf2*:GFP reporter levels with subsequent analysis of sgRNAs in the high *Klf2*:GFP gate by next-generation sequencing on HiSeq2500 (Illumina). Klf2 suppressor (gain-of-function) phenotype was determined as done previously for the loss-of function phenotype determination ranking genes based on the cumulative z-score from 3 highest scoring unique sgRNAs^11^.

### Shear stress stimulation

All shear stress experiments, unless otherwise indicated, were performed in parallel plate flow chambers and perfused within a pump and environmental control system as described^45^. Briefly, cells were seeded at 70–90% confluency on 10 μg/ml Fibronectin-coated glass slides for 48-72 h in complete medium. For shear stimulation, slides were mounted in custom-made 25×55 mm parallel plate shear chambers with 0.5 mm thick silicone gaskets and stimulated with 15 dynes/cm^2^ for LSS or 1 ± 4 dynes/cm^2^ for OSS in complete media. The media was maintained at 37°C and 5% CO_2_ with a heat gun and humidified bubbler, respectively. The orbital shaker method was used for mAb functional testing in vitro (Figures 6d, e, S7f, g). Well radius was divided into three equal parts, with the outermost part representing pulsatile LSS region (for measuring *Klf2:GFP* levels) and the innermost part representing disturbed shear region (for measuring VCAM1 levels).

### Animals and tissue preparation

The use of all proposed techniques followed Yale Environmental Health and Safety (EHS) regulations, and all animal work was approved by Yale Institutional Animal Care and Use Committee (IACUC) and Yale Animal Resource Center (YARC). Mice were maintained in a light-controlled and temperature-controlled environment with free access to food and water and all efforts were made to minimize animal suffering. *Pcdhg^fcon^*^3^ mice^24,25^ and *Cdh5Cre* mice^46^ are described elsewhere. *Pcdhg^fcon^*^3^ mice were a kind gift from Julie Lefebvre, University of Toronto, Canada. All mice in this study were on the C57BL/6J background. *Pcdhg^fcon^*^3^ and *Cdh5Cre* mice were maintained and bred as heterozygotes. Euthanasia was performed by an overdose of isoflurane inhalation and death was confirmed by subsequent cervical dislocation and/or removing vital organs and/or opening the chest cavity verified by the absence of cardiovascular function. Mice were perfused through the left ventricle with PBS and then 3.7% formaldehyde followed by tissue collection, as described. The heart and spinal column, with aorta and carotids attached, were removed, and fixed under gentle agitation for an additional 24 h at 4 °C, washed 3× with PBS and taken for further. For whole aorta *en face* prep, the isolated aortas were bisected along the lesser curvature and the aortic arch was bisected through the greater curvature as well. For aortic arch segment prep, the aortic arch was bisected through the greater curvature. Hearts (containing aortic roots) and carotids were allowed to sink in 30% sucrose in PBS overnight at 4°C, embedded in OCT compound (optimal cutting temperature compound, Sakura, 4583) and frozen on dry ice for sectioning. Tissue blocks were cut into 8-10 μm sections using a cryostat (Leica) and sections were stored at –80°C until use.

### Atherosclerosis and blood lipid analysis

To induce atherosclerosis, murine AAV8-PCSK9 adeno-associated virus (pAAV/D377Y-mPCSK9; 2×10^11^ PFU) produced by the Gene Therapy Program Vector Core at the University of Pennsylvania School of Medicine (Philadelphia, PA) was injected intraperitoneally. Mice were maintained on a high-fat diet (HFD; Clinton/Cybulsky high-fat rodent diet with regular casein and 1.25% added cholesterol; Research Diet, D12108c) for 16 weeks. Blood samples were collected from overnight starved mice, centrifuged at 13,000g at 4°C for 15 min and the supernatant (plasma) was extracted carefully and immediately stored at –80°C for total cholesterol and triglyceride (TAG) analysis. Plasma samples were stored at 4°C for <1 week for HDL-C analysis. Total plasma cholesterol was determined by cholesterol oxidase/colorimetric assay (Abcam; 65390). Plasma TAG levels were measured with a commercially available kit (Wako Pure Chemicals). Tissues were prepared as described above and analyzed as described below.

### Tissue analysis

For IF, OCT tissue sections were thawed and washed 3x with PBS to remove OCT. Cells were fixed with 3.7% formaldehyde for 15 min at ambient temperature, washed with and stored in PBS. Deidentified human specimens were deparaffinized in Histo-Clear (National Diagnostics, HS-200). Sections were progressively rehydrated before antigen retrieval for 30 min at 95°C in 1X Antigen Retrieval Buffer (Dako, 51699). Samples were incubated in perm-block buffer (5% donkey serum, 0.2% BSA, 0.3% Triton X-100 in PBS) for 1 h at room temperature, incubated with primary antibodies in perm-block overnight at 4°C, washed three times in perm-block and then incubated with alexafluor-conjugated secondary antibodies (ThermoFisher) at 1:1000 dilution in perm-block for 1 h at room temperature. Slides were washed 3x in perm-block and 3x in PBS before mounting in DAPI Fluoromount G (Southern Biotech; 0100–20). Images were acquired on a Leica SP8 confocal microscope with the Leica Application Suite software. Confocal stacks were flattened by maximum intensity z-projection in ImageJ. Post background subtraction, the mean fluorescence intensity (MFI) or nuclear intensity (with DAPI mask) was recorded. For aorta ORO staining, the whole aorta was opened longitudinally on a soft-bottomed silica dish, incubated with ORO solution (0.6% ORO in 60% isopropanol) with gentle rocking for 1 h at ambient temperature, washed in 60% isopropanol for 20 minutes, washed in dH2O 3x, mounted on slides with endothelium side up in OCT compound. Images were acquired with a digital microscopic camera (Leica DFC295). ORO staining on OCT tissue sections was done similarly. Quantitation of ORO positive area was done in ImageJ. Hematoxylin and eosin (H&E) staining on OCT tissue sections was done by Yale Research Histology Core using standard techniques. Plaque morphometric and vulnerability analysis was performed as described^47^. Plaque area was determined by ORO positive staining. For each plaque, the necrotic core (NC) area was defined as a clear area in the plaque that was H&E free, and the fibrous cap (FC) thickness was quantified by selecting the largest necrotic core and measuring the thinnest part of the cap.

### Cloning, Pcdhga9 ECD purification

Human Pcdhga9 mutants were generated by cloning PCR-amplified fragments into pBob-GFP vector (Addgene). For GFP-tagged constructs, fragments were cloned upstream and in-frame with the GFP ORF. For FLAG-tagged constructs, GFP was excised, and the fragments cloned with an added C-terminal FLAG tag using PCR. Mouse Pcdhga9 ECD was cloned in pCDNA3.1 vector (Invitrogen). Secreted Pcdhga9 ECD-FLAG-GST was purified from cell culture supernatant from transfected HEK293T cells by collecting medium at 48 h and 96 h after transfection. Supernatant was spun at 6000g, 4 °C for 15 min to remove debris, incubated with 200 µl washed Glutathione beads per 40 ml medium O/N at 4 °C with tumbling, washed 3x with 0.1% Triton X-100/PBS, and eluted by adding excess reduced Glutathione solution. The concentration of ECD was estimated by Coomassie staining compared to BSA standards.

### Generation and validation of mAbs and Pcdhga9 ECD-cell adhesion assay

1 mg purified Pcdhga9 ECD protein was used for monoclonal antibody (mAb) generation in rats (BiCell Scientific). mAbs were purified and concentrated from serum-free hybridoma cultures (BiCell Scientific). Low adhesion 96-well plates were coated with ECD (10 μg/ml) for 1h at ambient temperature, washed 3x with 0.1% Triton X-100/PBS and blocked with 1% heat denatured BSA in PBS for 1h at ambient temperature. These plates were used for testing IgG specificity and cell adhesion to ECD, as described below. For specificity, affinity, and amount of IgG, mAbs were added to the plates for 1h at ambient temperature with shaking, washed 3x with 0.1% Triton X-100 in PBS, incubated with secondary HRP ab (1:5k) for 1h at ambient temperature with shaking, washed 3x with 0.1% Triton X-100 in PBS and 3x with PBS, followed by addition of 100 μl TMB incubated for 15-30 min and absorbance measured at 605 nm, followed by addition of 100 ul 0.1N HCl and absorbance measured at 450 nm.

For cell adhesion assays, wells were washed 3x with complete EC medium followed by addition of *Klf2*:GFP MAECs in 100 μl total EC medium (with isotype control or test mAbs), incubated at 37 °C in the CO_2_ incubator for the indicated time and fixed by adding 33 μl of 16% PFA directly to the wells. Unadhered cells were removed by turning the plate upside down in a water bath and cells counted based on mCherry fluorescence.

### Partial Carotid Artery (PCA) Ligation model of accelerated experimental atherosclerosis

8-10 week-old *Apoe^-/-^* mice maintained on HFD for 1 week were anesthetized with ketamine and xylazine, and surgery was performed as described^48^. Briefly, 3 out of 4 branches of the left common carotid artery (left external carotid, internal carotid, and occipital artery) were ligated with sutures, with superior thyroid artery left intact. 2 μg mAb A9 or isotype control antibody was injected IP once every week for 2 weeks. Mice were euthanized with an overdose of isoflurane and perfused through the left ventricle with PBS and then 3.7% formaldehyde. Aortas with carotid arteries were isolated and imaged whole. Carotid arteries were embedded in OCT, sectioned at 10 μm, and IHC or immunofluorescence performed as described. LCA/RCA inner diameter ratio was calculated by measuring perimeter to avoid interference from changes in vessel morphology during mounting and handling. For testing mAb retention in vivo (half-life), 1 µg of mAbs (100 μl of 0.25 mg/ml in saline for a ∼20 g mouse) were injected IP in C57Bl/6 mice, 100 μl blood was collected via the retro-orbital route at the indicated times, centrifuged at 13,000g at 4°C for 15 min and the plasma removed and immediately stored at –80°C. mAb concentration was determined using a sandwich ELISA with immobilized anti-rat IgG (100 ng) as trap and secondary anti-rat IgG HRP antibody for detection, described above.

### Immunoblotting and immunoprecipitation (IP)

Cells were harvested and lysed in RIPA buffer (Roche) containing 1x Halt Protease Inhibitor Cocktail (ThermoFisher, 78429) and 1x PhosStop (Roche, 4906837001) for 30 min on ice, clarified at 13,000 g 4°C for 15 minutes, the supernatant transferred to new 1.5 ml tubes, 4x loading buffer (250mM Tris-HCl pH 6.8, 8% SDS, 40% glycerol, 20% β-mercaptoethanol, 0.008% bromophenol blue) added and the samples heated to 95°C for 5 min. Cell lysates were resolved by a 4-15% SDS Polyacrylamide gel electrophoresis, transferred to 0.2 μm nitrocellulose membranes which were blocked with 5% non-fat skim milk for 1h at ambient temperature and incubated with desired antibodies diluted in 5% BSA using a standard immunoblotting procedure and detection by ECL (Millipore). Images were quantified in ImageJ by densitometry and normalized to GAPDH or Tubulin loading controls. For immunoprecipitation (IP), GFP-trap (GTA-20; Chromotech) or FLAG M2 (Millipore Sigma, A2220) agarose was used. Lysates were harvested in 25 mM Tris, pH 7.4, 150 mM NaCl, 1% TritonX-100, 1x Halt Protease Inhibitor Cocktail (ThermoFisher, 78429) and 1x PhosStop (Roche, 4906837001), clarified at 13,000 g 4°C for 15 minutes, incubated with antibody-bound beads at 4°C for 2h to overnight. Beads were washed 3 times with 4°C lysis buffer and eluted with 2x protein sample buffer (for GFP-trap) or 3x FLAG peptide competition (for FLAG M2) and subjected to SDS-PAGE or Mass Spectrometry.

### RNA isolation, sequencing, and quantitative real-time PCR (qPCR)

Total RNA was extracted from cells with RNeasy Plus Mini Kit (Qiagen, 74136) according to the manufacturer’s instructions. RNA was quantified by NanoDrop, and RNA integrity was measured with an Agilent Bioanalyzer. Samples were subjected to RNA sequencing using Illumina NovaSeq 6000 (HiSeq paired-end, 100 bp). The base calling data from the sequencer were transferred into FASTQ files using bcl2fastq2 conversion software (version 2.20, Illumina). PartekFlow (a start-to-finish software analysis solution for next generation sequencing data applications) was used to determine differentially expressed genes (DEGs). For qPCR analysis, reverse transcription was performed with iScript™ Reverse Transcription Supermix for RT-qPCR (Bio-Rad). qPCR was performed real-time PCR with SsoAdvanced Universal SYBR Green Supermix (Biorad). The expression of target genes was normalized to GAPDH. RT-PCR primers are listed in Table S4.

### Quantification and Statistical Analysis

ImageJ software (version 1.51; National Institutes of Health, Bethesda, MD) was used for morphometric analysis. Graph preparation and statistical analysis was performed using GraphPad Prism 10.0 software (GraphPad software Inc.). Data were considered normally distributed, statistical significance was performed using Student t test for two groups comparison or one-way ANOVA with Tukey’s post hoc analysis for multiple groups comparison, as described in the figure legends. Data are presented as mean ± SEM. A p value less than 0.05 was considered significant (*p < 0.05, **p < 0.01, ***p < 0.001, ****p < 0.0001).

## Data and Resource Availability

Data and resources generated and/or analyzed during this study are either freely available or available from the corresponding author upon reasonable request.

## Author contributions

DJ performed most of the experiments, analyzed data and prepared figures. BGC performed the CRISPR screen and analysis. DJ and RC performed and analyzed PCA Ligation experiments. DJ, HD, and EM performed antibody injection experiments in mice. HD and PFT performed plasma lipid analysis experiments. CFH supervised lipid analysis experiments. MAS conceived of, supervised, and acquired funding for the project. DJ and MAS wrote the manuscript.

## Acknowledgments

The authors thank Julie Lefebvre (University of Toronto, Canada) for *Pcdhg^fcon3^* mice, Jiasheng Zhang (Yale University Animal Core) for PCA Ligation surgery, Dejian Zhao (Yale Center for Genome Analysis) for analysis of RNAseq data, Schwartz lab members for extensive discussions, Yale Center for Genome Analysis for RNAseq, Yale Keck Oligo Synthesis Resource and DNA Sequencing Core, and Harvard Center for Mass Spectrometry. This work was supported by National Institutes of Health grant R01 HL75092, a Leducq Trans-Atlantic Network Grant, and a Health Research consortium grant from Fundación Obra Social La Caixa (AtheroConvergence, HR20-00075) to MAS and American Heart Association grant #23POST1026109 to DJ.

## Online supplemental material

Online supplemental material includes 8 figures (Figures S1-S8), 4 tables (Tables S1-S4), and link to access RNAseq files.

