## Supplementary Information for "Gamma protocadherins in vascular endothelial cells inhibit Klf2/4 to promote atherosclerosis"

#### Supplementary figure legends

**Figure S1. Validation of Klf2 reporter.** (a) Immunoblot validation of LSS-mediated induction of *Klf2*:GFP in reporter MAECs; reporter is also sensitive to statin treatment, another potent inducer of Klf2 used as a positive control. (b) Rank-ordered candidate suppressors from the CRISPR screen to identify Klf2 modifiers (z<4 gray, z>4 blue), with validated cell-surface exposed candidates which are amenable to function neutralization marked in red, and known positive controls, Ccm2 and Pdcd10, marked in black. (c) qRT-PCR validation of siRNA mediated knockdown of Pcdhga9 in MAECs (Pcdhga9 si) (N=3). (d) Immunoblot validation of human Pcdhga9-FLAG overexpression (Hs Pcdhga9 OE) in MAECs. (e) Control si and Pcdhga9 si *Klf2* reporter MAECs were exposed to static (St), LSS, or oscillatory shear stress (OSS) for 16 h and immunoblotted for *Klf2*:GFP. (f) qRT-PCR for endogenous *Klf2* levels in Control si and Pcdhga9 si in Human Umbilical Vein Endothelial Cells (HUVECs) exposed to St or OSS for 16 h (N=3). Statistical analysis used one-way ANOVA.

**Figure S2. Protocadherin gamma (Pcdhg) gene cluster promotes inflammatory signaling.** (a) Validation of siRNA mediated knockdown of Pcdhg gene cluster in HUVECs using two different siRNAs targeting the common region in the 3' end, by qRT-PCR for Pcdhgc3, the highest expressed Pcdhg member in HUVECs (N=3). (b) qRT-PCR for the OSS induced pro-inflammatory mark *E-selectin (Sele)* in control and Pcdhg depleted HUVECs exposed to St, LSS and OSS for 16 h (N=3). (c) Immunoblot for VCAM1 in Control si or Pcdhg si HUVECs, after treatment with indicated doses of TNFα for 16 h (N=3). Graph: quantitation of VCAM1 normalized to Tubulin loading control. Statistical analysis used one-way ANOVA.

**Figure S3. Validation and blood lipid analysis of Pcdhg ECKO mouse.** (a) Generation of Pcdhg endothelial knockout (ECKO) by crossing $Pcdhg^{fcon3}$ with *Cdh5Cre*, and confirmation by genotyping PCR. wt: wild type for *Pcdhg* allele; flox: *Pcdhg floxed*; Cre: *Cdh5Cre*. (b, c) Analysis of progeny genotype showing no significant deviation from Mendelian ratio as tested by Chi-squared analysis. (d-f) Plasma triglycerides, cholesterol, HDL-C and body weights of male and female Control or Pcdhg ECKO mice injected with pCSK9-Adeno Associated Virus 8 (AAV8) and maintained on High Fat Diet (HFD) for 16 weeks, starved overnight before analysis. Statistical analysis was carried out using Student's t-test.

**Figure S4. Conserved ICD region is necessary and sufficient for Pcdhg function.** (a) Pcdhg mutants were expressed in HUVECs which were treated with St, LSS or OSS and immunoblotted for GFP, Klf4 and GAPDH(N=3). (b) GFP was imaged in HUVECs expressing the above mutants. (c) Multiple sequence alignment to test domain homology domains, using Pcdhga9 as an example, showing near-complete conservation in CCDs (highlighted in yellow). Percent conservation shown in the box. Scale bar: (c) 20 µm.

**Figure S5. Notch-dependent Klf2/4 regulation and Pcdhg-Notch interaction.** (a) RBPJ-Notch DNA-binding consensus motif. (b) RBPJ-Notch binding motifs in mouse and human *Klf2* and *Klf4* promoters. (c) NICD peptides detected in proteomic analysis of the IPs of full but not the ΔCCD Pcdhg mutant.

**Figure S6. Pcdhga9 blocking antibody generation and validation.** (a) Pcdhga9 ECD (ECD-FLAG-TEV-GST) protein run on a 10% Polyacrylamide SDS gel and visualized using Imperial Protein stain (Thermo Scientific). (b) ELISA for the 24 select high affinity monoclonal antibody (mAb) from clones labeled as A1-12 and B1-12 using ECD alone (GST cleaved off using TEV protease). (c) Adhesion of MAECs to ECD in the presence of Isotype control or mAbs A9, B1 and B4 (N=4). Graph: quantitation of percent total cells adhered to ECD. (d) Immunoblot with mAbs A9, B1, B4 shows detection of both ECD-FLAG-TEV-GST and ECD-FLAG. (e, f) *Klf2*:GFP reporter MAECs tested for *Klf2*:GFP reporter expression after 16h LSS (N=12) (e) or immunostained for VCAM1 when exposed to OSS for 16 h OSS (N=7) (f), in the presence of Isotype control or mAbs A9, B1 and B4 as indicated. Graphs: quantitation of *Klf2*:GFP and VCAM1 levels normalized to mCherry internal control. (g) Immunofluorescence with mAbs A9, B1, B4 showing highest signal from A9. Statistical analysis was carried out using one-way ANOVA. Scale bar: (c) 100 µm, (g) 20 µm. *p < 0.05, ns (not significant) > 0.05.

**Figure S7. Pcdhga9 mAb A9 validation.** (a) Immunoblot of Pcdhg knockdown cells with mAb A9. Purified ECD used as positive control. (b) Antibody half-life in vivo. A single dose of 2 µg of Isotype control or mAb A9 antibody was administered IP, and plasma levels of antibody were measured using an ELISA as described in Methods. mAb levels at 3h post injection were considered 100%. (c) LCA and RCA sections were stained for smooth muscle specific Acta2 (SMA) for marking plaque neointima (N=6). Graph: quantitation of the LCA to RCA inner diameter. (d-e) Plasma triglycerides and cholesterol in Isotype control or mAb A9 injected male and female Apoe$^{-/-}$ mice on HFD from figure 6h, i. Statistical analysis was carried out using Student's t-test. Scale bar: (c) 100 µm. *p < 0.05, ns (not significant) > 0.05.

**Figure S8. Pcdhg level correlates with atherosclerosis.** (a) Mouse carotids from the Partial Carotid Artery (PCA) Ligation model of accelerated atherosclerosis stained for Pcdhg and counter stained with DAPI to mark nuclei (N=3). RCA: Right Carotid Artery (control), LCA: Left Carotid Artery (atherosclerotic plaque. Graph: quantitation of Pcdhg staining intensity. (b) Commercial Pcdhg antibody and mAb A9 staining of retinal vasculature from Control and Pcdhg ECKO mouse. mAb A9 is specific to mouse Pcdhga9 and Pcdhg antibody targets the conserved region in 22 Pcdhg genes, also conserved between mouse and human. Statistical analysis was carried out using Student's t-test. Scale bar: (a) 100 µm, (b) 10 µm. *p < 0.05, **p < 0.01, ***p < 0.001, ns (not significant) > 0.05.

### Figure S1. Validation of Klf2 reporter

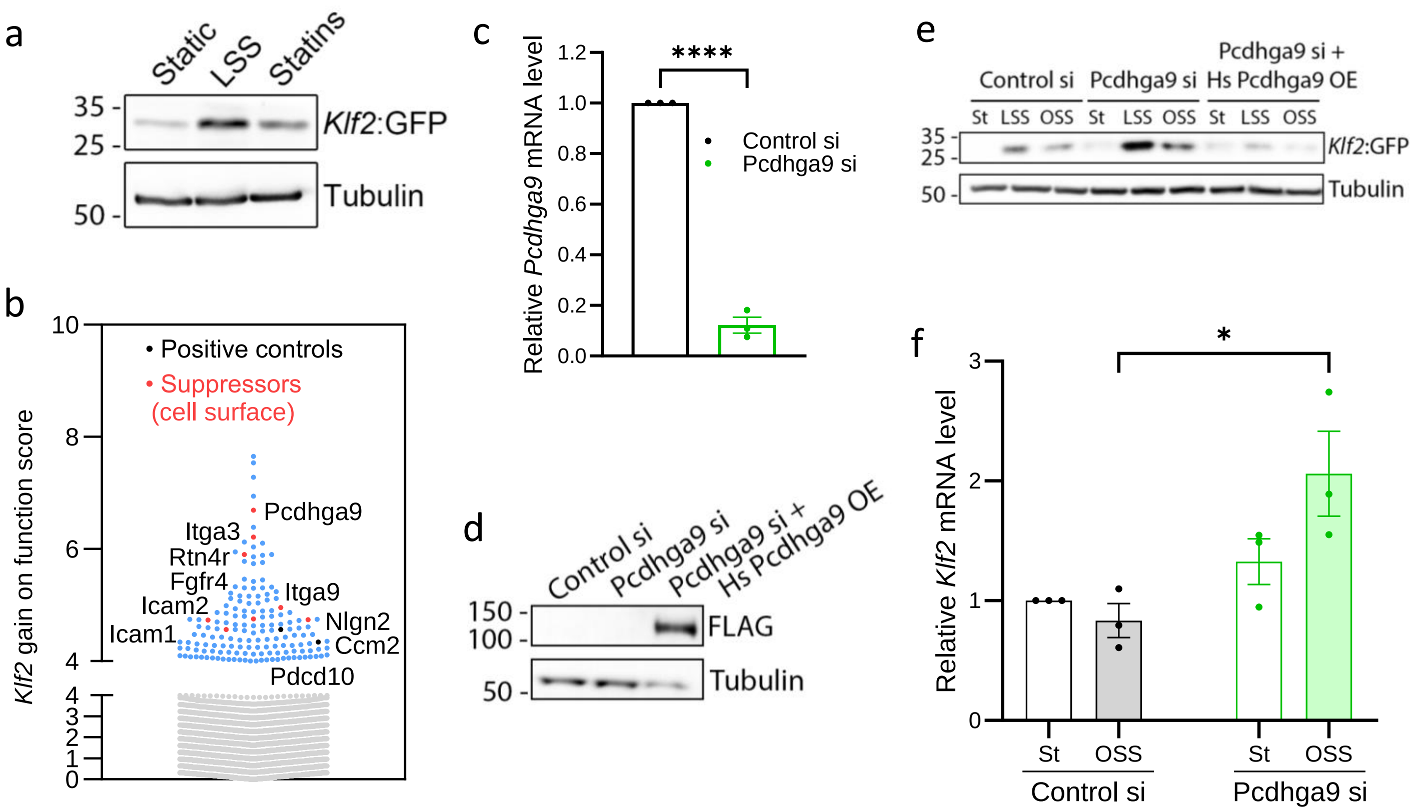

### Figure S2. Protocadherin gamma (Pcdhg) gene cluster promotes inflammatory signaling

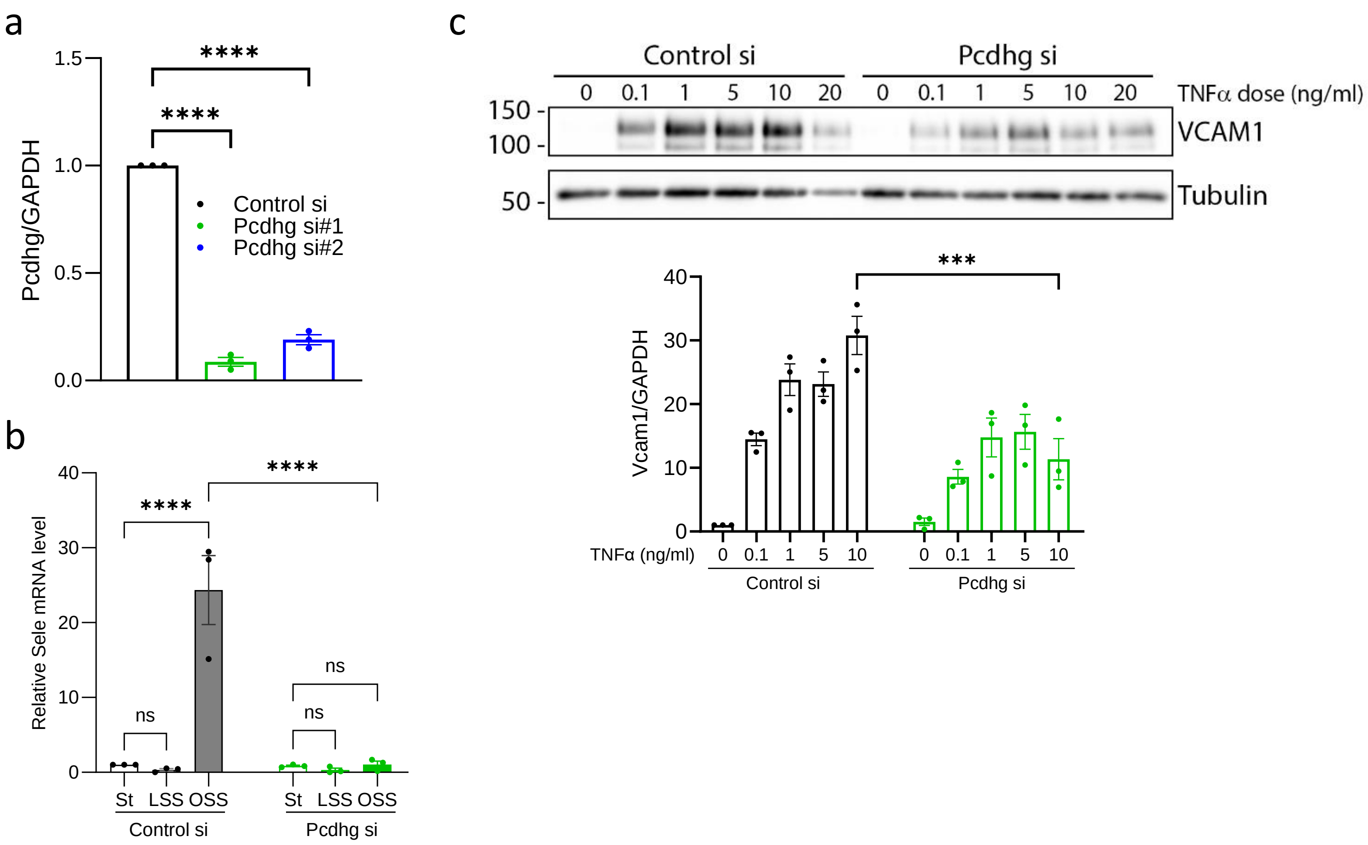

### Figure S3. Validation and blood lipid analysis of Pcdhg ECKO mouse

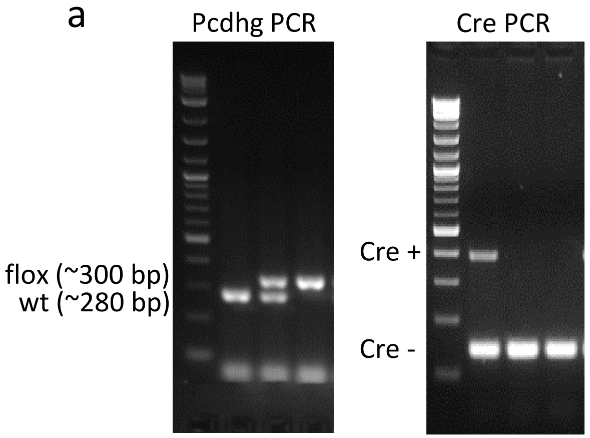


b

| Genotype/ Breeding # | Mut/Mut; Cre+/Cre- | WT/Mut; Cre+/Cre- | WT/WT; Cre+/Cre- | Mut/Mut; Cre-/Cre- | WT/Mut; Cre-/Cre- | WT/WT; Cre-/Cre- | Total |
|---|---|---|---|---|---|---|---|
| Breeding #1 | 2m 1f | 1m 1f | 0m 0f | 2m 1f | 3m 0f | 0m 0f | 8m 3f |
| Breeding #2 | 1m 0f | 1m 0f | 0m 1f | 0m 0f | 6m 1f | 0m 1f | 8m 3f |
| Breeding #3 | 1m 1f | 0m 2f | 0m 0f | 1m 0f | 0m 0f | 0m 1f | 2m 4f |
| Breeding #4 | 1m 1f | 1m 1f | 0m 0f | 1m 0f | 0m 2f | 1m 0f | 4m 4f |
| Breeding #5 | 0m 0f | 1m 2f | 0m 0f | 0m 1f | 0m 2f | 1m 0f | 2m 5f |
| Breeding #6 | 1m 1f | 1m 0f | 0m 0f | 0m 2f | 0m 1f | 0m 0f | 2m 4f |
| Breeding #7 | 0m 0f | 0m 1f | 1m 0f | 0m 1f | 1m 2f | 0m 1f | 2m 5f |
| Total | 6m 4f | 5m 7f | 1m 1f | 4m 5f | 10m 8f | 2m 3f | 28m 28f |

c

| df/χ2 | 0.995 | 0.99 | 0.975 | 0.95 | 0.90 |
|---|---|---|---|---|---|
| 1 | --- | --- | 0.001 | 0.004 | 0.016 |
| 2 | 0.010 | 0.020 | 0.051 | 0.103 | 0.211 |
| 3 | 0.072 | 0.115 | 0.216 | 0.352 | 0.584 |
| 4 | 0.207 | 0.297 | 0.484 | 0.711 | 1.064 |
| 5 | 0.412 | 0.554 | 0.831 | 1.145 | 1.610 |

df = 5; $\chi 2$ observed = 9.657534

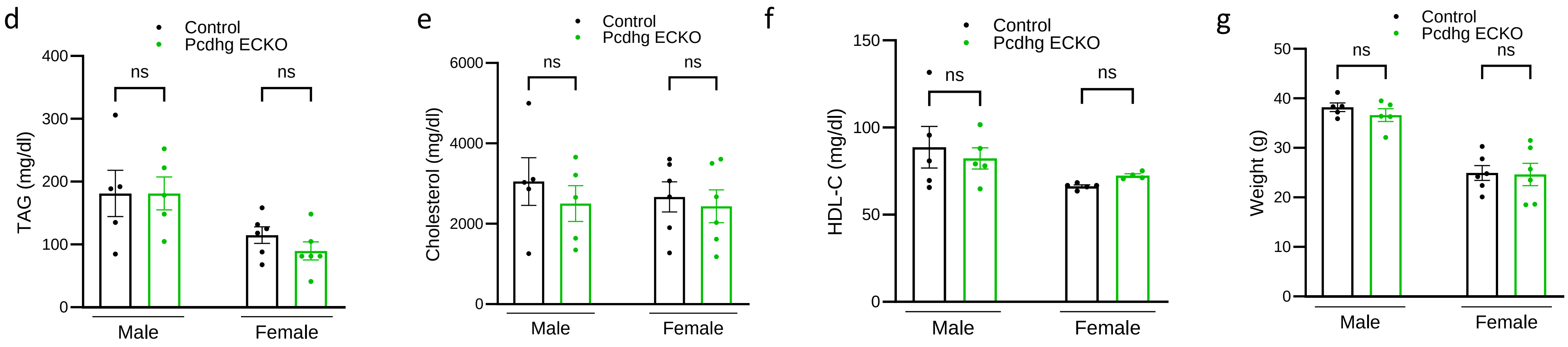

### Figure S4. Conserved ICD region is necessary and sufficient for Pcdhg function

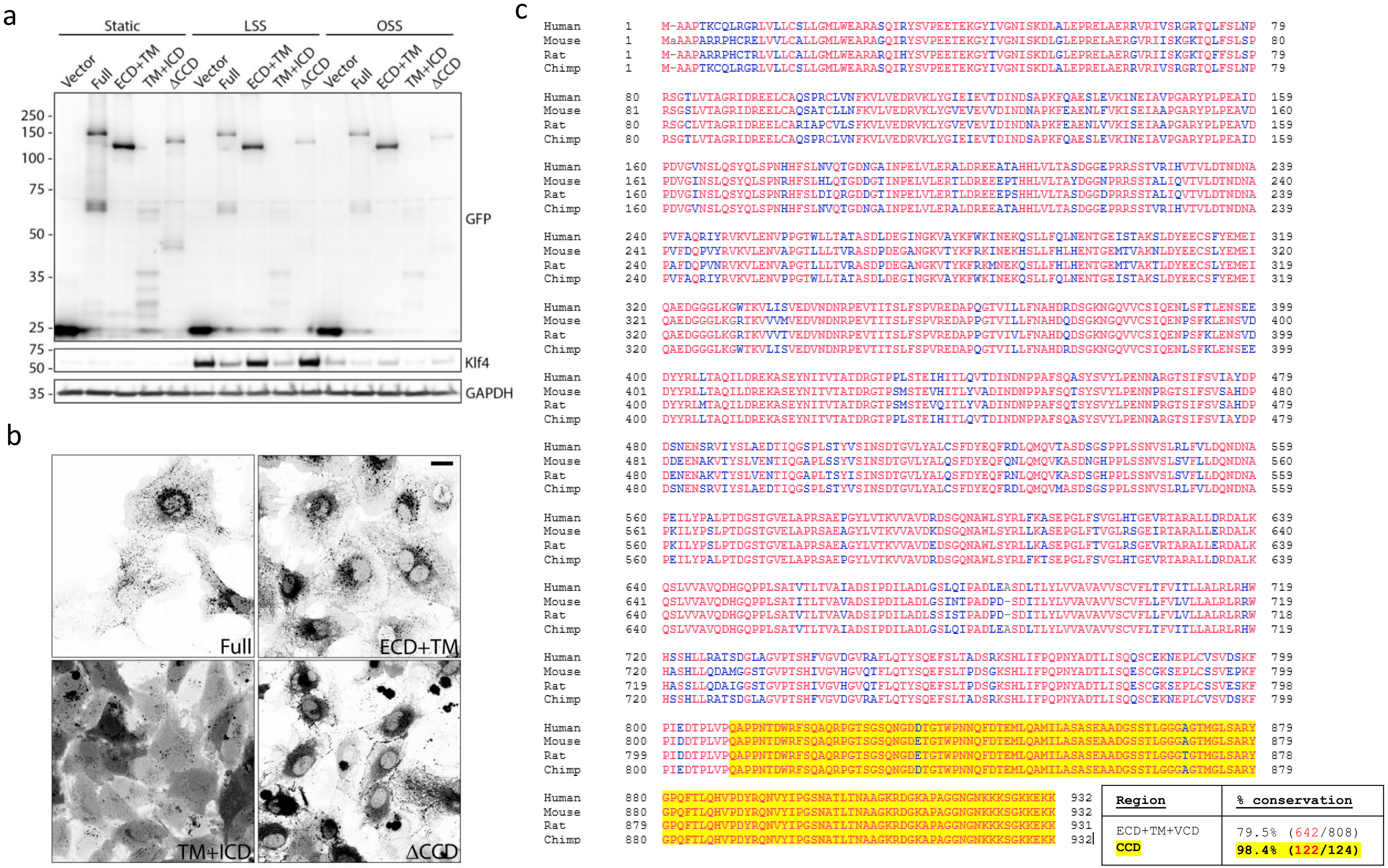

### Figure S5. Notch-dependent Klf2/4 regulation and Pcdhg-Notch interaction

a

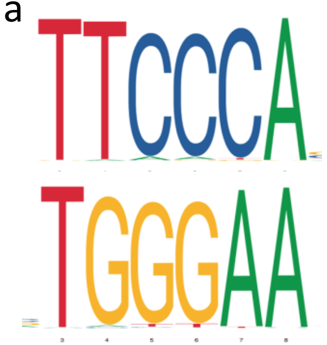


b

| Promoter regions | RBPJ motif location |
|---|---|
| Mm Klf2 promoter (1.7 kb) | -939 |
| Hs Klf2 promoter (2.2 kb) | -2204; -1688; -469 |
| Mm Klf4 promoter (1.6 kb) | -1065; -957; -736 |
| Hs Klf4 promoter (2.5 kb) | -2407; -1632; -1438 |

c

| | Vector IP | Full IP | ΔCCD IP |
|---|---|---|---|
| # of unique peptides | 0 | 6 | 0 |
| Peptide sequence | N/A | 1. RREPLGEDSVGLKPLKN<br>2. KFRFEEPVVLPDLDDQTDHRQ<br>3. RTGETALHLAARY<br>4. RMHDGTTPLILAARL<br>5. KVLLDHFANRD<br>6. RLLDEYNLVRS | N/A |

### Figure S6. Pcdhga9 blocking antibody generation and validation

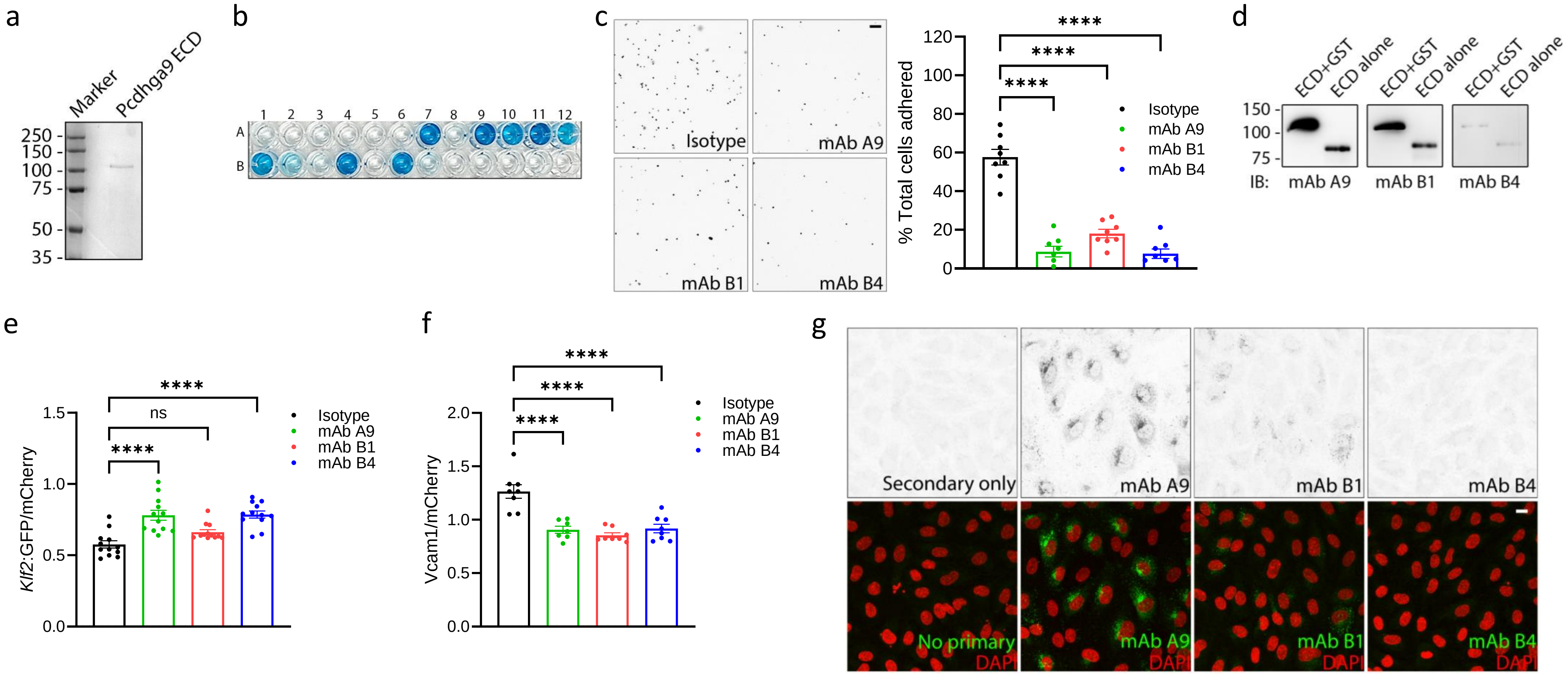

### Figure S7. Pcdhga9 mAb A9 validation

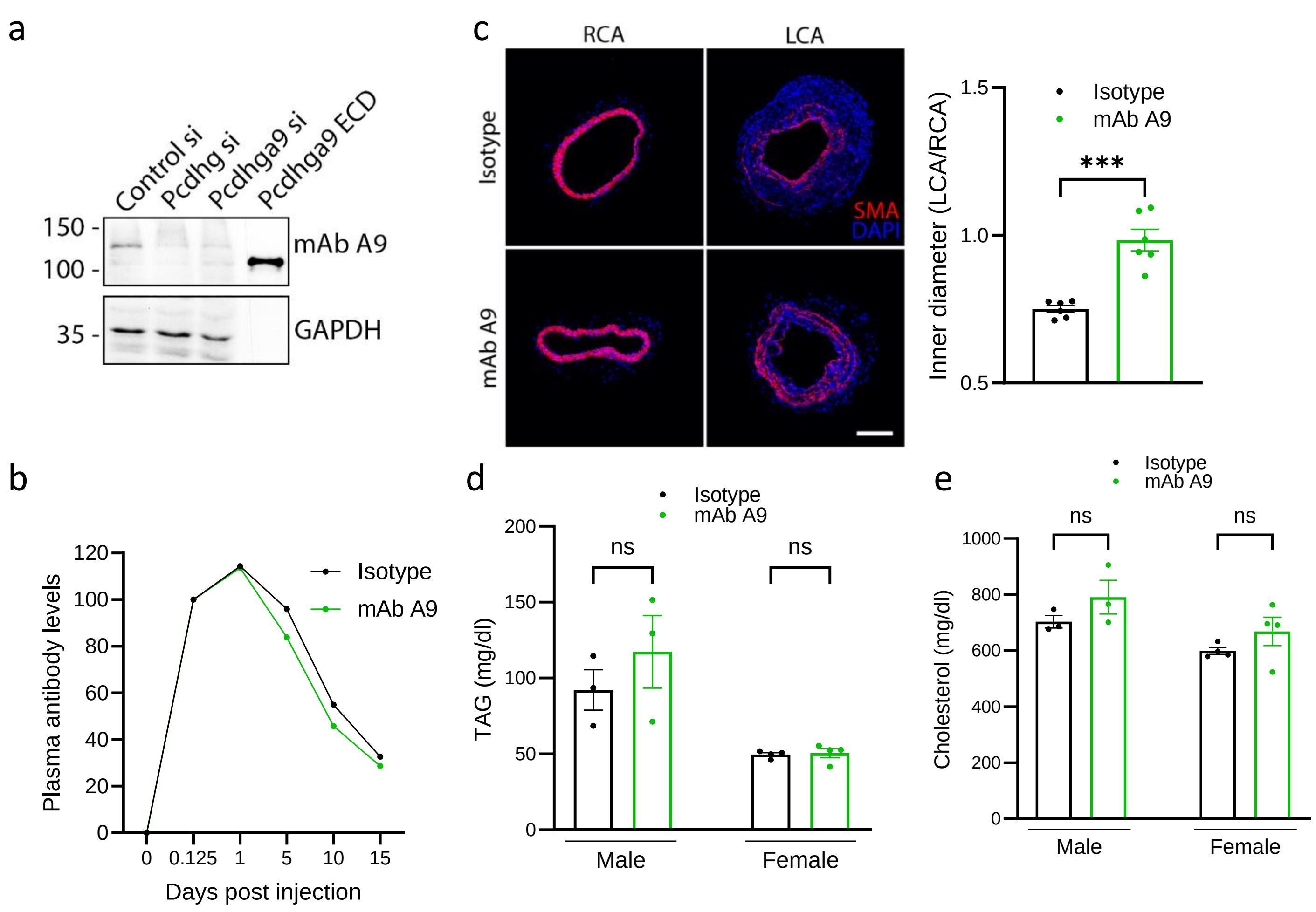

### Figure S8. Pcdhg level correlates with atherosclerosis

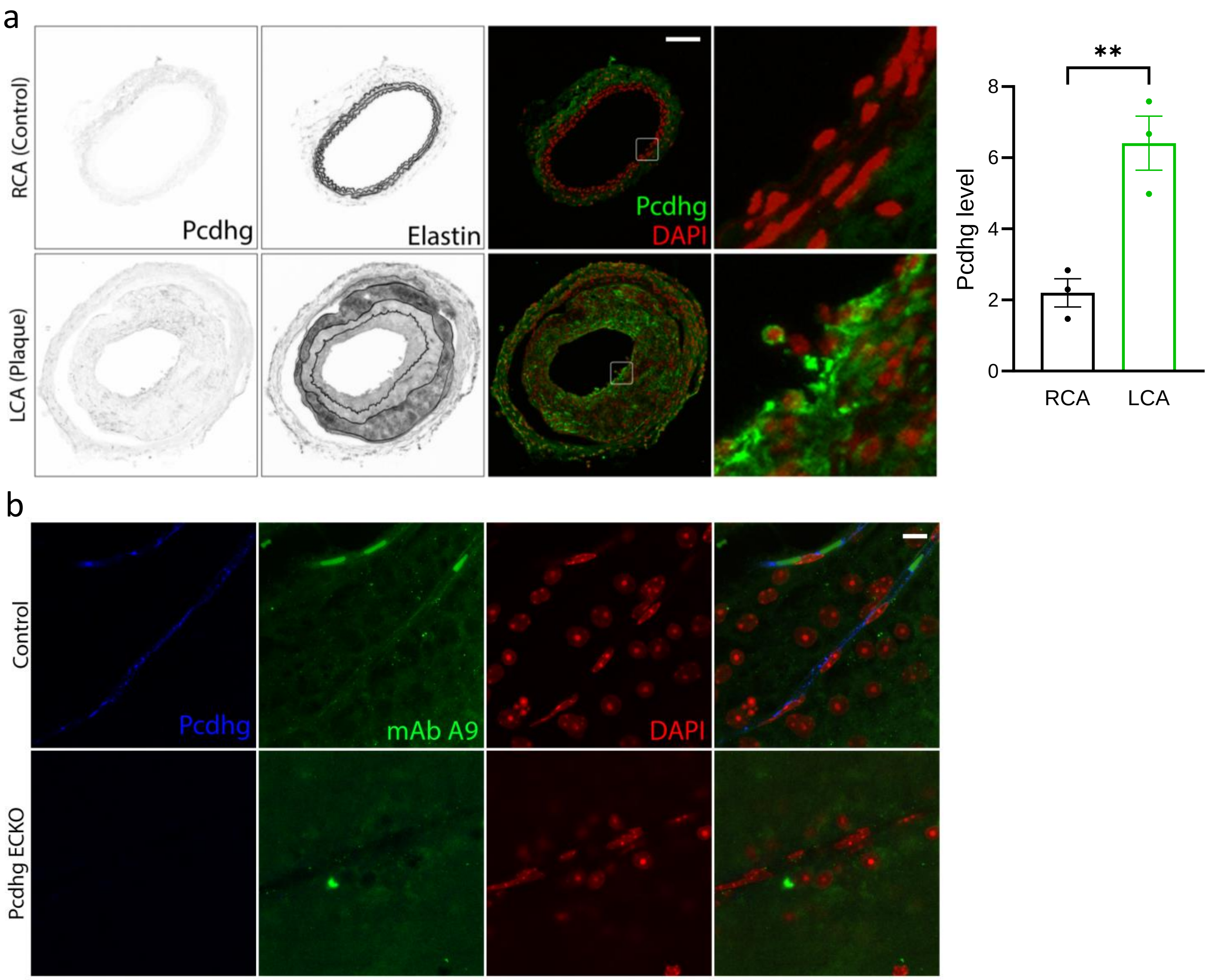

**Table S1a. Suppressors (cumulative z)**

| Gene | Cumulative z (top 3) |
|---|---|
| *Slc16a3* | *7.655257984* |
| *Cryl1* | *7.540379348* |
| *Pafah1b3* | *7.279220923* |
| *Polrmt* | *6.944511058* |
| ***Pcdhga9*** | ***6.689505294*** |
| *Vgll3* | *6.390476171* |
| *Itga3* | *6.21238783* |
| *Psmd3* | *6.127920702* |
| *Pif1* | *6.101544168* |
| *Abhd4* | *6.040343525* |
| *Ppp1r12c* | *5.950945245* |
| *Rtn4r* | *5.904293171* |
| *Ermap* | *5.899203615* |
| *Dcp2* | *5.889770257* |
| *Kremen1* | *5.862772626* |
| *Nadk* | *5.778027221* |
| *Ssbp1* | *5.759401317* |
| *B3gntl1* | *5.738200189* |
| *Hey1* | *5.465748602* |
| *Dbf4* | *5.447221667* |
| *Kdm3b* | *5.420450609* |
| *Crtc3* | *5.331007859* |
| *Siae* | *5.319192637* |
| *Avpi1* | *5.306922002* |
| *Litaf* | *5.30423513* |
| *Slc45a3* | *5.298138066* |
| *Scyl3* | *5.237637428* |
| *Metap1* | *5.21404302* |
| *Sh2d3c* | *5.190044825* |
| *Rpl35a* | *5.168159221* |
| *Osbpl1a* | *5.159717441* |
| *Gadd45g* | *5.155559372* |
| *Gpr137* | *5.117870504* |
| *Rogdi* | *5.115466004* |
| *Golph3l* | *5.091454623* |
| *L3mbtl2* | *5.071260731* |
| *Ltb* | *5.034670626* |
| *Stard10* | *5.029088032* |
| *Ginm1* | *4.98814457* |
| *Itga9* | *4.95604584* |
| *Mrps25* | *4.935085856* |
| *Dhx36* | *4.902533169* |
| *Ccnjl* | *4.897027381* |
| *Stt3b* | *4.896791823* |

| | |
|---|---|
| *Ap3m1* | *4.879159375* |
| *Pgm2* | *4.868837703* |
| *Bckdha* | *4.866650298* |
| *Eml1* | *4.857109184* |
| *Cidea* | *4.829771396* |
| *Tfap2a* | *4.80482265* |
| *Elmsan1* | *4.800698949* |
| *Zmym1* | *4.799581619* |
| *Atxn7l2* | *4.789635721* |
| *Nlgn2* | *4.751228498* |
| *Mcm4* | *4.746085498* |
| *Tmem104* | *4.745305578* |
| *Icam1* | *4.742356641* |
| *Slc35b4* | *4.736911925* |
| *Icam2* | *4.735612438* |
| *Ccdc92* | *4.729278734* |
| *Jazf1* | *4.701482173* |
| *Fbxo41* | *4.694825911* |
| *Rexo4* | *4.688675238* |
| *Bbs2* | *4.659084888* |
| *Hps4* | *4.65111877* |
| *Larp4b* | *4.647438713* |
| *Fam53b* | *4.646733355* |
| *Chac1* | *4.640751746* |
| *Pros1* | *4.630282139* |
| *Hsf2* | *4.616303712* |
| *Gsg2* | *4.6113801* |
| *Rhbdd1* | *4.601306075* |
| *Fkbp1a* | *4.593417482* |
| *Atg16l2* | *4.579352385* |
| *Sall2* | *4.572695452* |
| *Fgfr4* | *4.565617434* |
| *Pdcd10* | *4.564219138* |
| *Ankrd6* | *4.563438569* |
| *Tshz1* | *4.549193405* |
| *Sdccag3* | *4.54251268* |
| *Zbtb7b* | *4.53559488* |
| *Abhd11* | *4.502922792* |
| *Lsm11* | *4.474008096* |
| *B2m* | *4.47316319* |
| *Tsc1* | *4.463584287* |
| *Sepn1* | *4.462924022* |
| *Tstd3* | *4.462528141* |
| *Hhipl1* | *4.430386877* |
| *Prss35* | *4.429402898* |
| *Iqce* | *4.429146628* |
| *Apba3* | *4.40449467* |

| | |
|---|---|
| *Pan2* | *4.39949643* |
| *Hpcal1* | *4.389165078* |
| *Creg1* | *4.377320314* |
| *Ift122* | *4.377271773* |
| *Cdk5* | *4.366727504* |
| *Mospd3* | *4.356239949* |
| *Ccm2* | *4.340795791* |
| *Pygl* | *4.338790167* |
| *Mta1* | *4.3352845* |
| *Mipol1* | *4.31618398* |
| *Stk19* | *4.305754492* |
| *Pdcd5* | *4.296808879* |
| *Pik3r3* | *4.291346512* |
| *Bag5* | *4.283521035* |
| *Ptafr* | *4.281690295* |
| *Fbxo5* | *4.280230666* |
| *Fut10* | *4.272845338* |
| *F12* | *4.268111417* |
| *Kdm1b* | *4.26376406* |
| *Haus5* | *4.255414449* |
| *Tmem159* | *4.250094656* |
| *Cdca4* | *4.249033107* |
| *Agpat1* | *4.248183719* |
| *Selp* | *4.236192666* |
| *Arhgap19* | *4.235246115* |
| *Tshz3* | *4.226312498* |
| *Hnrnpf* | *4.210216667* |
| *Myl12a* | *4.201389827* |
| *Gchfr* | *4.188604759* |
| *Setdb1* | *4.182349248* |
| *Parn* | *4.182289899* |
| *Asb1* | *4.177633162* |
| *Shank1* | *4.177430028* |
| *Neurl4* | *4.159519608* |
| *Traf4* | *4.159456552* |
| *Myo5c* | *4.158875034* |
| *Itgb1bp1* | *4.146205708* |
| *Pfn1* | *4.14441859* |
| *Axin1* | *4.143596494* |
| *Lysmd1* | *4.135793279* |
| *Lrch1* | *4.133427274* |
| *Sema4b* | *4.1278137* |
| *Wwox* | *4.112018923* |
| *Gbx2* | *4.096009775* |
| *Fam124a* | *4.0893966* |
| *Ank2* | *4.08786855* |
| *Bicc1* | *4.083103592* |

| | |
|---|---|
| *Sox18* | *4.080849324* |
| *Dxo* | *4.075187467* |
| *Prcc* | *4.074998335* |
| *Folh1* | *4.071042599* |
| *Acr* | *4.063004297* |
| *Selo* | *4.062944078* |
| *Eif3g* | *4.062509291* |
| *Rab22a* | *4.055854204* |
| *Bcorl1* | *4.054921512* |
| *Mtg2* | *4.049239012* |
| *Ctif* | *4.048805549* |
| *Oaz3* | *4.036446561* |
| *Pls1* | *4.031352908* |
| *Med23* | *4.030781931* |
| *Utp18* | *4.030751193* |
| *Foxn2* | *4.027201095* |
| *Cxxc4* | *4.025822742* |
| *Zcchc9* | *4.023177396* |
| *Parp6* | *4.01938878* |
| *Ccp110* | *4.014257115* |
| *Cers6* | *4.00411303* |
| *F2rl1* | *4.000510052* |
| Tango6 | 3.998040349 |
| Glmn | 3.992825449 |
| Bdh1 | 3.989702029 |
| Elavl1 | 3.987529801 |
| Rps10 | 3.986678211 |
| Phf20 | 3.986179197 |
| Sh3bgr | 3.981393375 |
| Foxs1 | 3.977374001 |
| Grhl1 | 3.973828418 |
| Frrs1 | 3.973321475 |
| Sec11a | 3.969887531 |
| Zcchc14 | 3.969392067 |
| Mef2b | 3.968006844 |
| Arl10 | 3.957557271 |
| Tbc1d10b | 3.953250213 |
| Ifnar2 | 3.948753136 |
| Hectd3 | 3.940672672 |
| Glul | 3.933944466 |
| Adora2a | 3.930033942 |
| Ndrg1 | 3.923123792 |
| Gp1ba | 3.909901227 |
| Aldh1b1 | 3.907425396 |
| Loxl3 | 3.906389046 |
| Tmem9b | 3.900988724 |
| Zbtb44 | 3.895042432 |

| | |
|---|---|
| Nsmf | 3.890540446 |
| Haus2 | 3.889919105 |
| Calml4 | 3.889813351 |
| Stac | 3.885654553 |
| Zfp91 | 3.878419897 |
| Ascc1 | 3.876912961 |
| Lims2 | 3.874891426 |
| Crat | 3.863469326 |
| Sdf4 | 3.860480027 |
| Gpt2 | 3.858110558 |
| Kirrel | 3.85531223 |
| Ubl3 | 3.853937611 |
| Gpatch3 | 3.845288951 |
| Rhot2 | 3.84360257 |
| Dhx16 | 3.831053329 |
| Ubn2 | 3.828502829 |
| Plekhg1 | 3.822018175 |
| Athl1 | 3.818717491 |
| Psmc4 | 3.813800933 |
| Armc10 | 3.806085094 |
| Stard3 | 3.804071916 |
| Ap2a1 | 3.802891521 |
| Zc3h18 | 3.802033422 |
| Laptm4a | 3.797607837 |
| Jarid2 | 3.795206935 |
| Rinl | 3.793013982 |
| Rab6b | 3.788661748 |
| Il10rb | 3.78538361 |
| Chrd | 3.779065819 |
| Maml1 | 3.764717847 |
| Zbtb39 | 3.76137551 |
| Kif3c | 3.761373136 |
| Sertad2 | 3.761214304 |
| Ap3b1 | 3.759273597 |
| Usp33 | 3.758589772 |
| Atp6ap1l | 3.752168244 |
| Prdx6 | 3.746931334 |
| Vwa1 | 3.744613881 |
| Coprs | 3.744351174 |
| Trim11 | 3.739694702 |
| Efemp2 | 3.737965888 |
| Tctn1 | 3.737532424 |
| Emc1 | 3.736651276 |
| Atp2b4 | 3.736141048 |
| Pkdcc | 3.735686935 |
| Il15ra | 3.735463596 |
| Sphk2 | 3.729852012 |

| | |
|---|---|
| Armcx2 | 3.72753584 |
| Phf19 | 3.724029901 |
| Rce1 | 3.722275155 |
| Mrs2 | 3.722138807 |
| Pcif1 | 3.721769405 |
| Rpusd4 | 3.721010529 |
| Wdr60 | 3.716935526 |
| Tnfrsf12a | 3.716685125 |
| Stat3 | 3.714117653 |
| Paox | 3.713635639 |
| Vopp1 | 3.71333222 |
| Bcat2 | 3.70835197 |
| Fopnl | 3.704140665 |
| Doc2b | 3.702526862 |
| Fbrsl1 | 3.700778901 |
| Crispld2 | 3.698885447 |
| Slc16a4 | 3.694813972 |
| Timm13 | 3.693165653 |
| Pced1b | 3.692474194 |
| Tmem43 | 3.691996832 |
| Apbb3 | 3.69126688 |
| Chst7 | 3.690614572 |
| Eef1a1 | 3.688126607 |
| Eef1b2 | 3.685026593 |
| Nudt16l1 | 3.676873243 |
| Csnk1e | 3.676725497 |
| Ppt1 | 3.675843528 |
| Eng | 3.674595657 |
| Apex1 | 3.665840858 |
| Tnfsf4 | 3.661729579 |
| Psmb8 | 3.659298919 |
| Pak1ip1 | 3.657312721 |
| Slc7a8 | 3.656619659 |
| Commd8 | 3.655122334 |
| Rnf167 | 3.655065485 |
| Pias1 | 3.654184668 |
| Gga1 | 3.650339911 |
| Slc25a32 | 3.649948141 |
| Gng7 | 3.648379445 |
| Rnaseh2a | 3.646643478 |
| Dnajb14 | 3.646423077 |
| Sulf2 | 3.645556111 |
| Foxc2 | 3.643078349 |
| Atmin | 3.642635972 |
| Hif1an | 3.640962892 |
| Usp50 | 3.639192976 |
| Igflr1 | 3.634482354 |

| | |
|---|---|
| Fam214b | 3.630864187 |
| Rarb | 3.630260239 |
| Rbm4b | 3.627802461 |
| Tyro3 | 3.626936717 |
| Gem | 3.623259243 |
| Ptprm | 3.623247447 |
| Usp51 | 3.621837258 |
| Dsel | 3.621021758 |
| Sertad4 | 3.620075892 |
| Vgll4 | 3.618782098 |
| Ndrg3 | 3.614995602 |
| Abcd4 | 3.614327605 |
| Nudt2 | 3.611511263 |
| Klhl11 | 3.607773964 |
| Idh3g | 3.607263578 |
| Fxr1 | 3.606709287 |
| Spry4 | 3.604578307 |
| Gsto2 | 3.604321137 |
| Lss | 3.603180219 |
| Alkbh7 | 3.601004753 |
| Mrpl12 | 3.600734196 |
| Il1rl1 | 3.600699219 |
| Smim14 | 3.599921641 |
| Gusb | 3.599854737 |
| Kdm4d | 3.598954632 |
| Heg1 | 3.597256533 |
| Vps4a | 3.596277354 |
| Syce1l | 3.595686658 |
| Hdhd2 | 3.595339152 |
| Qdpr | 3.593676195 |
| Fem1b | 3.592816745 |
| Tnks1bp1 | 3.592675826 |
| Mrps14 | 3.584259199 |
| Gdap1 | 3.584137742 |
| Ncoa3 | 3.581892694 |
| Mybl1 | 3.578578404 |
| Shkbp1 | 3.569820112 |
| Pde10a | 3.568712779 |
| Dph1 | 3.562479287 |
| Nxt2 | 3.56140718 |
| Ssr1 | 3.560792964 |
| Fam43a | 3.559811036 |
| Hspbp1 | 3.558511935 |
| Sema6d | 3.557258882 |
| N4bp2l2 | 3.555761587 |
| Mettl10 | 3.55489401 |
| Adprhl1 | 3.553328269 |

| | |
|---|---|
| Gpr85 | 3.552168081 |
| Aen | 3.552030377 |
| Efna4 | 3.551467603 |
| Piga | 3.545520109 |
| Mgat5b | 3.545287037 |
| Vangl1 | 3.544926294 |
| Npas1 | 3.543388322 |
| Efnb2 | 3.541364147 |
| B3gat3 | 3.539777536 |
| Acin1 | 3.529205286 |
| Hbp1 | 3.526352332 |
| Clk3 | 3.525331413 |
| Cacnb1 | 3.521794609 |
| Glrx3 | 3.521607416 |
| Milr1 | 3.520641709 |
| Slc10a7 | 3.519800913 |
| Sin3a | 3.51945223 |
| Qtrt1 | 3.518528422 |
| Xkr8 | 3.518297941 |
| Tgfbr3 | 3.516658859 |
| Rab35 | 3.51652552 |
| Gon4l | 3.515279454 |
| Nyap1 | 3.513587702 |
| Tnfaip8l1 | 3.51330835 |
| Zbtb4 | 3.508123106 |
| Slc36a1 | 3.506128665 |
| Fam160b2 | 3.502390783 |
| Irak1bp1 | 3.502369521 |
| Itm2b | 3.501963856 |
| Smim12 | 3.499464507 |
| Stbd1 | 3.497317021 |
| Nptn | 3.496215135 |
| Myrip | 3.495785977 |
| Slc25a3 | 3.492701376 |
| Amer1 | 3.492279051 |
| Hs6st3 | 3.490704732 |
| Cul7 | 3.488730864 |
| Pik3r2 | 3.48687643 |
| Nos1ap | 3.483827317 |
| Eif1 | 3.481753791 |
| Klf8 | 3.48088229 |
| Gns | 3.476463329 |
| Ryk | 3.473661729 |
| Cmklr1 | 3.467780079 |
| Crtc2 | 3.464795356 |
| Anxa7 | 3.458377656 |
| Slc7a11 | 3.45664272 |

| | |
|---|---|
| Smc3 | 3.455797601 |
| Ssh3 | 3.455342585 |
| Snrk | 3.454011456 |
| Krt19 | 3.453241503 |
| Mpv17 | 3.452699207 |
| Atic | 3.451665936 |
| Prrg2 | 3.451250507 |
| Jak1 | 3.451160517 |
| Stk39 | 3.447431739 |
| Trim41 | 3.447033215 |
| Fkbp2 | 3.445818767 |
| Dsp | 3.445514524 |
| Spty2d1 | 3.443664941 |
| Sh3bp5 | 3.443583774 |
| B4galt7 | 3.441705257 |
| Ggn | 3.43965315 |
| Igf2bp2 | 3.438857659 |
| Sertad3 | 3.437000751 |
| Entpd7 | 3.433466541 |
| Rhof | 3.431464919 |
| Ddx59 | 3.426466319 |
| Pdgfa | 3.423931215 |
| Sdk1 | 3.421961222 |
| Exoc3 | 3.419366837 |
| Camkk1 | 3.418852799 |
| Kcnj14 | 3.417261699 |
| Egln1 | 3.416346953 |
| Kifc2 | 3.416039068 |
| Foxc1 | 3.415886852 |
| Zc3h8 | 3.415074048 |
| Ippk | 3.413165217 |
| Rsph4a | 3.412604102 |
| B3gnt5 | 3.410272979 |
| Lbr | 3.409708612 |
| Hcn3 | 3.404504812 |
| Notum | 3.398499956 |
| Tmem161a | 3.397153882 |
| Socs6 | 3.395035812 |
| Sox11 | 3.394113331 |
| Ndufb2 | 3.393336944 |
| Rp1l1 | 3.389932467 |
| Ubfd1 | 3.389584218 |
| Ankrd50 | 3.389379042 |
| Brca1 | 3.388810969 |
| Cstb | 3.385886146 |
| Cytl1 | 3.385794132 |
| Rnf180 | 3.385244255 |

| | |
|---|---|
| Cad | 3.38209258 |
| Prkce | 3.381815181 |
| Pddc1 | 3.381230012 |
| Cherp | 3.380633687 |
| Camk1d | 3.380226878 |
| Ccdc50 | 3.377652981 |
| Tle6 | 3.37702887 |
| Lrrc32 | 3.375880363 |
| Adprhl2 | 3.374986271 |
| Pank4 | 3.374262768 |
| Polr2l | 3.37014108 |
| Slc20a2 | 3.369965352 |
| Egln2 | 3.369030546 |
| Tbc1d7 | 3.368871002 |
| Spns2 | 3.368664895 |
| Ano5 | 3.367799773 |
| Klf13 | 3.367578116 |
| Ppp1r3d | 3.360798566 |
| Fuz | 3.35487269 |
| Frs2 | 3.354517257 |
| Mast4 | 3.354216037 |
| Prmt3 | 3.353148959 |
| Ttbk2 | 3.351953583 |
| Mmachc | 3.351874217 |
| Fip1l1 | 3.347137388 |
| Foxo6 | 3.342361652 |
| Grik4 | 3.342339298 |
| Pkp4 | 3.340557619 |
| Hltf | 3.338939188 |
| Lats1 | 3.338659908 |
| Ankrd33 | 3.337356105 |
| Gan | 3.336712477 |
| Ufsp2 | 3.336452492 |
| Prickle1 | 3.336062886 |
| Sirt6 | 3.330636655 |
| Dnm3 | 3.330167182 |
| Fah | 3.329081787 |
| Itpr1 | 3.326907349 |
| Iqch | 3.323986606 |
| Ppox | 3.322795459 |
| Calhm2 | 3.322472256 |
| Tmem97 | 3.319779589 |
| Myom3 | 3.319400517 |
| Pcna | 3.317559904 |
| Fat1 | 3.316359651 |
| Limk2 | 3.316237303 |
| Srgap2 | 3.315323863 |

| | |
|---|---|
| Lgalsl | 3.311871407 |
| Fbxw5 | 3.309163706 |
| Lrp11 | 3.307249546 |
| Coa6 | 3.306618063 |
| Prdm8 | 3.306551922 |
| Arhgap42 | 3.305872251 |
| Slbp | 3.30540712 |
| Prkcsh | 3.303691913 |
| Tmtc4 | 3.301406053 |
| Kank2 | 3.30034941 |
| Acvr2b | 3.29908542 |
| Uqcrfs1 | 3.299054514 |
| B4galt1 | 3.296562189 |
| Slc17a9 | 3.293879181 |
| Slc41a1 | 3.288416852 |
| Sec23a | 3.287403649 |
| Rhoq | 3.28525456 |
| Akap5 | 3.284408595 |
| Fam151b | 3.283831264 |
| Eme2 | 3.283795573 |
| Prr19 | 3.283186866 |
| Jrk | 3.281208879 |
| Primpol | 3.281178174 |
| Dnajc12 | 3.280483398 |
| Tceal8 | 3.277011557 |
| Klhl25 | 3.275951985 |
| Rhbdd3 | 3.275363853 |
| Tmem177 | 3.274264066 |
| Plxnb1 | 3.273698917 |
| Ctss | 3.272247636 |
| Tbc1d14 | 3.27035355 |
| Orai2 | 3.267424573 |
| Xrcc5 | 3.264942596 |
| Klhl22 | 3.264937389 |
| Oxct1 | 3.264779021 |
| Asf1a | 3.26254063 |
| Atf7ip | 3.262482513 |
| Fam213a | 3.261594769 |
| Rfxank | 3.255552762 |
| Ovca2 | 3.25533102 |
| Slc26a4 | 3.253418774 |
| Kcnj15 | 3.251331422 |
| Elk1 | 3.251293927 |
| Cxcl16 | 3.249763659 |
| Ercc4 | 3.248969774 |
| Engase | 3.24886489 |
| Daxx | 3.248148828 |

| | |
|---|---|
| Cdkn2d | 3.247375915 |
| Ncald | 3.246346769 |
| Rnft2 | 3.242517438 |
| Fgd5 | 3.241688134 |
| Cand2 | 3.241631783 |
| Arl6ip5 | 3.241064981 |
| Ankrd54 | 3.239303161 |
| Kbtbd11 | 3.238949151 |
| Nuak2 | 3.237707372 |
| Kdm2b | 3.235106023 |
| Ank3 | 3.235013907 |
| Mto1 | 3.234770914 |
| Ammecr1 | 3.234485019 |
| Tmem251 | 3.234238428 |
| Pxdn | 3.230802478 |
| Srgap1 | 3.229774043 |
| Rhoj | 3.226192364 |
| Nox4 | 3.224380924 |
| Fbxl20 | 3.223903249 |
| Apba1 | 3.222691851 |
| Fam222a | 3.222464167 |
| Dusp23 | 3.221862671 |
| Atp6v0a2 | 3.220684903 |
| Usp46 | 3.220273458 |
| Met | 3.219877798 |
| Slc27a1 | 3.218528868 |
| Cdhr3 | 3.216805565 |
| Mb21d2 | 3.216082778 |
| Ccdc134 | 3.216038976 |
| Tmem175 | 3.215753241 |
| Cdk5rap3 | 3.212235553 |
| Wnt5a | 3.211598747 |
| Il20rb | 3.209591272 |
| Wdr24 | 3.208653064 |
| Gnal | 3.208182634 |
| Kcnk6 | 3.204504242 |
| Mroh6 | 3.20398328 |
| Rprd1b | 3.20075786 |
| Pcdhb5 | 3.200682415 |
| Pde1c | 3.200421885 |
| Sdhaf1 | 3.200108949 |
| Lamc1 | 3.199976997 |
| Pcdhb7 | 3.1989512 |
| Rab11a | 3.198945038 |
| Leprot | 3.196070542 |
| Topbp1 | 3.195032969 |
| Inpp5d | 3.194582386 |

| | |
|---|---|
| Ost4 | 3.194308857 |
| Rps3 | 3.193597661 |
| Prdm1 | 3.193201055 |
| Jmy | 3.190964965 |
| Pnn | 3.190243358 |
| Vps26b | 3.188683835 |
| Nlrx1 | 3.187826465 |
| Cpne5 | 3.187247454 |
| Trmt6 | 3.185947758 |
| Fam131c | 3.182927434 |
| Pde3b | 3.182642558 |
| Nudt19 | 3.18115904 |
| Rfx3 | 3.179590741 |
| Pdha1 | 3.178888657 |
| Ssr2 | 3.1782585 |
| Ppif | 3.177671898 |
| Rpl36 | 3.177048538 |
| Smim13 | 3.176712629 |
| Vdac3 | 3.176552327 |
| Tamm41 | 3.175740504 |
| Tmem59l | 3.174211271 |
| Fbxo33 | 3.169664487 |
| Cdk6 | 3.164510685 |
| Lrrc8b | 3.164016413 |
| Enox1 | 3.160935539 |
| Srrd | 3.160752583 |
| Zbtb1 | 3.158254179 |
| Zfx | 3.158011591 |
| Txndc17 | 3.15757055 |
| Arpc5l | 3.156389458 |
| Ptdss2 | 3.156067925 |
| Txndc11 | 3.155214584 |
| Ufm1 | 3.154171211 |
| Eri3 | 3.153465003 |
| Thap11 | 3.153273572 |
| Exoc8 | 3.152545009 |
| Abca5 | 3.150178389 |
| Arrb2 | 3.150168818 |
| Fam189b | 3.148728295 |
| Nqo2 | 3.147936869 |
| Asf1b | 3.147217805 |
| Smg9 | 3.146800711 |
| Pcmtd1 | 3.145025827 |
| Tmem180 | 3.143931095 |
| Amotl1 | 3.143905712 |
| Gpr83 | 3.141682721 |
| Tmem200c | 3.14082421 |

| | |
|---|---|
| Cdk2ap1 | 3.140388927 |
| Arpc1a | 3.140379336 |
| Aurka | 3.139489939 |
| Rph3al | 3.13868787 |
| Fubp3 | 3.137600634 |
| Dmtf1 | 3.13691717 |
| Map3k11 | 3.134782372 |
| Znrd1 | 3.134459451 |
| Casp6 | 3.134422061 |
| Tmem109 | 3.133404744 |
| Akap1 | 3.132001355 |
| Ptma | 3.130338507 |
| Apod | 3.129920161 |
| Ggact | 3.128832248 |
| Sh3tc1 | 3.126027069 |
| Prrt2 | 3.125117349 |
| Ppic | 3.124190765 |
| Pak1 | 3.123911693 |
| Rpusd3 | 3.123144729 |
| Abhd17a | 3.122982439 |
| Per3 | 3.122216048 |
| Igfbp4 | 3.121353254 |
| Trim8 | 3.121041351 |
| Psmd2 | 3.120358822 |
| Arl2 | 3.120162005 |
| Scaf8 | 3.118978917 |
| Tacc1 | 3.118262293 |
| Shc2 | 3.116074061 |
| Hnmt | 3.114597097 |
| Slc33a1 | 3.114255048 |
| Senp1 | 3.109997836 |
| Raly | 3.106797315 |
| Ndufa5 | 3.105936362 |
| Pcdha7 | 3.10560374 |
| Gypc | 3.104161918 |
| Gps1 | 3.103083563 |
| Pdap1 | 3.102823979 |
| Srf | 3.099900531 |
| Ercc6l | 3.098311928 |
| Dnaja2 | 3.097215456 |
| Nr1d1 | 3.09617419 |
| Abhd16a | 3.09605732 |
| Kti12 | 3.095683727 |
| Ocrl | 3.094820871 |
| Ap5s1 | 3.093828006 |
| Tap1 | 3.093567171 |
| B3gnt7 | 3.093550438 |

| | |
|---|---|
| Degs1 | 3.09227344 |
| Tubal3 | 3.087870624 |
| Phf14 | 3.085534368 |
| Mfsd9 | 3.083166466 |
| Zswim4 | 3.082263072 |
| Dhx32 | 3.081091401 |
| Zer1 | 3.080914901 |
| Pdgfc | 3.080375151 |
| Sfxn3 | 3.080268617 |
| Tssc1 | 3.079639318 |
| Adam22 | 3.07935691 |
| Cenpj | 3.077006379 |
| Kdm5c | 3.076524602 |
| Dnajc4 | 3.076460504 |
| Tnpo3 | 3.076427544 |
| Ttyh3 | 3.07481572 |
| B3galt1 | 3.074388093 |
| Efna1 | 3.072718639 |
| Tmem106c | 3.072505092 |
| Setd5 | 3.071598292 |
| Tsen15 | 3.070822289 |
| Clcn7 | 3.070674341 |
| Ankrd44 | 3.069361117 |
| Ephx2 | 3.069272957 |
| Ercc5 | 3.066621955 |
| Arrdc1 | 3.066413357 |
| Kcnd1 | 3.065965636 |
| Bhlhe40 | 3.065661587 |
| Tdrd3 | 3.065186545 |
| Fam83d | 3.065165805 |
| Tnfrsf21 | 3.062925803 |
| Oscp1 | 3.061747924 |
| Git1 | 3.061634874 |
| Sfxn2 | 3.061566536 |
| Fam221a | 3.060780758 |
| Nelfcd | 3.060249904 |
| Rfx7 | 3.058305047 |
| Spint1 | 3.057732347 |
| Ubqln2 | 3.055786517 |
| Fam101b | 3.055053625 |
| Camta2 | 3.052972643 |
| Cmss1 | 3.052008631 |
| Nfatc4 | 3.050162955 |
| Tcea1 | 3.049037447 |
| Rfwd2 | 3.047475563 |
| Dse | 3.047352572 |
| Cmas | 3.045500966 |

| | |
|---|---|
| Satb1 | 3.045432144 |
| Ids | 3.042183935 |
| Sdad1 | 3.040123711 |
| Nnt | 3.039989166 |
| Pex5 | 3.037577219 |
| Bcas3 | 3.035226485 |
| Wdr37 | 3.0347456 |
| Hoxb4 | 3.034228261 |
| Rrnad1 | 3.034151081 |
| Cutc | 3.034074851 |
| Taf9b | 3.027433855 |
| Strada | 3.025800649 |
| Gnb5 | 3.024537713 |
| Pccb | 3.023858502 |
| Robo4 | 3.022857665 |
| Pak4 | 3.022506729 |
| Pcdhb13 | 3.022503019 |
| Phrf1 | 3.022456918 |
| Decr2 | 3.021088334 |
| Ube2e2 | 3.020331853 |
| Anxa4 | 3.020105081 |
| Kif3a | 3.019575998 |
| Igfbp2 | 3.019106157 |
| Serpine1 | 3.017978475 |
| Slc12a2 | 3.017248735 |
| Chuk | 3.016443359 |
| Asic1 | 3.01561627 |
| Ly75 | 3.015433449 |
| Top3a | 3.014973662 |
| Mtss1l | 3.014513295 |
| Gja4 | 3.014149125 |
| Micu1 | 3.01366186 |
| Mpp1 | 3.013353346 |
| Gpaa1 | 3.013067692 |
| Fdxr | 3.012355627 |
| Ttc28 | 3.011541952 |
| Scn10a | 3.010722447 |
| Tsr2 | 3.010458435 |
| Zcchc18 | 3.007190004 |
| Trmt1 | 3.005204564 |
| Dennd1b | 3.005199149 |
| Fam210b | 3.004722907 |
| Atraid | 3.003813524 |
| Mfsd10 | 3.001672159 |
| Ptprn2 | 2.999847719 |
| Rit1 | 2.999713651 |
| Psmd1 | 2.998892745 |

| | |
|---|---|
| Slc7a7 | 2.997506545 |
| Fam227a | 2.997328822 |
| Ctdspl | 2.997236895 |
| Kidins220 | 2.997208658 |
| Thumpd2 | 2.994543038 |
| Ndst2 | 2.994294959 |
| Usp44 | 2.993086236 |
| Itprip | 2.984781246 |
| Cox17 | 2.984698259 |
| Nudcd2 | 2.98446011 |
| Pwwp2a | 2.983501814 |
| Rfx1 | 2.98227776 |
| Sh3yl1 | 2.982227347 |
| Snx11 | 2.981902056 |
| Dusp18 | 2.981499769 |
| Itch | 2.97770189 |
| Nucb2 | 2.977398185 |
| Med1 | 2.976355246 |
| Dnajc22 | 2.974662888 |
| Gpr137c | 2.974297753 |
| Fkbp9 | 2.973037472 |
| Mrm1 | 2.972574504 |
| Tlcd2 | 2.972534572 |
| Uhrf2 | 2.972385993 |
| Mertk | 2.971851457 |
| Mpz | 2.968492719 |
| Arl14ep | 2.968416527 |
| Pyroxd2 | 2.967800956 |
| Mvd | 2.966925158 |
| Egr1 | 2.966140893 |
| Il17rd | 2.965381601 |
| Acvr1 | 2.965034747 |
| Atp8a1 | 2.964996734 |
| Zkscan1 | 2.963457464 |
| Chst2 | 2.962346704 |
| Tatdn1 | 2.961878241 |
| Lin7b | 2.956910471 |
| Psmd13 | 2.956341574 |
| Gla | 2.956251604 |
| Gng2 | 2.954153982 |
| Tomm34 | 2.954034978 |
| Opn3 | 2.952891898 |
| Prpf38a | 2.951151228 |
| Pdzd8 | 2.950208923 |
| Fam117b | 2.949879384 |
| Zc3h10 | 2.949765082 |
| Tgif1 | 2.949678185 |

| | |
|---|---|
| Dhrs7 | 2.949138895 |
| Hoxd3 | 2.949000317 |
| Mrpl43 | 2.94887407 |
| Lzic | 2.948839505 |
| Chml | 2.948342544 |
| Tbk1 | 2.947867526 |
| Bcap31 | 2.947237263 |
| Rasal3 | 2.946179796 |
| Cep152 | 2.945813651 |
| Ubqln1 | 2.945413627 |
| Eif2ak3 | 2.945345136 |
| Cmip | 2.945205906 |
| Mob1a | 2.944582668 |
| Paqr5 | 2.944182283 |
| Alpl | 2.943533599 |
| Dcaf8 | 2.942291375 |
| Gbp3 | 2.941029277 |
| Slc38a7 | 2.940224492 |
| Cbfa2t2 | 2.939723544 |
| Smc5 | 2.938073947 |
| Tjap1 | 2.937037529 |
| Ly6e | 2.93690695 |
| Bach1 | 2.93532292 |
| Lrrc73 | 2.935137979 |
| Cdc37l1 | 2.93504216 |
| Igf2bp1 | 2.933290639 |
| Nkrf | 2.933232348 |
| Tgm2 | 2.933166442 |
| Rbm12 | 2.931317474 |
| Elk4 | 2.930645556 |
| Afap1l1 | 2.929515111 |
| Junb | 2.927315551 |
| Park7 | 2.927154621 |
| Fgf11 | 2.927071368 |
| Zdhhc12 | 2.925339065 |
| Pitpna | 2.923492545 |
| Cd74 | 2.92312289 |
| Ociad2 | 2.921206843 |
| Armc5 | 2.920776167 |
| Isg20 | 2.920088641 |
| Dcaf5 | 2.919318472 |
| Urod | 2.918757255 |
| Fbxl14 | 2.918188212 |
| Klhdc8b | 2.917092221 |
| Zc3h13 | 2.916166807 |
| Tecr | 2.915081516 |
| Agmat | 2.914746157 |

| | |
|---|---|
| Morn3 | 2.914299226 |
| Flot1 | 2.914229227 |
| Tecta | 2.912609713 |
| Pdlim1 | 2.911910801 |
| Nup62 | 2.911396674 |
| Cdkl5 | 2.909830036 |
| Gtf2ird1 | 2.907794045 |
| Tnfrsf19 | 2.907403986 |
| Tmem120a | 2.907118219 |
| Galnt2 | 2.905751653 |
| Slc35c2 | 2.904883613 |
| Ankrd11 | 2.902380761 |
| Sntb2 | 2.902367064 |
| Tial1 | 2.901530043 |
| Adamts4 | 2.900645484 |
| Ccdc96 | 2.900340958 |
| Rhpn2 | 2.900095342 |
| Prkch | 2.898453692 |
| Flad1 | 2.8963579 |
| Mcoln1 | 2.896274916 |
| Dusp7 | 2.894230811 |
| Pold1 | 2.893932152 |
| Mtmr3 | 2.893079841 |
| Cntd1 | 2.891994562 |
| Crls1 | 2.891842595 |
| Men1 | 2.891544076 |
| Akr1a1 | 2.889997536 |
| Tigd2 | 2.889285006 |
| Mbip | 2.887647299 |
| Tctex1d1 | 2.887574464 |
| St6galnac2 | 2.887391498 |
| Kbtbd2 | 2.886674427 |
| Phlpp2 | 2.886303714 |
| Ttc37 | 2.885261145 |
| Zbtb21 | 2.884091219 |
| Bola3 | 2.883975958 |
| Phf3 | 2.883565242 |
| Ces4a | 2.883156934 |
| Gstm3 | 2.882748189 |
| Gldc | 2.879193492 |
| Ints5 | 2.878731873 |
| Sh3bp4 | 2.877420526 |
| Luc7l3 | 2.875987034 |
| Ptbp2 | 2.873914729 |
| Mast2 | 2.87362658 |
| Calcrl | 2.873469686 |
| Dab2 | 2.872303493 |

| | |
|---|---|
| Gys1 | 2.871571369 |
| Tlr4 | 2.870979179 |
| Cgnl1 | 2.870887753 |
| Ap3s2 | 2.868596407 |
| Rnf148 | 2.868419224 |
| Nt5m | 2.868316339 |
| Snrnp48 | 2.867492518 |
| Klhl17 | 2.86747973 |
| Nme4 | 2.865926994 |
| Hexim2 | 2.86573354 |
| Loxl2 | 2.863676228 |
| Atp6v1g2 | 2.863663354 |
| Cdc42ep2 | 2.86309677 |
| Tceal7 | 2.862632076 |
| Acot2 | 2.861696706 |
| Pdss2 | 2.861489567 |
| Fmo4 | 2.860429743 |
| Prkar1a | 2.86002102 |
| Ddx56 | 2.859751094 |
| Lage3 | 2.85917875 |
| Fam171a2 | 2.85872679 |
| Dnajb5 | 2.857298948 |
| Tpr | 2.856448908 |
| Mical1 | 2.855169872 |
| Pom121 | 2.853537811 |
| Mcm10 | 2.852177441 |
| Esd | 2.852074677 |
| Msl2 | 2.851732872 |
| Pqlc2 | 2.850950829 |
| Fbxl7 | 2.850492422 |
| Kank3 | 2.849949367 |
| Dok4 | 2.84960385 |
| Mpc1 | 2.849181725 |
| Snta1 | 2.847506592 |
| Crocc | 2.846823379 |
| Ddrgk1 | 2.846733042 |
| Sdcbp | 2.846248357 |
| Cdc6 | 2.84577044 |
| Pdlim3 | 2.844703513 |
| Tex15 | 2.844389726 |
| Usp27x | 2.844328699 |
| Mettl22 | 2.839784546 |
| Ddx3y | 2.839669271 |
| Gdpd5 | 2.839055961 |
| Chrna1 | 2.838617327 |
| Ubash3b | 2.838602569 |
| Appl2 | 2.838245511 |

| | |
|---|---|
| Acad10 | 2.834878385 |
| Ddr2 | 2.834453392 |
| Ppard | 2.834361353 |
| Avil | 2.832356729 |
| Anapc13 | 2.831390332 |
| Alas1 | 2.830781819 |
| Dusp12 | 2.830297713 |
| Kcns1 | 2.830279857 |
| Acadsb | 2.830123876 |
| Fam57a | 2.829120561 |
| Spcs3 | 2.828825139 |
| Mfap2 | 2.828289109 |
| Snf8 | 2.827603382 |
| Far1 | 2.827448479 |
| Ube2e3 | 2.827426136 |
| Ap1s1 | 2.827284493 |
| Rgag4 | 2.826415345 |
| Prex1 | 2.824663617 |
| Ankmy2 | 2.823164354 |
| Dcxr | 2.822897566 |
| Atxn7l3 | 2.822724699 |
| Slc44a1 | 2.822332776 |
| Dagla | 2.821817009 |
| Shisa2 | 2.820511276 |
| Adam10 | 2.820409494 |
| Ccdc112 | 2.820179502 |
| Desi2 | 2.819720555 |
| Arpc5 | 2.819174049 |
| Slc25a18 | 2.81850456 |
| Unc50 | 2.817845969 |
| Bre | 2.817375085 |
| Atp6ap2 | 2.817081072 |
| Pcdh15 | 2.81572806 |
| Arhgef7 | 2.812475752 |
| Them6 | 2.811573282 |
| Map2k3 | 2.811353048 |
| Ccne1 | 2.807447687 |
| Ankrd12 | 2.802735476 |
| Adcy4 | 2.801665012 |
| Plekhg3 | 2.801572597 |
| Etv1 | 2.800797446 |
| Guk1 | 2.800410945 |
| Dhtkd1 | 2.800294985 |
| Ly6g5c | 2.79960327 |
| Hivep3 | 2.798594164 |
| Hnrnpul1 | 2.797327006 |
| Mamdc2 | 2.796733462 |

| | |
|---|---|
| Bok | 2.796402515 |
| Wdr26 | 2.795603351 |
| Rpa1 | 2.795519561 |
| Styk1 | 2.795244524 |
| Parva | 2.792064814 |
| Acvrl1 | 2.791977817 |
| Tbce | 2.791465839 |
| Hnrnph1 | 2.789224083 |
| Thop1 | 2.788533495 |
| Lig4 | 2.786414879 |
| Bpgm | 2.786207906 |
| Adam15 | 2.785826135 |
| Suv420h2 | 2.785441038 |
| Wnt9a | 2.784114571 |
| Dpy19l2 | 2.783920989 |
| Mmd | 2.782632461 |
| Hscb | 2.781612025 |
| Pdhb | 2.781177506 |
| Itga10 | 2.779731612 |
| Il34 | 2.778397901 |
| Pcyt2 | 2.778134891 |
| Abcf2 | 2.777913246 |
| Prnp | 2.77766562 |
| Gpr162 | 2.777537983 |
| Gpr183 | 2.776447009 |
| Dlgap4 | 2.776032668 |
| Dusp4 | 2.775927738 |
| Fbxo45 | 2.775785214 |
| Dpp9 | 2.774079386 |
| Epcam | 2.773156919 |
| Arfgef2 | 2.773017057 |
| Lamb2 | 2.772167894 |
| Atxn7 | 2.772019799 |
| Mtf1 | 2.77196428 |
| Taf1a | 2.770346018 |
| Ube4b | 2.770085029 |
| Pigu | 2.769771308 |
| Tm6sf1 | 2.769371207 |
| Sun1 | 2.767803226 |
| Arhgef11 | 2.767778778 |
| Mamdc4 | 2.766891757 |
| Ncbp2 | 2.764710978 |
| Kat7 | 2.764707443 |
| Nlrp12 | 2.764026426 |
| Gpr68 | 2.763172355 |
| Sox13 | 2.76286994 |
| Ppp1cb | 2.762672732 |

| | |
|---|---|
| Gpam | 2.762641875 |
| Tuba4a | 2.760996389 |
| Trim59 | 2.759760977 |
| Creg2 | 2.759281016 |
| Wdr27 | 2.759265812 |
| Rab3gap2 | 2.759020939 |
| Arl6 | 2.7567931 |
| Ankrd36 | 2.756395591 |
| Metap1d | 2.756001121 |
| Stk10 | 2.753897348 |
| Tubb3 | 2.753107565 |
| Prpf18 | 2.752839026 |
| H6pd | 2.752174775 |
| Tmem186 | 2.75166316 |
| Bcl2l2 | 2.75034858 |
| Agap1 | 2.746921755 |
| Rasgef1a | 2.745942956 |
| Zbtb42 | 2.745138331 |
| Dctn5 | 2.745106288 |
| Preb | 2.744794049 |
| Thoc1 | 2.74471196 |
| Ncoa1 | 2.744360825 |
| Tinf2 | 2.742299061 |
| Sssca1 | 2.742257265 |
| Kctd11 | 2.740603124 |
| Chrac1 | 2.739598995 |
| Wfikkn1 | 2.73908666 |
| Lrp4 | 2.738654549 |
| Gtf2f1 | 2.738279438 |
| Tbcel | 2.738244724 |
| Anxa1 | 2.738219479 |
| Ift140 | 2.737291468 |
| Acot11 | 2.737070154 |
| Rnf181 | 2.735699831 |
| Ttc38 | 2.734107375 |
| Erich1 | 2.733801326 |
| Itpk1 | 2.733344762 |
| Pik3c2a | 2.733214461 |
| Utp23 | 2.730490973 |
| Lrrc28 | 2.729247466 |
| Hcfc1 | 2.728687051 |
| Itpka | 2.728615744 |
| Agpat3 | 2.727954311 |
| Asap1 | 2.727755346 |
| Tmed4 | 2.727336929 |
| Socs5 | 2.727132543 |
| Acot13 | 2.72614925 |

| | |
|---|---|
| Cdo1 | 2.725716116 |
| Naca | 2.725056068 |
| Gphn | 2.724875709 |
| Rims1 | 2.723776527 |
| Scn9a | 2.723628658 |
| Brap | 2.723214709 |
| Fbxo16 | 2.721430502 |
| Smim1 | 2.721360066 |
| Csnk1g3 | 2.720008425 |
| Pkig | 2.718829341 |
| Rab33a | 2.718142825 |
| Bdkrb2 | 2.717916926 |
| Apmap | 2.71674624 |
| Tmem170b | 2.716253194 |
| Igfbp6 | 2.715770258 |
| Esm1 | 2.715215043 |
| Klc4 | 2.715039584 |
| Dera | 2.71484315 |
| Arhgap24 | 2.714291023 |
| Chst12 | 2.714263095 |
| Cd163l1 | 2.713780092 |
| Fam214a | 2.713112401 |
| Ap5z1 | 2.71217949 |
| Cct6b | 2.710582225 |
| Dscr3 | 2.710304674 |
| Plekhn1 | 2.709052424 |
| Tmem173 | 2.70905234 |
| Gpatch1 | 2.707177041 |
| Tgfbr2 | 2.706639526 |
| Tmem234 | 2.705761673 |
| Rap2a | 2.705110009 |
| Jdp2 | 2.704232692 |
| Cep128 | 2.702576938 |
| Dhcr7 | 2.702265747 |
| Rdh11 | 2.701485392 |
| Hirip3 | 2.701189069 |
| Srxn1 | 2.698706791 |
| Arhgap33 | 2.698644116 |
| Usp15 | 2.698629786 |
| Lhfpl2 | 2.698197807 |
| Kat6a | 2.697782266 |
| Dvl1 | 2.697659132 |
| Ufd1l | 2.696855244 |
| Sccpdh | 2.696235041 |
| Asb6 | 2.695641646 |
| Gale | 2.69472595 |
| Nr3c2 | 2.69470423 |

| | |
|---|---|
| Snrnp25 | 2.691984792 |
| Eefsec | 2.691916665 |
| Pla2g15 | 2.691034914 |
| Adap1 | 2.690761428 |
| Zdhhc18 | 2.690757808 |
| Ipp | 2.690242442 |
| Zc3h12a | 2.689908561 |
| Pum1 | 2.688823065 |
| Csrnp1 | 2.688268707 |
| Lifr | 2.688068763 |
| Fam167b | 2.68789766 |
| Napepld | 2.687606901 |
| Trappc12 | 2.687286891 |
| Hoxd8 | 2.685508214 |
| Dedd2 | 2.685213573 |
| Hacl1 | 2.685062522 |
| Cdc23 | 2.684883076 |
| Rufy2 | 2.68454735 |
| Tlcd1 | 2.683986843 |
| Myct1 | 2.683741091 |
| Tmem100 | 2.682526439 |
| Rps2 | 2.682045905 |
| Lrrc8a | 2.681424776 |
| Celsr1 | 2.68135025 |
| Papd5 | 2.678548371 |
| Hoxc9 | 2.678512026 |
| Idua | 2.678044285 |
| Memo1 | 2.677309283 |
| Snapc2 | 2.675450723 |
| Mfsd6 | 2.67463692 |
| Zmat5 | 2.674328034 |
| Pnpla6 | 2.673426678 |
| Col8a2 | 2.673168497 |
| Gemin6 | 2.670061001 |
| Tesc | 2.669551689 |
| Sdr39u1 | 2.669096751 |
| Tie1 | 2.665035907 |
| Arhgef25 | 2.664874856 |
| Dna2 | 2.66421074 |
| Il17rc | 2.664076361 |
| Otub1 | 2.662473821 |
| Map3k5 | 2.661879563 |
| Larp1 | 2.661814503 |
| Card11 | 2.661804489 |
| Rps18 | 2.661560033 |
| Mrps21 | 2.661162254 |
| Mtmr12 | 2.659565748 |

| | |
|---|---|
| Sh3bgrl2 | 2.65919756 |
| Ilf2 | 2.657851881 |
| Ube2f | 2.657397487 |
| Pik3c2b | 2.656754125 |
| Cerk | 2.656356757 |
| Scp2 | 2.655689823 |
| Nono | 2.655669721 |
| Asphd2 | 2.655236248 |
| Gfi1 | 2.655069473 |
| Tmem68 | 2.653627131 |
| Nos1 | 2.65361065 |
| Atrx | 2.652939505 |
| Cd81 | 2.652722418 |
| Mkl1 | 2.651666023 |
| Chac2 | 2.651440271 |
| Sdf2l1 | 2.650754593 |
| Spire2 | 2.65073905 |
| Ganab | 2.650434915 |
| Smurf2 | 2.649711493 |
| Nat14 | 2.6488626 |
| Arl4d | 2.647487318 |
| Kcnk1 | 2.647116052 |
| Spag5 | 2.647052119 |
| Dennd6b | 2.645542107 |
| C2cd2l | 2.645421351 |
| Tesk1 | 2.644817723 |
| Glrx5 | 2.64367596 |
| Atp5h | 2.643374221 |
| Ntm | 2.641792976 |
| Qrsl1 | 2.641649464 |
| Vipas39 | 2.641241465 |
| Pstk | 2.640786813 |
| Kcnn1 | 2.639794019 |
| Otud7b | 2.638835838 |
| Chchd7 | 2.63845473 |
| Notch2 | 2.637350358 |
| Crip1 | 2.637289148 |
| Pbdc1 | 2.635935522 |
| Ak5 | 2.635257882 |
| Ccdc97 | 2.633990296 |
| Gtf2e1 | 2.633926912 |
| Acadvl | 2.633514918 |
| Chd1l | 2.633334388 |
| Dock6 | 2.631721043 |
| Capza1 | 2.631646106 |
| Klf4 | 2.630840401 |
| Spop | 2.630607272 |

| | |
|---|---|
| Atg2a | 2.628898903 |
| Tceb2 | 2.628795029 |
| Snw1 | 2.628645019 |
| Chmp1b | 2.628177157 |
| Tdrp | 2.624412699 |
| Ppp1r16b | 2.623019132 |
| Greb1 | 2.622400473 |
| Golm1 | 2.62162946 |
| Slc35b2 | 2.621624819 |
| Tom1l2 | 2.621117175 |
| Pld3 | 2.620173675 |
| Cox18 | 2.619958864 |
| Pla2g12a | 2.619388731 |
| Aimp1 | 2.619081865 |
| Ndrg4 | 2.618339001 |
| Thap7 | 2.617443659 |
| Mocs2 | 2.617151028 |
| Mgat2 | 2.61705788 |
| Mllt4 | 2.615723452 |
| Tpst2 | 2.61543375 |
| Uevld | 2.615020042 |
| Vcam1 | 2.61397688 |
| Ap1b1 | 2.611983275 |
| St8sia4 | 2.611827504 |
| Med24 | 2.611440019 |
| Bard1 | 2.610514466 |
| Plekhg2 | 2.610201013 |
| Dus1l | 2.60932617 |
| Erlec1 | 2.609318874 |
| Rrbp1 | 2.608454894 |
| Emc9 | 2.608262285 |
| Mmp17 | 2.607196674 |
| Pdxk | 2.605553206 |
| Gba2 | 2.604623111 |
| Zbtb48 | 2.601586591 |
| Tm4sf1 | 2.601181049 |
| Clasp1 | 2.598712324 |
| Matr3 | 2.597375305 |
| Usp18 | 2.597244728 |
| Gpr37 | 2.596998267 |
| H2afz | 2.596965812 |
| Ticam1 | 2.596834404 |
| C2cd4b | 2.595310677 |
| Acbd6 | 2.594803946 |
| Pitpnb | 2.593449958 |
| Mknk1 | 2.593322943 |
| Taf5l | 2.59266948 |

| | |
|---|---|
| Porcn | 2.592550387 |
| Noc2l | 2.590681883 |
| Caps2 | 2.589531306 |
| Sqstm1 | 2.589354258 |
| Ndufa3 | 2.589145998 |
| Jrkl | 2.587028767 |
| Suds3 | 2.585784495 |
| Tspan3 | 2.585623391 |
| Ogfod2 | 2.585428175 |
| Cul9 | 2.585289816 |
| Pag1 | 2.584451642 |
| Lamp2 | 2.584221251 |
| Cd44 | 2.583910793 |
| Hhip | 2.583096285 |
| Leprotl1 | 2.582194075 |
| Speg | 2.581373421 |
| Mvb12b | 2.579938302 |
| Mtif3 | 2.578875689 |
| Atf5 | 2.578440605 |
| Ankrd17 | 2.578182895 |
| Wnk4 | 2.578064616 |
| Aven | 2.576016834 |
| Thoc7 | 2.575959113 |
| E4f1 | 2.575698746 |
| Runx1 | 2.575161191 |
| Rsg1 | 2.575133178 |
| Uimc1 | 2.574888359 |
| Ccdc117 | 2.574442973 |
| Psma4 | 2.57423157 |
| Prep | 2.57286707 |
| Apcdd1 | 2.57283637 |
| Asphd1 | 2.572732704 |
| Sox2 | 2.572538986 |
| Dcaf13 | 2.572340214 |
| Ercc3 | 2.572218587 |
| Cast | 2.571035646 |
| Wdr18 | 2.570015411 |
| Smarcal1 | 2.570002688 |
| Ago4 | 2.569938199 |
| Nr1h3 | 2.569851852 |
| Xrcc1 | 2.569645641 |
| Pofut2 | 2.569050431 |
| Nop58 | 2.569010968 |
| Cd320 | 2.568649192 |
| Hspa12b | 2.565990916 |
| Dctpp1 | 2.564396353 |
| Tsc22d3 | 2.563714319 |

| | |
|---|---|
| Strn | 2.563551574 |
| Klhl18 | 2.56272009 |
| Nckap5 | 2.561502539 |
| Tspyl1 | 2.561227741 |
| Gipc2 | 2.560889848 |
| Iba57 | 2.560871034 |
| Nfyb | 2.560822364 |
| Vdac2 | 2.559832045 |
| Psmg4 | 2.559336994 |
| Fbxo43 | 2.558987323 |
| Trim46 | 2.558736294 |
| Dennd4b | 2.557508767 |
| Magi2 | 2.556630183 |
| Slirp | 2.55635408 |
| Atg3 | 2.55629603 |
| Emc7 | 2.55559806 |
| Sytl2 | 2.555364035 |
| Fabp5 | 2.554371776 |
| Mlf1 | 2.55416602 |
| Galnt13 | 2.553481363 |
| Psrc1 | 2.553188431 |
| Poc1b | 2.552923021 |
| Amn | 2.552665077 |
| Zfp36l1 | 2.551741572 |
| Pxmp4 | 2.550777642 |
| Dmtn | 2.550517649 |
| Map4k4 | 2.549302787 |
| Ifit2 | 2.548483422 |
| Pax6 | 2.548214003 |
| Cul4b | 2.547941784 |
| Ubc | 2.547806011 |
| Prdm2 | 2.547604509 |
| Tet3 | 2.547532968 |
| Spesp1 | 2.547311856 |
| Il18 | 2.546459925 |
| Dnajc3 | 2.545225208 |
| Dctd | 2.544819722 |
| Spata24 | 2.543320084 |
| Wdr61 | 2.542742745 |
| Rplp2 | 2.542204615 |
| C2cd4a | 2.54194439 |
| Fam124b | 2.541261485 |
| Rgp1 | 2.541023145 |
| Fam102a | 2.540846761 |
| Avl9 | 2.540633534 |
| Stk16 | 2.539605796 |
| Smn1 | 2.538618958 |

| | |
|---|---|
| Plvap | 2.538391556 |
| Srsf2 | 2.538057862 |
| Tpd52 | 2.537779723 |
| Got2 | 2.536281954 |
| Usp38 | 2.53601848 |
| Vhl | 2.535256719 |
| Arid3a | 2.534884437 |
| Bend7 | 2.534867083 |
| Lpl | 2.534695613 |
| Wars2 | 2.53428812 |
| Emp2 | 2.533987122 |
| Arhgef33 | 2.533780023 |
| Cldn10 | 2.533605575 |
| Ttll11 | 2.532948791 |
| Aar2 | 2.532918907 |
| Aldh1l2 | 2.532745572 |
| Slc12a4 | 2.532519377 |
| Cox5a | 2.531186071 |
| Dusp10 | 2.530609539 |
| Snx25 | 2.529982845 |
| Foxd2 | 2.529682061 |
| Hmgn2 | 2.529665021 |
| Ublcp1 | 2.528320741 |
| Sec14l2 | 2.528271695 |
| Coro1c | 2.528185662 |
| Ube2c | 2.527490724 |
| Igip | 2.527140495 |
| Ik | 2.524627099 |
| Tecpr2 | 2.52363096 |
| Xirp2 | 2.523511275 |
| Plekhb1 | 2.522418576 |
| Me3 | 2.520480097 |
| Eme1 | 2.519823112 |
| Ahnak | 2.519758632 |
| Crot | 2.51929057 |
| Ccpg1 | 2.518883977 |
| Ankrd24 | 2.518207626 |
| Pdcd6ip | 2.517848557 |
| Dnase1l1 | 2.517029675 |
| Tnfrsf14 | 2.516590337 |
| Gatad2b | 2.516340769 |
| Ago2 | 2.515751537 |
| Thap1 | 2.515657867 |
| Set | 2.515622375 |
| Mafb | 2.515312287 |
| Mapk3 | 2.514948585 |
| Tpcn2 | 2.514946522 |

| | |
|---|---|
| Cars | 2.51476645 |
| Nme7 | 2.514274664 |
| Dnajc17 | 2.513350731 |
| Wdr77 | 2.513090745 |
| Hyal1 | 2.508736994 |
| Tln1 | 2.507948605 |
| Prpsap1 | 2.507911823 |
| Tceanc2 | 2.507610777 |
| Cdh2 | 2.507348324 |
| Tead4 | 2.507103232 |
| Pebp1 | 2.506599695 |
| Asxl3 | 2.505829902 |
| Fads2 | 2.505329332 |
| Bcdin3d | 2.504724011 |
| Card9 | 2.503875446 |
| Sema3d | 2.503131673 |
| Med12 | 2.501968958 |
| Pogk | 2.501601289 |
| Poldip2 | 2.501287378 |
| Ccdc6 | 2.500824286 |
| Fbll1 | 2.500752084 |
| Grm8 | 2.500462693 |
| Mrpl51 | 2.499988524 |
| Thsd7b | 2.499404587 |
| Amot | 2.499377974 |
| Csmd2 | 2.498669764 |
| Psma3 | 2.498608498 |
| Tysnd1 | 2.498497307 |
| Nqo1 | 2.498074524 |
| Rhobtb1 | 2.497903963 |
| Uros | 2.497750279 |
| Pmvk | 2.496271045 |
| Rab11fip3 | 2.495497764 |
| Yes1 | 2.494955601 |
| Ppip5k1 | 2.494880936 |
| Kmt2c | 2.494632421 |
| Trib2 | 2.494335465 |
| Rabep1 | 2.493972015 |
| Zswim8 | 2.493030975 |
| Trim66 | 2.49284517 |
| Fbxl2 | 2.492726711 |
| Khnyn | 2.492363954 |
| Mad1l1 | 2.492013982 |
| Lamtor4 | 2.491328555 |
| Tspan14 | 2.491029128 |
| Neil2 | 2.490507333 |
| Mypop | 2.490465964 |

| | |
|---|---|
| Tspan6 | 2.48988606 |
| Pex6 | 2.489574843 |
| Itga8 | 2.489167336 |
| Ddx20 | 2.48831036 |
| Hnrnpk | 2.487944113 |
| Golga3 | 2.487321981 |
| Map2k6 | 2.487103401 |
| Bcl10 | 2.486562652 |
| Yars | 2.48600958 |
| Lrpap1 | 2.48591179 |
| Rrm1 | 2.48394629 |
| Mvk | 2.483109086 |
| Srebf1 | 2.482414058 |
| Ndufa12 | 2.481332876 |
| Cuta | 2.481319966 |
| Pip5k1c | 2.480993463 |
| Ddx55 | 2.479962161 |
| Atp6v1e2 | 2.478848126 |
| Npepl1 | 2.478808866 |
| Slc7a2 | 2.478536879 |
| Ech1 | 2.477622516 |
| Mtor | 2.477517345 |
| Rasgrp3 | 2.477274568 |
| Erh | 2.477148203 |
| Upf3a | 2.476995826 |
| Npas3 | 2.476794505 |
| Golga5 | 2.47579311 |
| Slc25a38 | 2.475195018 |
| Tm2d3 | 2.472864669 |
| Prrt3 | 2.472590307 |
| Apopt1 | 2.472236369 |
| Dynll2 | 2.471076549 |
| Calu | 2.468359719 |
| Mknk2 | 2.467767146 |
| Mpp2 | 2.465590283 |
| Zw10 | 2.46541922 |
| Khk | 2.464982808 |
| Ssr3 | 2.464230131 |
| Hs6st1 | 2.463135364 |
| Abca2 | 2.463069857 |
| Lrrc6 | 2.462204333 |
| Tbc1d17 | 2.461743888 |
| Hfm1 | 2.460753911 |
| Mxi1 | 2.460615485 |
| Mtmr7 | 2.460501405 |
| Naalad2 | 2.460113237 |
| Itgb3 | 2.460036937 |

| | |
|---|---|
| Scoc | 2.459827102 |
| Clcn6 | 2.459629645 |
| Pemt | 2.459195626 |
| Rnf25 | 2.459174406 |
| Muc4 | 2.459000728 |
| Mreg | 2.457806393 |
| Kdm5a | 2.457189076 |
| Entpd5 | 2.454940357 |
| Lrrfip2 | 2.454525288 |
| Kcnn4 | 2.454067463 |
| Slc16a1 | 2.453742995 |
| Trak1 | 2.453151883 |
| Ppih | 2.452615612 |
| Ube2s | 2.452161534 |
| Tada1 | 2.450728943 |
| Slc39a1 | 2.449768006 |
| Tff3 | 2.449368191 |
| Rpusd1 | 2.449226418 |
| Purb | 2.448331623 |
| Add3 | 2.447473768 |
| Pim2 | 2.447253146 |
| Ptprg | 2.44662033 |
| Cep97 | 2.446353916 |
| Sgcd | 2.445981444 |
| Vps13c | 2.44587415 |
| Limd1 | 2.445589566 |
| Mrpl55 | 2.445203982 |
| Fam131b | 2.444930276 |
| Dnajc30 | 2.444007825 |
| Cldn16 | 2.442997575 |
| Ndufa13 | 2.442516783 |
| Apob | 2.44219027 |
| Tmem158 | 2.442159511 |
| Tmem11 | 2.440802725 |
| Ak3 | 2.439882401 |
| Mall | 2.439738559 |
| Slc2a8 | 2.43877758 |
| Ssrp1 | 2.438734557 |
| Klhl42 | 2.437570244 |
| Atad1 | 2.436187309 |
| Lmbrd1 | 2.435580519 |
| Nudt7 | 2.435517083 |
| Mrpl39 | 2.435257565 |
| Lime1 | 2.433709487 |
| Fam122b | 2.432763302 |
| Fscn1 | 2.431984468 |
| Fam184a | 2.431245838 |

| | |
|---|---|
| Klhdc2 | 2.43104598 |
| Fam212a | 2.430942482 |
| Tmtc3 | 2.429362327 |
| Ephb1 | 2.428540661 |
| Flrt2 | 2.428484823 |
| Osbpl11 | 2.428244069 |
| Anp32b | 2.427390129 |
| Gcc2 | 2.427302178 |
| Iqcd | 2.426224677 |
| Tfpi | 2.425783441 |
| Etv4 | 2.42573317 |
| Elmo2 | 2.42539556 |
| Dync1li2 | 2.42509881 |
| Flna | 2.42462628 |
| Numb | 2.423963977 |
| Chmp1a | 2.423360101 |
| Atp6v1f | 2.422655743 |
| Lrrc3 | 2.422536712 |
| Snx21 | 2.4218987 |
| Gmpr | 2.420439512 |
| Phlda1 | 2.419140601 |
| Nsl1 | 2.418211509 |
| Ppp1r3f | 2.417812597 |
| Tymp | 2.417797463 |
| Limk1 | 2.417523808 |
| Fbn2 | 2.417269686 |
| Canx | 2.414050725 |
| Kcnj12 | 2.413838693 |
| Snurf | 2.413801629 |
| Spata7 | 2.411829735 |
| Itpr3 | 2.411426587 |
| Gss | 2.411204888 |
| Phactr1 | 2.411066146 |
| Med7 | 2.410925982 |
| Mplkip | 2.410816594 |
| Atg12 | 2.409933278 |
| Myo5b | 2.408650815 |
| Slc39a10 | 2.408557934 |
| Bmpr1a | 2.407274915 |
| Spef2 | 2.405678173 |
| Sod2 | 2.405105231 |
| Lurap1l | 2.404077726 |
| Creb3l4 | 2.40395317 |
| Scamp1 | 2.403771537 |
| Galk1 | 2.403770972 |
| Aagab | 2.403368347 |
| Tsg101 | 2.402689728 |

| | |
|---|---|
| Flrt3 | 2.402668252 |
| Gmppa | 2.402131109 |
| Msmp | 2.400688951 |
| Usp8 | 2.400403544 |
| Mfsd2b | 2.400288317 |
| Fam71f2 | 2.400170678 |
| St3gal2 | 2.400002746 |
| Ap2s1 | 2.399810706 |
| Pcdhb10 | 2.398901952 |
| Syt7 | 2.39826438 |
| Rpl36a | 2.397861437 |
| Smim3 | 2.397439841 |
| Poc5 | 2.394894287 |
| Ogg1 | 2.394820937 |
| Taf1c | 2.39460884 |
| Cnpy2 | 2.39451919 |
| Usp25 | 2.393315843 |
| Pgrmc2 | 2.393296343 |
| Bahcc1 | 2.390966308 |
| Pex10 | 2.390030742 |
| Hic2 | 2.389856795 |
| Yipf2 | 2.389422834 |
| Zbtb20 | 2.389410746 |
| Gas2l1 | 2.388694963 |
| Trim7 | 2.387761156 |
| Sbno2 | 2.38763396 |
| Chst10 | 2.386529368 |
| Fundc1 | 2.386348004 |
| Lpcat3 | 2.385862131 |
| Map2 | 2.385801204 |
| Dcun1d5 | 2.385025981 |
| Ldoc1 | 2.3846624 |
| Srsf9 | 2.384199855 |
| Snx9 | 2.384067349 |
| Lca5l | 2.383429161 |
| Mars2 | 2.38211768 |
| Ano10 | 2.381661231 |
| Med18 | 2.380503758 |
| Mrps27 | 2.379461801 |
| Mrps18b | 2.379303815 |
| Dusp22 | 2.378879242 |
| Bcl6 | 2.378687951 |
| Fsd1 | 2.37744555 |
| Mif4gd | 2.376385624 |
| Zfp69 | 2.376235271 |
| Mettl23 | 2.375570993 |
| Prpf3 | 2.375459467 |

| | |
|---|---|
| Angel2 | 2.375100923 |
| Pcdhb15 | 2.374749548 |
| Wdr31 | 2.373647687 |
| Timeless | 2.37157839 |
| Btg1 | 2.371165141 |
| Cebpz | 2.371137729 |
| Dtd1 | 2.37059058 |
| Sacm1l | 2.369455961 |
| Pomt2 | 2.369025011 |
| Fus | 2.368690462 |
| Clk4 | 2.368312676 |
| Prr12 | 2.366059996 |
| Bcl2l1 | 2.365841396 |
| Ptpn22 | 2.364285046 |
| Pnma1 | 2.362611192 |
| Yif1b | 2.362215291 |
| Hgsnat | 2.361837685 |
| S1pr3 | 2.360056569 |
| Syt17 | 2.359854637 |
| Slc25a12 | 2.359559004 |
| Cdk8 | 2.359554655 |
| Invs | 2.359476447 |
| Fam229b | 2.358952036 |
| Ccdc94 | 2.358426774 |
| Iffo2 | 2.357787939 |
| Med14 | 2.357338059 |
| Pdia5 | 2.35712511 |
| Ccdc122 | 2.356823371 |
| Ap2a2 | 2.356092603 |
| Asb9 | 2.355869809 |
| Clybl | 2.355625729 |
| Tnks | 2.354750149 |
| Scaf4 | 2.353933491 |
| Chtf8 | 2.35358974 |
| Nrip3 | 2.35348467 |
| Adprh | 2.352177772 |
| Ndufaf7 | 2.351920531 |
| Fbn1 | 2.351320407 |
| Nsdhl | 2.349820784 |
| Znf512b | 2.349589161 |
| Alox12 | 2.349195721 |
| Gna11 | 2.348645944 |
| Tmem258 | 2.347721038 |
| Nsun5 | 2.347102256 |
| Mrps9 | 2.345066924 |
| Slc9a5 | 2.3446227 |
| Slc7a6 | 2.344439097 |

| | |
|---|---|
| Bag3 | 2.344319551 |
| Fbxw9 | 2.343664301 |
| Kdr | 2.342768794 |
| Tmem79 | 2.341673187 |
| Suv39h1 | 2.340926228 |
| Kcns3 | 2.339768663 |
| Hipk2 | 2.33845225 |
| Epm2aip1 | 2.337353915 |
| Mmrn1 | 2.337233499 |
| Usp48 | 2.336403722 |
| Cbx7 | 2.336186039 |
| Lsm12 | 2.335545546 |
| Vat1l | 2.334963203 |
| Cdk1 | 2.334962901 |
| Dcun1d2 | 2.334068043 |
| Nphp3 | 2.333587873 |
| Cog1 | 2.333334477 |
| Mob1b | 2.33311129 |
| Sgsm1 | 2.332707503 |
| Camsap1 | 2.332114489 |
| Mbd4 | 2.33177076 |
| Oard1 | 2.331605823 |
| Jph3 | 2.331599968 |
| Aoc2 | 2.330631705 |
| Comt | 2.33015119 |
| Stk38 | 2.329847923 |
| Pawr | 2.329382037 |
| Akap12 | 2.329109956 |
| Abhd17b | 2.328671324 |
| Lpin1 | 2.327619628 |
| Epc2 | 2.327606161 |
| Tmem160 | 2.327480546 |
| Tcf7l1 | 2.326571994 |
| Inf2 | 2.325512568 |
| Eif1ad | 2.325328504 |
| Arl8b | 2.32514609 |
| Tbc1d23 | 2.324518644 |
| Ppp1r13b | 2.324442673 |
| Rps6ka3 | 2.324206688 |
| Emp3 | 2.324005779 |
| Psph | 2.323839169 |
| Ftsj2 | 2.323756856 |
| Palld | 2.322966797 |
| Tapbpl | 2.32245272 |
| Ehd1 | 2.321287606 |
| Fam20b | 2.320257429 |
| Itgb5 | 2.319926307 |

| | |
|---|---|
| Npl | 2.319867797 |
| Gtf2h4 | 2.319452373 |
| Cisd3 | 2.31848895 |
| Fam110a | 2.317634641 |
| Wdfy3 | 2.317586415 |
| Alad | 2.31745273 |
| Hus1 | 2.317332371 |
| Plcxd3 | 2.316547022 |
| Dguok | 2.316208239 |
| Smtn | 2.316183154 |
| Setd2 | 2.316109204 |
| Cdc42ep3 | 2.316064569 |
| Pdgfra | 2.316034845 |
| Adra2a | 2.31586767 |
| Daw1 | 2.314989866 |
| Adamts13 | 2.314833051 |
| Myo1f | 2.314306145 |
| Hemk1 | 2.314302017 |
| Usp13 | 2.313341033 |
| Gpr63 | 2.313272806 |
| Usp19 | 2.311549163 |
| Rps14 | 2.310824949 |
| Col14a1 | 2.310429601 |
| Shank3 | 2.310230404 |
| Dnah5 | 2.309592777 |
| Gabbr1 | 2.308312377 |
| Syt1 | 2.308077094 |
| Muc1 | 2.307402249 |
| Elk3 | 2.307339739 |
| Lpcat4 | 2.307210697 |
| Malsu1 | 2.307203115 |
| Vkorc1l1 | 2.306497532 |
| Rbmxl2 | 2.306062733 |
| Dhx57 | 2.305842968 |
| Prdm4 | 2.305676718 |
| Crlf3 | 2.30562615 |
| Maf1 | 2.304708787 |
| Zfyve28 | 2.304688522 |
| Ankib1 | 2.304511875 |
| Ip6k2 | 2.303696961 |
| Egflam | 2.303041015 |
| Col11a2 | 2.301565908 |
| Hagh | 2.299827595 |
| Phactr4 | 2.29963055 |
| Spin4 | 2.299089159 |
| Mtm1 | 2.298752315 |
| Cdkn2c | 2.298355998 |

| | |
|---|---|
| Usp39 | 2.297405483 |
| Zglp1 | 2.297330276 |
| Tnnt1 | 2.297303081 |
| Stox2 | 2.296145242 |
| Fbxw2 | 2.295935854 |
| Gnai2 | 2.295813762 |
| Nras | 2.295729974 |
| Ccdc13 | 2.295514133 |
| Slc35d2 | 2.295483712 |
| Mrpl50 | 2.295339974 |
| Acss2 | 2.295245938 |
| Bfsp1 | 2.292969731 |
| Sema6b | 2.292808364 |
| Ahsa2 | 2.292615193 |
| Pten | 2.29244419 |
| Rpl23a | 2.291700257 |
| Vimp | 2.291479039 |
| Usp36 | 2.290846055 |
| Wwc2 | 2.290094258 |
| Sugp2 | 2.289995691 |
| Bscl2 | 2.289825747 |
| Cyfip1 | 2.288941984 |
| Mrps18a | 2.288642949 |
| Sun2 | 2.287462844 |
| Trpt1 | 2.287457367 |
| Cdk20 | 2.286911475 |
| Adpgk | 2.286365821 |
| Alpk3 | 2.28603967 |
| Myl12b | 2.286006901 |
| Peak1 | 2.285418167 |
| Vps8 | 2.284521217 |
| Rftn1 | 2.283662613 |
| Pdzd11 | 2.283398819 |
| Pbx1 | 2.281528663 |
| Dgcr14 | 2.2809828 |
| Gng10 | 2.280478609 |
| Mesdc1 | 2.280096234 |
| Cmtm7 | 2.279745741 |
| Nob1 | 2.278993693 |
| Tnfaip3 | 2.278852854 |
| Acyp1 | 2.278791227 |
| Glipr2 | 2.278508066 |
| Ccdc169 | 2.278344516 |
| Nrg1 | 2.277262374 |
| Mrpl37 | 2.277063264 |
| Plcg2 | 2.277059702 |
| Cbs | 2.276809155 |

| | |
|---|---|
| Sema6a | 2.276713885 |
| St3gal5 | 2.276616964 |
| Fzd1 | 2.276149236 |
| Mrpl48 | 2.275804669 |
| Tmppe | 2.274998056 |
| Qsox2 | 2.274633035 |
| Pycard | 2.274627264 |
| Glg1 | 2.272111733 |
| Dhx58 | 2.271985417 |
| Btbd11 | 2.271971508 |
| Ar | 2.271956819 |
| Ccdc126 | 2.27188987 |
| Ndufs6 | 2.271833941 |
| Pqlc1 | 2.271775888 |
| Ptpn7 | 2.271761559 |
| Atf4 | 2.271199058 |
| Kctd20 | 2.271161063 |
| Slc9a6 | 2.271112021 |
| Cnnm4 | 2.270557459 |
| Prr7 | 2.270553234 |
| Six2 | 2.26932973 |
| Zswim5 | 2.268664721 |
| Plrg1 | 2.268406516 |
| B9d1 | 2.268061224 |
| Hcrtr1 | 2.267509153 |
| Pde3a | 2.267062747 |
| Htr1b | 2.26649367 |
| Scrn1 | 2.265873727 |
| Cdon | 2.26581313 |
| Sesn3 | 2.264161312 |
| Nxn | 2.263925667 |
| Araf | 2.263796725 |
| Taf6 | 2.263023316 |
| Scyl1 | 2.262505593 |
| Slc20a1 | 2.26221409 |
| Dnajc14 | 2.262141218 |
| Rhebl1 | 2.261717697 |
| Psmb9 | 2.260582944 |
| Rnf220 | 2.259183876 |
| Adat1 | 2.258334617 |
| Zfpl1 | 2.256875873 |
| Pappa | 2.256785539 |
| Me1 | 2.256217574 |
| Pfas | 2.255457366 |
| Ttn | 2.25482371 |
| Plekhf2 | 2.254425554 |
| Slc35g1 | 2.254422656 |

| | |
|---|---|
| Ccdc90b | 2.252827066 |
| Slc25a22 | 2.252641332 |
| Mvb12a | 2.251848028 |
| Polm | 2.251608902 |
| Ccnl2 | 2.250066934 |
| Dlst | 2.249964484 |
| Fzd5 | 2.249541918 |
| P4htm | 2.248982622 |
| Tprn | 2.248645501 |
| Panx1 | 2.248193518 |
| Mdp1 | 2.248136386 |
| Cep63 | 2.247974084 |
| Cdk5r1 | 2.247790144 |
| Plxnd1 | 2.246914719 |
| Paqr6 | 2.246784139 |
| Kctd6 | 2.24669547 |
| Slc2a6 | 2.24571767 |
| Psmd12 | 2.245578268 |
| Ddb2 | 2.24552328 |
| Nbea | 2.244618977 |
| Mtmr14 | 2.244366299 |
| Arhgap27 | 2.244301459 |
| Cyp27b1 | 2.244143675 |
| Atg7 | 2.243151429 |
| Fchsd2 | 2.242578461 |
| Dpysl2 | 2.242290489 |
| Cacng6 | 2.240816739 |
| Snx13 | 2.2405239 |
| Nog | 2.240334056 |
| Tmem67 | 2.238876971 |
| Tnfsf9 | 2.238660574 |
| Tnfrsf11b | 2.23830365 |
| Cobll1 | 2.238144224 |
| Cbwd1 | 2.237507348 |
| Cops7a | 2.237296979 |
| Gnb2l1 | 2.236882176 |
| Fitm2 | 2.236666065 |
| Rfc5 | 2.236617312 |
| Myadm | 2.236605893 |
| Tchp | 2.236381213 |
| Espl1 | 2.236357488 |
| Gpr180 | 2.236305426 |
| Bola2 | 2.236238062 |
| Rb1 | 2.2347123 |
| Cwf19l1 | 2.234429651 |
| Wdr78 | 2.234325416 |
| Glyr1 | 2.232915625 |

| | |
|---|---|
| Tsga10 | 2.232695936 |
| Mylk2 | 2.231380518 |
| Ndufa6 | 2.231198072 |
| Tbc1d25 | 2.230844404 |
| Rad21 | 2.230550204 |
| Smcr8 | 2.229500199 |
| Wtip | 2.229464646 |
| Eya3 | 2.229132029 |
| Mxra7 | 2.228838347 |
| Fbxo17 | 2.228660371 |
| Lpar4 | 2.227653188 |
| Adamts5 | 2.226987128 |
| Cdkl3 | 2.22658668 |
| Ogt | 2.226154901 |
| Racgap1 | 2.225342506 |
| Lztfl1 | 2.225089877 |
| Tenm3 | 2.224823324 |
| Gsk3a | 2.224506609 |
| Klhl21 | 2.224219223 |
| Spc24 | 2.223894475 |
| Adh5 | 2.22312159 |
| Pus7 | 2.223104392 |
| Chmp4b | 2.22301552 |
| Sema4a | 2.222461922 |
| Hdac7 | 2.222016657 |
| Vsig1 | 2.221899802 |
| Trdmt1 | 2.221752675 |
| Dtd2 | 2.221528151 |
| Dag1 | 2.220426339 |
| Hlx | 2.220346243 |
| Sptb | 2.219280124 |
| Cnot4 | 2.219225309 |
| Rnf146 | 2.218907516 |
| Slc35a2 | 2.218317242 |
| Fosb | 2.217780776 |
| Ruvbl2 | 2.217665158 |
| Rbbp4 | 2.217522502 |
| Zfp36 | 2.217505823 |
| Trip13 | 2.217407152 |
| S100a3 | 2.217309623 |
| Dgki | 2.21658874 |
| Acap3 | 2.216585223 |
| Ern1 | 2.216558288 |
| Slx1b | 2.216535688 |
| Dync1i2 | 2.214991416 |
| Gimap1 | 2.214803844 |
| Lrfn3 | 2.214689028 |

| | |
|---|---|
| Sumo1 | 2.214432909 |
| Tanc2 | 2.214069635 |
| Sos1 | 2.213905689 |
| Bora | 2.213550047 |
| Ddx27 | 2.21354447 |
| Atm | 2.213220875 |
| Msrb1 | 2.211214271 |
| Ankrd28 | 2.210671992 |
| Mthfd2l | 2.209948984 |
| Cdk5rap1 | 2.209856391 |
| Gabpb1 | 2.209854571 |
| Jam2 | 2.209837913 |
| Fbxo7 | 2.209593212 |
| Epha4 | 2.209554838 |
| Mdfi | 2.208163758 |
| Hspa8 | 2.207975714 |
| Gpr1 | 2.207891348 |
| Ncoa6 | 2.207880108 |
| Klhdc10 | 2.207813933 |
| Ccdc127 | 2.207499998 |
| Yap1 | 2.207315389 |
| Ndufa2 | 2.206980361 |
| Snapc4 | 2.206891252 |
| Myoz3 | 2.206625154 |
| Pgap1 | 2.205340317 |
| Sema7a | 2.204893587 |
| Cntf | 2.204588523 |
| Iars2 | 2.204282237 |
| Rabgef1 | 2.204125623 |
| Brf2 | 2.202764852 |
| Sec24b | 2.202715948 |
| Lctl | 2.202269817 |
| Cers4 | 2.201933328 |
| Cchcr1 | 2.201919351 |
| Igfbp5 | 2.201765128 |
| Cbll1 | 2.201273838 |
| Ercc2 | 2.200702983 |
| Fam168a | 2.200699198 |
| Nipal2 | 2.200616048 |
| Gpalpp1 | 2.200534411 |
| Klhl24 | 2.200303641 |
| Hsf4 | 2.200103652 |
| Slc4a2 | 2.199192906 |
| Ube2n | 2.198626094 |
| Ndufs8 | 2.197737951 |
| Sgms2 | 2.197542116 |
| Slc46a1 | 2.197004555 |

| | |
|---|---|
| Ears2 | 2.19621685 |
| Cyb561d1 | 2.195273855 |
| Pi16 | 2.194374266 |
| Ormdl1 | 2.193745888 |
| Scn1b | 2.193522313 |
| Samd11 | 2.193122315 |
| Ywhab | 2.192983316 |
| Slc6a6 | 2.191936079 |
| Ptch2 | 2.191482315 |
| Sepsecs | 2.191220864 |
| Rcor1 | 2.190679593 |
| Ostf1 | 2.190365087 |
| Arhgap28 | 2.190313881 |
| Psma1 | 2.189795966 |
| Myo1b | 2.189700494 |
| Mesdc2 | 2.189390796 |
| Acer2 | 2.188897725 |
| Gprc5d | 2.187425517 |
| Rras | 2.186914219 |
| Tma16 | 2.186813983 |
| L3mbtl3 | 2.186765018 |
| Slc22a23 | 2.186757629 |
| Ngrn | 2.186681455 |
| Xpo6 | 2.18625055 |
| Lrfn1 | 2.1861797 |
| Alpk1 | 2.185608805 |
| Nptx2 | 2.185352352 |
| Bcar3 | 2.184961083 |
| Ncbp1 | 2.18350237 |
| Ncoa4 | 2.182123141 |
| Mboat2 | 2.182054965 |
| Mettl18 | 2.181872199 |
| Lcn10 | 2.181251657 |
| Slain1 | 2.179928449 |
| Add1 | 2.179088563 |
| Adam23 | 2.178545156 |
| Irgq | 2.178398849 |
| Mlxip | 2.178163023 |
| Spin1 | 2.177207365 |
| Setd7 | 2.176918315 |
| Atp8b2 | 2.176624436 |
| Nbn | 2.17656273 |
| Ptk2 | 2.17615158 |
| Ankrd13b | 2.175882208 |
| Tmem128 | 2.17570611 |
| Cabyr | 2.175375608 |
| Spryd4 | 2.175295912 |

| | |
|---|---|
| Cbl | 2.173847841 |
| Leng8 | 2.17326037 |
| Rbm3 | 2.173084751 |
| Ttpal | 2.1727876 |
| Terf1 | 2.172460686 |
| Madd | 2.171414904 |
| Kcnab3 | 2.171014906 |
| Vps36 | 2.170647788 |
| Atrip | 2.169517913 |
| Tcf3 | 2.169049279 |
| Cyfip2 | 2.168961723 |
| Hps6 | 2.168802588 |
| Trmu | 2.168429171 |
| Cntrob | 2.168351964 |
| Hectd1 | 2.167418347 |
| Zc3hav1l | 2.16606095 |
| Npw | 2.165946531 |
| Alg5 | 2.16593314 |
| Acd | 2.165803457 |
| Mroh8 | 2.165697466 |
| Ercc8 | 2.165031739 |
| Cuedc1 | 2.164772073 |
| Slc16a13 | 2.164472916 |
| Ccdc80 | 2.164354726 |
| Cpox | 2.163933609 |
| Cldn23 | 2.163458088 |
| Rps21 | 2.163447244 |
| Iah1 | 2.162936696 |
| Isl1 | 2.162753566 |
| Mlx | 2.162407519 |
| Sac3d1 | 2.162378132 |
| Srr | 2.160766665 |
| Krit1 | 2.160713901 |
| Arl13b | 2.160654209 |
| Cdk5rap2 | 2.160155395 |
| Trpc3 | 2.158550203 |
| Usp4 | 2.158438492 |
| Dnah1 | 2.158217484 |
| Trappc2 | 2.157644399 |
| Mrpl35 | 2.157231675 |
| Krt80 | 2.15518966 |
| Trpv2 | 2.154863522 |
| Lancl2 | 2.154465499 |
| Cdca3 | 2.154421504 |
| Eif2ak2 | 2.154304607 |
| Tpk1 | 2.154166879 |
| Rai2 | 2.152814034 |

| | |
|---|---|
| Lrp1b | 2.152054126 |
| Ergic2 | 2.151631574 |
| Lcor | 2.150851656 |
| Txndc12 | 2.150719805 |
| Rpp40 | 2.148844852 |
| Stx6 | 2.148649467 |
| Wee1 | 2.14857351 |
| Fcho1 | 2.148043055 |
| Col16a1 | 2.147991692 |
| Enkur | 2.147964989 |
| Dnajb11 | 2.147648915 |
| Cnbp | 2.146345812 |
| Nme1 | 2.146343476 |
| Tefm | 2.146098722 |
| Eml4 | 2.145774963 |
| Map3k7 | 2.145525676 |
| Phc2 | 2.145372978 |
| Rnf138 | 2.145169877 |
| Tut1 | 2.144979534 |
| Dimt1 | 2.14469671 |
| Plcb4 | 2.14435102 |
| Hddc2 | 2.143974402 |
| Tmem98 | 2.143822595 |
| Mrvi1 | 2.14313301 |
| Tom1 | 2.142871476 |
| Cenpi | 2.142102457 |
| Dedd | 2.141893867 |
| Tomm40l | 2.141570107 |
| Kazald1 | 2.141499695 |
| Ndufb9 | 2.140069399 |
| Ap1g1 | 2.139582396 |
| Phb | 2.139101488 |
| Armcx3 | 2.138678328 |
| Fignl1 | 2.137868217 |
| Lrrc47 | 2.137402196 |
| Kdsr | 2.136803006 |
| Tmlhe | 2.136699973 |
| C1qtnf3 | 2.136495136 |
| Actl6a | 2.136164912 |
| Parp3 | 2.134241438 |
| Rab3ip | 2.134088937 |
| Nrcam | 2.133589392 |
| Bst2 | 2.133314 |
| Fndc4 | 2.131634319 |
| Mtdh | 2.131328917 |
| Prox1 | 2.130772842 |
| Ddhd1 | 2.130769171 |

| | |
|---|---|
| Uaca | 2.130481585 |
| Ushbp1 | 2.130347947 |
| Snx16 | 2.12981697 |
| Lst1 | 2.129654727 |
| Gmppb | 2.128709095 |
| Fubp1 | 2.128582942 |
| Scpep1 | 2.128357627 |
| Tdrd9 | 2.127442643 |
| Dennd5a | 2.127397992 |
| Ing5 | 2.127289734 |
| Osbpl10 | 2.126841833 |
| Immt | 2.126099037 |
| Nr2f1 | 2.126068136 |
| Rere | 2.125836286 |
| Eea1 | 2.125832421 |
| Trim58 | 2.125813095 |
| Psat1 | 2.125453364 |
| Rft1 | 2.125009565 |
| Ten1 | 2.124789566 |
| Foxp1 | 2.124453228 |
| Vps18 | 2.124419103 |
| Angptl1 | 2.124330237 |
| Mpp3 | 2.124100871 |
| Nop56 | 2.12336913 |
| Dzip1l | 2.123033877 |
| Tubgcp6 | 2.122356903 |
| Cnnm3 | 2.122319266 |
| Plag1 | 2.122241673 |
| Akap2 | 2.122158595 |
| Gorasp1 | 2.122019804 |
| Hvcn1 | 2.121957533 |
| Ccdc69 | 2.121297554 |
| Pdlim2 | 2.120878878 |
| Alkbh6 | 2.120662324 |
| Rcn2 | 2.120580306 |
| Nt5dc1 | 2.120202941 |
| Fbln5 | 2.120166047 |
| Mex3a | 2.119631083 |
| Parp10 | 2.11916982 |
| Eva1a | 2.119030449 |
| Glb1l2 | 2.117944986 |
| Trip4 | 2.117797268 |
| Kcmf1 | 2.116787514 |
| Med4 | 2.11654061 |
| Mfsd4 | 2.116375481 |
| Cenpu | 2.115998388 |
| Tmx4 | 2.115624359 |

| | |
|---|---|
| Rad54l | 2.114918094 |
| Mrps23 | 2.114724587 |
| Kif3b | 2.114437039 |
| Carhsp1 | 2.114111057 |
| Emc10 | 2.114078048 |
| Ypel2 | 2.1138557 |
| Mea1 | 2.113077932 |
| Spag9 | 2.112442984 |
| Rpap3 | 2.112226404 |
| Smoc1 | 2.11195878 |
| Foxn3 | 2.111727911 |
| Coro2b | 2.111377515 |
| Hist3h2a | 2.110576328 |
| Nars2 | 2.109626527 |
| Sfrp1 | 2.109537786 |
| Ier5l | 2.10896669 |
| Mrps22 | 2.108098484 |
| Klhl6 | 2.107521415 |
| Slc35a5 | 2.106430837 |
| Gbf1 | 2.105999586 |
| Amy2b | 2.105891086 |
| Plcd3 | 2.105610551 |
| Eif4a1 | 2.105164966 |
| Rassf8 | 2.1041602 |
| Pja1 | 2.104127027 |
| Ebna1bp2 | 2.103668642 |
| Wdr47 | 2.103633888 |
| Clcn5 | 2.103275059 |
| Wdr41 | 2.102901696 |
| Mfap3l | 2.102681423 |
| Id1 | 2.102549808 |
| Nit1 | 2.102534038 |
| Pard3 | 2.101436267 |
| Scap | 2.100507296 |
| Mrto4 | 2.099939702 |
| Cdip1 | 2.098005572 |
| Prmt2 | 2.097905828 |
| Brd7 | 2.097567332 |
| Col4a2 | 2.097539592 |
| Rchy1 | 2.097095756 |
| Hfe | 2.09641309 |
| Tek | 2.095888013 |
| Arid2 | 2.09537743 |
| Ctbp2 | 2.094273267 |
| Bola1 | 2.094154688 |
| Arsj | 2.094133149 |
| Marveld1 | 2.092777369 |

| | |
|---|---|
| Dennd5b | 2.092726626 |
| Eif5a | 2.092710631 |
| Angel1 | 2.092209252 |
| Rab2b | 2.092068353 |
| Pepd | 2.090514203 |
| Glrx2 | 2.089958087 |
| Prdm5 | 2.089315884 |
| Nek2 | 2.089053145 |
| Hdac3 | 2.088928506 |
| Setd4 | 2.088429285 |
| Tmem116 | 2.088211855 |
| B4galt2 | 2.088177029 |
| Wac | 2.087834816 |
| Atp11a | 2.087583421 |
| Pcsk6 | 2.087504019 |
| Fancf | 2.086379744 |
| Armc4 | 2.084514506 |
| Tmbim1 | 2.084070563 |
| Stambp | 2.083381945 |
| Plcb3 | 2.083262938 |
| Maz | 2.082785565 |
| Zfyve21 | 2.082715074 |
| Specc1l | 2.082266515 |
| Lysmd4 | 2.082148296 |
| Fam134c | 2.081296522 |
| Polh | 2.081267256 |
| Emd | 2.080738094 |
| Fam120a | 2.080510867 |
| Rbm41 | 2.080195464 |
| Gdpd3 | 2.08016402 |
| Haus1 | 2.08000113 |
| Fam171b | 2.079946065 |
| Hoxb7 | 2.079344619 |
| Arhgef12 | 2.078724818 |
| Rpl35 | 2.078570688 |
| Tep1 | 2.077833504 |
| Fam185a | 2.077640733 |
| Mtr | 2.07761449 |
| Ube2j2 | 2.076794537 |
| Atp6v0e2 | 2.076503361 |
| Pole | 2.076132324 |
| Mettl6 | 2.075471293 |
| Spag7 | 2.074698197 |
| Map6 | 2.073626783 |
| Vma21 | 2.07284073 |
| Angptl2 | 2.07280667 |
| Grasp | 2.07251584 |

| | |
|---|---|
| Usp12 | 2.072117575 |
| Prx | 2.072041109 |
| Cpne1 | 2.071075675 |
| Scarf1 | 2.070855498 |
| Pet100 | 2.069758295 |
| Sigmar1 | 2.069294168 |
| Mrps28 | 2.069189824 |
| Epdr1 | 2.068973842 |
| Crebzf | 2.068957031 |
| Procr | 2.068046794 |
| Cda | 2.067622953 |
| Tcfl5 | 2.067318303 |
| Srp9 | 2.065679583 |
| Mib2 | 2.065319208 |
| Pmaip1 | 2.064463562 |
| Dhrs11 | 2.063944098 |
| Kif11 | 2.063763022 |
| Dgkh | 2.06340953 |
| Ccdc103 | 2.063403145 |
| Cd164 | 2.063210565 |
| Scamp5 | 2.063122439 |
| Dzip1 | 2.06304287 |
| Homer3 | 2.0618657 |
| Zhx2 | 2.061730633 |
| Chm | 2.061666736 |
| Efna5 | 2.061552838 |
| Pgm1 | 2.060666777 |
| Tnpo2 | 2.059537074 |
| Cstf2 | 2.058802628 |
| Arhgef15 | 2.058704714 |
| Trmt10b | 2.057997348 |
| Ndn | 2.056938796 |
| Tpx2 | 2.056755149 |
| Reep5 | 2.055756563 |
| Kat2a | 2.055306991 |
| Isca1 | 2.055233333 |
| Atxn3 | 2.055070138 |
| Elof1 | 2.054904195 |
| Lrrc4b | 2.054711407 |
| Aldoc | 2.052823261 |
| Rnf19a | 2.052804772 |
| Tpgs2 | 2.052775772 |
| Gxylt1 | 2.052544749 |
| Atg4a | 2.051626497 |
| Bach2 | 2.051103399 |
| Cks1b | 2.050106877 |
| Cap1 | 2.049256941 |

| | |
|---|---|
| Uba7 | 2.048731928 |
| Nipbl | 2.048554549 |
| Lysmd2 | 2.04809899 |
| Stx7 | 2.047980912 |
| Aak1 | 2.047651675 |
| Msi2 | 2.047171571 |
| Cdk9 | 2.046960224 |
| Phf2 | 2.046944311 |
| Rnf39 | 2.046293709 |
| Arl4c | 2.045838164 |
| Znhit6 | 2.045740618 |
| Papd4 | 2.044742529 |
| Coa5 | 2.044516251 |
| Afap1 | 2.044486981 |
| Ctbp1 | 2.04442262 |
| Nsun4 | 2.043975069 |
| Rbm44 | 2.043596709 |
| Map4k3 | 2.042696936 |
| Pianp | 2.042261327 |
| Pcyt1a | 2.041187736 |
| Sema3f | 2.040667814 |
| Stx2 | 2.040555294 |
| Rassf3 | 2.039603474 |
| Trim65 | 2.039550163 |
| Itgbl1 | 2.039175383 |
| Aatf | 2.038596711 |
| Sergef | 2.038071728 |
| Arhgdig | 2.036830295 |
| Mkl2 | 2.036540882 |
| Foxj2 | 2.036534848 |
| Hdac11 | 2.03649934 |
| Mybbp1a | 2.036052267 |
| Dysf | 2.035490669 |
| Slc39a4 | 2.035102908 |
| Mfsd1 | 2.034853677 |
| Rmdn3 | 2.034668515 |
| Vbp1 | 2.034488771 |
| Dhh | 2.033666708 |
| Klhl2 | 2.033587804 |
| Syp | 2.033164089 |
| Ngfrap1 | 2.032180791 |
| Narfl | 2.03209485 |
| Psmb2 | 2.031801344 |
| Smarcb1 | 2.031749417 |
| Svip | 2.031604982 |
| Tln2 | 2.030554392 |
| Rbm4 | 2.030527942 |

| | |
|---|---|
| Lsg1 | 2.030414577 |
| Fth1 | 2.030230517 |
| Lamtor3 | 2.030126149 |
| Pibf1 | 2.029911702 |
| Syt5 | 2.029494405 |
| Med12l | 2.029475117 |
| Med16 | 2.02672308 |
| Tiam1 | 2.025462203 |
| Rnf216 | 2.025394481 |
| Lmo7 | 2.024791774 |
| Ddx26b | 2.024483948 |
| Nkx6-2 | 2.024176626 |
| Ppp1r37 | 2.023849684 |
| Ppp2r1b | 2.023687037 |
| Cdan1 | 2.023049833 |
| Rps6kb2 | 2.022934734 |
| Cds2 | 2.022745045 |
| Actr5 | 2.022096138 |
| Sipa1l3 | 2.021665015 |
| L2hgdh | 2.020842084 |
| Eif2b1 | 2.020719106 |
| Eid3 | 2.020626053 |
| Ivd | 2.020402323 |
| Lypd6 | 2.019965025 |
| Ccdc146 | 2.019947898 |
| Nap1l5 | 2.019911328 |
| Slain2 | 2.019646978 |
| Dtwd2 | 2.018613789 |
| Nanos3 | 2.018248239 |
| Fkbp7 | 2.017822961 |
| Cnpy3 | 2.017400764 |
| Fam107a | 2.017062608 |
| Tob1 | 2.016850317 |
| Cln8 | 2.016269893 |
| Snd1 | 2.016257297 |
| Fech | 2.015788915 |
| Ssr4 | 2.015577314 |
| Sft2d1 | 2.014985925 |
| Zfand4 | 2.014675389 |
| Knstrn | 2.014583874 |
| Lrch2 | 2.013919108 |
| Cdrt4 | 2.013826019 |
| Ulk1 | 2.011975289 |
| Mkks | 2.011892923 |
| Zwint | 2.011729435 |
| Senp2 | 2.010715007 |
| Arfgef1 | 2.009767271 |

| | |
|---|---|
| Grik5 | 2.008865037 |
| Aox1 | 2.008774088 |
| Shroom4 | 2.008714799 |
| Tmem233 | 2.008176845 |
| Zfpm1 | 2.008162885 |
| Ano8 | 2.007644606 |
| Fam104a | 2.005878259 |
| Map3k2 | 2.004368713 |
| Cish | 2.003838855 |
| Sema4c | 2.00363672 |
| Eogt | 2.003219373 |
| Id2 | 2.003212215 |
| Eif3d | 2.003018474 |
| Fig4 | 2.002823734 |
| Gimap8 | 2.001969404 |
| Nup160 | 2.001894722 |
| Dst | 2.001767327 |
| Rbbp9 | 2.001561807 |
| Pvrl3 | 2.001165298 |
| Mroh1 | 2.00098466 |
| Rbm28 | 2.000920015 |
| Gcc1 | 2.000572309 |
| Cep19 | 2.000143936 |
| Uggt1 | 1.999527305 |
| Rtn3 | 1.99946427 |
| Gar1 | 1.998759542 |
| Adamtsl4 | 1.998751991 |
| Ppm1a | 1.998666314 |
| Exosc5 | 1.998624155 |
| Ankrd40 | 1.998556376 |
| Ece1 | 1.998085988 |
| Klf9 | 1.997863523 |
| Creld2 | 1.997660565 |
| Ptrhd1 | 1.997310509 |
| Tsen2 | 1.997190178 |
| Dnajc19 | 1.996967152 |
| Nop10 | 1.996832465 |
| Crcp | 1.996570181 |
| Hspa1b | 1.996169997 |
| Ralgps2 | 1.995878367 |
| Arfgap1 | 1.995245044 |
| Tmem108 | 1.995069228 |
| Lcorl | 1.994280662 |
| Senp6 | 1.994152589 |
| Tbc1d9 | 1.994037951 |
| Prex2 | 1.993961314 |
| Commd9 | 1.993806854 |

| | |
|---|---|
| Msh3 | 1.993717657 |
| Arhgdib | 1.993361416 |
| Tst | 1.992663923 |
| Bmpr1b | 1.99231307 |
| S100a16 | 1.991663601 |
| Morn2 | 1.991462343 |
| Nin | 1.991069081 |
| Tmem184c | 1.990442415 |
| Actn4 | 1.990184238 |
| Ado | 1.989558283 |
| Klc1 | 1.989266207 |
| Cyb5r1 | 1.988814935 |
| Draxin | 1.988053866 |
| Kif18b | 1.987949644 |
| Yeats4 | 1.987829326 |
| Ccdc88b | 1.98773109 |
| Agbl3 | 1.987565188 |
| Socs7 | 1.987320808 |
| Megf9 | 1.987212669 |
| Urb1 | 1.987074974 |
| Ppie | 1.986945558 |
| Vps37b | 1.986775711 |
| Plekhj1 | 1.986609333 |
| Tufm | 1.98480325 |
| Rbm18 | 1.984620606 |
| Ganc | 1.984569691 |
| Pmpcb | 1.984334848 |
| Dnaja4 | 1.98360724 |
| Fam174b | 1.982928469 |
| Xaf1 | 1.982901533 |
| Slc5a2 | 1.981418637 |
| Zcchc6 | 1.981038121 |
| Skp2 | 1.980554614 |
| Dock9 | 1.980311751 |
| Pex26 | 1.980071446 |
| Pcyox1 | 1.979788636 |
| Itgb8 | 1.979336334 |
| Gopc | 1.979062726 |
| Sf3b5 | 1.977937781 |
| Lamtor5 | 1.97764177 |
| Chmp5 | 1.977454965 |
| F3 | 1.977408604 |
| Sp2 | 1.97715371 |
| Ccdc137 | 1.977083281 |
| Sp110 | 1.976546883 |
| Dll3 | 1.975861085 |
| Lrrk2 | 1.974972784 |

| | |
|---|---|
| Sacs | 1.974677975 |
| Nme2 | 1.974633167 |
| Epn1 | 1.974579626 |
| Rbms3 | 1.974439852 |
| Rrp9 | 1.974034183 |
| Dtx2 | 1.973957966 |
| Ercc1 | 1.973673387 |
| Zswim6 | 1.972630032 |
| Fbxl8 | 1.972127304 |
| Repin1 | 1.971720987 |
| Pfdn1 | 1.970937129 |
| Slc51a | 1.970438754 |
| Kcnf1 | 1.970385584 |
| Slc9a1 | 1.970097565 |
| Letmd1 | 1.969835498 |
| Rbms2 | 1.969769012 |
| Rad54l2 | 1.969362826 |
| Nop16 | 1.969234003 |
| Tmed9 | 1.969217993 |
| Uvssa | 1.969053434 |
| Dpm2 | 1.968983069 |
| Tmem204 | 1.968801117 |
| Ctsk | 1.968401271 |
| Cnrip1 | 1.967988585 |
| Slc25a19 | 1.967454983 |
| Slc38a10 | 1.966441745 |
| Cldn11 | 1.966332771 |
| Vamp7 | 1.966227608 |
| Calm2 | 1.966146041 |
| Slc43a1 | 1.965833408 |
| Anks1b | 1.965760777 |
| Tars | 1.965238796 |
| Rnf185 | 1.964668804 |
| Bex2 | 1.964489454 |
| Ccdc51 | 1.964157751 |
| Rem1 | 1.963830157 |
| Hdhd3 | 1.963239837 |
| Syt15 | 1.963233972 |
| Ssna1 | 1.961560588 |
| Rpap2 | 1.960924658 |
| Tmem129 | 1.960253631 |
| Sgsh | 1.960026482 |
| Plekhh2 | 1.959900114 |
| Snai2 | 1.959157263 |
| Pcnxl3 | 1.959098824 |
| Rundc3b | 1.958635312 |
| Nsmce4a | 1.958286856 |

| | |
|---|---|
| Adam17 | 1.958123372 |
| Dcaf7 | 1.957879183 |
| Hnrnpr | 1.957832991 |
| Zik1 | 1.956421166 |
| Tm9sf1 | 1.956389785 |
| Lypd5 | 1.956219262 |
| Arsb | 1.955924813 |
| Hic1 | 1.955801247 |
| Rps15 | 1.955463578 |
| Phf21a | 1.955291664 |
| Stpg1 | 1.954663078 |
| Mmgt1 | 1.954435225 |
| Arhgef37 | 1.954277379 |
| She | 1.954010832 |
| Chchd6 | 1.952381817 |
| Timm17b | 1.952266219 |
| Taf15 | 1.951138304 |
| Entpd3 | 1.949689347 |
| Ace | 1.94964357 |
| Slc22a18 | 1.949362927 |
| Ssu72 | 1.949291759 |
| Fundc2 | 1.948971628 |
| Plekha2 | 1.948948706 |
| Irf7 | 1.948614764 |
| Zfp2 | 1.94838979 |
| Arhgef9 | 1.94834226 |
| C1qtnf4 | 1.948331517 |
| Wdr53 | 1.948210878 |
| Fxyd5 | 1.947972874 |
| Murc | 1.947939595 |
| Arl6ip6 | 1.947687296 |
| Mfhas1 | 1.947487092 |
| Tgds | 1.947418644 |
| Pomc | 1.947303835 |
| Unc5b | 1.946717152 |
| Ddb1 | 1.946352851 |
| Dgcr8 | 1.946256444 |
| Ccz1 | 1.946060026 |
| Peg3 | 1.945465312 |
| Ggt5 | 1.945329378 |
| Lyrm9 | 1.944826884 |
| Aamp | 1.944474978 |
| Elp2 | 1.944456463 |
| Gprasp2 | 1.944261453 |
| Rab31 | 1.944018118 |
| Hebp1 | 1.943991646 |
| Fastkd3 | 1.94371368 |

| | |
|---|---|
| Rnf152 | 1.943594682 |
| Rgl3 | 1.943352028 |
| Gab1 | 1.942763439 |
| Pcdhb2 | 1.942245406 |
| Zfyve1 | 1.942184983 |
| Trafd1 | 1.941971806 |
| Osbpl2 | 1.941895281 |
| Ppp4r2 | 1.94173534 |
| Il17ra | 1.941576582 |
| Ntsr1 | 1.940985578 |
| Lmcd1 | 1.939754545 |
| Phldb3 | 1.939551361 |
| Becn1 | 1.939302988 |
| Sema3g | 1.938511929 |
| Mus81 | 1.937943089 |
| Frs3 | 1.93785196 |
| Rpp14 | 1.937744698 |
| Ppm1m | 1.93758778 |
| Rpp38 | 1.936680914 |
| Naa60 | 1.936545535 |
| Rnf170 | 1.935522855 |
| Apoo | 1.935085003 |
| Strap | 1.935058288 |
| Helb | 1.934734759 |
| Mthfd1 | 1.933378441 |
| Gnpat | 1.932350563 |
| Ttll7 | 1.932208216 |
| Cacnb3 | 1.931494461 |
| Sec24c | 1.931431592 |
| Kbtbd4 | 1.93142946 |
| Tmem60 | 1.931143732 |
| Ctdp1 | 1.930785699 |
| Mbd6 | 1.929857015 |
| Amdhd2 | 1.9298207 |
| Dhrs13 | 1.929630745 |
| Hoxb8 | 1.929357733 |
| Mzt1 | 1.929268516 |
| Rab36 | 1.928656494 |
| Pdk4 | 1.928396527 |
| Zscan25 | 1.928157123 |
| Wbp2 | 1.927679261 |
| Gpr160 | 1.927309778 |
| Parp11 | 1.927282673 |
| Gucy1b3 | 1.926964971 |
| Fdxacb1 | 1.926138302 |
| Pno1 | 1.925997555 |
| Rhobtb2 | 1.925656898 |

| | |
|---|---|
| Zranb2 | 1.925635742 |
| Nlrc5 | 1.924992956 |
| Cops4 | 1.924961283 |
| Ptp4a1 | 1.924557174 |
| Pdzd7 | 1.924414421 |
| Lrriq3 | 1.92431652 |
| Prickle2 | 1.924218846 |
| Rilp | 1.924195149 |
| Ube2q2 | 1.923986945 |
| Ano2 | 1.923247582 |
| Mlh3 | 1.922760244 |
| Usp40 | 1.922326129 |
| P2rx7 | 1.922299239 |
| Fan1 | 1.921918253 |
| S100a6 | 1.92152364 |
| Galnt7 | 1.921512504 |
| Gdf7 | 1.920758349 |
| Orc6 | 1.920742641 |
| Azi2 | 1.920689248 |
| Pura | 1.92060479 |
| Mical3 | 1.920596392 |
| Nolc1 | 1.920260487 |
| Msantd2 | 1.919835372 |
| Dstyk | 1.919548316 |
| Lrrc34 | 1.919064958 |
| Ccnb2 | 1.919047888 |
| Tppp3 | 1.919047591 |
| Frat2 | 1.918485702 |
| Ifit3 | 1.918469467 |
| Ubtf | 1.917954984 |
| Arrdc4 | 1.917724666 |
| Sf1 | 1.917371299 |
| Wdfy1 | 1.917265669 |
| Daam1 | 1.917233677 |
| Dus3l | 1.916387875 |
| Slc26a2 | 1.916184236 |
| Copb2 | 1.915989138 |
| Tor1b | 1.915171305 |
| Fam126a | 1.914689081 |
| Pla2g4c | 1.914193829 |
| Phlda3 | 1.913970507 |
| Rap1a | 1.91367168 |
| Rap1b | 1.913555765 |
| Umps | 1.913493885 |
| Alkbh2 | 1.913382847 |
| Zbtb5 | 1.913030126 |
| Mtfr1l | 1.912170034 |

| | |
|---|---|
| Pld6 | 1.912062972 |
| Ppp1r2 | 1.911881513 |
| Tnfrsf8 | 1.911726174 |
| Anks3 | 1.91164177 |
| Rpl38 | 1.911004762 |
| Cbln3 | 1.910983715 |
| Rfx8 | 1.910274829 |
| Mrpl36 | 1.910080021 |
| Pgf | 1.909034764 |
| Mcfd2 | 1.908915809 |
| Inpp5a | 1.908069668 |
| Itga5 | 1.908036627 |
| Atn1 | 1.908013923 |
| Slc30a1 | 1.907425964 |
| Crebl2 | 1.9063428 |
| Snx8 | 1.906083282 |
| Arhgap10 | 1.905217268 |
| Rpl24 | 1.905083354 |
| Ptger4 | 1.905035587 |
| Prmt5 | 1.90494737 |
| Clip1 | 1.904090691 |
| Tfip11 | 1.903243455 |
| Vegfc | 1.902457495 |
| Cyhr1 | 1.901436973 |
| Pex11a | 1.901078422 |
| Rpl11 | 1.900040357 |
| Eif6 | 1.899593056 |
| Mrpl24 | 1.899420358 |
| Atg14 | 1.898941809 |
| Hccs | 1.89878225 |
| Bag2 | 1.898372364 |
| Stk32c | 1.898290412 |
| Fancl | 1.898166847 |
| Oaf | 1.897182774 |
| Pomgnt2 | 1.896951422 |
| Ebpl | 1.896353291 |
| Mrpl32 | 1.896171009 |
| Malt1 | 1.895887421 |
| Tex30 | 1.895828818 |
| Sowahc | 1.895758906 |
| Polr2e | 1.895494075 |
| Sh3glb1 | 1.894248446 |
| Pet117 | 1.893844307 |
| Snx12 | 1.893322217 |
| Fgfbp3 | 1.893018754 |
| Asgr1 | 1.892220391 |
| Ranbp9 | 1.892103605 |

| | |
|---|---|
| Ptpn14 | 1.890903144 |
| Atf7 | 1.890532583 |
| Bicd2 | 1.889366439 |
| Chmp2a | 1.888167402 |
| Aimp2 | 1.887601641 |
| Irf2bp2 | 1.887096116 |
| Trpv1 | 1.887047573 |
| H3f3a | 1.886881794 |
| P2rx4 | 1.886530565 |
| Fgf18 | 1.886459608 |
| Ckap2 | 1.886307151 |
| Wbscr16 | 1.885053844 |
| Cmtm1 | 1.884828643 |
| Fkbp4 | 1.884740304 |
| Wnt4 | 1.88455723 |
| Tfg | 1.884145496 |
| Rhou | 1.88370757 |
| Txnrd3 | 1.883424058 |
| Pdf | 1.883231507 |
| Fndc3b | 1.882291234 |
| Lrrc23 | 1.881574686 |
| Plcd1 | 1.881570432 |
| Qrich1 | 1.881564094 |
| Cpq | 1.881469024 |
| Serinc5 | 1.880590965 |
| Usp7 | 1.880308966 |
| Maged1 | 1.879763459 |
| Prkd1 | 1.879408251 |
| Arnt | 1.879175935 |
| Polr3e | 1.879145637 |
| Dut | 1.878863328 |
| Cd47 | 1.878420134 |
| Lin9 | 1.878277711 |
| Mpzl1 | 1.877460657 |
| Myo7a | 1.877394604 |
| Ptpru | 1.876581405 |
| Axin2 | 1.876572986 |
| Uap1 | 1.87636907 |
| Fermt1 | 1.875966921 |
| Sys1 | 1.875863721 |
| Iqsec1 | 1.875088734 |
| Atg4b | 1.875048147 |
| Prss23 | 1.87482618 |
| Dennd3 | 1.874807757 |
| Smox | 1.8747022 |
| Tmem192 | 1.874268996 |
| Abcb8 | 1.874268937 |

| | |
|---|---|
| Slc9a9 | 1.874103713 |
| Svil | 1.87287588 |
| Palm | 1.872489117 |
| Jak3 | 1.872399371 |
| Casc3 | 1.871661094 |
| Nid1 | 1.871622741 |
| Hist1h2ac | 1.871171068 |
| Psmb1 | 1.870706893 |
| Whamm | 1.869976535 |
| Nr5a2 | 1.869237699 |
| Hecw2 | 1.868700335 |
| Nle1 | 1.868261886 |
| Adat3 | 1.867637812 |
| Sned1 | 1.86741158 |
| Cks2 | 1.86668146 |
| Pmf1 | 1.866549264 |
| Slc44a5 | 1.865411012 |
| Nprl2 | 1.865219343 |
| Gapdh | 1.864208229 |
| S1pr1 | 1.863794713 |
| Haus6 | 1.863644418 |
| Unkl | 1.863372855 |
| Cep170 | 1.863290672 |
| Parg | 1.862963942 |
| Wdr5b | 1.862902686 |
| Myd88 | 1.862824055 |
| Camk2n1 | 1.862694621 |
| Mtg1 | 1.86265534 |
| Mrpl9 | 1.862487991 |
| Tspan8 | 1.862474326 |
| Meis1 | 1.862423793 |
| Srrm2 | 1.862420708 |
| Prdx4 | 1.862018946 |
| Nudt8 | 1.861934594 |
| Cmtr1 | 1.861863502 |
| Cd274 | 1.861195541 |
| Ints8 | 1.861156661 |
| Faim | 1.860842901 |
| Cep164 | 1.860695855 |
| Upf1 | 1.859222277 |
| Nfat5 | 1.859220882 |
| Jade3 | 1.858688155 |
| Hsbp1 | 1.858055731 |
| Ddost | 1.857547285 |
| Slc22a5 | 1.856558808 |
| Vps9d1 | 1.855800156 |
| Phldb1 | 1.855581042 |

| | |
|---|---|
| Bgn | 1.854856733 |
| Luc7l2 | 1.854674173 |
| Ece2 | 1.854358031 |
| Mapk9 | 1.854253325 |
| Gtse1 | 1.854026891 |
| Rnf157 | 1.853814615 |
| Tnfsf18 | 1.853346674 |
| Pde7a | 1.852615727 |
| Aktip | 1.851197289 |
| Slc2a12 | 1.850688334 |
| Uba3 | 1.849345934 |
| Ankhd1 | 1.849030213 |
| Arhgef17 | 1.84861855 |
| Gzf1 | 1.848509787 |
| Atp5g1 | 1.848342342 |
| Ube2d1 | 1.848247755 |
| Cdk14 | 1.84815959 |
| Scg5 | 1.847154917 |
| Trappc9 | 1.847079545 |
| Dcaf6 | 1.846824562 |
| N4bp2 | 1.846542918 |
| Rcc2 | 1.846190977 |
| Nbeal2 | 1.845633437 |
| Sec22c | 1.845441375 |
| Dcaf12 | 1.844954308 |
| Celf2 | 1.843450305 |
| Naa40 | 1.843449979 |
| Fzd6 | 1.843251451 |
| Aatk | 1.84303975 |
| Nkain2 | 1.842908702 |
| Ankrd33b | 1.842888311 |
| Trappc5 | 1.842044962 |
| Ubr7 | 1.841931254 |
| Grb14 | 1.840739083 |
| Efhb | 1.84059453 |
| Chpf | 1.840594461 |
| Commd10 | 1.840416797 |
| Mrpl15 | 1.840334276 |
| Wipf1 | 1.840004584 |
| Clcn3 | 1.839933613 |
| Nrarp | 1.839907169 |
| Ifit1 | 1.839566575 |
| Clic5 | 1.839168993 |
| Mpp6 | 1.836332517 |
| Pabpc4 | 1.835543359 |
| Mapk11 | 1.835526325 |
| Slc25a29 | 1.835390431 |

| | |
|---|---|
| Egfl8 | 1.83526366 |
| Copz1 | 1.834926029 |
| Slc6a8 | 1.834834964 |
| Ptdss1 | 1.834817392 |
| Fjx1 | 1.834683079 |
| Nrxn3 | 1.834418675 |
| Stk24 | 1.83432146 |
| Dnmt1 | 1.833893461 |
| Pms2 | 1.833873216 |
| Pgap2 | 1.833818572 |
| Fam189a2 | 1.833744633 |
| Psmc6 | 1.8337201 |
| Gtpbp8 | 1.833391098 |
| Uck2 | 1.833075224 |
| Clec12b | 1.832829 |
| Eps8l2 | 1.831946831 |
| Acad9 | 1.83155026 |
| Ermp1 | 1.831270042 |
| Fiz1 | 1.82982575 |
| Areg | 1.829463835 |
| Cenpf | 1.828595537 |
| Mrpl40 | 1.828471695 |
| Hmg20b | 1.827180218 |
| Lmo2 | 1.826552239 |
| Lcn6 | 1.826359626 |
| Scai | 1.826013169 |
| Ttc7b | 1.825995795 |
| Bbs9 | 1.825965495 |
| Wdtc1 | 1.825940577 |
| Snx29 | 1.824826505 |
| Map3k12 | 1.824644532 |
| Tfrc | 1.824142735 |
| Map7d2 | 1.823561295 |
| Trak2 | 1.823398353 |
| Nr2f2 | 1.82185688 |
| Fam162a | 1.82182915 |
| Abracl | 1.821776173 |
| Trim32 | 1.821631418 |
| Endod1 | 1.821519411 |
| Uhmk1 | 1.82137297 |
| Gtpbp10 | 1.821182009 |
| Tmem117 | 1.821115031 |
| Abce1 | 1.820371656 |
| Letm1 | 1.820142832 |
| Arsa | 1.819806153 |
| Rgl1 | 1.819658915 |
| Arhgap32 | 1.818476831 |

| | |
|---|---|
| Cd55 | 1.817977244 |
| Cul5 | 1.817731517 |
| Pmpca | 1.817125142 |
| Mrpl33 | 1.815857902 |
| Vamp3 | 1.815648017 |
| Ppia | 1.815407075 |
| Map3k13 | 1.815199946 |
| Lrrc42 | 1.81511719 |
| Luzp2 | 1.814844701 |
| Uhrf1bp1 | 1.81450595 |
| Trmt5 | 1.814246681 |
| Pcdh18 | 1.814005806 |
| Me2 | 1.813482702 |
| Las1l | 1.813101941 |
| Chd3 | 1.812749903 |
| Irf6 | 1.812395116 |
| Kcnma1 | 1.811933916 |
| Tmbim4 | 1.81181224 |
| Slc30a9 | 1.811599196 |
| Pdia4 | 1.811057912 |
| Itgav | 1.810874343 |
| Amigo2 | 1.810546195 |
| Man2c1 | 1.810503242 |
| Ehhadh | 1.809908995 |
| Fis1 | 1.809481937 |
| Cdc123 | 1.809423169 |
| Kansl2 | 1.809086956 |
| Klc2 | 1.809004475 |
| Psmd7 | 1.808625499 |
| Cnn2 | 1.808610934 |
| Alg2 | 1.808563724 |
| Ube2o | 1.808365804 |
| Ppp3cc | 1.80831045 |
| Kbtbd7 | 1.804429368 |
| Smarca4 | 1.803922461 |
| Ppp2r3a | 1.803807066 |
| Wiz | 1.803571221 |
| Cdc42ep1 | 1.803452729 |
| Tubb4b | 1.802993385 |
| Cnppd1 | 1.802743488 |
| Ift57 | 1.802628088 |
| Rangap1 | 1.80246623 |
| Axl | 1.801564974 |
| Slc23a2 | 1.801507853 |
| Tmem86a | 1.801027703 |
| Fam76a | 1.80098793 |
| Caap1 | 1.800689455 |

| | |
|---|---|
| Parp4 | 1.80045221 |
| S100a11 | 1.800370151 |
| Smtnl2 | 1.800047002 |
| Ptgds | 1.798964466 |
| Myzap | 1.798048407 |
| Igf1r | 1.797312858 |
| Ube2l6 | 1.797081031 |
| Id3 | 1.796654862 |
| Pabpc1 | 1.796510553 |
| Nagpa | 1.796201469 |
| Ctgf | 1.796032964 |
| Nanp | 1.795248465 |
| Slc35b1 | 1.793971282 |
| Tmem106a | 1.793445639 |
| Rhbdf2 | 1.793238282 |
| Patl1 | 1.792736225 |
| Syvn1 | 1.792723646 |
| Eid1 | 1.792655661 |
| Tbxas1 | 1.792559241 |
| Cd109 | 1.792331841 |
| Tmcc3 | 1.79191903 |
| Mgat1 | 1.791580979 |
| Mobp | 1.791236948 |
| Socs2 | 1.790841567 |
| Ergic1 | 1.790625084 |
| Eps8l1 | 1.789942376 |
| Nr4a2 | 1.789655174 |
| Cep41 | 1.789618305 |
| Baz1a | 1.789396019 |
| Sh3pxd2a | 1.789222911 |
| Aldh1a3 | 1.789189301 |
| Ldb2 | 1.789158145 |
| Adamts7 | 1.789119601 |
| Tyms | 1.788409732 |
| Fbln2 | 1.787972777 |
| Spcs1 | 1.787713998 |
| Pxn | 1.78732475 |
| Nrip2 | 1.787155731 |
| Il11 | 1.78686274 |
| Mrps26 | 1.786653554 |
| Acy1 | 1.786485887 |
| Itga4 | 1.786196627 |
| Clec10a | 1.785439973 |
| Cyp2s1 | 1.784584605 |
| Tnfrsf1a | 1.784404422 |
| Uty | 1.783895884 |
| Lrrc43 | 1.783834237 |

| | |
|---|---|
| Il27ra | 1.783734684 |
| Fez2 | 1.783628609 |
| Cdkal1 | 1.783356135 |
| Gjc2 | 1.783345402 |
| Gabarapl2 | 1.782428987 |
| Tmem259 | 1.782285505 |
| Rabgap1l | 1.782097041 |
| Lias | 1.781857476 |
| Papln | 1.781412546 |
| Atg13 | 1.781131685 |
| Psmc3 | 1.780681311 |
| Utp15 | 1.780551208 |
| Calr | 1.780497393 |
| Ttc39b | 1.780433321 |
| Zfyve19 | 1.780210045 |
| Fam3c | 1.779516302 |
| Arhgap12 | 1.779157977 |
| Ykt6 | 1.778537163 |
| Tfcp2 | 1.778363444 |
| Extl3 | 1.777807833 |
| Meox2 | 1.777786075 |
| Psmd6 | 1.777738413 |
| Tcf20 | 1.777615653 |
| Chrna5 | 1.777439785 |
| Plaur | 1.777088099 |
| Lrrn1 | 1.776933311 |
| Polr2d | 1.77686703 |
| Gnb1l | 1.77649066 |
| Itgae | 1.776060467 |
| Mprip | 1.775918902 |
| Fancd2 | 1.775917538 |
| Amigo3 | 1.775751148 |
| Mapre2 | 1.775437464 |
| Ggt7 | 1.775178607 |
| Mmp24 | 1.775142563 |
| Ccdc66 | 1.774629609 |
| Llgl1 | 1.774449559 |
| Tmbim6 | 1.774292865 |
| Ino80b | 1.773842687 |
| Foxf1 | 1.773841186 |
| Sez6l2 | 1.773815979 |
| Slc35d1 | 1.773433927 |
| Foxred1 | 1.772696737 |
| Brd9 | 1.772118412 |
| Pdrg1 | 1.771893117 |
| Unc13d | 1.771840502 |
| Fbln1 | 1.771423946 |

| | |
|---|---|
| Cyyr1 | 1.771072414 |
| Hnrnph2 | 1.77012539 |
| Brms1l | 1.769938064 |
| Otud6b | 1.769687308 |
| Alms1 | 1.769599793 |
| Cep250 | 1.769246088 |
| Furin | 1.769049269 |
| Tssk6 | 1.768853437 |
| Isyna1 | 1.768662721 |
| Triobp | 1.767905906 |
| Atp6v1c1 | 1.767551206 |
| Actr8 | 1.767355077 |
| Pbx3 | 1.766668446 |
| Eml5 | 1.76598799 |
| Cwc25 | 1.764729739 |
| Hmgcr | 1.764579003 |
| Gramd1b | 1.763323195 |
| Has2 | 1.762091435 |
| Sh3rf2 | 1.761817646 |
| Abcb10 | 1.761566046 |
| Taf4b | 1.761051006 |
| Col4a3bp | 1.759929403 |
| Pigx | 1.758786472 |
| Sh2d5 | 1.758607384 |
| Osgep | 1.758482799 |
| Ninj2 | 1.758147361 |
| Nr1h2 | 1.757576836 |
| Fzd4 | 1.757372713 |
| Rtcb | 1.7571655 |
| Itfg2 | 1.757042646 |
| Srm | 1.75697642 |
| Bves | 1.756548934 |
| Kmt2d | 1.756010621 |
| Eln | 1.755431566 |
| Srsf6 | 1.754951256 |
| Tubb6 | 1.754499352 |
| Aldh7a1 | 1.753986886 |
| Pde4a | 1.753642039 |
| Syf2 | 1.753540919 |
| Srp19 | 1.752502188 |
| Arhgef39 | 1.752480759 |
| Ttc23 | 1.752341664 |
| Ptar1 | 1.752299347 |
| Creb5 | 1.752010779 |
| Mylk | 1.751819417 |
| Klhl12 | 1.751705373 |
| Angpt2 | 1.750169959 |

| | |
|---|---|
| Scrn3 | 1.750126057 |
| Trim25 | 1.749594516 |
| Tmem42 | 1.748929561 |
| Zzef1 | 1.748664719 |
| Elmod3 | 1.748448346 |
| Lonrf1 | 1.748180045 |
| Ndor1 | 1.747805233 |
| Gnpda1 | 1.747797354 |
| Eif4g3 | 1.747751609 |
| Mettl15 | 1.747580799 |
| Fibcd1 | 1.747369028 |
| Cmtm6 | 1.747091109 |
| Scand1 | 1.747046117 |
| Apln | 1.746818789 |
| Pi4k2b | 1.74630299 |
| Minos1 | 1.746282791 |
| Chn2 | 1.745990416 |
| Itfg1 | 1.745658733 |
| Slc39a3 | 1.745438996 |
| 42623 | 1.74433429 |
| Ptcd3 | 1.744216422 |
| Rag1 | 1.744116172 |
| Asah2 | 1.74351435 |
| Fbxo8 | 1.74342688 |
| Ppm1n | 1.743284851 |
| Pfdn5 | 1.743201382 |
| Pigt | 1.743103266 |
| Mut | 1.743003858 |
| Elf1 | 1.742605215 |
| Shroom2 | 1.742541383 |
| Ccbl2 | 1.741909301 |
| Mlkl | 1.741733216 |
| Anks6 | 1.741502788 |
| Rundc1 | 1.741133932 |
| Gle1 | 1.740847211 |
| Btg2 | 1.740184604 |
| Fbf1 | 1.739340404 |
| Btbd7 | 1.739333481 |
| Vps28 | 1.738942507 |
| Nicn1 | 1.738480529 |
| Mcl1 | 1.738401436 |
| Zbtb18 | 1.738375742 |
| Pik3cg | 1.737996488 |
| Gadd45a | 1.737932561 |
| Psen2 | 1.737373224 |
| Hoxa9 | 1.73700067 |
| Ccdc124 | 1.735725883 |

| | |
|---|---|
| Mov10l1 | 1.73540538 |
| Tmem33 | 1.735125804 |
| Cep104 | 1.733861025 |
| Poln | 1.733689347 |
| Fam173b | 1.733609705 |
| Pmm1 | 1.733585026 |
| Glis3 | 1.733529686 |
| Rac3 | 1.73340111 |
| Oxsm | 1.733393501 |
| Rnd3 | 1.733314908 |
| Rab10 | 1.73325481 |
| Nrd1 | 1.733188545 |
| Uhrf1 | 1.731919569 |
| Tsnax | 1.731734143 |
| Twist2 | 1.73142598 |
| Tarsl2 | 1.731254122 |
| Tspan13 | 1.731175871 |
| Hmox1 | 1.730939378 |
| Cby1 | 1.730911517 |
| Atg10 | 1.730702135 |
| Dhx40 | 1.730159297 |
| Fastkd5 | 1.72972734 |
| Thbs1 | 1.729678705 |
| Cntln | 1.728872374 |
| Ttc13 | 1.728416965 |
| Sirt3 | 1.727852266 |
| Mbnl1 | 1.727851701 |
| Camkk2 | 1.727298283 |
| Rbfox2 | 1.727049633 |
| Hnrnpdl | 1.726839515 |
| Fbxo9 | 1.726367364 |
| Scn8a | 1.726170019 |
| Col12a1 | 1.725201497 |
| Heca | 1.724892973 |
| Mcm3ap | 1.724646173 |
| Sp3 | 1.723770818 |
| Megf6 | 1.723669599 |
| Sirt1 | 1.723472135 |
| Ppil4 | 1.723372776 |
| Nckap1 | 1.722583879 |
| Runx1t1 | 1.72203538 |
| Fbxo4 | 1.721796191 |
| Recql4 | 1.721454232 |
| Catsper2 | 1.721034705 |
| Chtf18 | 1.72091473 |
| Pold4 | 1.720633188 |
| Clstn2 | 1.720194198 |

| | |
|---|---|
| Smyd2 | 1.719411468 |
| Nhej1 | 1.718909404 |
| Abcd1 | 1.718510504 |
| Ralgapa1 | 1.718140094 |
| Hilpda | 1.718002781 |
| Cep85 | 1.717960604 |
| Tinagl1 | 1.717489782 |
| Klhl26 | 1.717017879 |
| Fsip1 | 1.716996072 |
| Timm50 | 1.716969495 |
| Panx2 | 1.716862786 |
| Spry1 | 1.716647862 |
| Sec13 | 1.716512165 |
| Mien1 | 1.716365372 |
| Stk11 | 1.716002697 |
| Lpar1 | 1.716001315 |
| Col8a1 | 1.715655087 |
| Wnk1 | 1.715569274 |
| Naa30 | 1.715250285 |
| Rrp1 | 1.715142482 |
| Bsg | 1.714668006 |
| Rbm26 | 1.714224027 |
| Chrna3 | 1.7138114 |
| Csf3 | 1.713452722 |
| Zbtb38 | 1.713362486 |
| Mrps16 | 1.712685725 |
| Nhlrc2 | 1.712323729 |
| Atp13a1 | 1.712137095 |
| Pde6g | 1.712113899 |
| Coch | 1.712092279 |
| Wsb2 | 1.711991627 |
| Zcwpw1 | 1.711985699 |
| Clcf1 | 1.711954414 |
| Ppp1r9b | 1.710987376 |
| Xrcc6bp1 | 1.710202449 |
| Nbeal1 | 1.710157521 |
| Slc43a3 | 1.710032735 |
| Fcrlb | 1.709791695 |
| Cspp1 | 1.709535588 |
| Gpr137b | 1.709299731 |
| Eif3f | 1.708548415 |
| Ddit3 | 1.708391001 |
| Arhgdia | 1.707805042 |
| Kcnip4 | 1.707794815 |
| Zc3h3 | 1.707191926 |
| Srsf10 | 1.706757544 |
| Ndufb6 | 1.706554096 |

| | |
|---|---|
| Syn2 | 1.70600613 |
| Phkg2 | 1.705545292 |
| Akap10 | 1.705374916 |
| Sgpp1 | 1.705374139 |
| Gtf3c6 | 1.705368201 |
| Jam3 | 1.705019105 |
| Rbm34 | 1.704555036 |
| Ccdc57 | 1.704542518 |
| Rtn2 | 1.704380851 |
| Cdpf1 | 1.704122783 |
| Slc25a15 | 1.703990963 |
| Npdc1 | 1.703792435 |
| Maml2 | 1.703295352 |
| Trappc4 | 1.703287132 |
| Mapk8ip1 | 1.703205791 |
| Stat5a | 1.70259409 |
| Tmem107 | 1.702481548 |
| Tmco3 | 1.70223341 |
| Ppp2r2d | 1.702153906 |
| Oaz1 | 1.702129047 |
| Zdhhc7 | 1.701271802 |
| Nfia | 1.701239111 |
| Slc26a6 | 1.701126238 |
| Ccdc157 | 1.701052418 |
| Ralgps1 | 1.700992497 |
| Ccdc14 | 1.700938105 |
| Rasip1 | 1.700524937 |
| Dym | 1.69975426 |
| Eif4b | 1.699547715 |
| Ankrd13c | 1.699546572 |
| Maea | 1.699322091 |
| Tbc1d12 | 1.699192756 |
| Mapkbp1 | 1.698854402 |
| Shpk | 1.698653074 |
| Baz2a | 1.698072414 |
| Mgat5 | 1.697629034 |
| Tmem167b | 1.697435769 |
| Tyw1 | 1.697265459 |
| Anapc11 | 1.697103228 |
| Ccni | 1.697007308 |
| Rpusd2 | 1.696839937 |
| Klhdc1 | 1.696675353 |
| Mad2l1 | 1.69654952 |
| Vapb | 1.695357242 |
| Tdp2 | 1.695052576 |
| Pcolce | 1.694417063 |
| Rad51ap1 | 1.693662621 |

| | |
|---|---|
| Pi4ka | 1.693251131 |
| E2f7 | 1.692721773 |
| Cep135 | 1.692684295 |
| Tmem132a | 1.692594513 |
| Nup188 | 1.692501248 |
| Usp28 | 1.692368125 |
| Col17a1 | 1.692234973 |
| Stau2 | 1.692229048 |
| Tmem199 | 1.692172071 |
| H3f3b | 1.691970995 |
| Reln | 1.691388956 |
| Dpp8 | 1.69038221 |
| Fam174a | 1.690369198 |
| Actr10 | 1.689747031 |
| Mrps34 | 1.68953667 |
| Mnt | 1.688970536 |
| Tmem57 | 1.688145041 |
| Pwwp2b | 1.688023513 |
| Ccdc28a | 1.68787523 |
| Fgf2 | 1.68609221 |
| Cox4i1 | 1.685995278 |
| Mtrf1 | 1.685961941 |
| Tubgcp2 | 1.685447611 |
| Polr3a | 1.684730173 |
| Ch25h | 1.684265408 |
| Atox1 | 1.684182334 |
| Dapk2 | 1.683786341 |
| Xylt2 | 1.683351594 |
| Kif16b | 1.683159199 |
| Sdhaf2 | 1.682153764 |
| Sik1 | 1.681619838 |
| Ankrd26 | 1.680803894 |
| Fkbp8 | 1.680563091 |
| Tmsb10 | 1.680447354 |
| Htr2b | 1.680330459 |
| Nktr | 1.680223315 |
| Alg6 | 1.679920089 |
| Setx | 1.679679684 |
| Fbxl4 | 1.679339384 |
| Mex3d | 1.678408377 |
| Cysltr2 | 1.677458749 |
| Sdha | 1.67722421 |
| Ptges3 | 1.676948497 |
| Huwe1 | 1.676755109 |
| Taf1b | 1.676596074 |
| Slc25a17 | 1.676558042 |
| Atoh8 | 1.676343552 |

| | |
|---|---|
| Gtf3c2 | 1.676307122 |
| Ptp4a3 | 1.676104246 |
| Irak3 | 1.675992923 |
| Scube2 | 1.675976323 |
| Mtx3 | 1.675555091 |
| Pqlc3 | 1.675180296 |
| Naa20 | 1.674975658 |
| Nelfb | 1.674804664 |
| Elp5 | 1.674802561 |
| Dcaf17 | 1.674613957 |
| Arsk | 1.674355828 |
| Krt10 | 1.67428034 |
| Nxpe3 | 1.674005216 |
| Samd1 | 1.672929066 |
| Pdcd7 | 1.672229277 |
| Ahr | 1.671666731 |
| Slc25a1 | 1.671201935 |
| Sp1 | 1.670215503 |
| Gltp | 1.669777528 |
| Htra3 | 1.669511285 |
| Sec31b | 1.669083845 |
| Rgl2 | 1.667903302 |
| Isy1 | 1.66764535 |
| Ubtd1 | 1.666782147 |
| Arl3 | 1.666663929 |
| Mrps10 | 1.666538121 |
| Zfand2b | 1.666525528 |
| Smarcd3 | 1.665740053 |
| Wdr48 | 1.665729636 |
| Cops7b | 1.66554236 |
| Lsm10 | 1.664844086 |
| Pdcd2 | 1.66448598 |
| Flnb | 1.664361571 |
| Slc5a10 | 1.664320556 |
| Fam73a | 1.664005682 |
| Mdm1 | 1.663441739 |
| Gpr107 | 1.66340296 |
| Mblac2 | 1.662986819 |
| Cryzl1 | 1.662748687 |
| Cbln2 | 1.662720526 |
| Eid2b | 1.660658666 |
| Gdap2 | 1.660240525 |
| Cryz | 1.659377373 |
| Tbcb | 1.659227229 |
| Stxbp5 | 1.659025692 |
| Itgb4 | 1.658904003 |
| Fkbp1b | 1.658854433 |

| | |
|---|---|
| Hps1 | 1.658552385 |
| Gba | 1.658181687 |
| Gja5 | 1.657607635 |
| Rcbtb2 | 1.65721439 |
| Ranbp6 | 1.656815753 |
| Tomm22 | 1.656502762 |
| Rfc1 | 1.656466441 |
| Yaf2 | 1.656185937 |
| Eif4e3 | 1.654730334 |
| Klf6 | 1.654414367 |
| Eci2 | 1.653327866 |
| Pde2a | 1.653020926 |
| Dhx15 | 1.653003801 |
| Rnf111 | 1.652986531 |
| Manf | 1.652307733 |
| Trim35 | 1.652294643 |
| R3hcc1l | 1.651816903 |
| Hyal3 | 1.651651893 |
| Frat1 | 1.651293306 |
| Edem2 | 1.651107056 |
| Tnfaip2 | 1.649675824 |
| Glis2 | 1.64952143 |
| Smyd3 | 1.6492836 |
| Bricd5 | 1.648399621 |
| Tmem134 | 1.648396938 |
| Tmem30a | 1.648279288 |
| Pop4 | 1.647182707 |
| Fbxo30 | 1.647145571 |
| Taok2 | 1.646182768 |
| Tmem201 | 1.64587919 |
| Klk6 | 1.645088634 |
| Xrn1 | 1.644802136 |
| Arfrp1 | 1.64436155 |
| Iqcc | 1.643474945 |
| St7l | 1.642856362 |
| Pqbp1 | 1.642627295 |
| Ndufb5 | 1.642510587 |
| Gtf2h2 | 1.642426004 |
| Rad52 | 1.642199887 |
| Serpine2 | 1.641945324 |
| Pgls | 1.641703089 |
| Gskip | 1.641434546 |
| Slc45a1 | 1.641385076 |
| Tnk2 | 1.641173689 |
| Tuba1c | 1.640969395 |
| Lama4 | 1.64052996 |
| Il6st | 1.640287083 |

| | |
|---|---|
| Rbmx | 1.638748188 |
| Abhd5 | 1.637976647 |
| R3hdm4 | 1.636963066 |
| Adck3 | 1.636378227 |
| Slc38a9 | 1.636322388 |
| Dab2ip | 1.635610201 |
| Arfgap2 | 1.635482801 |
| Selplg | 1.635457675 |
| Tldc1 | 1.635347269 |
| Plaa | 1.63523001 |
| Dpyd | 1.634629923 |
| Bud13 | 1.634086703 |
| Nupl2 | 1.633787156 |
| Cep120 | 1.633109551 |
| Eml2 | 1.633089927 |
| Cyba | 1.632922757 |
| Slc4a7 | 1.632912568 |
| Fdx1 | 1.632378666 |
| Tspan11 | 1.631658125 |
| Dnase1l3 | 1.631557249 |
| Etv5 | 1.631378475 |
| Crlf1 | 1.630931004 |
| Nhlrc1 | 1.630307411 |
| Elp4 | 1.629883117 |
| Fbxl13 | 1.629565053 |
| Grhpr | 1.62941392 |
| D2hgdh | 1.628897196 |
| Abcg2 | 1.628824723 |
| Ecscr | 1.62787144 |
| Jagn1 | 1.627553961 |
| Dpcd | 1.626267011 |
| Fhit | 1.625704068 |
| Rab5c | 1.625302445 |
| Gnl1 | 1.625279913 |
| Tnfsf10 | 1.62437325 |
| Slc25a37 | 1.624284983 |
| Fbxl18 | 1.623540492 |
| Eml6 | 1.622920311 |
| Clip2 | 1.622536712 |
| Dpp3 | 1.622250114 |
| Ntrk2 | 1.622202677 |
| Papss1 | 1.62136777 |
| Cfdp1 | 1.621138172 |
| Ikbkg | 1.621009833 |
| Ttc1 | 1.620849287 |
| Scaf11 | 1.620812847 |
| Sclt1 | 1.62071592 |

| | |
|---|---|
| Usb1 | 1.620159344 |
| Cdkn1b | 1.619829213 |
| Clk2 | 1.618920547 |
| Mpzl3 | 1.618777527 |
| Pdzrn3 | 1.618565592 |
| Gcnt1 | 1.61847292 |
| Lrrc45 | 1.618159174 |
| Gtf2a1 | 1.617283697 |
| Cdkn2aip | 1.617170973 |
| Naglu | 1.617101375 |
| Mbnl2 | 1.616272322 |
| Nelfe | 1.615990466 |
| Gabarap | 1.615896307 |
| Sf3b3 | 1.615598962 |
| Tepp | 1.614946064 |
| Hp1bp3 | 1.614155204 |
| Slc30a3 | 1.613983598 |
| Ezr | 1.61340553 |
| Ccdc86 | 1.61312876 |
| Fry | 1.612815847 |
| Sox4 | 1.612761827 |
| Enc1 | 1.612515338 |
| Oxld1 | 1.61219245 |
| Rgs2 | 1.611292638 |
| Adm2 | 1.610499471 |
| Zmat3 | 1.610456024 |
| Unc93b1 | 1.610441018 |
| Dph7 | 1.610208392 |
| Vps13b | 1.609666163 |
| Mpdz | 1.609478742 |
| Elovl5 | 1.609286707 |
| Eif3i | 1.608326063 |
| Oma1 | 1.607912388 |
| Vasn | 1.606316949 |
| Tmem38a | 1.605766688 |
| Psme2 | 1.605507587 |
| Tmem255b | 1.604098338 |
| Cox16 | 1.60402297 |
| Nup93 | 1.603692798 |
| Lmtk2 | 1.603619802 |
| Aldh3b1 | 1.603271024 |
| Gen1 | 1.603263158 |
| Gcat | 1.603211973 |
| Slc2a9 | 1.602137962 |
| Flywch2 | 1.60201016 |
| Sfmbt1 | 1.601888769 |
| Ccdc59 | 1.601253083 |

| | |
|---|---|
| Dhps | 1.601119088 |
| Zbtb17 | 1.601034289 |
| Plekha7 | 1.600481395 |
| Hddc3 | 1.600480774 |
| Ndufb8 | 1.600386029 |
| Fam135a | 1.60032936 |
| Parp12 | 1.600085513 |
| Rnf115 | 1.599769474 |
| Cnn3 | 1.599743619 |
| Abhd17c | 1.599723352 |
| Chd9 | 1.599606858 |
| Pop7 | 1.598803524 |
| Atat1 | 1.598444526 |
| Rab13 | 1.598320083 |
| Rpl7l1 | 1.598219164 |
| Usp45 | 1.59810223 |
| Wbp1 | 1.597651822 |
| Ppp2ca | 1.597190547 |
| Nrip1 | 1.597180281 |
| Scube3 | 1.597027598 |
| Frmd4b | 1.597008545 |
| Mllt11 | 1.596723839 |
| Nfe2l2 | 1.596473464 |
| Usp34 | 1.596287432 |
| Kat2b | 1.595500914 |
| Ankrd49 | 1.595139777 |
| Fkbp14 | 1.594686723 |
| Polr2c | 1.594670711 |
| Cpsf3l | 1.594614133 |
| E2f2 | 1.594544792 |
| Ndufb4 | 1.594532692 |
| Kmt2b | 1.594015542 |
| Galnt10 | 1.593569239 |
| Bmx | 1.593495558 |
| Etfdh | 1.593412184 |
| Slco2a1 | 1.593344076 |
| Gprasp1 | 1.593229609 |
| Clasrp | 1.592993367 |
| Nfic | 1.592966188 |
| Zbtb10 | 1.59260562 |
| Col24a1 | 1.592474007 |
| Ndufa11 | 1.591232768 |
| Pla2r1 | 1.591097529 |
| Strn3 | 1.590956886 |
| Zswim3 | 1.58990271 |
| Pced1a | 1.589879441 |
| Fam19a2 | 1.58959864 |

| | |
|---|---|
| Nr2c2ap | 1.589527866 |
| Prpf4b | 1.588828132 |
| Ythdc1 | 1.588755476 |
| Bloc1s5 | 1.588658581 |
| Gstm4 | 1.588637698 |
| Dnajc21 | 1.588382922 |
| Afmid | 1.588159429 |
| Nat1 | 1.58791891 |
| Odf3b | 1.587525561 |
| Elf2 | 1.587006141 |
| Klf11 | 1.586586752 |
| Atp1b1 | 1.586554795 |
| Dhrs9 | 1.585911156 |
| Slu7 | 1.585426733 |
| Vps11 | 1.585212757 |
| Plbd2 | 1.585145169 |
| Ppp1r3c | 1.585112008 |
| Gfer | 1.58500597 |
| Rcan3 | 1.584963668 |
| Atad3a | 1.584947954 |
| Senp7 | 1.584493242 |
| Gadl1 | 1.583681445 |
| Tmem206 | 1.583635639 |
| Gas8 | 1.583255592 |
| Grk5 | 1.583212388 |
| St6galnac4 | 1.583173328 |
| Rc3h1 | 1.582861449 |
| Tbl1x | 1.58281276 |
| Adcy1 | 1.582557226 |
| Utp20 | 1.582486312 |
| Mov10 | 1.582469551 |
| Tdrd7 | 1.582081827 |
| Tmem165 | 1.581903299 |
| Impa2 | 1.58098978 |
| Slc12a8 | 1.580918582 |
| Sin3b | 1.580602495 |
| Mnat1 | 1.580533808 |
| Kiss1 | 1.580480257 |
| Asxl1 | 1.57937779 |
| Ap1m1 | 1.579154111 |
| Tmem62 | 1.578930943 |
| Mllt6 | 1.578897004 |
| Mdm4 | 1.578531746 |
| Ssb | 1.578006121 |
| Pptc7 | 1.57784691 |
| Deaf1 | 1.577419383 |
| Nfix | 1.577363815 |

| | |
|---|---|
| Mnd1 | 1.577128987 |
| Stx1a | 1.576952487 |
| Wdr45b | 1.576833296 |
| Vapa | 1.576504347 |
| Neil3 | 1.57605241 |
| Nek10 | 1.576007189 |
| Ccdc85b | 1.575854421 |
| Cd101 | 1.575423427 |
| Dync1i1 | 1.574234822 |
| Dnmbp | 1.573970325 |
| Dnajb12 | 1.573870361 |
| Bnip2 | 1.573847693 |
| Cirbp | 1.572183973 |
| Gnb4 | 1.572121299 |
| Paqr3 | 1.571879613 |
| Nedd4l | 1.571545166 |
| Timm17a | 1.571080476 |
| Map3k7cl | 1.570881201 |
| Manbal | 1.570429709 |
| Rnps1 | 1.569994842 |
| Vps41 | 1.56995896 |
| Armc7 | 1.569875599 |
| Bsdc1 | 1.569713972 |
| Tbc1d10a | 1.568725167 |
| Suclg1 | 1.568571399 |
| Fnta | 1.567899613 |
| Nop9 | 1.567834769 |
| Fkrp | 1.567715772 |
| Cnot7 | 1.567298471 |
| Col5a1 | 1.567228208 |
| Dnajc9 | 1.566685181 |
| Eif1b | 1.566558096 |
| Eif1ax | 1.566431239 |
| Ccl28 | 1.566326698 |
| Gbp2 | 1.566263304 |
| Pdlim4 | 1.565692193 |
| Twf1 | 1.565647697 |
| Noa1 | 1.565069928 |
| Mcur1 | 1.565001297 |
| Vsig2 | 1.564762038 |
| Acsl3 | 1.564684151 |
| Rnf139 | 1.564643338 |
| Shfm1 | 1.56438612 |
| Samhd1 | 1.564153185 |
| Smpd1 | 1.563822894 |
| Serinc2 | 1.563647846 |
| Cdk12 | 1.563591548 |

| | |
|---|---|
| Zcchc24 | 1.563442245 |
| Wdr45 | 1.56340618 |
| Sptan1 | 1.563289077 |
| Aftph | 1.563208481 |
| Tmtc1 | 1.562705048 |
| Nhlrc3 | 1.561216816 |
| Trim24 | 1.560997146 |
| Dot1l | 1.560390154 |
| Aip | 1.560341355 |
| Phtf1 | 1.559798879 |
| Zmiz2 | 1.559589313 |
| Rab3c | 1.559572267 |
| Iqsec2 | 1.559452188 |
| Tbl3 | 1.558915898 |
| Ksr1 | 1.557726696 |
| Bcl7b | 1.557688733 |
| Nek11 | 1.557226421 |
| Rhoc | 1.557095819 |
| Abcc5 | 1.556996957 |
| Pard6a | 1.556821011 |
| Apeh | 1.556275606 |
| Psd | 1.55623802 |
| Eef2k | 1.556132514 |
| Rps12 | 1.555810097 |
| Pecam1 | 1.555808238 |
| Farsb | 1.555521927 |
| Gmip | 1.555051065 |
| Pa2g4 | 1.554924334 |
| Zbtb41 | 1.554727313 |
| Eva1b | 1.554470537 |
| Prr13 | 1.554296445 |
| Tmem51 | 1.553890852 |
| Ttc12 | 1.553204193 |
| Akap11 | 1.552949245 |
| Fhl2 | 1.552735942 |
| Tagln | 1.552366673 |
| Mal2 | 1.552000098 |
| Ift80 | 1.551940252 |
| Mbp | 1.551823064 |
| Tspan10 | 1.551703077 |
| Usp6nl | 1.550933003 |
| Bhmt2 | 1.550027425 |
| Arpc3 | 1.549933615 |
| Tbxa2r | 1.549857259 |
| Rxrb | 1.549758462 |
| A1bg | 1.549645139 |
| Ube2i | 1.549524468 |

| | |
|---|---|
| Tbc1d20 | 1.549313611 |
| Gprin1 | 1.549080532 |
| Mon1a | 1.547176933 |
| Ppfia3 | 1.547106107 |
| Ccnd1 | 1.546963607 |
| Ikzf4 | 1.546431954 |
| Phyhd1 | 1.546278802 |
| Bsn | 1.545747244 |
| Ptpn11 | 1.545310398 |
| Fut11 | 1.545182957 |
| Tvp23b | 1.545027924 |
| Klhl4 | 1.544814721 |
| Mdga1 | 1.544653734 |
| Gpx4 | 1.544536988 |
| Bloc1s6 | 1.544474893 |
| Wasf1 | 1.544156424 |
| Chtop | 1.54390696 |
| Rest | 1.543395122 |
| Cactin | 1.543359981 |
| Pde6d | 1.542431432 |
| Zfyve27 | 1.542298975 |
| Cacul1 | 1.542010812 |
| Rpl39 | 1.541270713 |
| Ybx3 | 1.540668627 |
| Pnpla7 | 1.539445179 |
| Rcn1 | 1.539062496 |
| Zhx3 | 1.538949326 |
| Paxbp1 | 1.538718294 |
| Bcl11a | 1.538596605 |
| Qrich2 | 1.538497744 |
| Ubac2 | 1.538159675 |
| Dfna5 | 1.536801214 |
| Gmpr2 | 1.536691754 |
| Dcaf4 | 1.536264385 |
| Pnrc2 | 1.535863145 |
| Prepl | 1.535832338 |
| Hsd17b10 | 1.535546695 |
| Insig1 | 1.535491478 |
| Baiap3 | 1.534397824 |
| Gid8 | 1.533459199 |
| Disp1 | 1.533395731 |
| Shroom1 | 1.53335755 |
| Skiv2l2 | 1.533356861 |
| Rpn2 | 1.533292372 |
| Tmx3 | 1.533232028 |
| Lyve1 | 1.533100295 |
| Pcyt1b | 1.532242318 |

| | |
|---|---|
| Ercc6 | 1.531771698 |
| Cxcl12 | 1.531602462 |
| Recql5 | 1.531256851 |
| Kcnn3 | 1.531008868 |
| Nae1 | 1.530698361 |
| Smad3 | 1.530583878 |
| Mtl5 | 1.530494367 |
| Klf15 | 1.53019559 |
| Ndst1 | 1.529751754 |
| Blmh | 1.528596182 |
| Slc7a5 | 1.528439533 |
| Cabin1 | 1.528377386 |
| Rbm48 | 1.528233592 |
| Skida1 | 1.528033435 |
| Akap8 | 1.528009861 |
| Ppp1r18 | 1.527556879 |
| Stat5b | 1.527396787 |
| Cotl1 | 1.527386053 |
| Ssh2 | 1.526865203 |
| Mrpl34 | 1.526691982 |
| Nov | 1.526514371 |
| Rabggtb | 1.526344947 |
| Gcdh | 1.526100693 |
| Zdhhc15 | 1.526058695 |
| Ruvbl1 | 1.526049064 |
| Slc9b2 | 1.525659806 |
| Nrk | 1.524585575 |
| Dnah8 | 1.524266265 |
| Uba52 | 1.524135976 |
| Pknox1 | 1.523810963 |
| Glb1l | 1.523410554 |
| Mdk | 1.522945936 |
| Bin3 | 1.522740028 |
| Rec8 | 1.522302613 |
| Clta | 1.521699826 |
| Rps17 | 1.521426305 |
| Celsr2 | 1.521378869 |
| Akt3 | 1.521243781 |
| Rnd2 | 1.520638593 |
| Fam229a | 1.520355341 |
| Myo1c | 1.519735945 |
| Ifitm2 | 1.519314256 |
| Szt2 | 1.519250237 |
| Rufy3 | 1.51895875 |
| Fam126b | 1.518738506 |
| Ckap2l | 1.51871387 |
| Slc45a4 | 1.518573231 |

| | |
|---|---|
| Dcaf15 | 1.518557916 |
| Chrna7 | 1.518080669 |
| Lin54 | 1.517949373 |
| Rnpepl1 | 1.517836867 |
| Aida | 1.517692851 |
| Tspyl2 | 1.517285581 |
| Cgrrf1 | 1.517112122 |
| Rpn1 | 1.516782152 |
| Nup35 | 1.516565778 |
| Cep89 | 1.516218335 |
| Mrpl47 | 1.515432834 |
| Gatsl3 | 1.515352844 |
| Lrrc24 | 1.514920397 |
| Tap2 | 1.514285768 |
| Pde1a | 1.514252009 |
| Psmd11 | 1.512914205 |
| Dync1h1 | 1.512728621 |
| Pin1 | 1.512334616 |
| Gne | 1.51174249 |
| Far2 | 1.511719674 |
| Pex7 | 1.511603635 |
| Ube2w | 1.511492437 |
| Nrbf2 | 1.511337247 |
| Tram1l1 | 1.51069085 |
| Mamstr | 1.510375588 |
| Peli1 | 1.510233821 |
| Ndufb10 | 1.508856544 |
| Lrba | 1.508830486 |
| Nacad | 1.508749625 |
| Eif2b2 | 1.507686445 |
| Rtf1 | 1.507165762 |
| Rhoh | 1.506559904 |
| Gga3 | 1.506065808 |
| Sike1 | 1.505295106 |
| Lin7c | 1.505193129 |
| Unc119b | 1.50513606 |
| Mfsd12 | 1.50467155 |
| Fasn | 1.50465659 |
| Mrps7 | 1.504555918 |
| Lrp10 | 1.504555417 |
| Tenm4 | 1.504268572 |
| Capzb | 1.504090057 |
| Birc3 | 1.503266956 |
| Dicer1 | 1.503236605 |
| Ppp1r15b | 1.502540014 |
| Zfp64 | 1.502103455 |
| Mpp4 | 1.501712093 |

| | |
|---|---|
| Lhpp | 1.501656028 |
| Ubac1 | 1.5015439 |
| Gnl3l | 1.501341762 |
| Sdf2 | 1.501322118 |
| Tmem127 | 1.500787433 |
| Cpeb3 | 1.499934435 |
| Acaa2 | 1.499808495 |
| Ppp6c | 1.499758166 |
| Mafk | 1.499059756 |
| Spg21 | 1.498899269 |
| Crnkl1 | 1.498880425 |
| Scara3 | 1.498747221 |
| Tubg2 | 1.498125375 |
| Schip1 | 1.49691906 |
| Twsg1 | 1.496269432 |
| Tor1aip1 | 1.495943469 |
| Optn | 1.495613133 |
| Gimap6 | 1.495193954 |
| Cdkn3 | 1.494624678 |
| Armc8 | 1.493848285 |
| Ptprh | 1.493786477 |
| Vps33a | 1.49302014 |
| Gtpbp6 | 1.492683469 |
| Bckdk | 1.491948265 |
| Znhit1 | 1.491761563 |
| Coro2a | 1.491752807 |
| Rab20 | 1.490962108 |
| Cbx5 | 1.490733861 |
| Arl8a | 1.489917257 |
| Agap3 | 1.48960214 |
| Cyp26b1 | 1.489297383 |
| Ier3 | 1.488534604 |
| C4b | 1.488031962 |
| Cox5b | 1.487800321 |
| Dbt | 1.487721403 |
| Ntmt1 | 1.487630771 |
| Rtn4rl1 | 1.486886562 |
| Ndufv2 | 1.486821071 |
| Ankar | 1.486642293 |
| Coro6 | 1.486134958 |
| Aplp1 | 1.485898825 |
| Rapgef1 | 1.485573409 |
| Zwilch | 1.485434067 |
| Ube3b | 1.485365686 |
| Xpnpep1 | 1.485200847 |
| Klhl7 | 1.485154395 |
| Epha7 | 1.484917647 |

| | |
|---|---|
| Kras | 1.484366899 |
| Pfdn4 | 1.484256815 |
| Cat | 1.484232937 |
| Sema3b | 1.483106163 |
| Trit1 | 1.483099763 |
| Fam114a2 | 1.483003273 |
| Vwf | 1.482987197 |
| Pax8 | 1.482916599 |
| Ap1ar | 1.4824781 |
| Actr1b | 1.482031635 |
| Ttc39c | 1.481985582 |
| B4galt4 | 1.481937872 |
| Arpp19 | 1.481468728 |
| Gtpbp4 | 1.480532416 |
| Ccdc115 | 1.480458878 |
| Tmem55a | 1.480313534 |
| Dgat1 | 1.480032882 |
| Pdia3 | 1.479778912 |
| Atp10d | 1.479297312 |
| Pigm | 1.479098142 |
| Fam111a | 1.478862062 |
| Abi1 | 1.478796312 |
| Scnm1 | 1.478772239 |
| Ly96 | 1.478396455 |
| Pola2 | 1.478360339 |
| Batf2 | 1.478323359 |
| Zdhhc14 | 1.478282635 |
| Sirt7 | 1.478064697 |
| Pitpnc1 | 1.477976841 |
| Prdm10 | 1.477782749 |
| Rps6kb1 | 1.477679513 |
| Cdk7 | 1.477403884 |
| Nasp | 1.477188779 |
| Drap1 | 1.476800389 |
| Plec | 1.47659587 |
| Srsf4 | 1.476564685 |
| Coasy | 1.476277768 |
| Spire1 | 1.475754317 |
| Npas2 | 1.475336195 |
| Myoz2 | 1.475071711 |
| Tspan1 | 1.474773646 |
| Eif4enif1 | 1.474723833 |
| Irak1 | 1.474602903 |
| Wdr74 | 1.473973688 |
| Ibtk | 1.473077424 |
| Zdhhc24 | 1.472829538 |
| Adcy6 | 1.471918783 |

| | |
|---|---|
| Dohh | 1.471902712 |
| B4galt6 | 1.471534802 |
| Med28 | 1.471315177 |
| Trmt11 | 1.471265308 |
| Cped1 | 1.470739753 |
| Mettl3 | 1.470705517 |
| Fyn | 1.470296357 |
| Pou6f1 | 1.470254417 |
| Ifitm3 | 1.469864718 |
| Rsl24d1 | 1.469364101 |
| Osmr | 1.469272241 |
| Cc2d2a | 1.468997729 |
| Acly | 1.468928525 |
| Cdc7 | 1.468545642 |
| Mmp2 | 1.468213613 |
| Nr4a1 | 1.467671356 |
| Foxp4 | 1.4674752 |
| Mat2b | 1.467255275 |
| Cadm1 | 1.467237472 |
| Cd79b | 1.465836359 |
| Fam3a | 1.465556981 |
| Brwd1 | 1.464103547 |
| Slc4a3 | 1.463836164 |
| B3galt6 | 1.463650548 |
| Sptbn2 | 1.463163882 |
| Spata1 | 1.462463937 |
| Mageh1 | 1.462231617 |
| Slc29a1 | 1.462200516 |
| Btbd6 | 1.461781674 |
| Rabep2 | 1.461674084 |
| Tomm40 | 1.461469053 |
| Cyp4x1 | 1.461317908 |
| Susd5 | 1.461042573 |
| Cyp2r1 | 1.460839236 |
| Ranbp2 | 1.460623108 |
| Mks1 | 1.459506846 |
| Hgf | 1.459256792 |
| Bambi | 1.45877175 |
| Arl5a | 1.458692024 |
| Pir | 1.458267895 |
| B3gnt9 | 1.458254655 |
| Pnpla8 | 1.457511668 |
| Zdhhc3 | 1.457327804 |
| Cpvl | 1.457206532 |
| Mbd3 | 1.457146206 |
| Tceanc | 1.457074098 |
| Rrp7a | 1.45697634 |

| | |
|---|---|
| Cflar | 1.456548809 |
| Xpo5 | 1.456480368 |
| Map1a | 1.455993608 |
| Sec31a | 1.455365138 |
| Naa15 | 1.455097013 |
| Sema5b | 1.455037911 |
| Med11 | 1.454942638 |
| Vac14 | 1.454925213 |
| Gpsm2 | 1.454854904 |
| Fam105a | 1.454360363 |
| Kdm8 | 1.454128658 |
| Coq2 | 1.45410355 |
| Polr3gl | 1.453394614 |
| Fbxo31 | 1.452793649 |
| Wdr25 | 1.45252475 |
| Atxn7l1 | 1.452318253 |
| Simc1 | 1.452007835 |
| Fam132a | 1.451697839 |
| Dpy19l4 | 1.451158618 |
| Kdm5b | 1.450531197 |
| Lnx2 | 1.450059327 |
| Nckipsd | 1.449541945 |
| Ndufv1 | 1.449450184 |
| Ehbp1 | 1.448835011 |
| Pparg | 1.448787275 |
| Nrros | 1.448634786 |
| Tlr1 | 1.448338978 |
| Pcdh7 | 1.448088309 |
| Ccdc84 | 1.447706342 |
| Lats2 | 1.447692915 |
| Tlr6 | 1.447358964 |
| Ppara | 1.447319596 |
| Hps3 | 1.447291129 |
| Ap4s1 | 1.447287353 |
| Pithd1 | 1.447258065 |
| Kdelc1 | 1.446985892 |
| Obscn | 1.446922709 |
| Cwc27 | 1.446831689 |
| Ska3 | 1.446675716 |
| Hrct1 | 1.446277357 |
| Tmem121 | 1.446137955 |
| Ccnt2 | 1.445851024 |
| Ogdhl | 1.445638414 |
| Ttc32 | 1.445425418 |
| Snrnp70 | 1.445175786 |
| Cerkl | 1.445104503 |
| Espn | 1.445057596 |

| | |
|---|---|
| Fermt3 | 1.444852291 |
| Ankle1 | 1.444691251 |
| Smarcc1 | 1.444608103 |
| Dhrs3 | 1.443375765 |
| Sox7 | 1.442811811 |
| Tmem86b | 1.442637343 |
| Mrap2 | 1.442346905 |
| Tbc1d22b | 1.442227056 |
| Dgkd | 1.441904586 |
| Slfn5 | 1.441463678 |
| Naa35 | 1.441152655 |
| Llgl2 | 1.440964938 |
| Lyplal1 | 1.440950817 |
| Pafah1b1 | 1.440863594 |
| Csnk1g2 | 1.440559266 |
| Cenpb | 1.440432656 |
| Dbi | 1.439986405 |
| Cd302 | 1.439448128 |
| Rab6a | 1.439288481 |
| Dusp15 | 1.439026631 |
| Cstf1 | 1.439013571 |
| Mgat4a | 1.438771534 |
| Rrs1 | 1.438523966 |
| Rbak | 1.437979441 |
| Il3ra | 1.437838755 |
| Lims1 | 1.437741207 |
| Cnp | 1.437382369 |
| Cdc42bpb | 1.437252749 |
| Map9 | 1.437074885 |
| Arfip2 | 1.436875522 |
| Tada3 | 1.436686014 |
| Slc30a4 | 1.436579749 |
| Iqcb1 | 1.436435535 |
| Celf1 | 1.436413474 |
| Ptprcap | 1.436387124 |
| Ap2m1 | 1.436206793 |
| Tnpo1 | 1.436061918 |
| Per2 | 1.435520512 |
| Armc2 | 1.435340836 |
| Arl6ip1 | 1.434630661 |
| Senp5 | 1.434389886 |
| Tox2 | 1.433969697 |
| Snrpc | 1.433706861 |
| Casp12 | 1.433537505 |
| Pdpk1 | 1.433495015 |
| Tulp3 | 1.433400407 |
| Plxna1 | 1.433393837 |

| | |
|---|---|
| Dr1 | 1.433337449 |
| Smo | 1.433125516 |
| Fam32a | 1.43298611 |
| Srrm3 | 1.432582045 |
| Abcg4 | 1.43227447 |
| Ass1 | 1.431739022 |
| Smad9 | 1.431647815 |
| Aplp2 | 1.431556855 |
| Ap1g2 | 1.431418189 |
| Rell2 | 1.430489047 |
| Abhd14a | 1.430094877 |
| Ostm1 | 1.428852771 |
| Thra | 1.428529912 |
| Grin3b | 1.428494407 |
| Traf7 | 1.428343734 |
| Adamtsl1 | 1.42834339 |
| Ankrd34a | 1.428207417 |
| Pcdh10 | 1.427960424 |
| Hsd17b11 | 1.427150299 |
| Mlec | 1.426822572 |
| Hspe1 | 1.425951049 |
| Cggbp1 | 1.425349907 |
| Zcchc4 | 1.425266216 |
| Scyl2 | 1.425189175 |
| Aass | 1.424848433 |
| Gabpb2 | 1.424590868 |
| Flot2 | 1.423925497 |
| Erlin1 | 1.423203842 |
| Irs1 | 1.423127738 |
| Fkbp5 | 1.423064252 |
| Arl1 | 1.422717559 |
| G2e3 | 1.422514498 |
| Lrrc16a | 1.422322164 |
| Vwce | 1.422232696 |
| Mest | 1.422153788 |
| Vps13d | 1.422097416 |
| Wdr7 | 1.421781031 |
| Prosc | 1.421709923 |
| Ap1s2 | 1.421455715 |
| Dzip3 | 1.421225768 |
| Dido1 | 1.421012016 |
| Pcdh9 | 1.420365119 |
| Acp2 | 1.420359343 |
| Mccc2 | 1.42023024 |
| Zbed4 | 1.420009571 |
| Sgol2 | 1.419860423 |
| Polr1a | 1.419837466 |

| | |
|---|---|
| Spsb1 | 1.419727721 |
| Atxn1 | 1.419261593 |
| Sdc4 | 1.419121839 |
| Rpl13a | 1.41889542 |
| Stard3nl | 1.418722691 |
| Usf2 | 1.418578018 |
| Dpysl4 | 1.417574865 |
| Trio | 1.417113775 |
| Cnpy4 | 1.415819867 |
| Ccdc34 | 1.415673009 |
| Nfkbid | 1.415657989 |
| Ndufa4 | 1.415592559 |
| Rpl26 | 1.414034481 |
| Retsat | 1.413909827 |
| Eif2a | 1.413069227 |
| Trim14 | 1.412831452 |
| Pisd | 1.411309626 |
| Dazap2 | 1.411013121 |
| Wbp1l | 1.410490556 |
| Ppp2cb | 1.41047606 |
| Vamp4 | 1.410099412 |
| Fau | 1.410072483 |
| Dnmt3a | 1.409896049 |
| Cndp2 | 1.409766306 |
| Nup37 | 1.409140157 |
| Plagl1 | 1.409088414 |
| Mbtps1 | 1.408974947 |
| Stxbp2 | 1.408433214 |
| Smdt1 | 1.40839815 |
| Ascc3 | 1.408363453 |
| Cenpl | 1.408180041 |
| Dscc1 | 1.408038533 |
| Chpt1 | 1.407949312 |
| Uckl1 | 1.407605105 |
| Nfkbiz | 1.407018306 |
| Dis3 | 1.406291845 |
| Evi5 | 1.405087665 |
| Lin7a | 1.40483599 |
| Tbl1xr1 | 1.404736953 |
| Col4a6 | 1.403584724 |
| Osgin1 | 1.403388677 |
| Adrm1 | 1.403310759 |
| Ell | 1.403064816 |
| Slc25a45 | 1.402051594 |
| Pradc1 | 1.402016772 |
| Ccdc85c | 1.401109128 |
| Epc1 | 1.400859129 |

| | |
|---|---|
| Plod1 | 1.400231081 |
| Baiap2l1 | 1.400204753 |
| Tbc1d13 | 1.399568377 |
| Tmem81 | 1.398461143 |
| Morn4 | 1.39789823 |
| Mrps30 | 1.39769169 |
| Snrnp27 | 1.396817566 |
| Mphosph10 | 1.396625998 |
| Laptm5 | 1.396146437 |
| Ptprb | 1.395167778 |
| Mmp7 | 1.395008149 |
| Plxna4 | 1.394751717 |
| Alg13 | 1.394535882 |
| Hes6 | 1.393465821 |
| Rpap1 | 1.393130053 |
| Col4a5 | 1.392918989 |
| Naga | 1.392845552 |
| Kri1 | 1.392811536 |
| Nt5e | 1.392760615 |
| Sec61g | 1.392731116 |
| Chchd2 | 1.392420732 |
| Abcc3 | 1.39101297 |
| Klhl13 | 1.390996963 |
| Anp32a | 1.390835666 |
| Pml | 1.390572264 |
| Fam72a | 1.390001119 |
| Ccdc85a | 1.389310235 |
| Mtmr9 | 1.389220515 |
| Pacs2 | 1.388711459 |
| Apom | 1.387672646 |
| Coa3 | 1.386850574 |
| Nme6 | 1.386415955 |
| Txnrd1 | 1.386241809 |
| Actr1a | 1.386128462 |
| Abcc1 | 1.3855091 |
| Plk1 | 1.384900749 |
| Sh3kbp1 | 1.384427355 |
| Ahrr | 1.384193407 |
| Haus7 | 1.38417168 |
| Lhfp | 1.384160863 |
| Lin52 | 1.383478294 |
| Snx27 | 1.383344427 |
| Rap2b | 1.383060424 |
| Ttll1 | 1.38241247 |
| Tmem50a | 1.382319569 |
| Med26 | 1.381882616 |
| Cgref1 | 1.38171183 |

| | |
|---|---|
| Rnmtl1 | 1.381546192 |
| Ier2 | 1.381409173 |
| Kcnc4 | 1.381242067 |
| Sox6 | 1.381110703 |
| Kif17 | 1.381076922 |
| Pdxdc1 | 1.379978688 |
| Mpst | 1.379194098 |
| Rpl14 | 1.379113019 |
| Nckap5l | 1.378419663 |
| Gtf2h3 | 1.377458488 |
| Sigirr | 1.377392963 |
| Aurkaip1 | 1.377198242 |
| Trnp1 | 1.377091611 |
| Rbm25 | 1.376184838 |
| Galnt3 | 1.375870329 |
| Postn | 1.375505952 |
| Thrb | 1.375233626 |
| Bbs7 | 1.37490664 |
| Mrpl38 | 1.374818832 |
| Tbc1d30 | 1.374031476 |
| Wnk3 | 1.374029067 |
| Tmem242 | 1.3737421 |
| Psd3 | 1.373713348 |
| Smim7 | 1.37299543 |
| Ptrh2 | 1.372567461 |
| Harbi1 | 1.372395808 |
| Gpx1 | 1.371946834 |
| Rbpj | 1.371826879 |
| Sdcbp2 | 1.371781051 |
| Cep170b | 1.371578572 |
| Msx1 | 1.371302158 |
| Cpsf1 | 1.371223555 |
| Ndufb3 | 1.370943071 |
| Apold1 | 1.370579727 |
| Bloc1s1 | 1.370492483 |
| Gpc5 | 1.370475365 |
| Gpn3 | 1.370427684 |
| Glrx | 1.369785453 |
| Sil1 | 1.369579008 |
| Sub1 | 1.369297029 |
| Morf4l2 | 1.369249275 |
| Abca4 | 1.369185208 |
| Dkk2 | 1.369051091 |
| Orc2 | 1.369020793 |
| Tnfrsf25 | 1.368778168 |
| Numa1 | 1.368009241 |
| Kpna2 | 1.367890137 |

| | |
|---|---|
| Rpl7a | 1.367737309 |
| Cul2 | 1.366440466 |
| Lgals1 | 1.366306963 |
| Slc2a10 | 1.365799815 |
| Arf6 | 1.365785197 |
| Plekha8 | 1.36509427 |
| Fbxo3 | 1.364982549 |
| Hdac5 | 1.36461716 |
| Prkdc | 1.36442736 |
| Lmbr1 | 1.363987339 |
| Rtca | 1.363965371 |
| Appbp2 | 1.363559024 |
| Sec62 | 1.362764736 |
| Ap2b1 | 1.362310186 |
| Foxo3 | 1.362223171 |
| Hsp90aa1 | 1.362051415 |
| Med13l | 1.360945126 |
| Sema6c | 1.359733853 |
| Tnfaip8l3 | 1.359522426 |
| Dyrk1b | 1.359513311 |
| Mrpl54 | 1.358860526 |
| Nek1 | 1.35854654 |
| Rps6ka4 | 1.358160212 |
| Rnf19b | 1.357005864 |
| Sipa1 | 1.356564016 |
| Mid1ip1 | 1.35566941 |
| Msantd3 | 1.355649297 |
| Pfkfb3 | 1.355425159 |
| Gucy1a2 | 1.355278081 |
| Mettl16 | 1.35473078 |
| Add2 | 1.354515453 |
| Mecom | 1.354229245 |
| Wdr59 | 1.353873501 |
| Tro | 1.353583785 |
| Cecr5 | 1.353297373 |
| Idh3a | 1.353178892 |
| Ubald2 | 1.353128587 |
| Cln6 | 1.352917048 |
| Eri2 | 1.352367459 |
| Antxr2 | 1.352251866 |
| Gstcd | 1.35180783 |
| Dnah11 | 1.351625508 |
| Gstm2 | 1.351017284 |
| Dmwd | 1.350991149 |
| Sox8 | 1.350861984 |
| Pcyox1l | 1.350795679 |
| Glt8d1 | 1.350705268 |

| | |
|---|---|
| Timp3 | 1.350318148 |
| Rdh13 | 1.349622882 |
| Anxa3 | 1.349621162 |
| Katna1 | 1.349267035 |
| Crtc1 | 1.349260276 |
| Nt5c3b | 1.349241466 |
| Morn1 | 1.34840161 |
| Pank3 | 1.347537496 |
| Fzr1 | 1.347114805 |
| Steap1 | 1.346811512 |
| Mmp15 | 1.346699411 |
| Bcas2 | 1.346502666 |
| Trove2 | 1.346268895 |
| Ager | 1.345916176 |
| Ddn | 1.345910666 |
| Prkrir | 1.345896144 |
| Yipf3 | 1.345752617 |
| Fgfr1op | 1.345704691 |
| Gse1 | 1.345594592 |
| Lrrc36 | 1.345348306 |
| Coq10a | 1.345161803 |
| Ubr2 | 1.344770101 |
| Tnfrsf11a | 1.344172414 |
| Clpp | 1.343704264 |
| Fancb | 1.34355058 |
| Thyn1 | 1.343548919 |
| Alg8 | 1.343253095 |
| Tsc22d4 | 1.342869082 |
| Eftud1 | 1.342818783 |
| P4ha1 | 1.34271926 |
| Tmem19 | 1.342013978 |
| Dcp1b | 1.341871934 |
| Zfr | 1.341719382 |
| Plac8 | 1.341684078 |
| Rps19 | 1.340919953 |
| Dfnb59 | 1.340299747 |
| Ninl | 1.339831934 |
| Scamp4 | 1.339615469 |
| Hist1h4j | 1.339536461 |
| Kntc1 | 1.339208651 |
| Samd8 | 1.338867348 |
| Dph3 | 1.338118112 |
| Rnh1 | 1.338020489 |
| Adamts10 | 1.33783863 |
| Fut8 | 1.337733985 |
| Jkamp | 1.337243951 |
| Mbd2 | 1.336956864 |

| | |
|---|---|
| Tal1 | 1.336536148 |
| Smc2 | 1.336414547 |
| Ciita | 1.336046304 |
| Lrfn4 | 1.335615175 |
| Abi3 | 1.335369093 |
| Ict1 | 1.335153053 |
| Fam50b | 1.335110141 |
| Fxyd6 | 1.3349763 |
| Stub1 | 1.334552369 |
| Disc1 | 1.3345153 |
| Esyt2 | 1.334380217 |
| Golgb1 | 1.334215581 |
| Wbp4 | 1.333857467 |
| Dtl | 1.333856514 |
| Psmd5 | 1.333698347 |
| Ehbp1l1 | 1.333387145 |
| Gabre | 1.33317042 |
| Ppp2r5d | 1.332777896 |
| Sel1l | 1.332413937 |
| Hes2 | 1.332116829 |
| Abcb6 | 1.331957083 |
| Pdk1 | 1.331909923 |
| Xpo1 | 1.331869443 |
| Cdk11b | 1.331745103 |
| Nr3c1 | 1.33153223 |
| Npm3 | 1.331237572 |
| Zfp92 | 1.33079437 |
| Ube2v2 | 1.330464875 |
| Ireb2 | 1.330302649 |
| Atg9a | 1.329907566 |
| Zswim7 | 1.329864445 |
| Gpr19 | 1.329693636 |
| Mief1 | 1.329677783 |
| Ppp6r2 | 1.329341013 |
| Katnb1 | 1.329194659 |
| Psmc3ip | 1.328674025 |
| Mrps18c | 1.328191819 |
| Ifngr2 | 1.328123568 |
| Psap | 1.327628458 |
| Upk3bl | 1.327581649 |
| Rhod | 1.327552684 |
| Eif2s1 | 1.327197067 |
| Caprin2 | 1.326557009 |
| Chd4 | 1.326517552 |
| Slc9a8 | 1.326094589 |
| Slc8b1 | 1.325029858 |
| Kif24 | 1.324607947 |

| | |
|---|---|
| Nrep | 1.324548314 |
| Rhoa | 1.323930105 |
| Rhbdd2 | 1.323866336 |
| Trap1 | 1.322994248 |
| Mrc2 | 1.322192262 |
| Armcx5 | 1.32209431 |
| Mfn2 | 1.321610116 |
| Mef2a | 1.321356246 |
| Clptm1l | 1.321207859 |
| Ngly1 | 1.321035013 |
| Ptprd | 1.320805364 |
| Zbtb12 | 1.320707274 |
| Npr2 | 1.320508752 |
| Uba1 | 1.319008608 |
| Serpinb2 | 1.318932919 |
| Cers1 | 1.318414223 |
| Ccnyl1 | 1.318409545 |
| Medag | 1.318188419 |
| Elfn1 | 1.317971635 |
| Fam189a1 | 1.317088449 |
| Gna12 | 1.316811961 |
| Pold2 | 1.31675713 |
| Tatdn3 | 1.316737021 |
| Tmem87b | 1.316406834 |
| Zbtb7a | 1.316376489 |
| Itgax | 1.315875371 |
| Gas2 | 1.31553603 |
| Mms19 | 1.314799442 |
| Bend6 | 1.314791957 |
| Tspo | 1.314757039 |
| Sec61a2 | 1.313747838 |
| Myc | 1.313641741 |
| Phc1 | 1.31334826 |
| Tll1 | 1.313166305 |
| Slc35e1 | 1.313130927 |
| Tsc2 | 1.312495026 |
| Rbm6 | 1.312491838 |
| Magix | 1.312341186 |
| Gnaq | 1.312000145 |
| Ywhaz | 1.311539875 |
| Gemin8 | 1.310687455 |
| Dpf3 | 1.310337233 |
| Cenpt | 1.310240631 |
| Poglut1 | 1.309497405 |
| Saysd1 | 1.309430605 |
| Dmd | 1.309251848 |
| Mogs | 1.309209531 |

| | |
|---|---|
| Klhdc9 | 1.30850122 |
| Cd68 | 1.308178612 |
| Dlat | 1.306564556 |
| Pam16 | 1.306270446 |
| Msrb3 | 1.305956726 |
| Nudt9 | 1.305747149 |
| Cdk2ap2 | 1.30543119 |
| Traf5 | 1.305179761 |
| Dusp1 | 1.305178568 |
| Acox2 | 1.305073784 |
| Thsd4 | 1.304325392 |
| Drosha | 1.304034261 |
| Cadm3 | 1.303785221 |
| Mtch2 | 1.303643049 |
| Ints10 | 1.302917511 |
| Wtap | 1.302707471 |
| Pigp | 1.302535627 |
| Crybg3 | 1.3025071 |
| Smc1a | 1.302295544 |
| Exoc5 | 1.302034146 |
| Veph1 | 1.301707825 |
| Micall1 | 1.301485742 |
| Gja1 | 1.301363126 |
| Hk1 | 1.30053524 |
| Six5 | 1.300512597 |
| Atpaf2 | 1.300116202 |
| Phka2 | 1.299852763 |
| Dcaf10 | 1.299502901 |
| Sprtn | 1.299370571 |
| Sat1 | 1.299342052 |
| Ubtd2 | 1.29934049 |
| Phtf2 | 1.299063434 |
| Gpr161 | 1.298095303 |
| Dnajb2 | 1.297615454 |
| Vrk2 | 1.297435051 |
| Ulk4 | 1.296503198 |
| Pdgfb | 1.296387696 |
| Wdyhv1 | 1.296183959 |
| Igf2 | 1.295952658 |
| Rcor3 | 1.295758343 |
| Crtap | 1.295571859 |
| Vldlr | 1.295350871 |
| Cntn4 | 1.295243005 |
| Rad51b | 1.294892005 |
| Stard5 | 1.294745219 |
| Aldh18a1 | 1.294667563 |
| Gfpt1 | 1.294458207 |

| | |
|---|---|
| Nup153 | 1.294432043 |
| Atp11b | 1.294315071 |
| Srsf7 | 1.294098229 |
| Ttc21b | 1.293861257 |
| Isca2 | 1.293761923 |
| Tpgs1 | 1.293626384 |
| Mgp | 1.293521318 |
| Dmxl1 | 1.293456804 |
| Tipin | 1.29318138 |
| Dhdds | 1.293157522 |
| Ctnnb1 | 1.293123913 |
| Fkbp10 | 1.292529433 |
| Gsta4 | 1.292362859 |
| Cep112 | 1.292025916 |
| Tceb3 | 1.291427544 |
| Slc25a46 | 1.291266778 |
| Ddx49 | 1.290883309 |
| Ano1 | 1.290816568 |
| Ptchd4 | 1.290764918 |
| Atp1b3 | 1.290603225 |
| Kansl3 | 1.290438486 |
| Ints3 | 1.290147616 |
| Ddx23 | 1.28986886 |
| Elf4 | 1.289833063 |
| Cpa4 | 1.289429292 |
| Ythdf1 | 1.289275937 |
| Sdc2 | 1.289066252 |
| Serpina5 | 1.288954018 |
| Ulk2 | 1.288864827 |
| Wdr92 | 1.288846204 |
| Med15 | 1.288678201 |
| Erbb2 | 1.288669063 |
| Hist1h2bh | 1.288606306 |
| Micall2 | 1.287998165 |
| Impact | 1.28783535 |
| Sat2 | 1.287575347 |
| Cetn3 | 1.286498505 |
| Nln | 1.286473733 |
| Shisa4 | 1.286232133 |
| Mier2 | 1.285709781 |
| Tnk1 | 1.285341082 |
| Ltbr | 1.284957515 |
| Thoc6 | 1.284594518 |
| Gstz1 | 1.284099937 |
| Rbpms | 1.283646689 |
| Plagl2 | 1.283435479 |
| Herc4 | 1.282894482 |

| | |
|---|---|
| Scamp2 | 1.282538566 |
| Rin1 | 1.28250086 |
| Tcp11l2 | 1.282487393 |
| Aaed1 | 1.28232367 |
| Haus3 | 1.282255073 |
| Lipt1 | 1.281701918 |
| Podxl2 | 1.281532832 |
| Psmd10 | 1.281481524 |
| Med22 | 1.280831308 |
| Tmco6 | 1.280469933 |
| Nfyc | 1.280330869 |
| Nlrp3 | 1.280220406 |
| Gdf6 | 1.279200377 |
| Rassf10 | 1.27911471 |
| Cables2 | 1.278813153 |
| Atpif1 | 1.278570481 |
| Ppme1 | 1.278548733 |
| Pcnx | 1.277845767 |
| Ephb2 | 1.277714316 |
| Arfgap3 | 1.277684751 |
| Rab11fip2 | 1.27753916 |
| Ccdc121 | 1.277452496 |
| Insr | 1.277148077 |
| Rgcc | 1.275434457 |
| Timp1 | 1.275427473 |
| Mavs | 1.275247622 |
| Ubb | 1.274423896 |
| Ccng2 | 1.274366168 |
| Nsa2 | 1.274320886 |
| Manea | 1.274086218 |
| Trim6 | 1.273614811 |
| Slc29a4 | 1.27352203 |
| Ube3c | 1.273104841 |
| Mapk13 | 1.272999323 |
| Slc25a51 | 1.272879924 |
| Tmod2 | 1.27247878 |
| Nck1 | 1.272390658 |
| Snap91 | 1.272316749 |
| Atf6b | 1.27228744 |
| Zfand3 | 1.271695225 |
| Mcee | 1.271601356 |
| Usp42 | 1.271238204 |
| Nlrc3 | 1.271175452 |
| Slc41a3 | 1.271034235 |
| Pgm3 | 1.270894982 |
| Sparc | 1.270765728 |
| Pigf | 1.270076713 |

| | |
|---|---|
| Mei1 | 1.269868907 |
| Wdr46 | 1.269845498 |
| Ewsr1 | 1.269644551 |
| Hmgb1 | 1.269489951 |
| Apip | 1.26942297 |
| Pikfyve | 1.269313966 |
| Il18r1 | 1.269176721 |
| L3mbtl1 | 1.268798165 |
| Acot7 | 1.268429475 |
| Rab3a | 1.268428871 |
| Scrn2 | 1.268210941 |
| Tbca | 1.268064118 |
| Mrpl13 | 1.268050375 |
| Taok3 | 1.267162284 |
| Mphosph9 | 1.266331788 |
| Cdc42se1 | 1.266028643 |
| Lsm5 | 1.265958632 |
| Rai1 | 1.26556642 |
| Cbx4 | 1.265247981 |
| Pdss1 | 1.265027855 |
| Hsph1 | 1.264874355 |
| Surf2 | 1.264524589 |
| Zdhhc9 | 1.264501975 |
| Aarsd1 | 1.264449792 |
| Fkbp11 | 1.264436508 |
| Spred1 | 1.264113267 |
| Spice1 | 1.263782289 |
| Ilf3 | 1.263181232 |
| Trmt44 | 1.262738236 |
| Spata2l | 1.262440305 |
| Steap3 | 1.262420097 |
| Srrt | 1.262338847 |
| Pih1d2 | 1.262014663 |
| Hif1a | 1.261738338 |
| Cd3eap | 1.261175889 |
| Phf13 | 1.261144332 |
| Cept1 | 1.261134816 |
| Gstm1 | 1.261121777 |
| Phyh | 1.260903887 |
| Kdelr1 | 1.260675735 |
| Fam35a | 1.260492895 |
| Jmjd6 | 1.260435154 |
| Fggy | 1.259130381 |
| Trpc1 | 1.259017039 |
| Nebl | 1.258954067 |
| Sap30l | 1.258213495 |
| Armc9 | 1.258056507 |

| | |
|---|---|
| Dars2 | 1.257695564 |
| Tmem182 | 1.257215088 |
| Ino80d | 1.256887943 |
| Prim2 | 1.25672766 |
| Frk | 1.256637429 |
| P4hb | 1.256568775 |
| Copb1 | 1.256099254 |
| Sh3bgrl3 | 1.255607998 |
| Ftsj3 | 1.25459728 |
| Klhl15 | 1.254003331 |
| Rnf11 | 1.25387178 |
| Gpatch2 | 1.253587167 |
| Rfc4 | 1.253470331 |
| Enoph1 | 1.253227761 |
| Pigv | 1.253215797 |
| Rab27a | 1.253046898 |
| Mapk12 | 1.252451099 |
| Il1r1 | 1.251874311 |
| Mcm6 | 1.251284861 |
| Yipf6 | 1.251240065 |
| Mcat | 1.250928288 |
| Stk17b | 1.250841001 |
| Fancg | 1.250732639 |
| Cecr6 | 1.24957746 |
| Itpr2 | 1.249437669 |
| Clspn | 1.249425162 |
| Zc4h2 | 1.249231602 |
| Mettl14 | 1.2489397 |
| Zdhhc4 | 1.248720449 |
| Cdc27 | 1.248510641 |
| Rbm15b | 1.248441075 |
| Usp2 | 1.248346212 |
| Dlg1 | 1.248136116 |
| Lurap1 | 1.247770703 |
| Appl1 | 1.24774302 |
| Ldoc1l | 1.247557058 |
| Rpa2 | 1.246653993 |
| Inpp1 | 1.246637385 |
| Atrn | 1.246574179 |
| Plxna2 | 1.246495178 |
| Rps27a | 1.246134539 |
| Ppp1r12a | 1.24605495 |
| Stat4 | 1.245964288 |
| Pltp | 1.24577939 |
| Zfp90 | 1.245718238 |
| Hsp90b1 | 1.245500746 |
| Coq9 | 1.244916728 |

| | |
|---|---|
| Anapc15 | 1.244896562 |
| Fam160a2 | 1.244371866 |
| Dlgap1 | 1.244243738 |
| Acsl4 | 1.243546356 |
| Capza2 | 1.243543554 |
| Cmtm3 | 1.24260176 |
| Traf6 | 1.242287547 |
| Tra2a | 1.242055382 |
| Wdr20 | 1.242042888 |
| Tsr1 | 1.2420208 |
| Klhdc3 | 1.241689637 |
| Dus4l | 1.241317066 |
| Sepw1 | 1.241253005 |
| Slc16a5 | 1.241191177 |
| Ell3 | 1.241121162 |
| Gas2l3 | 1.240991537 |
| Stard6 | 1.24079019 |
| Rnasel | 1.24043135 |
| Il7r | 1.240430292 |
| Slit2 | 1.239989257 |
| Gjd3 | 1.239904261 |
| Abca1 | 1.239728723 |
| Pank1 | 1.239724092 |
| Tmem136 | 1.239514596 |
| Fam65a | 1.239451296 |
| Ppp6r1 | 1.23901026 |
| Raf1 | 1.238669201 |
| Vash2 | 1.238167979 |
| Fbxo38 | 1.237582081 |
| Fam63a | 1.237551943 |
| Chaf1a | 1.237430044 |
| Ppm1h | 1.237365143 |
| Tmem200b | 1.236682511 |
| Ankra2 | 1.236238177 |
| Nubp1 | 1.236237431 |
| Ppp4c | 1.236165347 |
| Mpv17l2 | 1.235995987 |
| Phgdh | 1.235948129 |
| Cul4a | 1.235714188 |
| Trappc6a | 1.235430768 |
| Rtn1 | 1.235054714 |
| Rictor | 1.234747556 |
| St8sia6 | 1.234018946 |
| Frem1 | 1.233888339 |
| Commd6 | 1.233745501 |
| Ppip5k2 | 1.233739051 |
| Slc35a1 | 1.233353177 |

| | |
|---|---|
| Pigq | 1.233338325 |
| Foxp2 | 1.233316879 |
| Lum | 1.233232838 |
| Atp6v1d | 1.232801345 |
| Spata21 | 1.232778795 |
| Cenpq | 1.232559537 |
| Arhgef2 | 1.232297453 |
| Ralgds | 1.231474289 |
| C1rl | 1.231470801 |
| Klhl28 | 1.231230332 |
| Rps7 | 1.230963963 |
| Aasdh | 1.230865455 |
| Olfm2 | 1.230527841 |
| Gata3 | 1.230400303 |
| Rnf5 | 1.230147555 |
| Cgn | 1.229867615 |
| Polr2a | 1.229617166 |
| Akr1b10 | 1.229147189 |
| Ets2 | 1.228772914 |
| Alg12 | 1.228665042 |
| Uqcr11 | 1.228262904 |
| Fpgs | 1.227991354 |
| Banp | 1.227631247 |
| Rragd | 1.227429256 |
| Shroom3 | 1.226559095 |
| Tbpl1 | 1.226513843 |
| Ccnh | 1.226078642 |
| Map4 | 1.224873771 |
| Tecpr1 | 1.224833468 |
| Smg8 | 1.224815671 |
| Tcirg1 | 1.223886359 |
| Hoxa3 | 1.223791771 |
| Pcmt1 | 1.223198501 |
| Fibp | 1.222854806 |
| Ndufaf4 | 1.222830457 |
| Pja2 | 1.222315733 |
| Tomm20 | 1.221552528 |
| Psip1 | 1.22146025 |
| Apex2 | 1.220981539 |
| Csgalnact1 | 1.220776667 |
| Vsig10 | 1.220727339 |
| Zadh2 | 1.220694261 |
| Carkd | 1.220390437 |
| Crebrf | 1.220328184 |
| Pik3ap1 | 1.219524912 |
| Hells | 1.219419237 |
| Dzank1 | 1.219018412 |

| | |
|---|---|
| Xiap | 1.218959039 |
| Net1 | 1.218751423 |
| Ctnnbip1 | 1.218686285 |
| Htra2 | 1.218502516 |
| Psma7 | 1.218434497 |
| Plekho2 | 1.218389003 |
| Pcsk5 | 1.218384601 |
| Cnst | 1.217913141 |
| Garem | 1.21721009 |
| Tmem50b | 1.216786406 |
| Ptgs1 | 1.216491575 |
| Rps15a | 1.216434188 |
| Mapk8 | 1.21641459 |
| Ctsz | 1.215907084 |
| Pknox2 | 1.215242054 |
| Aldh16a1 | 1.21509126 |
| Rdh14 | 1.214254993 |
| Fn1 | 1.213802723 |
| Rps6ka1 | 1.213588251 |
| Ankrd53 | 1.213519276 |
| Kcnh7 | 1.212983005 |
| Vamp1 | 1.212929001 |
| Nipsnap1 | 1.211613857 |
| Ankrd9 | 1.211589643 |
| Smg6 | 1.210156199 |
| Cdk10 | 1.209770579 |
| Ube2h | 1.2097659 |
| Vcan | 1.209557633 |
| Tmem80 | 1.209398134 |
| Ep400 | 1.209144457 |
| Smad1 | 1.209077546 |
| Ogfr | 1.208396599 |
| Tuba8 | 1.208314629 |
| Bcap29 | 1.208210549 |
| Ncl | 1.20774681 |
| Gapvd1 | 1.207309587 |
| Rps24 | 1.207156214 |
| Lrp3 | 1.207097985 |
| Gsn | 1.20699603 |
| Atp6v1g1 | 1.206666582 |
| Btn2a2 | 1.206528236 |
| Ccdc113 | 1.206189848 |
| Shc3 | 1.205959852 |
| Cuzd1 | 1.205746593 |
| Efhd2 | 1.205606691 |
| Dyx1c1 | 1.205437942 |
| Snrpa1 | 1.204562499 |

| | |
|---|---|
| Dap3 | 1.204059283 |
| Fchsd1 | 1.203914357 |
| Ak1 | 1.203850729 |
| Pdcd6 | 1.203537225 |
| Mtbp | 1.203495094 |
| Stard8 | 1.203227155 |
| Tpm4 | 1.203164724 |
| Rttn | 1.202920456 |
| Cstf2t | 1.202810787 |
| Nceh1 | 1.202779378 |
| Tbcd | 1.202692359 |
| Safb | 1.202478546 |
| Tspan12 | 1.202390857 |
| Pnpo | 1.202047517 |
| Ramp2 | 1.20197363 |
| Ptges2 | 1.20096754 |
| Cth | 1.200050692 |
| Kctd3 | 1.199068066 |
| Fam207a | 1.198980821 |
| Rsad2 | 1.198825274 |
| Tor3a | 1.1986803 |
| Ppp2r5e | 1.198538742 |
| Dtwd1 | 1.197962848 |
| Ifi44 | 1.197920662 |
| Abt1 | 1.197770349 |
| Emcn | 1.197646256 |
| Smarcd1 | 1.197413914 |
| Rnaseh1 | 1.197221424 |
| Ddx28 | 1.197185775 |
| Camk2b | 1.196837561 |
| Bmp6 | 1.196760303 |
| Aldh3a2 | 1.196587511 |
| Fam49b | 1.195880728 |
| Eapp | 1.194722902 |
| Filip1l | 1.194328505 |
| Pgp | 1.193384596 |
| Tax1bp1 | 1.193197342 |
| Myo19 | 1.19318601 |
| Nudcd1 | 1.192991745 |
| Sdc1 | 1.192660376 |
| Igsf3 | 1.192269846 |
| Wrap53 | 1.191862189 |
| Zc3h6 | 1.191082285 |
| Dlg4 | 1.190992569 |
| Rbl2 | 1.190884685 |
| Dnaaf2 | 1.1908747 |
| Gadd45gip1 | 1.190818857 |

| | |
|---|---|
| Crygs | 1.189682851 |
| Spon1 | 1.189516592 |
| Asna1 | 1.189511128 |
| Rffl | 1.189129886 |
| Dhx37 | 1.188585074 |
| Chst3 | 1.188028491 |
| Klf10 | 1.187461521 |
| Wrnip1 | 1.187325997 |
| Mpi | 1.186925047 |
| Cd93 | 1.186704305 |
| Dixdc1 | 1.186411145 |
| Camk2g | 1.18631079 |
| Zmym5 | 1.186296066 |
| Ehmt1 | 1.185953271 |
| Mettl21a | 1.184169237 |
| Trib1 | 1.183744967 |
| Ube2a | 1.183701621 |
| Rpl17 | 1.182844823 |
| Gnai3 | 1.182267334 |
| Napg | 1.181774505 |
| Mcc | 1.181352259 |
| Dpf2 | 1.181117996 |
| C2cd2 | 1.180744148 |
| Cxcl2 | 1.180358799 |
| Slc31a2 | 1.180358352 |
| Kpna1 | 1.180169238 |
| Nod1 | 1.179822543 |
| Nr1i3 | 1.179470479 |
| Dll4 | 1.179339522 |
| Fstl1 | 1.179283526 |
| Uprt | 1.178077912 |
| Pomp | 1.176997655 |
| Mcm9 | 1.176971083 |
| Pde7b | 1.176789105 |
| Rnf44 | 1.176695015 |
| Tacstd2 | 1.17667677 |
| Tnrc6a | 1.176662972 |
| Glb1 | 1.176087813 |
| Zbtb45 | 1.175948297 |
| Acsf2 | 1.175918672 |
| Dtnbp1 | 1.175898003 |
| Nfkbib | 1.175299786 |
| Brd2 | 1.175271747 |
| Papss2 | 1.175217185 |
| Eif4e2 | 1.174567639 |
| Exoc6 | 1.174150781 |
| Pcbd2 | 1.174116223 |

| | |
|---|---|
| Tacr2 | 1.173737527 |
| Hspa13 | 1.173701464 |
| Unk | 1.173665542 |
| C2cd3 | 1.173511037 |
| Eftud2 | 1.173374618 |
| Ift88 | 1.173317862 |
| Ss18l1 | 1.173196348 |
| Wdr73 | 1.173146699 |
| Usp22 | 1.172889082 |
| Ankrd42 | 1.172526074 |
| Aurkc | 1.17251054 |
| Mycbp | 1.172324003 |
| Slc35a3 | 1.171305275 |
| Rassf4 | 1.171212878 |
| Cdkn1c | 1.171046675 |
| Tgfb2 | 1.171006898 |
| Sdhb | 1.170087577 |
| Prkcd | 1.168953237 |
| Ralgapa2 | 1.168613925 |
| Ephx4 | 1.168467179 |
| Baiap2 | 1.167843504 |
| Col5a3 | 1.167797732 |
| Rragb | 1.167783845 |
| Pygb | 1.167778279 |
| Lgr4 | 1.167645346 |
| Pck2 | 1.167120049 |
| Gbp4 | 1.165340063 |
| Atp2a2 | 1.165260502 |
| Dnajc25 | 1.165056807 |
| Fam63b | 1.164978877 |
| Stx3 | 1.16461102 |
| Parp16 | 1.163813243 |
| Nfe2l1 | 1.16323414 |
| Ptp4a2 | 1.163159704 |
| Srd5a3 | 1.163070342 |
| Ing1 | 1.16282749 |
| Tmem200a | 1.162798907 |
| Crk | 1.162735041 |
| Fbxo48 | 1.162619022 |
| Pde5a | 1.162508556 |
| Pars2 | 1.162291582 |
| Ptk7 | 1.162061995 |
| Mns1 | 1.162005812 |
| Nipal1 | 1.161852127 |
| Hk2 | 1.161435129 |
| Gfod1 | 1.161247776 |
| Cebpd | 1.160441105 |

| | |
|---|---|
| Mctp2 | 1.160271604 |
| Akap13 | 1.159714963 |
| Dgkz | 1.15965994 |
| Tstd1 | 1.158975629 |
| Tmem223 | 1.15885191 |
| Anapc7 | 1.158832844 |
| Gtf2h1 | 1.15876582 |
| Prr5 | 1.158062935 |
| Traf2 | 1.158018119 |
| Tada2a | 1.157715997 |
| Rbks | 1.157397765 |
| Xrra1 | 1.157320681 |
| Psmc2 | 1.157283237 |
| Ncam2 | 1.15726747 |
| Snx19 | 1.155837762 |
| Lrrc46 | 1.155738192 |
| Acbd4 | 1.155345281 |
| Dlg5 | 1.154463055 |
| Ap3d1 | 1.154190797 |
| Usp37 | 1.154034417 |
| Ube2q1 | 1.153783375 |
| Zbtb37 | 1.152547263 |
| Adcy7 | 1.152085166 |
| Lrrcc1 | 1.150636279 |
| Rell1 | 1.150431787 |
| Tmtc2 | 1.150180479 |
| Gnpnat1 | 1.150166355 |
| Sfxn1 | 1.150123835 |
| Ap4b1 | 1.149802542 |
| B4galt5 | 1.149731038 |
| Dgcr2 | 1.148649575 |
| Cd276 | 1.148538962 |
| Ccdc53 | 1.148403922 |
| Dnajc16 | 1.148323935 |
| Fstl5 | 1.148057695 |
| Gatsl2 | 1.14803555 |
| Gpr153 | 1.147900846 |
| Ephb4 | 1.147619115 |
| Mycl | 1.147428745 |
| Rbm10 | 1.146938748 |
| Ep300 | 1.145992734 |
| Nipsnap3b | 1.145854038 |
| Ttc36 | 1.144878678 |
| Synj2bp | 1.144440062 |
| Ndufaf3 | 1.144309485 |
| Csnk2a2 | 1.144289957 |
| Cxcl1 | 1.14426672 |

| | |
|---|---|
| Ntng1 | 1.143977085 |
| Prnd | 1.142872636 |
| Tm7sf2 | 1.142288563 |
| Gpm6b | 1.14220726 |
| Trip11 | 1.14217492 |
| Fam188a | 1.142004601 |
| Eef1g | 1.141937714 |
| Hexa | 1.141788343 |
| Birc6 | 1.14105696 |
| Ralgapb | 1.141044122 |
| L3mbtl4 | 1.140927279 |
| Hist1h2bj | 1.139793682 |
| Nmt1 | 1.139681043 |
| Pfdn2 | 1.139656623 |
| Iqgap3 | 1.139576953 |
| Blcap | 1.139350074 |
| Gpr135 | 1.139197307 |
| Rps27l | 1.139018596 |
| Dclk2 | 1.138982279 |
| Sec16b | 1.138876121 |
| Kdm3a | 1.138570535 |
| Tram1 | 1.138433064 |
| Fanca | 1.138123477 |
| Anxa6 | 1.137245054 |
| Rnf219 | 1.136864469 |
| Atr | 1.136634424 |
| Mgll | 1.136392879 |
| Pctp | 1.136275988 |
| Golga7 | 1.135977576 |
| Mki67 | 1.135763012 |
| Cdc40 | 1.135575431 |
| N4bp2l1 | 1.135122604 |
| Helz2 | 1.13448021 |
| Moap1 | 1.134457397 |
| Inpp4a | 1.13422231 |
| Filip1 | 1.13412719 |
| Kit | 1.133743216 |
| Vstm4 | 1.133682312 |
| Tbc1d2b | 1.133554663 |
| Spats2l | 1.133474009 |
| Iqsec3 | 1.13346803 |
| Pwp1 | 1.133171504 |
| Mpnd | 1.132904409 |
| Enox2 | 1.132130253 |
| Stc2 | 1.131905934 |
| Tmem8b | 1.131864029 |
| Faf1 | 1.131775199 |

| | |
|---|---|
| Gprin3 | 1.131685386 |
| Lars2 | 1.131182125 |
| Rbfa | 1.131018868 |
| Gpatch2l | 1.13090552 |
| Gabrp | 1.13044069 |
| Kcnq1 | 1.130109385 |
| Morf4l1 | 1.129524676 |
| Loxl4 | 1.128926412 |
| Tctex1d2 | 1.128801879 |
| Entpd6 | 1.128507749 |
| Srrm1 | 1.128342849 |
| Magi1 | 1.127882292 |
| Ikzf5 | 1.127615894 |
| Slc16a11 | 1.127035625 |
| Tmed1 | 1.127023131 |
| Hmgn5 | 1.126894066 |
| Rad1 | 1.126854535 |
| Hsd17b7 | 1.125979211 |
| Cux1 | 1.125973173 |
| Sap18 | 1.125960166 |
| Erf | 1.125462561 |
| Cycs | 1.124925932 |
| Kctd17 | 1.124439274 |
| Tspan18 | 1.124393558 |
| Cdc26 | 1.124240615 |
| Neto2 | 1.124235391 |
| Wdr1 | 1.123812443 |
| Bcat1 | 1.122994522 |
| Cnot3 | 1.12276555 |
| Hoxb6 | 1.122716952 |
| Atl1 | 1.122662281 |
| Btg3 | 1.121911225 |
| Tmf1 | 1.121743434 |
| Tmem185b | 1.121664 |
| Rnf182 | 1.121547763 |
| Snai1 | 1.121523383 |
| Akirin2 | 1.121423562 |
| Rabac1 | 1.121396614 |
| Ivns1abp | 1.120797191 |
| Arcn1 | 1.120533123 |
| Atp6v1h | 1.120161065 |
| Bphl | 1.119496972 |
| Magohb | 1.119287761 |
| Hapln1 | 1.119266322 |
| Pus3 | 1.119037119 |
| Zbtb8os | 1.118971365 |
| Dapk3 | 1.118919092 |

| | |
|---|---|
| Pias4 | 1.118902673 |
| Atp5c1 | 1.118598567 |
| Bbs1 | 1.118066839 |
| Kctd5 | 1.117714059 |
| Mrpl41 | 1.117654783 |
| Fzd3 | 1.117417083 |
| Cxcl11 | 1.11738871 |
| Gpr108 | 1.117003589 |
| Hs3st1 | 1.11669308 |
| Tcf12 | 1.11667295 |
| Cdr2l | 1.11643521 |
| Pdp1 | 1.115837836 |
| Cpm | 1.115134373 |
| Higd2a | 1.114900485 |
| Slc25a10 | 1.114760904 |
| Maoa | 1.114402687 |
| Tcta | 1.11369014 |
| Nek4 | 1.113164754 |
| Rap2c | 1.112961046 |
| Mdh1b | 1.112702663 |
| Atf6 | 1.112507595 |
| Rel | 1.111906715 |
| Ubxn1 | 1.111902118 |
| Ttc9 | 1.111808123 |
| Cystm1 | 1.111585898 |
| Lbx2 | 1.111547739 |
| Etfa | 1.111149255 |
| Katnbl1 | 1.110433784 |
| Aldh6a1 | 1.11032203 |
| Immp2l | 1.109358043 |
| Tiparp | 1.109265636 |
| Slc38a2 | 1.109036609 |
| Hdlbp | 1.108869158 |
| Spock2 | 1.108749757 |
| Mga | 1.107994791 |
| Smpdl3a | 1.107867604 |
| Lamtor2 | 1.107615779 |
| Tssc4 | 1.107391502 |
| Nap1l3 | 1.107378439 |
| Ptpn23 | 1.107346818 |
| Frmd4a | 1.106670854 |
| Gstp1 | 1.106591068 |
| Snrnp35 | 1.106490773 |
| Antxr1 | 1.106488548 |
| Gprc5b | 1.106324943 |
| Emp1 | 1.106250656 |
| Arhgap44 | 1.106204144 |

| | |
|---|---|
| Bmf | 1.106189485 |
| Perp | 1.106144329 |
| Nedd8 | 1.105809459 |
| Bmper | 1.105422511 |
| Aptx | 1.103784347 |
| Vps33b | 1.103448812 |
| Casp2 | 1.103218658 |
| Lrrc41 | 1.10274207 |
| Birc5 | 1.102623029 |
| Mthfs | 1.102513544 |
| Lamp1 | 1.102425063 |
| Il17re | 1.102360493 |
| Psma6 | 1.102265052 |
| Nthl1 | 1.101821393 |
| Dip2a | 1.101736863 |
| Gtf2h5 | 1.101653768 |
| Ipo8 | 1.101635566 |
| Ptgir | 1.101632795 |
| Bnip3 | 1.100339709 |
| Mrpl45 | 1.10027239 |
| Aprt | 1.099978132 |
| Gatad1 | 1.099739546 |
| Os9 | 1.099589302 |
| Galnt6 | 1.099314101 |
| Itga2 | 1.099104608 |
| Swsap1 | 1.09886748 |
| Casp4 | 1.098775038 |
| Ranbp1 | 1.098726755 |
| Emc4 | 1.097905028 |
| Hmbox1 | 1.097176088 |
| Nudt13 | 1.096487099 |
| Erc1 | 1.096443158 |
| 42436 | 1.09629446 |
| Chrne | 1.096197183 |
| Efhd1 | 1.096153751 |
| Hdac10 | 1.095996262 |
| Usp47 | 1.095804364 |
| Mon2 | 1.09547802 |
| Prpf31 | 1.095281316 |
| Dis3l | 1.095256873 |
| Dgkq | 1.095220934 |
| Fbxo6 | 1.095128311 |
| P2ry2 | 1.095020388 |
| Ttf1 | 1.094934231 |
| Rnd1 | 1.094782816 |
| Taf5 | 1.094137344 |
| Slc17a5 | 1.093933108 |

| | |
|---|---|
| Stx16 | 1.093856152 |
| Eif4ebp2 | 1.093848213 |
| Lgals9 | 1.093657578 |
| Med27 | 1.093564222 |
| Galnt11 | 1.093484414 |
| Nrgn | 1.093336354 |
| Ddx11 | 1.093136279 |
| Synrg | 1.093088719 |
| Fam120c | 1.092361425 |
| Satb2 | 1.092270127 |
| Smad2 | 1.092257855 |
| Ube2z | 1.091998241 |
| Mark3 | 1.091883731 |
| Akt1 | 1.091257517 |
| Tmem44 | 1.0910874 |
| Adal | 1.090619987 |
| Pvrl1 | 1.090507551 |
| Depdc5 | 1.090165873 |
| Gnb1 | 1.089733743 |
| Spred3 | 1.089415744 |
| Ralb | 1.089145297 |
| Sumo2 | 1.08850309 |
| Fnip2 | 1.088355039 |
| Qpctl | 1.088342228 |
| Csrp1 | 1.088215625 |
| Gng12 | 1.088059122 |
| Mrpl4 | 1.087638063 |
| Cd99l2 | 1.087353574 |
| Ptgr2 | 1.087264902 |
| Otogl | 1.086785529 |
| Usp30 | 1.086141939 |
| Irf3 | 1.085987316 |
| Btf3 | 1.08584546 |
| Nde1 | 1.085774468 |
| Fam131a | 1.08565918 |
| Gsr | 1.085622969 |
| Taz | 1.085616395 |
| Gars | 1.085196369 |
| Spry2 | 1.085018136 |
| Lmna | 1.085006535 |
| Cmpk2 | 1.085004372 |
| Mtfmt | 1.084891009 |
| Wbscr27 | 1.084794703 |
| Krt15 | 1.084773688 |
| Psmb3 | 1.084252478 |
| Mfi2 | 1.083288121 |
| Ropn1l | 1.082883532 |

| | |
|---|---|
| Fam117a | 1.082813859 |
| Fbxo36 | 1.082748299 |
| Prrt1 | 1.082717902 |
| Lpp | 1.082635298 |
| Ppt2 | 1.082578681 |
| Col1a2 | 1.082550212 |
| Rbbp8 | 1.081842177 |
| Capn5 | 1.081010157 |
| Rps13 | 1.08037986 |
| Klhl3 | 1.080134338 |
| Tpbg | 1.080054205 |
| Nacc2 | 1.07960464 |
| Arhgap29 | 1.079435716 |
| Pgbd1 | 1.078137315 |
| Rnf4 | 1.077420956 |
| Snrpb | 1.077311621 |
| Galt | 1.076849853 |
| Bahd1 | 1.076331239 |
| Nrm | 1.07585238 |
| Ptpn12 | 1.075409915 |
| Kif14 | 1.075314171 |
| Tmx2 | 1.075204292 |
| Irf9 | 1.075202369 |
| Spata18 | 1.074338293 |
| Dbn1 | 1.074153758 |
| Piezo1 | 1.073892304 |
| Gdf15 | 1.073751088 |
| Tars2 | 1.073488109 |
| Ctdspl2 | 1.07345308 |
| Miip | 1.073113682 |
| Lrrk1 | 1.072934264 |
| Sh2d4a | 1.072857935 |
| Prkacb | 1.072753113 |
| Psmg3 | 1.071262038 |
| Vwa9 | 1.071031898 |
| Hoxa4 | 1.070802802 |
| Rev3l | 1.070706586 |
| Aamdc | 1.070503811 |
| Slc25a16 | 1.069843074 |
| Cyp2u1 | 1.069684698 |
| Cox6b1 | 1.06914661 |
| Snx22 | 1.068823517 |
| Slc9a3r1 | 1.068747338 |
| Strip2 | 1.068703046 |
| Chsy1 | 1.068158137 |
| Traf3ip1 | 1.067979525 |
| Wasl | 1.067628754 |

| | |
|---|---|
| Fuk | 1.067626654 |
| Npr3 | 1.067454012 |
| Arl14epl | 1.067103442 |
| Il17d | 1.067037758 |
| Amfr | 1.066961597 |
| Rpl39l | 1.066889375 |
| Hmgb3 | 1.06687521 |
| Kdm4a | 1.06656076 |
| Kif26b | 1.065855815 |
| Elmod1 | 1.065595524 |
| Ociad1 | 1.065510792 |
| Bmi1 | 1.064742148 |
| Bnc1 | 1.064658056 |
| Tnrc6b | 1.064461773 |
| Nubpl | 1.064269882 |
| Cdcp1 | 1.064169503 |
| Fosl2 | 1.064157831 |
| Pcbp1 | 1.064120117 |
| Sema4g | 1.063043477 |
| Msantd4 | 1.062121956 |
| Acbd7 | 1.061997649 |
| Tubgcp3 | 1.061315523 |
| Pcgf3 | 1.060414986 |
| Slc15a4 | 1.060392628 |
| Emc2 | 1.059851509 |
| Casp7 | 1.059498131 |
| Lrp5 | 1.059442815 |
| Ddhd2 | 1.059413719 |
| Gxylt2 | 1.059334412 |
| Myeov2 | 1.05932644 |
| Ppan | 1.058713125 |
| Cnot10 | 1.0583758 |
| Hiat1 | 1.058093293 |
| Slc9a7 | 1.057678835 |
| Rpl19 | 1.057468401 |
| Copa | 1.057179606 |
| Trappc1 | 1.056848092 |
| Psme3 | 1.056626622 |
| Slc9a3r2 | 1.056535188 |
| Igfbp7 | 1.056483139 |
| Rab27b | 1.056090722 |
| Lcmt2 | 1.055538918 |
| Zmiz1 | 1.054620201 |
| Rnf41 | 1.054605412 |
| Tmsb15a | 1.054263247 |
| Timm10 | 1.054204411 |
| Wdpcp | 1.053394729 |

| | |
|---|---|
| Metrn | 1.053310997 |
| Itsn1 | 1.053077781 |
| Kxd1 | 1.053077143 |
| Pinlyp | 1.052902525 |
| Fgfr1 | 1.05266924 |
| Rusc1 | 1.052664064 |
| Zeb1 | 1.05223567 |
| Yeats2 | 1.052146078 |
| Acot8 | 1.051889281 |
| Bag4 | 1.051581802 |
| Spopl | 1.051476529 |
| Abcc10 | 1.051288759 |
| Ldb1 | 1.050606777 |
| Npat | 1.050540151 |
| Efemp1 | 1.05034253 |
| Srebf2 | 1.049927826 |
| Fhod1 | 1.04985621 |
| Ecd | 1.04907122 |
| Fahd2a | 1.048805 |
| Psmb4 | 1.048205715 |
| Ogfod1 | 1.047725397 |
| Adamts6 | 1.04738396 |
| Adprm | 1.047300643 |
| Hhex | 1.047296387 |
| Snx32 | 1.047177993 |
| Efcab2 | 1.047094699 |
| Ubl7 | 1.046826662 |
| Jade1 | 1.046515244 |
| Smpdl3b | 1.046509364 |
| Prickle3 | 1.046293488 |
| Ppwd1 | 1.04591496 |
| Sec23b | 1.045863706 |
| Prkab1 | 1.045743878 |
| Cyb5rl | 1.045360075 |
| Eif3l | 1.045308643 |
| Sox15 | 1.045035857 |
| Dcn | 1.044989729 |
| Slc25a34 | 1.044892785 |
| Stxbp1 | 1.044406509 |
| Bhlhb9 | 1.043980892 |
| Plp2 | 1.043791236 |
| Casc4 | 1.043679414 |
| Cbr3 | 1.043330479 |
| Npbwr1 | 1.04287292 |
| Ctsh | 1.042580903 |
| St7 | 1.042473055 |
| Bfar | 1.042406527 |

| | |
|---|---|
| Gramd1a | 1.042155543 |
| Imp4 | 1.042155257 |
| Ptrh1 | 1.041924812 |
| Urm1 | 1.041018251 |
| Cltb | 1.040588298 |
| Gnb3 | 1.040023769 |
| Ncapg2 | 1.039929305 |
| Tpd52l2 | 1.039620935 |
| Dld | 1.039271317 |
| Ctdnep1 | 1.039171244 |
| Ogdh | 1.038932933 |
| Slc52a2 | 1.038883042 |
| Scube1 | 1.038845268 |
| Tmem140 | 1.038040526 |
| Rbm5 | 1.038030795 |
| Cdk16 | 1.037986927 |
| Rabif | 1.037928153 |
| Dnm1 | 1.037866203 |
| Tns1 | 1.037831744 |
| Hlcs | 1.037400634 |
| Caskin2 | 1.036700264 |
| Pop1 | 1.036458781 |
| Ccnl1 | 1.036403719 |
| Dlgap5 | 1.03631168 |
| Coil | 1.036284762 |
| Chmp2b | 1.03591254 |
| Rock2 | 1.03565121 |
| Agfg2 | 1.034916412 |
| Il13ra1 | 1.034766183 |
| Alg9 | 1.034500852 |
| Aqp1 | 1.034339727 |
| Nabp2 | 1.034227863 |
| Nfu1 | 1.034118889 |
| Trim44 | 1.033840501 |
| Dcbld1 | 1.033302714 |
| Bnc2 | 1.033270118 |
| Notch1 | 1.033251813 |
| Katnal1 | 1.033001399 |
| Fam89a | 1.032739339 |
| Phyhip | 1.032558449 |
| Als2 | 1.032545562 |
| Afap1l2 | 1.032041846 |
| Nipa2 | 1.031834726 |
| Nfya | 1.031655176 |
| Tpmt | 1.031548232 |
| Rprd1a | 1.03120849 |
| Ahsa1 | 1.031043979 |

| | |
|---|---|
| Osbpl7 | 1.030960229 |
| Pik3cd | 1.030131912 |
| Anxa2 | 1.029394122 |
| Wdr34 | 1.028714161 |
| Zswim1 | 1.028510135 |
| Pycrl | 1.028509669 |
| Plekha1 | 1.028255364 |
| Fzd9 | 1.028140733 |
| Slc24a1 | 1.027730033 |
| Rbbp7 | 1.027495297 |
| Ttc27 | 1.027234687 |
| Ptgfrn | 1.027229401 |
| Ptbp3 | 1.026289103 |
| Slc27a5 | 1.025942395 |
| Zbtb49 | 1.025941519 |
| Pih1d1 | 1.02571742 |
| Pusl1 | 1.025472049 |
| Ift172 | 1.025399565 |
| Rbm27 | 1.023575258 |
| Angpt1 | 1.023335628 |
| Sdsl | 1.022989084 |
| Stk40 | 1.022782518 |
| Dnajc1 | 1.022337613 |
| Tbc1d31 | 1.022055468 |
| Fam73b | 1.022013715 |
| Nxph3 | 1.021836573 |
| Mettl5 | 1.021506843 |
| Erap1 | 1.02058993 |
| Erp27 | 1.020575708 |
| Pcid2 | 1.020176417 |
| Dennd2d | 1.019341453 |
| Rint1 | 1.019208272 |
| Dnajc28 | 1.019033162 |
| Lrp12 | 1.018871515 |
| Ppm1k | 1.018757829 |
| Bet1 | 1.018569172 |
| Arhgap5 | 1.018075263 |
| Sphk1 | 1.017475368 |
| Cntnap3 | 1.017334189 |
| Det1 | 1.017171754 |
| Fam107b | 1.016900038 |
| G6pc3 | 1.016772472 |
| Slc38a4 | 1.016472781 |
| Rbm45 | 1.016394966 |
| Rab5a | 1.016046819 |
| Rnf141 | 1.01580652 |
| Foxa1 | 1.015471976 |

| | |
|---|---|
| Mysm1 | 1.015358772 |
| Ndufaf1 | 1.015234522 |
| St6galnac1 | 1.01516133 |
| Dusp3 | 1.015019673 |
| Med10 | 1.014402824 |
| Znrf1 | 1.014131445 |
| Selm | 1.013551408 |
| Rnft1 | 1.013046641 |
| Slc27a4 | 1.012859754 |
| Slc44a2 | 1.012835882 |
| Edn1 | 1.012315503 |
| Mmadhc | 1.012144559 |
| Rbl1 | 1.012119289 |
| Fbp1 | 1.012010387 |
| Cyb561d2 | 1.01198802 |
| Acss1 | 1.011720993 |
| Fam129b | 1.011507869 |
| Fbxo27 | 1.011432216 |
| Cenpa | 1.011005206 |
| Agl | 1.010955362 |
| Ccdc61 | 1.010587481 |
| Tmub2 | 1.010456656 |
| Gng11 | 1.009933168 |
| Eps15l1 | 1.009727933 |
| Sh3bp2 | 1.009713692 |
| Brk1 | 1.009665339 |
| Plod2 | 1.008960104 |
| Cc2d1b | 1.008743107 |
| Hnrnpm | 1.008352008 |
| Kctd13 | 1.00830205 |
| Ankrd13a | 1.008296788 |
| Lamb3 | 1.007921566 |
| Dnajb9 | 1.007448946 |
| Dnttip2 | 1.007411945 |
| Map1s | 1.006985614 |
| Enpp1 | 1.006667119 |
| Alkbh5 | 1.006314118 |
| Lsm2 | 1.005355348 |
| Tril | 1.004544875 |
| Snx33 | 1.004288758 |
| Specc1 | 1.004203427 |
| Gaa | 1.004129433 |
| Adck5 | 1.003880297 |
| Dapk1 | 1.003748688 |
| Stag1 | 1.003680467 |
| Mcmbp | 1.00307582 |
| Ccnb1 | 1.002562705 |

| | |
|---|---|
| Mettl17 | 1.002435341 |
| Pmepa1 | 1.002303327 |
| Mgmt | 1.002200092 |
| Nucks1 | 1.002018269 |
| Letm2 | 1.001749818 |
| Ube2b | 1.000688155 |
| Heatr1 | 0.999619146 |
| Hbegf | 0.999260842 |
| Pigw | 0.999098113 |
| Tubgcp4 | 0.998947968 |
| Vezt | 0.998883501 |
| Atp5g2 | 0.998610508 |
| Pik3ca | 0.998355815 |
| Hs2st1 | 0.997863347 |
| Dcun1d4 | 0.997730195 |
| Klf3 | 0.997092077 |
| Srp14 | 0.99697002 |
| Csnk2b | 0.996899995 |
| Ovgp1 | 0.996422333 |
| Mamld1 | 0.996170663 |
| Col23a1 | 0.995761925 |
| Psmc1 | 0.995402226 |
| Ywhaq | 0.99539428 |
| Ctsd | 0.995366339 |
| Ccar1 | 0.995347678 |
| Arhgap20 | 0.995234577 |
| Trmt12 | 0.995164857 |
| Rrm2 | 0.994969785 |
| Hax1 | 0.994710286 |
| Man2a2 | 0.99455453 |
| Cyp20a1 | 0.993253147 |
| S100a10 | 0.993128779 |
| Dph6 | 0.992408797 |
| Smap1 | 0.992261452 |
| Sh3tc2 | 0.992114527 |
| Txlng | 0.99201471 |
| Tmem214 | 0.991759006 |
| Zfp62 | 0.991731554 |
| Rnasek | 0.991464819 |
| Srsf5 | 0.991391167 |
| Masp1 | 0.991295028 |
| Armcx6 | 0.991264495 |
| Ufc1 | 0.990668074 |
| Tia1 | 0.990137074 |
| Ccdc82 | 0.990114559 |
| Cited4 | 0.989975041 |
| Gas6 | 0.989655702 |

| | |
|---|---|
| Cox7b | 0.989594889 |
| Mrpl49 | 0.989438968 |
| Necab1 | 0.989261548 |
| Pitrm1 | 0.989235793 |
| Ncf2 | 0.988538893 |
| Pecr | 0.988045359 |
| Trim39 | 0.98719264 |
| Ngef | 0.986708895 |
| Gemin2 | 0.986149787 |
| Pdk2 | 0.986073852 |
| Zmpste24 | 0.986034263 |
| Zcchc8 | 0.986016701 |
| Slc39a8 | 0.985350425 |
| Upf3b | 0.985343156 |
| Tada2b | 0.984984773 |
| Ninj1 | 0.984929106 |
| Sncg | 0.984117735 |
| Chd1 | 0.983860204 |
| H2afx | 0.982623729 |
| Slc25a11 | 0.981912206 |
| Trmt10c | 0.981773782 |
| Mmp11 | 0.981695718 |
| Swap70 | 0.981180834 |
| Nkapl | 0.981141155 |
| Anapc10 | 0.980687557 |
| Zbtb24 | 0.980684076 |
| Chd8 | 0.980656363 |
| Synm | 0.980549072 |
| Gab2 | 0.97952823 |
| Sorbs2 | 0.979452392 |
| Phf6 | 0.978974467 |
| Gatm | 0.978862297 |
| Agpat5 | 0.978687418 |
| Edc3 | 0.978651841 |
| Commd2 | 0.978360215 |
| Rpl21 | 0.978051754 |
| Ghr | 0.977783774 |
| Nub1 | 0.977317863 |
| Drp2 | 0.976864212 |
| Rab3b | 0.976862055 |
| Zc3h15 | 0.975920482 |
| Pbrm1 | 0.975683793 |
| Mrrf | 0.975349268 |
| Mad2l1bp | 0.975132574 |
| Acaca | 0.974983869 |
| Dap | 0.974817371 |
| Rps20 | 0.974648539 |

| | |
|---|---|
| Abca7 | 0.974164913 |
| Tbc1d1 | 0.974111938 |
| Pik3c3 | 0.974102185 |
| Leo1 | 0.973931098 |
| Kifc3 | 0.973785497 |
| Akt2 | 0.973320338 |
| Ppl | 0.973305935 |
| Hnrnpc | 0.972833382 |
| Cmtm8 | 0.972684713 |
| Ccdc181 | 0.972349475 |
| Churc1 | 0.972103498 |
| Gpr4 | 0.971915095 |
| Pclo | 0.971796497 |
| Fads3 | 0.971708928 |
| Mon1b | 0.971488288 |
| Pcf11 | 0.971251191 |
| Mgst2 | 0.971193382 |
| Astn2 | 0.970963862 |
| Dclre1a | 0.970897688 |
| Irf2 | 0.970858297 |
| Nol6 | 0.970794331 |
| Slc4a5 | 0.970379188 |
| Parm1 | 0.969664143 |
| Aga | 0.969529534 |
| B3gnt2 | 0.969436511 |
| Ndufb7 | 0.968978702 |
| Fbxo25 | 0.968697974 |
| Cthrc1 | 0.968396921 |
| Henmt1 | 0.9682512 |
| Rapgef4 | 0.967712206 |
| Ptx3 | 0.967615229 |
| Btbd2 | 0.967571758 |
| Atp13a3 | 0.967417178 |
| Pcbp4 | 0.967409634 |
| Nedd1 | 0.967091452 |
| Ddx47 | 0.966758936 |
| Cit | 0.966645891 |
| Rbm24 | 0.966562012 |
| Ube3a | 0.966338516 |
| Trappc10 | 0.96624649 |
| Ttc33 | 0.966188313 |
| Cldn15 | 0.965244688 |
| Fuom | 0.96476782 |
| Slc4a1ap | 0.964511707 |
| Bcar1 | 0.964404543 |
| Dnah17 | 0.963900964 |
| Copg1 | 0.963767121 |

| | |
|---|---|
| Fam46c | 0.963687516 |
| Hspa5 | 0.963525359 |
| Rnf14 | 0.962991521 |
| Tfap4 | 0.961854773 |
| Tspan15 | 0.961634843 |
| Samd4b | 0.961084933 |
| Lama2 | 0.959898877 |
| Tmed7 | 0.959585158 |
| Cox7a2l | 0.95958486 |
| Tm2d1 | 0.958925043 |
| Cpsf3 | 0.9587638 |
| Mtmr2 | 0.958713741 |
| Atl3 | 0.958453059 |
| Fkbp15 | 0.958024012 |
| Rnf166 | 0.957762368 |
| Hras | 0.957435061 |
| Popdc3 | 0.957399502 |
| Fastkd2 | 0.957200113 |
| Grk6 | 0.956665974 |
| Map7d1 | 0.955775654 |
| Nkap | 0.955724405 |
| Prkaca | 0.955616632 |
| Ebf4 | 0.955142087 |
| Pde8b | 0.955094629 |
| Gtf3c1 | 0.954609552 |
| Cyp27a1 | 0.954526382 |
| Znrf2 | 0.954213557 |
| Klhl5 | 0.953901189 |
| Cbr1 | 0.953807223 |
| Bcl6b | 0.953559872 |
| Paxip1 | 0.953382521 |
| Robo1 | 0.953356164 |
| Znrf3 | 0.952854106 |
| Atxn1l | 0.952847312 |
| Itpkc | 0.952609936 |
| Actr3 | 0.952569528 |
| Mast3 | 0.951984892 |
| Ablim1 | 0.951605314 |
| Stk33 | 0.950654645 |
| Anxa9 | 0.950416905 |
| Rrp15 | 0.950084868 |
| Dock8 | 0.950012078 |
| Yipf5 | 0.949714048 |
| Ttll12 | 0.949643098 |
| Hint2 | 0.949573581 |
| Klf2 | 0.949566124 |
| Fbl | 0.949166632 |

| | |
|---|---|
| Samsn1 | 0.948879961 |
| Bbs5 | 0.948569172 |
| Tmem154 | 0.948350157 |
| Fgfr2 | 0.948247927 |
| Ppp1r32 | 0.948223194 |
| Oplah | 0.946917208 |
| Gmcl1 | 0.946908002 |
| Wdr5 | 0.946671882 |
| Ebf1 | 0.94611396 |
| Rnpep | 0.946081078 |
| Bcl7c | 0.945641139 |
| Rad23a | 0.945426728 |
| Elovl4 | 0.944844906 |
| Chadl | 0.944404979 |
| Samd5 | 0.944179207 |
| Mblac1 | 0.944172573 |
| Stx18 | 0.943869567 |
| Fyttd1 | 0.943537773 |
| Tagln2 | 0.94307768 |
| Llph | 0.942979367 |
| Dusp6 | 0.942898969 |
| Mtch1 | 0.942665225 |
| Ypel3 | 0.942327376 |
| Asap3 | 0.941922164 |
| Trmt2b | 0.941795942 |
| Uqcrb | 0.941501358 |
| Ola1 | 0.941133761 |
| Slk | 0.940779815 |
| Ccno | 0.940691615 |
| Ctnna3 | 0.940635391 |
| Pde4dip | 0.940512336 |
| Cdh6 | 0.940322771 |
| Nr4a3 | 0.940308745 |
| Ndel1 | 0.940218255 |
| Sntb1 | 0.940109707 |
| Nav2 | 0.939604712 |
| Kif20b | 0.939547538 |
| Tank | 0.939409424 |
| Ccdc12 | 0.939188072 |
| Rasa3 | 0.93914837 |
| Tmem70 | 0.939038577 |
| Psme1 | 0.938602494 |
| Mid2 | 0.938520019 |
| Itgb1 | 0.938491158 |
| Ambra1 | 0.938416338 |
| Ubn1 | 0.937689352 |
| Ncln | 0.937474777 |

| | |
|---|---|
| Uap1l1 | 0.937351573 |
| Aff4 | 0.936882646 |
| Dph5 | 0.935875969 |
| Flt1 | 0.935590106 |
| E2f4 | 0.935543401 |
| Inca1 | 0.935475756 |
| Ocel1 | 0.935274858 |
| Alkbh3 | 0.935267419 |
| Mylip | 0.934568176 |
| Hnrnpul2 | 0.934466841 |
| Cib2 | 0.934247358 |
| Slc18a2 | 0.93418901 |
| Mob2 | 0.934152715 |
| Top1mt | 0.93363443 |
| Shmt1 | 0.933440301 |
| Iqcg | 0.933115213 |
| Magee1 | 0.932817931 |
| Snrpa | 0.932801566 |
| Sumf1 | 0.932567613 |
| Ick | 0.931109442 |
| Eif4e | 0.93106957 |
| Rarres1 | 0.930538814 |
| Mink1 | 0.92984352 |
| Ubiad1 | 0.929766885 |
| Mcm5 | 0.929249617 |
| Zfand1 | 0.928485183 |
| Aff3 | 0.927497649 |
| Fbxo32 | 0.927391503 |
| Cacna2d1 | 0.927306502 |
| Actr6 | 0.927196372 |
| Gtf2f2 | 0.927062639 |
| Etfb | 0.926982696 |
| Mettl25 | 0.926978323 |
| Tspan5 | 0.926725789 |
| Tti2 | 0.926703794 |
| Nhs | 0.926666412 |
| Glod4 | 0.926636214 |
| Spc25 | 0.926036424 |
| Palmd | 0.926034819 |
| Atp11c | 0.925992622 |
| Tnfsf13 | 0.92594897 |
| Snap29 | 0.925910277 |
| Trim21 | 0.925637863 |
| Ctxn1 | 0.925621536 |
| Ythdf3 | 0.924771778 |
| Tsku | 0.923899483 |
| Dhrs7b | 0.92383716 |

| | |
|---|---|
| Tmem216 | 0.923673605 |
| Vps37c | 0.923366875 |
| St6gal1 | 0.923241022 |
| Tmem110 | 0.922851976 |
| Cmbl | 0.922742861 |
| Tmem222 | 0.922562492 |
| Pvrl2 | 0.922454242 |
| Chic2 | 0.922197871 |
| Arrb1 | 0.922042067 |
| Fbxw8 | 0.921980397 |
| Rasl11a | 0.921976968 |
| Chd6 | 0.92160675 |
| Csgalnact2 | 0.921039932 |
| Snap23 | 0.921002611 |
| Map2k4 | 0.920919945 |
| Taf9 | 0.920737504 |
| Zbtb33 | 0.920178337 |
| Dnajc11 | 0.919527759 |
| Aebp1 | 0.919424757 |
| Ubap2 | 0.91883064 |
| Tex264 | 0.918541708 |
| Cnep1r1 | 0.91841202 |
| Cebpa | 0.918250805 |
| Psmb10 | 0.918239991 |
| Ell2 | 0.918041806 |
| Asb7 | 0.917785669 |
| Dusp28 | 0.91771442 |
| Emc8 | 0.91761681 |
| Dennd4a | 0.917045447 |
| Mta3 | 0.916906926 |
| Obsl1 | 0.916878175 |
| Ppfia1 | 0.916817087 |
| Rbmx2 | 0.91658772 |
| Exog | 0.916245307 |
| Tmem184b | 0.916228105 |
| Coq7 | 0.915743649 |
| Heyl | 0.915670122 |
| Sult1b1 | 0.915124722 |
| Htr1d | 0.91508159 |
| Kat8 | 0.914755783 |
| Rgs3 | 0.914607861 |
| Fbxo21 | 0.914378009 |
| Gipc1 | 0.9130547 |
| Sema4d | 0.913048834 |
| Arhgap4 | 0.912790691 |
| Usp11 | 0.912640608 |
| Fmnl1 | 0.912519772 |

| | |
|---|---|
| Zfhx3 | 0.912421529 |
| Reep1 | 0.912213245 |
| Lactb | 0.911804873 |
| Olfml2a | 0.911347473 |
| Wdr11 | 0.911086142 |
| Vps29 | 0.911055394 |
| Ptrf | 0.910995392 |
| S100pbp | 0.910901059 |
| Aspscr1 | 0.910609963 |
| Prpsap2 | 0.910278623 |
| Npm1 | 0.910259159 |
| Keap1 | 0.909671609 |
| Plcxd2 | 0.909115514 |
| Drg2 | 0.908514261 |
| Arhgef28 | 0.908431781 |
| Nus1 | 0.908009203 |
| Bnip3l | 0.907885437 |
| Zdhhc13 | 0.906853732 |
| Incenp | 0.906789977 |
| Trappc13 | 0.906679211 |
| Prkg1 | 0.906158327 |
| Tor1a | 0.906028844 |
| Nbl1 | 0.905822188 |
| Exoc6b | 0.905321898 |
| Slc6a15 | 0.905006056 |
| Rps23 | 0.904789327 |
| Ect2 | 0.904694477 |
| Gpr155 | 0.904474926 |
| Robo3 | 0.904128063 |
| Dchs1 | 0.903715619 |
| Plekhg4 | 0.903188791 |
| Trmt10a | 0.903162472 |
| Fn3k | 0.903149847 |
| Lcmt1 | 0.903136807 |
| Unc13b | 0.902953892 |
| Dda1 | 0.902773846 |
| Arhgap1 | 0.901956495 |
| Srsf12 | 0.901600876 |
| Zfp1 | 0.901397173 |
| Cluap1 | 0.901333906 |
| Champ1 | 0.900972763 |
| Ints2 | 0.900587832 |
| Hgs | 0.900505895 |
| Chfr | 0.900367059 |
| Rin2 | 0.899511342 |
| Heatr6 | 0.898997777 |
| Nol10 | 0.898800763 |

| | |
|---|---|
| Cdc45 | 0.898756054 |
| Fzd8 | 0.898708346 |
| Map3k14 | 0.898442148 |
| Prrc2b | 0.898233242 |
| Sirt2 | 0.898023025 |
| 42438 | 0.897863088 |
| Dnaaf3 | 0.897242855 |
| Ciapin1 | 0.897145858 |
| Arhgef1 | 0.897039023 |
| Zmym6 | 0.89683695 |
| Mbd5 | 0.896757431 |
| Tmed6 | 0.896678275 |
| Twistnb | 0.8965831 |
| Foxo4 | 0.896068454 |
| Pacs1 | 0.895586886 |
| Gsdmd | 0.895461051 |
| Arap1 | 0.89488509 |
| Amacr | 0.893380765 |
| Fmn1 | 0.893035284 |
| Tmem17 | 0.8930084 |
| Gpd1l | 0.89280954 |
| Phlpp1 | 0.892220536 |
| Rad17 | 0.892102456 |
| Ano7 | 0.891817259 |
| Jmjd8 | 0.891783291 |
| Ephx1 | 0.891559098 |
| Flt4 | 0.891231237 |
| B3gnt8 | 0.891093985 |
| Nhp2 | 0.891057426 |
| Nrn1 | 0.890913928 |
| Abcc2 | 0.890768569 |
| Man2b1 | 0.890724347 |
| Snx10 | 0.890561542 |
| Mrpl1 | 0.890557907 |
| Glce | 0.890197653 |
| Dnajc6 | 0.890159645 |
| Ube2e1 | 0.890088175 |
| Trpv4 | 0.888997631 |
| Agk | 0.888881694 |
| Exo1 | 0.888390459 |
| Tor2a | 0.888265645 |
| Nipsnap3a | 0.888183124 |
| Sidt2 | 0.888090481 |
| Hapln3 | 0.886901849 |
| Fam228b | 0.886874228 |
| Zfat | 0.886607376 |
| Fam171a1 | 0.885943006 |

| | |
|---|---|
| Fam195a | 0.885341839 |
| Plekha5 | 0.885215972 |
| Sp8 | 0.88514781 |
| Dram2 | 0.885006736 |
| Cep55 | 0.884607469 |
| Slc2a3 | 0.884410634 |
| Soga3 | 0.883907809 |
| Slc14a1 | 0.883858513 |
| Pdcd2l | 0.883635299 |
| Cenpo | 0.883436363 |
| Mrps17 | 0.882238801 |
| Ciao1 | 0.882087864 |
| Cd82 | 0.881935793 |
| E2f3 | 0.88186123 |
| Surf6 | 0.881796561 |
| Pbxip1 | 0.88155735 |
| Rbp1 | 0.880205978 |
| Frg1 | 0.880163499 |
| Ddx31 | 0.879994407 |
| Mfsd5 | 0.879653549 |
| Lman2 | 0.87924501 |
| Cdk2 | 0.878489884 |
| Pnmal1 | 0.878346308 |
| Ltv1 | 0.877975545 |
| Trim68 | 0.87778189 |
| Il31ra | 0.877596397 |
| Nfatc1 | 0.877457913 |
| Paqr7 | 0.877209371 |
| Dync1li1 | 0.876933137 |
| Nek7 | 0.876689347 |
| Arhgap23 | 0.87637353 |
| Anln | 0.876371011 |
| Impdh1 | 0.876190151 |
| Ttc21a | 0.876086974 |
| Nfkbie | 0.875784429 |
| Rnf144a | 0.875715275 |
| Susd1 | 0.875665196 |
| Actg1 | 0.875195325 |
| Sorbs3 | 0.874343871 |
| Rab38 | 0.874318215 |
| Fam53a | 0.874305635 |
| Epha2 | 0.874126033 |
| Slc5a6 | 0.873971291 |
| Spry3 | 0.873946903 |
| Cwc22 | 0.87355469 |
| Aig1 | 0.873347395 |
| Dmap1 | 0.872397872 |

| | |
|---|---|
| Lox | 0.871690889 |
| Dbr1 | 0.871157937 |
| Snip1 | 0.870729972 |
| Cct6a | 0.87021485 |
| Angptl4 | 0.869874925 |
| Vps54 | 0.869815995 |
| Tub | 0.869724696 |
| Piezo2 | 0.86947758 |
| Rad51c | 0.869256004 |
| Stard7 | 0.868911385 |
| Golim4 | 0.86823202 |
| Tbx1 | 0.868209147 |
| Zdhhc23 | 0.868015064 |
| Trmt13 | 0.867873105 |
| Zbtb11 | 0.867562625 |
| Gramd3 | 0.867477402 |
| Eif4a2 | 0.867474823 |
| Nudt16 | 0.867171495 |
| Fbxo2 | 0.867017864 |
| Usp35 | 0.86606495 |
| Hey2 | 0.865695646 |
| Ciz1 | 0.865270964 |
| Fgfr3 | 0.865188135 |
| H2afj | 0.865038674 |
| Ahcy | 0.86419223 |
| Lyl1 | 0.864180199 |
| Dck | 0.863385591 |
| Stab1 | 0.863362624 |
| Klhl36 | 0.863008002 |
| Csad | 0.862493782 |
| Gtf2e2 | 0.862417644 |
| Eva1c | 0.862083314 |
| Golga4 | 0.861241759 |
| Gabarapl1 | 0.86104195 |
| Ptprj | 0.86052513 |
| Mid1 | 0.859955481 |
| Timp2 | 0.859709522 |
| Nlrp14 | 0.859552068 |
| Chn1 | 0.859416707 |
| Zc2hc1a | 0.8579004 |
| Chst15 | 0.857321157 |
| Adamts3 | 0.856923644 |
| Entpd1 | 0.856851669 |
| Ccnf | 0.856478489 |
| Rpl18 | 0.856403535 |
| Ifne | 0.855444182 |
| Ccdc106 | 0.855300573 |

| | |
|---|---|
| Klhl8 | 0.855259808 |
| Mettl4 | 0.855012107 |
| Them4 | 0.854621727 |
| Praf2 | 0.854420954 |
| Il15 | 0.853924111 |
| Yrdc | 0.853637944 |
| Lrrc14 | 0.853359278 |
| Msrb2 | 0.853212962 |
| Agtrap | 0.852799745 |
| Pcsk1 | 0.852797404 |
| Gfpt2 | 0.852794563 |
| Wbscr22 | 0.852214083 |
| Mtx1 | 0.851385802 |
| Gk5 | 0.850903664 |
| Txndc15 | 0.85086125 |
| Dtna | 0.850769129 |
| Btrc | 0.850655477 |
| Il6 | 0.850422766 |
| Fastk | 0.850274001 |
| Smarcad1 | 0.850192246 |
| Slc35g2 | 0.849698083 |
| Snx6 | 0.849682813 |
| Trabd | 0.849542064 |
| St3gal6 | 0.849385014 |
| Serp1 | 0.849093792 |
| Fbxo18 | 0.848854288 |
| Rabggta | 0.848701013 |
| Adrb2 | 0.848696967 |
| Tcp1 | 0.848568943 |
| Ly6g5b | 0.848270216 |
| Rfxap | 0.848190129 |
| Wdr6 | 0.848112264 |
| Timm10b | 0.847134564 |
| Tnrc6c | 0.846745788 |
| Snx5 | 0.846640279 |
| Aco1 | 0.846595842 |
| Rsf1 | 0.846519527 |
| Ctnnd1 | 0.846446344 |
| Pick1 | 0.845203741 |
| Abtb2 | 0.845179068 |
| F2rl3 | 0.84478686 |
| Syngap1 | 0.844722029 |
| Get4 | 0.844189031 |
| Inpp4b | 0.844057656 |
| Colec12 | 0.843842326 |
| Ctnnbl1 | 0.843679759 |
| Fam84b | 0.843588485 |

| | |
|---|---|
| Atp5l | 0.843280087 |
| Ptpn18 | 0.843087832 |
| Taf1d | 0.842851756 |
| Ctsf | 0.842316005 |
| Matn2 | 0.842200441 |
| Dok1 | 0.842127195 |
| Rtkn | 0.842078767 |
| Hdac8 | 0.841341987 |
| Naf1 | 0.84122632 |
| Cep78 | 0.84099189 |
| Prps2 | 0.840968086 |
| Parpbp | 0.840896323 |
| Ss18 | 0.840293429 |
| Nap1l4 | 0.839474278 |
| Ncapg | 0.839352061 |
| Pias3 | 0.839300154 |
| Pex11b | 0.839261633 |
| Ccdc144b | 0.839104919 |
| Fgd4 | 0.838311901 |
| Serac1 | 0.838248553 |
| Elac2 | 0.838217645 |
| Foxk1 | 0.838162995 |
| Gpank1 | 0.837984649 |
| Synj2 | 0.837933181 |
| Fam53c | 0.837610315 |
| Fam210a | 0.837433819 |
| Fbxl15 | 0.837367965 |
| Wdr33 | 0.837182195 |
| Txndc5 | 0.83712193 |
| Ccdc47 | 0.836991896 |
| Ssbp4 | 0.83692121 |
| Atp6v0a1 | 0.836757025 |
| Myh15 | 0.836731917 |
| Peli2 | 0.83615276 |
| Rab40b | 0.835965727 |
| Nup54 | 0.835846526 |
| Selt | 0.835647157 |
| Pigl | 0.835597961 |
| Nmnat3 | 0.835541883 |
| Ythdf2 | 0.835520047 |
| Cldn5 | 0.835224071 |
| Ppp5c | 0.834813067 |
| Fbxw11 | 0.834292426 |
| Ankrd13d | 0.834125578 |
| Grsf1 | 0.833226698 |
| Pole2 | 0.833192143 |
| Ranbp10 | 0.832732524 |

| | |
|---|---|
| Brd8 | 0.832635416 |
| Acvr1b | 0.832624095 |
| Adnp | 0.832610139 |
| Nfatc2 | 0.832587075 |
| 42619 | 0.832228153 |
| Gltpd2 | 0.83176492 |
| Vars2 | 0.831478291 |
| Nexn | 0.831352753 |
| Tbc1d8b | 0.830730801 |
| Kctd7 | 0.830329054 |
| Dtx4 | 0.830193287 |
| Kansl1l | 0.828636567 |
| Ppp1r14b | 0.828629719 |
| Sptlc3 | 0.828488358 |
| Lrrc8e | 0.828328485 |
| Golt1b | 0.827694009 |
| Fgd1 | 0.827439464 |
| Atg5 | 0.827185304 |
| Sec61b | 0.827160298 |
| Dhx34 | 0.827047314 |
| Spa17 | 0.827044619 |
| Ntn1 | 0.826862918 |
| Eny2 | 0.825728069 |
| Ift81 | 0.825715626 |
| Med19 | 0.825021751 |
| Ccdc3 | 0.824718767 |
| Armc6 | 0.824462339 |
| Rrp1b | 0.824064586 |
| Pstpip2 | 0.823955249 |
| Gucd1 | 0.823622303 |
| Ccl7 | 0.822622998 |
| Mcam | 0.822483677 |
| Zyg11b | 0.822408355 |
| Clec1a | 0.822397166 |
| Scly | 0.822200369 |
| Cdkn2a | 0.822194354 |
| Mrpl44 | 0.822038232 |
| Npr1 | 0.821805182 |
| Atp6ap1 | 0.821469935 |
| Kcnmb1 | 0.821420483 |
| Thap3 | 0.820778704 |
| Nmi | 0.82039493 |
| Larp4 | 0.820068113 |
| Srpk2 | 0.820061234 |
| Tcf7 | 0.819673467 |
| Scn5a | 0.819606202 |
| Ccdc87 | 0.818793563 |

| | |
|---|---|
| Arhgef40 | 0.818100094 |
| Tbkbp1 | 0.817958132 |
| Usf1 | 0.81754607 |
| Emg1 | 0.817481875 |
| Mcu | 0.817432204 |
| Armcx4 | 0.817428498 |
| Ipo5 | 0.817111057 |
| Tbc1d2 | 0.816298945 |
| Smim15 | 0.815947765 |
| Slc15a3 | 0.815831088 |
| Ier3ip1 | 0.815446472 |
| Renbp | 0.815395493 |
| Anp32e | 0.815046442 |
| Reep2 | 0.814994668 |
| Arntl | 0.813969932 |
| Bdh2 | 0.813791489 |
| Ube2d3 | 0.813300927 |
| Fstl3 | 0.813173464 |
| Cfh | 0.813154727 |
| Agbl5 | 0.81303174 |
| Polr2f | 0.813003235 |
| Gdf1 | 0.812997711 |
| Pdlim5 | 0.812701554 |
| Slc25a40 | 0.812627761 |
| Eps15 | 0.812325234 |
| Aifm1 | 0.812060735 |
| Gdpgp1 | 0.812002123 |
| Cldn7 | 0.811576558 |
| Bzw2 | 0.811508885 |
| Szrd1 | 0.811399397 |
| Ypel5 | 0.811151485 |
| Aim1 | 0.811140039 |
| Cope | 0.81112828 |
| Lpin2 | 0.810165706 |
| Uba2 | 0.809480121 |
| Dynlrb1 | 0.809397389 |
| Zc3hav1 | 0.808967157 |
| Cdc42ep4 | 0.808810054 |
| Tnks2 | 0.808596086 |
| Snx7 | 0.808591052 |
| Bms1 | 0.808390166 |
| Rhobtb3 | 0.808088275 |
| Phospho1 | 0.807934933 |
| Eda2r | 0.807837162 |
| Man2a1 | 0.80723637 |
| Rpl23 | 0.806541646 |
| Opn1sw | 0.80625059 |

| | |
|---|---|
| Syn1 | 0.806178334 |
| Prcp | 0.805889729 |
| Zfyve26 | 0.804886758 |
| Acadm | 0.804733961 |
| Inpp5e | 0.80431338 |
| Atp5g3 | 0.804047667 |
| Ano6 | 0.803384775 |
| Phf12 | 0.803259137 |
| Eepd1 | 0.803234424 |
| Ccdc71 | 0.803230083 |
| Ankzf1 | 0.803116292 |
| Spint2 | 0.802859751 |
| Snx1 | 0.802796299 |
| Ipo4 | 0.802138584 |
| Ccne2 | 0.801852608 |
| Dtx3 | 0.801660741 |
| Ptbp1 | 0.801621195 |
| Ccna1 | 0.800922747 |
| Rnf169 | 0.800920904 |
| Aaas | 0.800826786 |
| Nfkb1 | 0.800802514 |
| Uba6 | 0.800563363 |
| Abi2 | 0.799743717 |
| Cdh24 | 0.799720039 |
| Mrpl19 | 0.799336242 |
| Ccnd2 | 0.799172569 |
| Uspl1 | 0.798569029 |
| Hsd17b1 | 0.798454626 |
| Hdx | 0.798436822 |
| Fcf1 | 0.797999614 |
| Efcab5 | 0.797692393 |
| Lmo4 | 0.797657102 |
| Camkmt | 0.79715699 |
| Ddx3x | 0.797156377 |
| Oser1 | 0.796931637 |
| Mtmr1 | 0.796320558 |
| Lrrc27 | 0.796306081 |
| Fdx1l | 0.795965515 |
| Sash1 | 0.795903492 |
| Cideb | 0.795848987 |
| Man1c1 | 0.795370137 |
| Smchd1 | 0.795088863 |
| Xpot | 0.794730636 |
| Klf12 | 0.794518867 |
| Camk1 | 0.794283703 |
| Prkab2 | 0.794209222 |
| Vamp5 | 0.794104353 |

| | |
|---|---|
| N6amt2 | 0.793351232 |
| Fam103a1 | 0.793260657 |
| C1d | 0.793073923 |
| Slitrk4 | 0.793061369 |
| Ppat | 0.792989998 |
| Hipk1 | 0.792952916 |
| Ccbl1 | 0.7927521 |
| Jtb | 0.792745483 |
| Hexim1 | 0.792681189 |
| Zkscan8 | 0.79264722 |
| Xpc | 0.791688255 |
| Arid1a | 0.791642817 |
| Srgn | 0.791433046 |
| Myl6 | 0.791237275 |
| Gsap | 0.790793655 |
| Wwtr1 | 0.789841671 |
| Sc5d | 0.78953579 |
| S100a4 | 0.78941253 |
| Cntn3 | 0.789256174 |
| Cry2 | 0.789209091 |
| Fam60a | 0.789014516 |
| Hsdl2 | 0.787921416 |
| Gnl2 | 0.787751438 |
| Arid5b | 0.787669174 |
| Ccna2 | 0.786631162 |
| Pabpc5 | 0.786549111 |
| Nagk | 0.786365866 |
| Cnksr3 | 0.786051581 |
| Erg | 0.78517547 |
| Lrwd1 | 0.784824641 |
| Cln3 | 0.784412032 |
| Snapc1 | 0.784269826 |
| Egfl7 | 0.784204668 |
| Jak2 | 0.784075395 |
| Cul1 | 0.78399475 |
| Disp2 | 0.783776347 |
| Tubg1 | 0.783590089 |
| Cenpw | 0.78346064 |
| Dkk1 | 0.7831807 |
| Mrpl46 | 0.782835593 |
| P2ry1 | 0.78244852 |
| Aggf1 | 0.782114086 |
| Atp5sl | 0.781970293 |
| Psma5 | 0.781756673 |
| Ttc30b | 0.781566093 |
| Wdr4 | 0.781214514 |
| Sdccag8 | 0.780452414 |

| | |
|---|---|
| Zcchc7 | 0.780361938 |
| Nrsn2 | 0.780150664 |
| Tdg | 0.779934341 |
| Lrp6 | 0.779120828 |
| Ehd3 | 0.778414944 |
| Pabpc4l | 0.778242209 |
| Adarb1 | 0.778130625 |
| Zgpat | 0.776836394 |
| Gpm6a | 0.776466651 |
| Bzw1 | 0.77603126 |
| Rab1b | 0.775971056 |
| Nrbp1 | 0.775768246 |
| Vps45 | 0.775476389 |
| Lrig3 | 0.775279753 |
| Prr11 | 0.775209937 |
| Abhd15 | 0.774977274 |
| Plekhm1 | 0.774636036 |
| Cdc34 | 0.774511206 |
| Traf1 | 0.774488878 |
| Micu2 | 0.774419913 |
| Bex4 | 0.774191801 |
| Polr1d | 0.773833359 |
| Ip6k1 | 0.773250011 |
| Kdm5d | 0.772779791 |
| Casp8ap2 | 0.772620365 |
| Telo2 | 0.772371518 |
| Sult1a1 | 0.771892624 |
| Slc19a2 | 0.771758235 |
| Erp44 | 0.771484131 |
| Tmod1 | 0.770746456 |
| Tnr | 0.770385763 |
| Anpep | 0.77038384 |
| Suv39h2 | 0.769033916 |
| Ktn1 | 0.768853977 |
| Rsad1 | 0.767806202 |
| Pspc1 | 0.767718189 |
| Bag6 | 0.76751948 |
| Rab11fip1 | 0.767084541 |
| Ighmbp2 | 0.766764313 |
| Ldhb | 0.766608955 |
| Wdr36 | 0.766455739 |
| Paics | 0.766412667 |
| Acp6 | 0.766406935 |
| Spink5 | 0.766402019 |
| Grin2a | 0.76634934 |
| Sppl2a | 0.766345798 |
| Pop5 | 0.76621488 |

| | |
|---|---|
| Mllt3 | 0.765974298 |
| Grin2c | 0.764777234 |
| Rad23b | 0.764460102 |
| Cpeb4 | 0.764402747 |
| Adipor1 | 0.764059127 |
| Krt7 | 0.763687866 |
| Cyb561a3 | 0.763534533 |
| Klf7 | 0.763353359 |
| Clu | 0.763247784 |
| Tef | 0.762676054 |
| Dnm1l | 0.762618857 |
| Atp5j | 0.762374683 |
| Sf3a2 | 0.762367633 |
| Pald1 | 0.762171715 |
| Osbpl6 | 0.760669874 |
| Cox20 | 0.760640742 |
| Pdk3 | 0.760616003 |
| Tango2 | 0.76032253 |
| Slc25a21 | 0.759973826 |
| Pllp | 0.759820221 |
| Dusp5 | 0.759531583 |
| Hdgf | 0.759093561 |
| Chordc1 | 0.758898367 |
| Tmem39b | 0.758858178 |
| Cfb | 0.75885436 |
| Nbas | 0.758731106 |
| Cdyl2 | 0.758253104 |
| Gpx8 | 0.758044401 |
| Spred2 | 0.757795066 |
| Dolpp1 | 0.757252059 |
| Brd3 | 0.757130715 |
| Tes | 0.756879068 |
| Psen1 | 0.756517539 |
| H2afy | 0.756269889 |
| Pwp2 | 0.755751231 |
| Vps35 | 0.755473889 |
| Cpt1b | 0.755212894 |
| Cant1 | 0.754647158 |
| Zmym2 | 0.753698515 |
| Surf1 | 0.753396557 |
| Ercc6l2 | 0.753190968 |
| Tmed2 | 0.753018946 |
| Large | 0.752960691 |
| Rnf2 | 0.752943902 |
| Ube2r2 | 0.75254672 |
| Enthd2 | 0.752483209 |
| Ccdc30 | 0.751960784 |

| | |
|---|---|
| Ctc1 | 0.751754717 |
| Jmjd1c | 0.751727415 |
| Fuca2 | 0.751669289 |
| Cdc16 | 0.751615236 |
| Ndufb11 | 0.751215092 |
| Tcf25 | 0.750987462 |
| Abca3 | 0.750815325 |
| Bdnf | 0.750773313 |
| Col6a2 | 0.750506198 |
| Calcoco2 | 0.74976721 |
| Klrg1 | 0.749265452 |
| Pus1 | 0.749190964 |
| Fzd7 | 0.749056245 |
| Scaper | 0.748928203 |
| Mgat4b | 0.748629671 |
| Fanci | 0.748589329 |
| Blm | 0.747915291 |
| Rad9b | 0.747892158 |
| Hyls1 | 0.747788179 |
| Akip1 | 0.747717607 |
| Ccdc25 | 0.747309754 |
| Sbno1 | 0.746269901 |
| Yod1 | 0.746217635 |
| Eml3 | 0.745909248 |
| Rwdd3 | 0.745789545 |
| Scaf1 | 0.745730936 |
| Mrpl18 | 0.74535381 |
| Pes1 | 0.745189112 |
| Lpcat1 | 0.744600689 |
| Erbb2ip | 0.744194961 |
| Pcnp | 0.744040826 |
| Xpo4 | 0.743642097 |
| Zfp28 | 0.743107989 |
| Thsd7a | 0.742651197 |
| Aph1b | 0.742534211 |
| Qpct | 0.742346774 |
| Ttc26 | 0.741086456 |
| Tsfm | 0.741059964 |
| Stra13 | 0.740848978 |
| Tnc | 0.740292763 |
| Ttc17 | 0.739888063 |
| Setd8 | 0.739540627 |
| Nf2 | 0.739500901 |
| Slc25a53 | 0.739292098 |
| Tmem246 | 0.739245546 |
| Spock1 | 0.738960257 |
| Atp2b1 | 0.73870565 |

| | |
|---|---|
| Nupr1 | 0.738614326 |
| Slc25a30 | 0.738439797 |
| Syde2 | 0.738359152 |
| Cltc | 0.738321142 |
| Mri1 | 0.738320207 |
| Xpa | 0.738164995 |
| Rcan2 | 0.737827122 |
| Ccdc120 | 0.737660124 |
| Otud4 | 0.737579413 |
| Ptpn9 | 0.737272686 |
| Wdsub1 | 0.736999467 |
| Cyth2 | 0.736447162 |
| Tm7sf3 | 0.735914132 |
| Ceacam19 | 0.735874766 |
| Atp2a1 | 0.735796603 |
| Ccdc40 | 0.735685507 |
| Eef1e1 | 0.735481303 |
| Kif2a | 0.735227942 |
| Ptpn6 | 0.735055494 |
| Evi5l | 0.734696129 |
| Fam13b | 0.734484687 |
| Atp6v1e1 | 0.734265121 |
| 42437 | 0.734023357 |
| Emc3 | 0.733354592 |
| Commd7 | 0.733200496 |
| Uri1 | 0.733198639 |
| Mppe1 | 0.732779876 |
| Sertad1 | 0.732672666 |
| Dnaja3 | 0.732569313 |
| Nek8 | 0.732187949 |
| Smug1 | 0.731832385 |
| Vill | 0.73171046 |
| Mob3a | 0.731536869 |
| Nol3 | 0.731271098 |
| Coro1b | 0.730955849 |
| Neb | 0.730580661 |
| Tbc1d32 | 0.730214912 |
| Abcc4 | 0.730145306 |
| Nrg3 | 0.730091609 |
| Taf13 | 0.729976463 |
| Sfr1 | 0.729760348 |
| Fhdc1 | 0.729010205 |
| Epsti1 | 0.728942127 |
| Prpf19 | 0.728315326 |
| Gnas | 0.728298043 |
| Mis18a | 0.728121548 |
| Rplp0 | 0.728083897 |

| | |
|---|---|
| Fam195b | 0.727723527 |
| Ptov1 | 0.727498167 |
| Ehmt2 | 0.727399284 |
| Ccin | 0.727234487 |
| Ccdc109b | 0.726798386 |
| Cdc14a | 0.726785646 |
| Sbk1 | 0.726734792 |
| Fktn | 0.726730827 |
| Pfkm | 0.726440551 |
| Prr16 | 0.726397133 |
| Kars | 0.726344863 |
| Aspm | 0.726194373 |
| Btd | 0.725596359 |
| Lrif1 | 0.725476066 |
| Sv2a | 0.724514256 |
| Ikbkap | 0.724371652 |
| Naaladl1 | 0.724179859 |
| Dact1 | 0.723947518 |
| Mgea5 | 0.723625033 |
| Rev1 | 0.723226312 |
| Sec24d | 0.723139358 |
| Marcksl1 | 0.72308988 |
| Rpl37a | 0.72307813 |
| Ikbip | 0.722890141 |
| Hoxd4 | 0.722712486 |
| Tmem54 | 0.722131281 |
| Gcsh | 0.721983862 |
| Rdh10 | 0.721761785 |
| Nmt2 | 0.721143814 |
| Gpc1 | 0.720732376 |
| Ppfia2 | 0.720217991 |
| Cox14 | 0.720207826 |
| Alg14 | 0.719408527 |
| Mrpl53 | 0.719368518 |
| Arhgap6 | 0.719229997 |
| Smad6 | 0.718916196 |
| Zfp30 | 0.718697488 |
| Taco1 | 0.718337405 |
| Cox6c | 0.717614007 |
| Zfp41 | 0.714911541 |
| Akap9 | 0.714698135 |
| Rptor | 0.71458979 |
| Med21 | 0.714097499 |
| Safb2 | 0.713756797 |
| Prkd3 | 0.713591922 |
| Rnf187 | 0.712881331 |
| Recql | 0.712626121 |

| | |
|---|---|
| Tmeff1 | 0.712606178 |
| Fam118b | 0.712545252 |
| Hspa9 | 0.712022646 |
| Med29 | 0.71201889 |
| Yipf4 | 0.711885905 |
| Cdkl1 | 0.711879054 |
| Mphosph8 | 0.710519801 |
| Acad11 | 0.710456539 |
| Eif2b3 | 0.709172515 |
| Mcm2 | 0.708657345 |
| Mrps6 | 0.708401316 |
| Ndufa7 | 0.707849538 |
| Cttn | 0.707841189 |
| Gclm | 0.706315725 |
| Gpr176 | 0.706255042 |
| Dync2h1 | 0.7058299 |
| Jund | 0.705496962 |
| Il16 | 0.704968372 |
| Rps29 | 0.704947652 |
| Kat5 | 0.704783137 |
| Dhx9 | 0.70475167 |
| Card10 | 0.704605503 |
| Ccdc36 | 0.704459916 |
| Napa | 0.704041233 |
| E2f1 | 0.70388587 |
| Stmn3 | 0.703478683 |
| Camk2d | 0.703473608 |
| Sptbn1 | 0.703116801 |
| Ext1 | 0.702709137 |
| Tpra1 | 0.702344437 |
| Prpf40b | 0.70102831 |
| Kdm7a | 0.700777293 |
| Sqle | 0.700690897 |
| Zdhhc17 | 0.700336303 |
| Qars | 0.700324211 |
| Abhd6 | 0.70004384 |
| Mapkap1 | 0.699096523 |
| Acap1 | 0.697797385 |
| Ranbp17 | 0.697720223 |
| Lancl1 | 0.697229189 |
| R3hcc1 | 0.695872421 |
| Capn1 | 0.695710491 |
| Kmt2e | 0.695613589 |
| Mapre3 | 0.695108985 |
| Ssbp2 | 0.694814009 |
| Tfpi2 | 0.694740571 |
| Rfwd3 | 0.694017776 |

| | |
|---|---|
| Adrbk2 | 0.693332834 |
| Chsy3 | 0.692867257 |
| Palb2 | 0.692458301 |
| Arap3 | 0.692404704 |
| Mms22l | 0.692320108 |
| Mpzl2 | 0.692263032 |
| Dkk3 | 0.691348152 |
| Abr | 0.691300453 |
| Inhba | 0.691034151 |
| Fam134a | 0.690795639 |
| Ankmy1 | 0.690748988 |
| Ubap1 | 0.69028545 |
| Kif21a | 0.689956409 |
| Meaf6 | 0.689499205 |
| Apc2 | 0.689010366 |
| Intu | 0.688816179 |
| Zcchc3 | 0.688745862 |
| Rnf207 | 0.688531034 |
| Rpl27a | 0.688475311 |
| Krcc1 | 0.688446191 |
| Gsto1 | 0.688129707 |
| Tceal1 | 0.68787164 |
| Nprl3 | 0.687672146 |
| Ate1 | 0.687624373 |
| Fblim1 | 0.687528094 |
| Ptgs2 | 0.687481969 |
| Diexf | 0.687366858 |
| Podxl | 0.686706123 |
| Gipc3 | 0.686166122 |
| Odf2 | 0.685891059 |
| Tmem143 | 0.68553109 |
| Faf2 | 0.68518941 |
| Tulp4 | 0.685147838 |
| Tmem163 | 0.684916911 |
| Ranbp3 | 0.684877118 |
| Plek2 | 0.684755076 |
| Glipr1 | 0.684713175 |
| Capn11 | 0.684636862 |
| Coq3 | 0.684265937 |
| Il18bp | 0.683827857 |
| Cpsf7 | 0.683650968 |
| Tsta3 | 0.683375299 |
| Lcat | 0.683126662 |
| Ints12 | 0.683035566 |
| Ppil6 | 0.682438619 |
| Ankrd39 | 0.682242725 |
| Mapk8ip3 | 0.680975273 |

| | |
|---|---|
| Ago1 | 0.680638561 |
| Gadd45b | 0.680458351 |
| Mtpap | 0.680126175 |
| Tspan31 | 0.680019341 |
| Kdelr2 | 0.6796764 |
| Aars | 0.67956507 |
| Sufu | 0.679557167 |
| Ldlrap1 | 0.67936024 |
| Zbtb3 | 0.679044362 |
| Mmp25 | 0.678713458 |
| Mical2 | 0.678366541 |
| Mcm3 | 0.678305272 |
| Rnls | 0.677874905 |
| Sdhd | 0.67778142 |
| Ezh2 | 0.677704581 |
| Usp3 | 0.677678353 |
| Peli3 | 0.677606032 |
| Sgce | 0.676724554 |
| Tspan17 | 0.676295968 |
| Fam109a | 0.675732577 |
| Bivm | 0.675658093 |
| Rdx | 0.675309185 |
| Snx15 | 0.675126993 |
| Nsmce2 | 0.675028972 |
| Ppm1l | 0.674287397 |
| Ckap4 | 0.673722389 |
| Map1lc3a | 0.673359058 |
| Kpna3 | 0.673300855 |
| Vrk3 | 0.673276928 |
| Dusp8 | 0.672328674 |
| Rasa1 | 0.671845931 |
| Tpm2 | 0.671672039 |
| Galnt12 | 0.67072171 |
| Kptn | 0.670616393 |
| Unc13a | 0.670570722 |
| Tpcn1 | 0.67053776 |
| Spsb3 | 0.670244971 |
| Wdr55 | 0.670161674 |
| Tpp1 | 0.67006487 |
| Rnf34 | 0.669661557 |
| Sgpl1 | 0.669091067 |
| Rilpl1 | 0.668922783 |
| Capns1 | 0.668033081 |
| Nt5dc2 | 0.667358131 |
| Dnajc2 | 0.667349758 |
| Evi2b | 0.666681059 |
| Gpatch4 | 0.6665798 |

| | |
|---|---|
| Prune2 | 0.665683011 |
| Crtac1 | 0.665164837 |
| Sirpa | 0.664992115 |
| Hook2 | 0.664963206 |
| Cdk17 | 0.664540741 |
| Hars2 | 0.663866525 |
| Polr2i | 0.66371909 |
| Fntb | 0.663217583 |
| Ino80c | 0.663188044 |
| Ccs | 0.66306973 |
| Fam220a | 0.662808477 |
| Lgals8 | 0.662692834 |
| Styxl1 | 0.662459322 |
| Traf3 | 0.662119942 |
| Qsox1 | 0.662089968 |
| Lap3 | 0.661949728 |
| Rbbp6 | 0.66165799 |
| Sgol1 | 0.660935782 |
| Thg1l | 0.660636761 |
| Sh3d19 | 0.660558832 |
| Ttc8 | 0.660415941 |
| Icmt | 0.659921473 |
| Ipo13 | 0.659919636 |
| Ppil1 | 0.659755838 |
| Lyar | 0.659651525 |
| Nkx3-1 | 0.659613794 |
| Rasd1 | 0.659499682 |
| Wipi1 | 0.6586877 |
| Pola1 | 0.658608725 |
| Pitx1 | 0.657876554 |
| Pcnt | 0.657712193 |
| Kctd1 | 0.657563829 |
| Ablim2 | 0.657375038 |
| Ppp2r3c | 0.657359989 |
| Fadd | 0.656528779 |
| Atp5a1 | 0.656477027 |
| Aqp11 | 0.656296242 |
| Cpe | 0.656106961 |
| Ccdc130 | 0.655581052 |
| Map1b | 0.655283111 |
| Meis2 | 0.654843268 |
| Adamts15 | 0.654763137 |
| Ifi44l | 0.654673382 |
| Nphp4 | 0.654671549 |
| Banf1 | 0.654531869 |
| Urb2 | 0.654481777 |
| Aacs | 0.653954791 |

| | |
|---|---|
| Rars | 0.653428332 |
| Ppp2r5b | 0.653418842 |
| Nadk2 | 0.6528692 |
| Gli3 | 0.652504058 |
| Msl1 | 0.651459585 |
| Steap2 | 0.651019485 |
| Ehd2 | 0.650728743 |
| Atp5e | 0.649554836 |
| Ajuba | 0.649521426 |
| Ift46 | 0.649471197 |
| Coq4 | 0.648708742 |
| Hmga2 | 0.648457972 |
| Snca | 0.64790795 |
| Kctd16 | 0.647833001 |
| Chchd5 | 0.647448896 |
| Ube2g2 | 0.647243798 |
| Mrps31 | 0.647202952 |
| Prrg1 | 0.647190207 |
| Rnf10 | 0.647025222 |
| Slc2a13 | 0.646269135 |
| Sar1a | 0.645734012 |
| Eaf2 | 0.645629955 |
| Ank1 | 0.645625249 |
| Rab34 | 0.645477748 |
| Usp31 | 0.645419022 |
| Tcf4 | 0.645342324 |
| Crispld1 | 0.645171192 |
| Hnrnpa0 | 0.645017935 |
| Wdr75 | 0.644916323 |
| Marcks | 0.644564045 |
| Hrasls | 0.644496035 |
| Cdca8 | 0.644202073 |
| Akap8l | 0.644120848 |
| Mynn | 0.64348334 |
| Pou2f1 | 0.643376563 |
| Plekhm2 | 0.643316474 |
| Rragc | 0.642931144 |
| Rnf125 | 0.642766889 |
| Gper1 | 0.642618658 |
| Shc1 | 0.64223274 |
| Pi4kb | 0.642196207 |
| Slc25a43 | 0.641970575 |
| Coq10b | 0.641969278 |
| Nts | 0.640909445 |
| Atf1 | 0.640800735 |
| Sart3 | 0.640757191 |
| Nans | 0.6398889 |

| | |
|---|---|
| Nusap1 | 0.639656803 |
| Exosc6 | 0.639346293 |
| Dpf1 | 0.639229182 |
| Rcl1 | 0.638665374 |
| Trib3 | 0.637932457 |
| Sass6 | 0.637794944 |
| Siah2 | 0.637772736 |
| Slc16a9 | 0.637554042 |
| Adhfe1 | 0.636693612 |
| Fcho2 | 0.635973602 |
| Ttc39a | 0.635952888 |
| Grn | 0.635715397 |
| Kifc1 | 0.635053576 |
| Bub3 | 0.634636282 |
| Nfs1 | 0.634133834 |
| Med9 | 0.634078202 |
| Mocos | 0.634005237 |
| Isg20l2 | 0.633946951 |
| Rfesd | 0.633063949 |
| Nedd9 | 0.632905663 |
| Rab11fip5 | 0.632879187 |
| Abhd2 | 0.632263362 |
| Hcfc1r1 | 0.632104368 |
| Decr1 | 0.631879155 |
| Htatsf1 | 0.631723052 |
| Ofd1 | 0.630820998 |
| Atp8b1 | 0.630678844 |
| Hat1 | 0.630310728 |
| Cuedc2 | 0.630304867 |
| Gigyf2 | 0.630236006 |
| Slc6a9 | 0.629938664 |
| St3gal1 | 0.629463537 |
| Lig1 | 0.629402921 |
| Wdr62 | 0.628978887 |
| Fam222b | 0.628131445 |
| Rpl4 | 0.627729503 |
| Tardbp | 0.627592163 |
| Ahcyl2 | 0.627555579 |
| Mpg | 0.627228593 |
| Cyb5r3 | 0.627092416 |
| Slc25a36 | 0.626978051 |
| Plekhb2 | 0.62692536 |
| Hspb6 | 0.626794203 |
| Rcn3 | 0.626599844 |
| Sipa1l2 | 0.626098254 |
| Prss53 | 0.626067769 |
| Bcs1l | 0.62601339 |

| | |
|---|---|
| F11r | 0.625028807 |
| Tmem2 | 0.625010921 |
| Pdcd11 | 0.624850418 |
| Rhbdf1 | 0.624655238 |
| Smap2 | 0.623715242 |
| Ascc2 | 0.623668905 |
| Csmd1 | 0.623584618 |
| Pvr | 0.622547511 |
| Plscr1 | 0.622431127 |
| Arhgef10 | 0.622197165 |
| Ak7 | 0.622167932 |
| Dact3 | 0.622036444 |
| Kcnip3 | 0.621836264 |
| Derl1 | 0.621492007 |
| Tmem35 | 0.62147603 |
| Smyd5 | 0.621158902 |
| Zhx1 | 0.621141478 |
| Adnp2 | 0.620881197 |
| Kynu | 0.619618481 |
| Xkr6 | 0.619298697 |
| Snrpg | 0.619271276 |
| Prmt6 | 0.618532282 |
| Jph1 | 0.618256226 |
| Mthfsd | 0.617593312 |
| Cct2 | 0.61657669 |
| Sec24a | 0.616511196 |
| Tmem47 | 0.616367239 |
| Pi4k2a | 0.615870472 |
| Larp7 | 0.615694815 |
| Sptssa | 0.615333125 |
| Fbxo46 | 0.614850971 |
| Tmcc1 | 0.614564857 |
| Rer1 | 0.614531557 |
| Cd2bp2 | 0.614527865 |
| Gsk3b | 0.614340838 |
| Cln5 | 0.614266086 |
| Ccdc39 | 0.614233553 |
| Ankrd1 | 0.614221948 |
| Gemin4 | 0.613591172 |
| Dyrk4 | 0.613526053 |
| Actn2 | 0.613152943 |
| Lbh | 0.612622211 |
| Ppp1r8 | 0.612101915 |
| Tti1 | 0.611899473 |
| Cxxc1 | 0.611881618 |
| Kif9 | 0.610838865 |
| Sav1 | 0.609909573 |

| | |
|---|---|
| Rgmb | 0.609647538 |
| Mmp14 | 0.609037908 |
| Pp2d1 | 0.608998507 |
| Tmem161b | 0.608942637 |
| Uchl1 | 0.608932753 |
| Glrb | 0.607623099 |
| Nphp1 | 0.606707936 |
| Aph1a | 0.605826512 |
| Mrps35 | 0.6057451 |
| Ccnj | 0.60570156 |
| Rexo1 | 0.60544083 |
| Herc2 | 0.605239263 |
| B4galt3 | 0.604749209 |
| Cfl2 | 0.604299927 |
| Tollip | 0.603987351 |
| Lrrc20 | 0.603966886 |
| Taldo1 | 0.603934158 |
| Cxcr4 | 0.603503089 |
| Tnip3 | 0.60337595 |
| Gga2 | 0.603256764 |
| Nup155 | 0.602241345 |
| Ubox5 | 0.602211713 |
| Slc8a1 | 0.602088547 |
| Atp9b | 0.601898312 |
| Trim23 | 0.601734844 |
| Lamb1 | 0.601409355 |
| Ticam2 | 0.601132432 |
| Plscr4 | 0.601034352 |
| Extl2 | 0.600775393 |
| Rnf24 | 0.600335725 |
| Chek2 | 0.599962925 |
| Mtmr11 | 0.599320705 |
| Homez | 0.599186739 |
| Mgrn1 | 0.598944804 |
| Nnat | 0.598938842 |
| Smc6 | 0.59866683 |
| Nudt10 | 0.598478629 |
| Nckap1l | 0.598264406 |
| Hira | 0.598256317 |
| Ppp1r16a | 0.597996653 |
| Tpm3 | 0.596975018 |
| Dis3l2 | 0.59652482 |
| Dlc1 | 0.596222508 |
| Sh2b1 | 0.595957364 |
| Wrap73 | 0.595942991 |
| 42614 | 0.595779324 |
| Tbc1d24 | 0.595436041 |

| | |
|---|---|
| Rfng | 0.594328535 |
| Lonp2 | 0.594285222 |
| Tgfbr1 | 0.594156928 |
| Asb3 | 0.593508447 |
| Gltscr1l | 0.593473368 |
| Vprbp | 0.593343778 |
| Ppm1b | 0.593331495 |
| Mboat7 | 0.592988124 |
| Nhsl2 | 0.59290099 |
| Crmp1 | 0.592760636 |
| Timm9 | 0.592644396 |
| Brd4 | 0.592028083 |
| Slc50a1 | 0.591494427 |
| Ntpcr | 0.590936326 |
| Habp4 | 0.59062446 |
| Dhcr24 | 0.590342896 |
| Ahctf1 | 0.590156523 |
| Iqgap1 | 0.590109174 |
| Cecr2 | 0.590108232 |
| Dhx35 | 0.58961048 |
| Arhgap25 | 0.589368643 |
| Trerf1 | 0.588978962 |
| Fars2 | 0.588618109 |
| Trip12 | 0.58858558 |
| Osbpl3 | 0.588355229 |
| Ocln | 0.588217767 |
| Rpl27 | 0.588026408 |
| Clip3 | 0.587715728 |
| Mier1 | 0.587331492 |
| Nup214 | 0.586999049 |
| Uchl3 | 0.586931746 |
| Layn | 0.586896696 |
| Src | 0.586465908 |
| Stk35 | 0.586336387 |
| Fmnl2 | 0.586244139 |
| Bcl7a | 0.585941222 |
| Pon2 | 0.585419442 |
| Sp6 | 0.585396893 |
| Nrl | 0.585034858 |
| Mlip | 0.584947932 |
| Thoc5 | 0.58492477 |
| Uqcc2 | 0.584862283 |
| Cables1 | 0.584845747 |
| Cyld | 0.58483296 |
| Notch3 | 0.58468596 |
| Nudt15 | 0.583786957 |
| Inppl1 | 0.583764802 |

| | |
|---|---|
| Ddx51 | 0.583693162 |
| Exosc9 | 0.583164036 |
| Rps19bp1 | 0.583135871 |
| Crkl | 0.58265594 |
| Crem | 0.58255777 |
| Tmem39a | 0.582347485 |
| Arap2 | 0.582238866 |
| Spata5l1 | 0.581823496 |
| Rab9b | 0.581802441 |
| Slc30a5 | 0.581174894 |
| Ext2 | 0.581160247 |
| Kcnmb3 | 0.580879754 |
| Rab15 | 0.580861354 |
| Galm | 0.580754876 |
| Cpne8 | 0.580573164 |
| Zfp3 | 0.57899115 |
| Eno2 | 0.57822327 |
| Exoc1 | 0.578185544 |
| Tsc22d2 | 0.577865111 |
| Taf10 | 0.577198905 |
| Spr | 0.576934045 |
| Riok3 | 0.576602732 |
| Galnt1 | 0.576406595 |
| Tbc1d4 | 0.576122491 |
| Clic1 | 0.574769216 |
| Pthlh | 0.573798155 |
| Pex2 | 0.573321736 |
| Mitd1 | 0.572623025 |
| Zbtb6 | 0.572552394 |
| Rpp25 | 0.571990826 |
| Sgms1 | 0.571708167 |
| Mtss1 | 0.571548103 |
| Mastl | 0.571491216 |
| Dhx30 | 0.571299926 |
| Nfib | 0.570999648 |
| Trim36 | 0.570757739 |
| B3galnt1 | 0.570699757 |
| Rars2 | 0.570415915 |
| Fuca1 | 0.570238161 |
| Nova2 | 0.569697331 |
| Mrps33 | 0.569526257 |
| Pycr1 | 0.569357458 |
| Plcb2 | 0.568403876 |
| Efnb1 | 0.568108804 |
| Mcph1 | 0.56714441 |
| Necap1 | 0.566953338 |
| Pgpep1 | 0.566717443 |

| | |
|---|---|
| Fank1 | 0.566444991 |
| Mrpl16 | 0.56642902 |
| Atg2b | 0.566271802 |
| Suox | 0.566139182 |
| Psmb7 | 0.565981575 |
| Pnma2 | 0.565751242 |
| Scarb2 | 0.564725563 |
| Pip5k1a | 0.564506887 |
| Rpl28 | 0.564426379 |
| Idh3b | 0.564254664 |
| Dpm1 | 0.563479163 |
| Amd1 | 0.563239242 |
| Nmd3 | 0.563114464 |
| Zscan12 | 0.562797847 |
| Cfl1 | 0.562701439 |
| Snrnp200 | 0.562022443 |
| Rbm22 | 0.561942147 |
| Syt14 | 0.561653169 |
| Rnf26 | 0.561651038 |
| H2afv | 0.559561451 |
| Prh1 | 0.559319883 |
| Relb | 0.559007881 |
| Fpgt | 0.558892022 |
| Bmp2 | 0.55883031 |
| Mapk1 | 0.558223926 |
| Ipo11 | 0.557888145 |
| Dusp11 | 0.557275647 |
| Ccdc24 | 0.557203667 |
| Cops5 | 0.556747867 |
| Parp2 | 0.55673624 |
| Prrc1 | 0.556549817 |
| Hoxa11 | 0.556001424 |
| Dgat2 | 0.555690515 |
| Echdc3 | 0.555542099 |
| Trub2 | 0.55549114 |
| Rab23 | 0.554322321 |
| Tspan2 | 0.55410557 |
| Cog4 | 0.554031457 |
| Stx8 | 0.553782282 |
| Echs1 | 0.553768646 |
| Exosc10 | 0.553269637 |
| Erlin2 | 0.552785564 |
| Mycbp2 | 0.551823557 |
| Hnrnpa3 | 0.551372187 |
| Prr3 | 0.550737361 |
| Msh6 | 0.55062711 |
| Dnajb1 | 0.550132149 |

| | |
|---|---|
| Fance | 0.550116059 |
| Polr2m | 0.549894203 |
| Wsb1 | 0.549711479 |
| Kctd2 | 0.549706808 |
| Lca5 | 0.549371919 |
| Bcl9 | 0.549278216 |
| Rpp21 | 0.549213954 |
| Fra10ac1 | 0.548958261 |
| Tbp | 0.548731427 |
| Elovl6 | 0.548605622 |
| Msra | 0.548522074 |
| Tmc7 | 0.547514616 |
| Chrnb1 | 0.547460535 |
| Pdgfrl | 0.54706676 |
| Prkaa1 | 0.546835047 |
| Zmynd8 | 0.546803899 |
| Lsm14a | 0.54630064 |
| Ppp1r9a | 0.545993022 |
| Cd34 | 0.545918323 |
| Prkd2 | 0.54576938 |
| Mif | 0.545711679 |
| Hhat | 0.545286551 |
| Gtf2b | 0.544830036 |
| Sh3pxd2b | 0.54415024 |
| Dlg2 | 0.543891963 |
| Acsl1 | 0.543825919 |
| Nt5c | 0.543715348 |
| Nol12 | 0.543682165 |
| Pgap3 | 0.543150798 |
| Polr2k | 0.542985073 |
| Thsd1 | 0.542595612 |
| Tmem237 | 0.54231072 |
| Slc35c1 | 0.542301541 |
| Paf1 | 0.542248666 |
| C1qtnf6 | 0.542179617 |
| Xbp1 | 0.542019952 |
| Fam109b | 0.541858375 |
| Col5a2 | 0.541215818 |
| Prkar1b | 0.539280306 |
| Slc25a39 | 0.53904813 |
| Stip1 | 0.538748771 |
| Samd14 | 0.53816617 |
| Pold3 | 0.538024502 |
| Auh | 0.537936037 |
| Laptm4b | 0.537882512 |
| Nmb | 0.537382585 |
| Rap1gds1 | 0.537381931 |

| | |
|---|---|
| Neurl2 | 0.537127586 |
| Ergic3 | 0.537079562 |
| Med17 | 0.536291515 |
| Htr7 | 0.536207668 |
| Rnaseh2b | 0.535718994 |
| C2cd5 | 0.535611348 |
| Arnt2 | 0.534888611 |
| Sri | 0.534744974 |
| Asxl2 | 0.534419149 |
| Lgals3 | 0.534291461 |
| Jun | 0.534246224 |
| Ski | 0.533839944 |
| Taf2 | 0.533724558 |
| Tjp1 | 0.533652276 |
| Cnot6l | 0.533483353 |
| Cep68 | 0.533400834 |
| Stk4 | 0.533177313 |
| Cstf3 | 0.532963154 |
| Ak2 | 0.532890469 |
| Ndufaf6 | 0.532865691 |
| Megf10 | 0.532703527 |
| Ybx1 | 0.531980634 |
| Eif3k | 0.531555778 |
| Tmem219 | 0.53136368 |
| Elp3 | 0.531244759 |
| Taf3 | 0.53115528 |
| Herpud1 | 0.530673042 |
| Oip5 | 0.530634966 |
| Gpc2 | 0.530605249 |
| Lrig2 | 0.530565108 |
| Ppp2r5a | 0.530402289 |
| Col18a1 | 0.529778879 |
| Calm3 | 0.529215738 |
| Sh2b2 | 0.528945017 |
| Blvra | 0.528820991 |
| Nol8 | 0.528269379 |
| Vps37d | 0.528059467 |
| Ccng1 | 0.527771795 |
| Ankrd10 | 0.527441328 |
| Gmeb1 | 0.527378054 |
| Slx4ip | 0.527326364 |
| Ptpre | 0.526844212 |
| Acyp2 | 0.526529792 |
| C1qbp | 0.525874763 |
| Rpl13 | 0.52557032 |
| Phf10 | 0.524992714 |
| Spdl1 | 0.524039678 |

| | |
|---|---|
| Sobp | 0.523928789 |
| Pnkp | 0.5231732 |
| Hyal2 | 0.522848629 |
| Dsn1 | 0.52222403 |
| Slc25a35 | 0.521994058 |
| Ampd3 | 0.521973788 |
| Hmgn3 | 0.521963556 |
| Phykpl | 0.521948665 |
| Cpne3 | 0.521759215 |
| Acsm3 | 0.521671367 |
| Pkia | 0.521605795 |
| Slc3a2 | 0.520849478 |
| Smim20 | 0.519973683 |
| Zfhx2 | 0.519288068 |
| Mapkapk3 | 0.519116504 |
| Usp1 | 0.518758903 |
| Ptprr | 0.518287632 |
| Wdhd1 | 0.517732133 |
| Cab39 | 0.517469839 |
| Clp1 | 0.517402639 |
| Rsc1a1 | 0.517287469 |
| Il2rb | 0.517199607 |
| Abhd10 | 0.517144985 |
| Npepps | 0.517026974 |
| Wwp1 | 0.516155633 |
| Smu1 | 0.515941977 |
| Htt | 0.515485771 |
| Zscan22 | 0.515124335 |
| Phf23 | 0.514914494 |
| Tmcc2 | 0.514791265 |
| Mettl20 | 0.514772463 |
| Nacc1 | 0.514304668 |
| Tmem217 | 0.513775153 |
| Smg5 | 0.513220246 |
| Arhgef16 | 0.512680317 |
| Slc1a5 | 0.51262579 |
| Cog7 | 0.512365182 |
| E2f5 | 0.511344435 |
| Fam76b | 0.511068663 |
| Pcgf5 | 0.511016034 |
| Isg15 | 0.509910195 |
| Heph | 0.509078 |
| Rapgef3 | 0.50898206 |
| Cpxm2 | 0.508934015 |
| A4galt | 0.508881873 |
| Cbx2 | 0.508794483 |
| Tapt1 | 0.508224217 |

| | |
|---|---|
| Ei24 | 0.507388038 |
| Rusc2 | 0.507258209 |
| Ggh | 0.507147875 |
| Snx30 | 0.506672218 |
| Zcchc11 | 0.506616836 |
| Spats2 | 0.506574957 |
| Tmub1 | 0.506532552 |
| Sumo3 | 0.506267951 |
| Pacsin2 | 0.506170617 |
| Tfdp1 | 0.505212702 |
| Ap5b1 | 0.504880351 |
| Adi1 | 0.504195905 |
| Ctage5 | 0.504132033 |
| Cep85l | 0.503994333 |
| Pde9a | 0.50301524 |
| Slc35e3 | 0.502743026 |
| Pfkp | 0.502726788 |
| Zscan18 | 0.502341559 |
| Rbm17 | 0.502116167 |
| Plekhh1 | 0.501916664 |
| Wasf3 | 0.501553976 |
| Ier5 | 0.501335774 |
| Dnajc10 | 0.500990123 |
| Gtdc1 | 0.500902311 |
| Pomgnt1 | 0.500588043 |
| Spag1 | 0.500389576 |
| Nrp1 | 0.500185767 |
| Amigo1 | 0.499847572 |
| Cnot2 | 0.499678844 |
| Lmln | 0.499633978 |
| Plgrkt | 0.499559554 |
| Hsf1 | 0.499388542 |
| Snapc3 | 0.499023845 |
| Unc45a | 0.498882728 |
| Eif2d | 0.498727027 |
| Grb10 | 0.498630899 |
| Slc46a3 | 0.498566721 |
| Tle3 | 0.498401078 |
| Got1 | 0.498015268 |
| Mafg | 0.497975593 |
| Rrp12 | 0.497811127 |
| Trim13 | 0.497665623 |
| Ppp1r7 | 0.497574937 |
| Dazap1 | 0.497427821 |
| Sh3bp1 | 0.497315911 |
| Rab28 | 0.497192791 |
| Selk | 0.496825702 |

| | |
|---|---|
| Mlf2 | 0.496433145 |
| Gtf2ird2 | 0.496253189 |
| Ccdc142 | 0.496192467 |
| Cenpm | 0.495701383 |
| Rcbtb1 | 0.49558132 |
| Xrcc6 | 0.495127932 |
| Plekhg6 | 0.494844113 |
| Nadsyn1 | 0.494841888 |
| Isoc1 | 0.492837828 |
| Cxcl3 | 0.492749832 |
| Cabp4 | 0.492623528 |
| Dtnb | 0.491976546 |
| Shc4 | 0.491909159 |
| Ccdc74a | 0.491589559 |
| Dexi | 0.49142757 |
| Sugt1 | 0.491229777 |
| Evc | 0.490115034 |
| Rapgef6 | 0.489873109 |
| Prdx3 | 0.489304709 |
| Clptm1 | 0.489260462 |
| Clec2d | 0.489204444 |
| Fmo5 | 0.488780596 |
| Frmd3 | 0.48865485 |
| Kctd21 | 0.488373221 |
| Tmed10 | 0.488363934 |
| Pex1 | 0.488361845 |
| Ttll4 | 0.488274663 |
| Rpl8 | 0.487964757 |
| Pigb | 0.487309511 |
| Mybl2 | 0.487128724 |
| Sft2d2 | 0.487100508 |
| Tmem150a | 0.486805266 |
| Lamtor1 | 0.486777434 |
| Dnaja1 | 0.486247966 |
| Dusp19 | 0.48598017 |
| Polr3d | 0.485377928 |
| Alkbh8 | 0.485121967 |
| Nsmaf | 0.484951187 |
| Nmnat2 | 0.483512038 |
| Adcy3 | 0.482943115 |
| Tmem88 | 0.482735914 |
| Psmc5 | 0.482448859 |
| Gatc | 0.481448526 |
| Snph | 0.481116086 |
| Oxsr1 | 0.480883106 |
| Hnrnpll | 0.480794806 |
| Dpy19l3 | 0.480304022 |

| | |
|---|---|
| Uhrf1bp1l | 0.479964799 |
| Rsl1d1 | 0.479877164 |
| Slc38a5 | 0.479449304 |
| Amz2 | 0.4794325 |
| Ugdh | 0.479297297 |
| Mx1 | 0.479199357 |
| Scrib | 0.478993958 |
| Cmc1 | 0.478272889 |
| Aqp3 | 0.478186398 |
| Tax1bp3 | 0.478079266 |
| Parp9 | 0.478058125 |
| Fgfrl1 | 0.477921429 |
| Rpl9 | 0.477815152 |
| Strn4 | 0.477017885 |
| Abcb7 | 0.476605638 |
| Rab24 | 0.476533984 |
| Gabpa | 0.476510139 |
| Neil1 | 0.476288209 |
| Fam169a | 0.475288342 |
| Nav1 | 0.474763834 |
| Dmpk | 0.474089923 |
| Mss51 | 0.473240343 |
| Uba5 | 0.473093016 |
| Cercam | 0.472896492 |
| Git2 | 0.472732667 |
| Eid2 | 0.471608935 |
| Kif26a | 0.471551803 |
| Usp54 | 0.47141064 |
| Mrpl42 | 0.471135579 |
| Utp3 | 0.47104753 |
| E2f8 | 0.470594392 |
| Zdhhc20 | 0.470430283 |
| Cd36 | 0.470153432 |
| Slc16a2 | 0.469799555 |
| Mapk6 | 0.46975576 |
| Hnrnpab | 0.46972833 |
| Ide | 0.469695187 |
| Tmem26 | 0.469370314 |
| Hiatl1 | 0.468207642 |
| Ywhag | 0.467469288 |
| Rnf130 | 0.467300436 |
| Mdc1 | 0.467282502 |
| Gmeb2 | 0.467100033 |
| Lemd2 | 0.467050213 |
| Txn2 | 0.466615698 |
| Snrpf | 0.466501902 |
| Stau1 | 0.466495904 |

| | |
|---|---|
| Inip | 0.466481721 |
| Itga1 | 0.466057513 |
| Kif2c | 0.465044843 |
| Ndnl2 | 0.464346203 |
| Krtcap2 | 0.46362157 |
| Dnajb6 | 0.462540175 |
| Eif4ebp1 | 0.46195093 |
| Gna13 | 0.461914626 |
| Sptlc1 | 0.461714218 |
| Tnrc18 | 0.461272884 |
| Chmp7 | 0.460878676 |
| Wdfy2 | 0.459790277 |
| Hspb11 | 0.45952154 |
| Tmc4 | 0.459454321 |
| Polg2 | 0.45921841 |
| Map7 | 0.459089658 |
| Rufy1 | 0.459065417 |
| Samm50 | 0.458222047 |
| Slc44a4 | 0.458008637 |
| Orc3 | 0.457972238 |
| Evc2 | 0.457661309 |
| Tlk2 | 0.457150386 |
| Shisa3 | 0.457067177 |
| Mapk14 | 0.456426437 |
| Caprin1 | 0.45617136 |
| Sec23ip | 0.456087102 |
| Psmd14 | 0.456037162 |
| Lacc1 | 0.453917515 |
| Pfkfb4 | 0.452751569 |
| Oprl1 | 0.452687373 |
| Pcgf2 | 0.45267105 |
| Sharpin | 0.451782724 |
| Pygo2 | 0.450969928 |
| Ammecr1l | 0.44995679 |
| Tmem147 | 0.449804405 |
| Ccdc93 | 0.449802781 |
| Hspd1 | 0.449075691 |
| Wdr91 | 0.448494012 |
| Wdr70 | 0.448222942 |
| Mrps2 | 0.447989115 |
| Romo1 | 0.447882612 |
| Rab11b | 0.447774816 |
| Thumpd3 | 0.447752542 |
| Mycn | 0.447736378 |
| Tnip1 | 0.447727867 |
| Vamp8 | 0.447631467 |
| Atf7ip2 | 0.447564844 |

| | |
|---|---|
| Dclre1c | 0.447115685 |
| Ccdc149 | 0.447058487 |
| Iqgap2 | 0.446971949 |
| Lman1 | 0.446455925 |
| Itm2a | 0.446414959 |
| Rps9 | 0.44639376 |
| Snn | 0.446341209 |
| Zscan26 | 0.446242066 |
| Mis12 | 0.446111731 |
| Trim47 | 0.445314503 |
| Mansc1 | 0.445228472 |
| Sgtb | 0.444300316 |
| Abhd13 | 0.444131301 |
| Eef2 | 0.443765816 |
| Ankrd55 | 0.443552252 |
| Wipf2 | 0.443461236 |
| Itm2c | 0.443030179 |
| Apbb2 | 0.442834479 |
| Hsd3b7 | 0.442831426 |
| Dmxl2 | 0.442776859 |
| Baz2b | 0.442698917 |
| Nelfa | 0.442591482 |
| Rnf149 | 0.442083156 |
| Eif2b4 | 0.441880501 |
| Eif3b | 0.441858795 |
| Slc12a9 | 0.44163117 |
| Emc6 | 0.441355594 |
| Strip1 | 0.441312801 |
| Kif13b | 0.4408078 |
| Mtrf1l | 0.440755146 |
| Tmem231 | 0.440047478 |
| Bank1 | 0.440024762 |
| Polr1b | 0.439909553 |
| Fermt2 | 0.439804733 |
| Atp6v0d1 | 0.439684983 |
| Pask | 0.439545483 |
| Gmps | 0.438824277 |
| Dock1 | 0.438570396 |
| Mia3 | 0.438493964 |
| Wdr44 | 0.43808594 |
| Sox17 | 0.437709419 |
| Kctd18 | 0.437604817 |
| Sh2b3 | 0.437459844 |
| Rwdd1 | 0.437326766 |
| Ptpdc1 | 0.437103112 |
| G3bp1 | 0.43691012 |
| Rab33b | 0.436814459 |

| | |
|---|---|
| Mboat1 | 0.436696604 |
| Pigh | 0.436457498 |
| Cenph | 0.436363907 |
| Stat1 | 0.436066135 |
| Fbxl3 | 0.435951393 |
| Tcf19 | 0.435943458 |
| Nags | 0.435540105 |
| Samd10 | 0.435432627 |
| Fgf5 | 0.435258836 |
| Smyd4 | 0.435231937 |
| Eif3a | 0.435188816 |
| Tmem245 | 0.435150966 |
| Tmem198b | 0.4349895 |
| Alg10b | 0.434988068 |
| Mkln1 | 0.434977301 |
| Neo1 | 0.434564101 |
| Hspa1l | 0.4338202 |
| Gltscr1 | 0.43363555 |
| Msh2 | 0.43358829 |
| Ctr9 | 0.433421662 |
| Fam199x | 0.432944896 |
| Etv3 | 0.432475539 |
| Mier3 | 0.431868412 |
| Prkag1 | 0.431360297 |
| Aste1 | 0.431330469 |
| Ngdn | 0.430936956 |
| Kbtbd8 | 0.430394642 |
| Adam12 | 0.430359836 |
| Fgf16 | 0.430009699 |
| Pou4f1 | 0.429902028 |
| Rnf123 | 0.429531122 |
| Fam155a | 0.429470974 |
| Pdia6 | 0.429055964 |
| Tpp2 | 0.428567859 |
| Abtb1 | 0.428349729 |
| Carf | 0.427005909 |
| Cldn14 | 0.426987483 |
| Mrps15 | 0.426980956 |
| Dram1 | 0.426931329 |
| Rnf103 | 0.426903362 |
| Prkra | 0.425827209 |
| Stoml2 | 0.425823192 |
| Nfatc3 | 0.425236188 |
| Aff2 | 0.424607284 |
| Trex1 | 0.424181233 |
| N4bp3 | 0.42406513 |
| Mmp28 | 0.423911267 |

| | |
|---|---|
| Piwil4 | 0.423753435 |
| Orc5 | 0.423500266 |
| Wbp5 | 0.423446982 |
| Hadhb | 0.423298669 |
| Scamp3 | 0.422732998 |
| Tmco4 | 0.422725607 |
| Arl2bp | 0.422343834 |
| Cfhr1 | 0.422302705 |
| Cc2d1a | 0.421779554 |
| Pcdhb14 | 0.421423863 |
| Exo5 | 0.421117927 |
| Nfil3 | 0.420912902 |
| Mrpl52 | 0.420463597 |
| Ddx10 | 0.420073134 |
| Tacc2 | 0.42006044 |
| Slc40a1 | 0.419907552 |
| Smco4 | 0.419508683 |
| Tnfaip1 | 0.419306442 |
| Fam219b | 0.419228767 |
| Setd1a | 0.419216813 |
| Myh10 | 0.418433962 |
| Mtmr6 | 0.418098872 |
| Tmem141 | 0.418024136 |
| Zkscan4 | 0.417347726 |
| Ubqln4 | 0.416667058 |
| Ccm2l | 0.416451393 |
| Ints9 | 0.416295337 |
| Polr3h | 0.41620106 |
| Mlycd | 0.416038881 |
| Tmem101 | 0.415591235 |
| Tpt1 | 0.415457235 |
| Nufip1 | 0.414910453 |
| Msh5 | 0.4141257 |
| Adck2 | 0.413950957 |
| Dvl3 | 0.413688947 |
| Dus2 | 0.413463009 |
| Tns3 | 0.413231709 |
| Tbccd1 | 0.413129708 |
| Wdr83 | 0.412984344 |
| Cpa3 | 0.412560827 |
| Polr3c | 0.412343112 |
| Pskh1 | 0.412072918 |
| Slc25a13 | 0.411929761 |
| Basp1 | 0.411921212 |
| Rad51d | 0.411110978 |
| Krba1 | 0.411026091 |
| Acox1 | 0.411021053 |

| | |
|---|---|
| Amph | 0.410697202 |
| Apc | 0.410450188 |
| Tfdp2 | 0.409779951 |
| Ino80 | 0.409545684 |
| Ecm1 | 0.409486478 |
| Zbtb14 | 0.409096601 |
| Nab1 | 0.408883766 |
| Ubr1 | 0.408415571 |
| Orai1 | 0.408343227 |
| Dclre1b | 0.407959908 |
| Ncoa7 | 0.407951059 |
| Oraov1 | 0.407939791 |
| Mgst3 | 0.407153645 |
| Atp6v1c2 | 0.407021976 |
| Slc48a1 | 0.406860281 |
| Trim56 | 0.406685973 |
| Dnajc8 | 0.406041239 |
| Art4 | 0.405937819 |
| Med6 | 0.405831793 |
| Ccdc7 | 0.405128061 |
| Rom1 | 0.404119937 |
| Ikbke | 0.403806351 |
| Immp1l | 0.403604791 |
| Cops2 | 0.403562204 |
| Lyrm4 | 0.403169893 |
| Serbp1 | 0.403146184 |
| Gch1 | 0.402451751 |
| Efcab14 | 0.402219905 |
| Zp3 | 0.402206686 |
| Ets1 | 0.402143188 |
| Enkd1 | 0.401821648 |
| Mbtd1 | 0.401712089 |
| Nop14 | 0.40108136 |
| Maml3 | 0.400834254 |
| Naaa | 0.400662745 |
| Fbxl5 | 0.400222262 |
| Tmem218 | 0.400020031 |
| Lyst | 0.399965879 |
| Gnpda2 | 0.399353701 |
| Fos | 0.399352565 |
| Scarb1 | 0.399107076 |
| Ache | 0.398842815 |
| Ulbp1 | 0.397934162 |
| Hist2h2ac | 0.397813895 |
| Nos3 | 0.397616514 |
| Ccdc153 | 0.397036214 |
| Zranb3 | 0.396400548 |

| | |
|---|---|
| Senp8 | 0.396032056 |
| Blvrb | 0.395564702 |
| Tyw5 | 0.395503627 |
| Prc1 | 0.395461422 |
| Rnf122 | 0.394817578 |
| Spryd7 | 0.394280799 |
| Gtf3c4 | 0.392916272 |
| Tab1 | 0.392878988 |
| Nipal3 | 0.392583129 |
| Limch1 | 0.392431387 |
| Nlgn3 | 0.392383575 |
| Ang | 0.391910258 |
| Tmem183a | 0.391792316 |
| Tfeb | 0.391717163 |
| Prpf40a | 0.391450986 |
| Cd151 | 0.391004297 |
| Tmem203 | 0.390874837 |
| Brpf1 | 0.390351038 |
| Chchd3 | 0.390236345 |
| Rae1 | 0.390092443 |
| Mzf1 | 0.389169801 |
| Myo1e | 0.389006724 |
| Lsmem1 | 0.388754284 |
| Snx18 | 0.388733848 |
| Ssx2ip | 0.388515966 |
| Phex | 0.388425015 |
| Cxxc5 | 0.388091111 |
| Slc22a17 | 0.387463068 |
| Apaf1 | 0.387083119 |
| Myo1d | 0.386313351 |
| Itpripl2 | 0.385530389 |
| Myo10 | 0.384938016 |
| Ephb6 | 0.384804486 |
| B3gat1 | 0.384730735 |
| Utrn | 0.384222896 |
| Ttc14 | 0.384203969 |
| Sdpr | 0.383790707 |
| Slc25a20 | 0.383394188 |
| Nrg2 | 0.38293717 |
| Tspan4 | 0.382929423 |
| Nif3l1 | 0.382664539 |
| Hexb | 0.382058146 |
| Mrgbp | 0.381886919 |
| Hspa1a | 0.381865411 |
| Esrra | 0.381174686 |
| Gata6 | 0.381066055 |
| Pigg | 0.380880321 |

| | |
|---|---|
| Scarf2 | 0.380808941 |
| Fam173a | 0.380742116 |
| Dnal4 | 0.379954212 |
| Rprd2 | 0.379386731 |
| Mthfd1l | 0.378672902 |
| Adss | 0.378509431 |
| Paip2 | 0.378240017 |
| Ggct | 0.377273915 |
| Slc2a1 | 0.377215988 |
| Tmem55b | 0.376029323 |
| Tmem120b | 0.375765379 |
| Acer3 | 0.375550469 |
| Bpnt1 | 0.375257643 |
| Fam134b | 0.374932293 |
| Acat1 | 0.374917407 |
| Slc7a6os | 0.374352251 |
| Dnpep | 0.373637163 |
| Tnxb | 0.373452751 |
| Ifrd1 | 0.373385328 |
| Znhit3 | 0.37302094 |
| Kcnab2 | 0.372754405 |
| Epas1 | 0.372683831 |
| Nudcd3 | 0.37222138 |
| Nfe2l3 | 0.372037434 |
| Mak16 | 0.371929244 |
| Commd1 | 0.37167913 |
| Paqr8 | 0.371634514 |
| Acsl5 | 0.371004118 |
| Slc31a1 | 0.370642552 |
| Uxt | 0.369953806 |
| Otub2 | 0.369472379 |
| Ltbp2 | 0.369204982 |
| Chst11 | 0.369171852 |
| Mxd3 | 0.36914207 |
| Top1 | 0.369069903 |
| Lmbrd2 | 0.369029117 |
| Dennd1a | 0.368672479 |
| Cnot8 | 0.367943785 |
| Fem1a | 0.367483639 |
| Mmp16 | 0.367186678 |
| Rab3d | 0.367079657 |
| Hs6st2 | 0.366378248 |
| Ung | 0.366065167 |
| Syap1 | 0.365738872 |
| Tox4 | 0.365390802 |
| Fam64a | 0.365353201 |
| Rnf150 | 0.364549386 |

| | |
|---|---|
| Sulf1 | 0.364529278 |
| Pink1 | 0.363945539 |
| Tmem53 | 0.363752138 |
| Pafah1b2 | 0.363699937 |
| Fn3krp | 0.363471886 |
| Acta2 | 0.36278166 |
| Atp6v0b | 0.362656635 |
| Ttc9c | 0.36263317 |
| Pkd1 | 0.362620489 |
| Fam98a | 0.361777917 |
| Dcp1a | 0.361684229 |
| Nr2f6 | 0.361659694 |
| Sspn | 0.361556623 |
| B4galnt4 | 0.36131539 |
| Fut4 | 0.361040659 |
| Chpf2 | 0.360600175 |
| Tubgcp5 | 0.360309219 |
| Cluh | 0.36013724 |
| Noc3l | 0.360056506 |
| Usmg5 | 0.359252653 |
| Unc119 | 0.358626737 |
| Btbd3 | 0.358194137 |
| Btk | 0.358141179 |
| Zbtb9 | 0.357530099 |
| Efr3b | 0.357424866 |
| Qser1 | 0.357100618 |
| Exosc8 | 0.357066925 |
| Rnf144b | 0.356966265 |
| Wdr12 | 0.356947789 |
| Ago3 | 0.356143083 |
| Ist1 | 0.355816656 |
| Fsd1l | 0.355575113 |
| Xrcc2 | 0.355471812 |
| Paqr4 | 0.355439612 |
| Myh9 | 0.35528813 |
| Foxl1 | 0.355258147 |
| Zbed3 | 0.354563663 |
| Sf3a3 | 0.354301694 |
| Rgs20 | 0.354285697 |
| Rpl37 | 0.354197161 |
| Kif7 | 0.353535987 |
| Igsf8 | 0.353420586 |
| Usp32 | 0.353019121 |
| Brpf3 | 0.352925195 |
| Psmg1 | 0.352790694 |
| Cox19 | 0.352378784 |
| Mark1 | 0.352295971 |

| | |
|---|---|
| Cpd | 0.351663754 |
| Acap2 | 0.351599796 |
| Lpar6 | 0.351518037 |
| Col4a1 | 0.350982456 |
| Rab30 | 0.350717491 |
| Chchd10 | 0.349989308 |
| Tube1 | 0.349635253 |
| Ccdc91 | 0.349203907 |
| Sco2 | 0.348841262 |
| Tnik | 0.348826819 |
| Vps51 | 0.348800538 |
| Ppig | 0.348624805 |
| Rwdd2b | 0.347604983 |
| Lman2l | 0.346966665 |
| Hcls1 | 0.345674263 |
| Sord | 0.345472608 |
| Arl6ip4 | 0.345164949 |
| Tceb1 | 0.345013699 |
| N4bp1 | 0.344853288 |
| Fen1 | 0.344764474 |
| Irf2bp1 | 0.344489388 |
| Negr1 | 0.34429053 |
| Rpsa | 0.343886191 |
| Vegfa | 0.343426732 |
| Fmnl3 | 0.343268425 |
| Slmap | 0.342486387 |
| Psmd8 | 0.342065314 |
| Mrps5 | 0.342037116 |
| Rpl3 | 0.341683567 |
| Aif1l | 0.341579056 |
| Stx17 | 0.340320227 |
| Exd2 | 0.340260693 |
| Gca | 0.339362678 |
| Dctn1 | 0.338496509 |
| Sell | 0.338436426 |
| Arhgef6 | 0.338287798 |
| Rnf145 | 0.338222128 |
| Mad2l2 | 0.338142782 |
| Gcnt4 | 0.337164762 |
| Clic3 | 0.337103328 |
| Scn3b | 0.336633284 |
| Spata5 | 0.33625929 |
| Kif21b | 0.33608258 |
| Lat2 | 0.335671488 |
| Clec3b | 0.335156463 |
| Tom1l1 | 0.334634361 |
| Rab4a | 0.33363355 |

| | |
|---|---|
| Oxr1 | 0.333094889 |
| Stap2 | 0.332831115 |
| Ptpn21 | 0.332576994 |
| Krt8 | 0.332390869 |
| Nfx1 | 0.332368883 |
| Slco1c1 | 0.332319502 |
| Shmt2 | 0.33231882 |
| Ccdc77 | 0.3321142 |
| Snx17 | 0.332037088 |
| Tob2 | 0.331990174 |
| Hoxc6 | 0.33176761 |
| Hbs1l | 0.331656217 |
| Gucy1a3 | 0.331243975 |
| Fas | 0.331189285 |
| Wdfy4 | 0.3309049 |
| Etf1 | 0.330430289 |
| Pgam1 | 0.330424144 |
| Impa1 | 0.3299784 |
| Fbxo10 | 0.328632876 |
| Kansl1 | 0.328309537 |
| Cyr61 | 0.327497288 |
| Fras1 | 0.327344459 |
| Tonsl | 0.326975963 |
| Myo9a | 0.326717344 |
| Rapgef5 | 0.326619745 |
| Arv1 | 0.326361377 |
| Abca9 | 0.326053091 |
| Cinp | 0.325064817 |
| Ufsp1 | 0.324539886 |
| Agpat2 | 0.324517177 |
| Asns | 0.324474046 |
| Cmtr2 | 0.324359903 |
| Brix1 | 0.324139211 |
| Ptprs | 0.324067519 |
| Dok3 | 0.323793932 |
| Tmem230 | 0.323347085 |
| Rin3 | 0.323259855 |
| Ltbp4 | 0.321847082 |
| Ankrd7 | 0.32180183 |
| Bbs4 | 0.321672403 |
| Timm21 | 0.321562272 |
| Pdcl | 0.321138042 |
| Ephb3 | 0.320703476 |
| Yme1l1 | 0.320570576 |
| Tyk2 | 0.32050102 |
| Mkrn2 | 0.319699372 |
| Clcn2 | 0.319417471 |

| | |
|---|---|
| Pitpnm3 | 0.319193831 |
| Sumf2 | 0.318279645 |
| Ccsap | 0.317796868 |
| Arf1 | 0.317712496 |
| Prtg | 0.316840935 |
| Inadl | 0.316799463 |
| Limd2 | 0.316618007 |
| Sppl2b | 0.31629119 |
| Mst1 | 0.316280991 |
| Cacfd1 | 0.316235529 |
| Pex14 | 0.315654206 |
| Dok6 | 0.315593157 |
| Syt11 | 0.31553711 |
| Tox | 0.314986143 |
| Armc1 | 0.314689345 |
| Ak4 | 0.314572068 |
| Galnt18 | 0.314266022 |
| Usp9y | 0.314188279 |
| Dhodh | 0.313255179 |
| Rps27 | 0.313246038 |
| Nosip | 0.313235829 |
| A2m | 0.312872071 |
| Impdh2 | 0.312868494 |
| Rasl10a | 0.31270887 |
| Sarnp | 0.31254956 |
| Auts2 | 0.311994517 |
| Mcrs1 | 0.311507453 |
| Ric8b | 0.311382048 |
| Spata13 | 0.311129231 |
| Plekhf1 | 0.311030027 |
| Zscan2 | 0.310606621 |
| Grin2d | 0.310507541 |
| Gdpd1 | 0.310013168 |
| Itgb3bp | 0.309769806 |
| Tfpt | 0.309674891 |
| Phpt1 | 0.309220161 |
| Rpl22 | 0.309197074 |
| Rad18 | 0.308506403 |
| Riok1 | 0.308467109 |
| Nipa1 | 0.308152521 |
| Anapc16 | 0.308030024 |
| Kat6b | 0.307956515 |
| Farp2 | 0.307842816 |
| Nup88 | 0.306854294 |
| Mtap | 0.306748464 |
| Lxn | 0.306640683 |
| Eno3 | 0.306466754 |

| | |
|---|---|
| Mpp5 | 0.305984755 |
| Ctsc | 0.305899204 |
| B9d2 | 0.304899445 |
| Nuak1 | 0.304230731 |
| Pkn2 | 0.304229847 |
| Hoxb3 | 0.304197851 |
| Nomo1 | 0.30408927 |
| Shh | 0.303839217 |
| Camta1 | 0.3036274 |
| Arhgap39 | 0.303433278 |
| Atp9a | 0.3033518 |
| Hspa2 | 0.302855276 |
| Rnf126 | 0.30251465 |
| Fam175b | 0.302290431 |
| Gngt2 | 0.302244635 |
| Mtf2 | 0.302184424 |
| Plcl2 | 0.301810658 |
| Bcl2 | 0.301338558 |
| Gimap7 | 0.301229376 |
| Trim3 | 0.301016187 |
| Pnpt1 | 0.300876047 |
| Sik3 | 0.300662444 |
| Ust | 0.300446774 |
| Stk11ip | 0.299819509 |
| Ing4 | 0.299628802 |
| Snrnp40 | 0.299460697 |
| Smarcc2 | 0.29843999 |
| Lrpprc | 0.2978589 |
| Idh2 | 0.296411777 |
| Nnmt | 0.296142852 |
| Rb1cc1 | 0.296121545 |
| Pik3ip1 | 0.296119245 |
| Sgsm3 | 0.295160174 |
| Bcam | 0.294753921 |
| Gpat2 | 0.294624738 |
| Opa1 | 0.294138345 |
| Ensa | 0.294045989 |
| Ncor1 | 0.293962195 |
| Atp5b | 0.293406836 |
| Tm9sf2 | 0.293013129 |
| Msto1 | 0.29288392 |
| Srp68 | 0.292765332 |
| Usp21 | 0.292618139 |
| Ddx5 | 0.292254232 |
| Hist1h2bn | 0.29195634 |
| Dlg3 | 0.291662567 |
| Grina | 0.291501407 |

| | |
|---|---|
| Cct7 | 0.291436268 |
| Ryr3 | 0.289944009 |
| Cdh11 | 0.289059402 |
| Tigd5 | 0.28874171 |
| Sync | 0.288474689 |
| Col27a1 | 0.288086819 |
| Hnrnpa1 | 0.287862952 |
| Anxa5 | 0.287128479 |
| Plxna3 | 0.286724109 |
| Lyn | 0.286379767 |
| St6galnac6 | 0.286368245 |
| Ddx60 | 0.285834255 |
| Rspry1 | 0.285685949 |
| Ctbs | 0.285632319 |
| Lama5 | 0.285183372 |
| Tgfb1i1 | 0.284597886 |
| Ctsa | 0.283895532 |
| Cd46 | 0.28378277 |
| Srbd1 | 0.283381388 |
| Loxl1 | 0.283203039 |
| Orai3 | 0.282820993 |
| Ilvbl | 0.282124184 |
| Alg3 | 0.282029088 |
| Havcr2 | 0.281386215 |
| Cask | 0.281374833 |
| Ppp1r14a | 0.281010979 |
| Crim1 | 0.280855013 |
| Pkmyt1 | 0.280419871 |
| Cald1 | 0.279826322 |
| Mtif2 | 0.279355139 |
| Nudt1 | 0.279197591 |
| Rbm39 | 0.278326109 |
| Zufsp | 0.278144051 |
| Clmn | 0.278048361 |
| Ddx1 | 0.278015959 |
| Pcmtd2 | 0.277914734 |
| Smagp | 0.277479065 |
| Gcfc2 | 0.276247577 |
| Dgka | 0.275488951 |
| Lancl3 | 0.275085576 |
| Nup210 | 0.274821371 |
| Amh | 0.274740613 |
| Senp3 | 0.274517778 |
| Ctrl | 0.274500619 |
| Ampd2 | 0.274104566 |
| Atp13a2 | 0.27407622 |
| Ap3m2 | 0.273988123 |

| | |
|---|---|
| Reep6 | 0.273924644 |
| Nes | 0.27390885 |
| Ttc19 | 0.2728703 |
| Lrrfip1 | 0.272666995 |
| Serpinb9 | 0.272466048 |
| Ppp1r12b | 0.271552869 |
| Tmem209 | 0.271544329 |
| Bmp1 | 0.271330394 |
| Fam172a | 0.270050146 |
| Mgst1 | 0.269176551 |
| Cyb5d2 | 0.2688055 |
| Trmt61a | 0.268610596 |
| Slco3a1 | 0.268353084 |
| Ankrd37 | 0.267814173 |
| Mrpl23 | 0.267471412 |
| Ugcg | 0.266862135 |
| Bet1l | 0.266282975 |
| Srpx2 | 0.265347107 |
| Pan3 | 0.265206786 |
| Gprc5a | 0.264973774 |
| Abcc6 | 0.264376653 |
| Plekho1 | 0.263955322 |
| Pappa2 | 0.263609314 |
| Zdbf2 | 0.263280349 |
| Suv420h1 | 0.262982874 |
| Dnali1 | 0.262879893 |
| Cbx8 | 0.26266472 |
| Mrpl22 | 0.26257885 |
| Ubr4 | 0.262115348 |
| Apoa1bp | 0.262013627 |
| Sars2 | 0.261944918 |
| Tapbp | 0.261418304 |
| Dtx3l | 0.261295334 |
| Chd2 | 0.261198157 |
| Scx | 0.261128449 |
| Magoh | 0.261089202 |
| Rraga | 0.260385452 |
| Dpagt1 | 0.259931884 |
| Ctsb | 0.258545422 |
| Phldb2 | 0.258399078 |
| Gnrh1 | 0.257979628 |
| Nfkbia | 0.257586869 |
| Zfand5 | 0.257374794 |
| Casp9 | 0.256700188 |
| Nptx1 | 0.256694774 |
| Pprc1 | 0.256667385 |
| Tsr3 | 0.256221242 |

| | |
|---|---|
| Elac1 | 0.256217649 |
| Syne1 | 0.256030632 |
| Pigs | 0.255828399 |
| Ppp1cc | 0.255293259 |
| Rbm19 | 0.255191422 |
| Sorl1 | 0.254847268 |
| Pcbd1 | 0.254099458 |
| Shisa5 | 0.253915336 |
| Sart1 | 0.253205863 |
| Rpp25l | 0.25285355 |
| Cdca2 | 0.252712453 |
| Brms1 | 0.252609295 |
| Hadh | 0.25217633 |
| Nsd1 | 0.251902129 |
| Hist1h4i | 0.251606421 |
| Psme4 | 0.251531245 |
| Prpf39 | 0.251176068 |
| Plekhh3 | 0.250638921 |
| Trmt2a | 0.249808485 |
| Lypla2 | 0.249336432 |
| Casp3 | 0.249142906 |
| Jmjd4 | 0.248404105 |
| Epor | 0.248142234 |
| Srpk1 | 0.24750304 |
| Atp2a3 | 0.247497329 |
| Mfn1 | 0.246969247 |
| Pggt1b | 0.246937333 |
| Prkx | 0.246875484 |
| Mtmr10 | 0.246501921 |
| Cnih4 | 0.246484425 |
| Tmem102 | 0.245759362 |
| Ccdc65 | 0.245007996 |
| Esyt1 | 0.244790558 |
| Ntn4 | 0.244405667 |
| Fam193b | 0.244153672 |
| Rab43 | 0.243700633 |
| Efr3a | 0.243623774 |
| Nsmce1 | 0.243236571 |
| Stk25 | 0.243038212 |
| Pkn3 | 0.24282181 |
| Myo18a | 0.242716408 |
| Trim37 | 0.242530751 |
| Dpp7 | 0.242142434 |
| Gpatch8 | 0.241506715 |
| Cep57l1 | 0.241462578 |
| Rgs12 | 0.240957937 |
| St3gal4 | 0.240500099 |

| | |
|---|---|
| Arhgef19 | 0.240211448 |
| Trnau1ap | 0.239750978 |
| Trps1 | 0.238692959 |
| Narf | 0.238549435 |
| Bbs12 | 0.238282494 |
| Khdrbs1 | 0.237748835 |
| Mxd1 | 0.237707547 |
| Guf1 | 0.237140962 |
| Casz1 | 0.236855017 |
| Ppm1f | 0.236637128 |
| Ogfod3 | 0.236224073 |
| Spg20 | 0.235464209 |
| Tbx18 | 0.234650927 |
| Vps39 | 0.234645784 |
| Klhdc4 | 0.234518879 |
| Selenbp1 | 0.233927304 |
| Tmed5 | 0.233873538 |
| Trim69 | 0.233570574 |
| Pld1 | 0.233449072 |
| Hexdc | 0.233191235 |
| Pnkd | 0.233009242 |
| Cab39l | 0.232958948 |
| Tmem18 | 0.232759864 |
| Cap2 | 0.232053494 |
| Ttc3 | 0.231863837 |
| Myo5a | 0.230696429 |
| Triml2 | 0.230491781 |
| Polb | 0.230310217 |
| Bcl2l13 | 0.229330663 |
| Amn1 | 0.229042771 |
| Bri3bp | 0.229025895 |
| Nup50 | 0.228181249 |
| Stk36 | 0.22784571 |
| Poc1a | 0.226738947 |
| Sars | 0.226341136 |
| Rqcd1 | 0.225898087 |
| Trappc6b | 0.225460348 |
| Sfi1 | 0.225288969 |
| Nfrkb | 0.224890458 |
| Myom2 | 0.224245954 |
| Spast | 0.224175841 |
| Tuba1a | 0.223236555 |
| Rpia | 0.223215806 |
| Pcsk4 | 0.22280167 |
| Cnih3 | 0.222384402 |
| Phlda2 | 0.222068047 |
| Rpf1 | 0.221908728 |

| | |
|---|---|
| Cdh5 | 0.221627031 |
| Hnrnpa2b1 | 0.220721799 |
| Ddx18 | 0.220698322 |
| Slc25a14 | 0.219920576 |
| Aldh2 | 0.219544881 |
| Nudt12 | 0.219433548 |
| Skil | 0.217978712 |
| Zfp82 | 0.217020527 |
| Pias2 | 0.216969256 |
| Vps72 | 0.216436431 |
| Rab3gap1 | 0.216240785 |
| Dars | 0.216054844 |
| Wrb | 0.215960591 |
| Atp5d | 0.215536424 |
| Rcor2 | 0.215377518 |
| Ncaph2 | 0.21493407 |
| Gtpbp1 | 0.214845125 |
| Hpdl | 0.21467235 |
| Pfkl | 0.214584515 |
| Gbas | 0.214434023 |
| Itsn2 | 0.214421099 |
| Tatdn2 | 0.21430371 |
| Sh3bgrl | 0.21394721 |
| Hmg20a | 0.213345561 |
| Ubxn8 | 0.21288874 |
| Clmp | 0.212395453 |
| Pxk | 0.21238256 |
| Rcc1 | 0.21219959 |
| Mcm8 | 0.211994746 |
| Fam168b | 0.211722627 |
| Atp7a | 0.211487642 |
| Cdk13 | 0.211130986 |
| Ndufs5 | 0.210262826 |
| Fkbpl | 0.209825368 |
| Bmp2k | 0.209240012 |
| Pkd2 | 0.20806598 |
| Hmces | 0.207411593 |
| Clpx | 0.2065598 |
| M6pr | 0.206376612 |
| Kcnn2 | 0.206284278 |
| N6amt1 | 0.205836615 |
| Pfkfb2 | 0.20580229 |
| Col3a1 | 0.205776277 |
| Psenen | 0.205693603 |
| Naa16 | 0.205287577 |
| Nek6 | 0.204906549 |
| Prdm11 | 0.204008767 |

| | |
|---|---|
| Gin1 | 0.203920754 |
| Foxred2 | 0.203640259 |
| Igf2bp3 | 0.202860164 |
| Zdhhc5 | 0.202791817 |
| Zkscan7 | 0.20260989 |
| Thoc3 | 0.202411776 |
| Eif3c | 0.202270029 |
| Tbl2 | 0.202224007 |
| Ube2m | 0.201906996 |
| Rad50 | 0.201463468 |
| Rpain | 0.200569156 |
| Utp11l | 0.200416105 |
| Srpx | 0.200123578 |
| Alkbh4 | 0.199897045 |
| Abl2 | 0.198128592 |
| Zdhhc6 | 0.198116314 |
| Syne2 | 0.197992159 |
| Ran | 0.19769793 |
| Nsfl1c | 0.197313402 |
| 42434 | 0.196763328 |
| Cdc25c | 0.19676045 |
| Epm2a | 0.196624813 |
| Ash1l | 0.196433801 |
| Babam1 | 0.195779479 |
| Chek1 | 0.195707634 |
| Mfap3 | 0.195457634 |
| Klhl35 | 0.194196766 |
| Herc6 | 0.194075557 |
| Bod1 | 0.19392212 |
| Gclc | 0.193677399 |
| Dusp14 | 0.193337678 |
| Sqrdl | 0.193294578 |
| Rimklb | 0.192808898 |
| Cd37 | 0.192792155 |
| Arhgap17 | 0.192688179 |
| Ptcd2 | 0.192658715 |
| Kcnj2 | 0.192636724 |
| Usp20 | 0.192525399 |
| Hdac4 | 0.191671047 |
| Wasf2 | 0.191656043 |
| Rasa2 | 0.191361874 |
| Irak4 | 0.191341955 |
| Ndufc2 | 0.191120166 |
| Opa3 | 0.190727249 |
| Fam132b | 0.190591108 |
| Il1b | 0.190460129 |
| Bdp1 | 0.190173354 |

| | |
|---|---|
| Ddah2 | 0.189654994 |
| Mtfr1 | 0.189426769 |
| Ifngr1 | 0.189073321 |
| Plekha6 | 0.188808116 |
| Adora2b | 0.18816384 |
| Tmem63a | 0.187760947 |
| Slc12a7 | 0.187005501 |
| Ispd | 0.18684291 |
| Ddx50 | 0.186488193 |
| Serinc1 | 0.186347045 |
| Fmr1 | 0.186119767 |
| Dock4 | 0.185911119 |
| Ydjc | 0.185196868 |
| Rab26 | 0.18495271 |
| Cyb5b | 0.184415967 |
| Polr1c | 0.183597676 |
| Tbc1d19 | 0.181301871 |
| Egfr | 0.181190887 |
| Prpf38b | 0.180993911 |
| Rnf135 | 0.180806979 |
| Uqcrc1 | 0.18064072 |
| Grwd1 | 0.180254698 |
| Trpc4ap | 0.18019323 |
| Prkcz | 0.180182293 |
| Nop2 | 0.180121507 |
| Eif5 | 0.17996867 |
| Lrp1 | 0.17994635 |
| Rftn2 | 0.179558766 |
| Fcer1g | 0.17929091 |
| Ap4m1 | 0.179028317 |
| Ipmk | 0.178623219 |
| Necap2 | 0.178479589 |
| Phospho2 | 0.178400789 |
| Ncoa2 | 0.177920838 |
| Slc7a1 | 0.17749335 |
| Pex16 | 0.177433767 |
| Fbxl19 | 0.177078572 |
| Psmd4 | 0.176700782 |
| Epn2 | 0.176004397 |
| Ints7 | 0.175899626 |
| Sfxn5 | 0.175152008 |
| Notch4 | 0.174742016 |
| Cbr4 | 0.174605118 |
| Mis18bp1 | 0.174165831 |
| Sgip1 | 0.172260914 |
| Pdik1l | 0.172109082 |
| Cul3 | 0.172070251 |

| | |
|---|---|
| Fes | 0.171093483 |
| Pigc | 0.170757577 |
| Hoxd9 | 0.170727024 |
| Ttk | 0.170632183 |
| Ptpn1 | 0.170321787 |
| Gigyf1 | 0.170251282 |
| Txndc16 | 0.169936544 |
| Bcor | 0.169911032 |
| Cenpe | 0.169497263 |
| Arid1b | 0.167913908 |
| Gng5 | 0.167882218 |
| Enpp2 | 0.167818343 |
| Serpind1 | 0.167584017 |
| Cacybp | 0.167570255 |
| Riok2 | 0.167108319 |
| Nova1 | 0.16702616 |
| Mmel1 | 0.166949685 |
| Tmem41a | 0.1661171 |
| Pspn | 0.166035142 |
| Serinc3 | 0.165139235 |
| Ecsit | 0.165075675 |
| Adssl1 | 0.164550645 |
| Cdkn1a | 0.164391477 |
| Adck4 | 0.164362717 |
| Smarca2 | 0.164312117 |
| Cops3 | 0.164049471 |
| Bop1 | 0.163701549 |
| Adam32 | 0.163686797 |
| Ptms | 0.162236871 |
| Arg2 | 0.161999282 |
| Kctd12 | 0.161480744 |
| Abi3bp | 0.160871293 |
| Khdrbs3 | 0.160843058 |
| Pip4k2b | 0.160770659 |
| Rsu1 | 0.160135995 |
| Lypd1 | 0.160025355 |
| Sec16a | 0.160022808 |
| Crip2 | 0.16000688 |
| Clk1 | 0.159793617 |
| Tk1 | 0.159546745 |
| Ddah1 | 0.159288657 |
| Mrfap1 | 0.159284009 |
| Rasal2 | 0.159239203 |
| Csrnp2 | 0.158549734 |
| Greb1l | 0.157782137 |
| Brsk2 | 0.157479881 |
| Asun | 0.157388941 |

| | |
|---|---|
| Spns1 | 0.157150406 |
| Ifitm10 | 0.156819748 |
| Lrp8 | 0.15677862 |
| Zc3hc1 | 0.156708331 |
| Agrn | 0.15669446 |
| Gspt1 | 0.156644231 |
| Pde4d | 0.156545084 |
| Tbrg4 | 0.155878981 |
| Ppa2 | 0.155821645 |
| Dyrk2 | 0.155534016 |
| Pinx1 | 0.155490417 |
| Txnl4a | 0.155271303 |
| Abl1 | 0.155031979 |
| Rpl15 | 0.154408757 |
| Lasp1 | 0.154304764 |
| Adm | 0.154183679 |
| Pex3 | 0.153979363 |
| Farsa | 0.153372118 |
| Tgfb1 | 0.153101829 |
| Vav3 | 0.152063181 |
| Pde12 | 0.152055173 |
| Cep76 | 0.151628627 |
| Fbln7 | 0.151014315 |
| Arid5a | 0.150954795 |
| Tlk1 | 0.150753532 |
| Ntng2 | 0.150743083 |
| Alx1 | 0.150596465 |
| Fam216a | 0.150145757 |
| Arglu1 | 0.149785487 |
| Ttl | 0.149685161 |
| Trappc2l | 0.14950832 |
| Elmo3 | 0.147921401 |
| Slc26a11 | 0.147873041 |
| Cebpb | 0.147683487 |
| Rnf38 | 0.14753755 |
| Vezf1 | 0.147464308 |
| Gtf2a2 | 0.147222482 |
| Cav1 | 0.146165489 |
| Igf2r | 0.145683397 |
| Rps16 | 0.145379948 |
| Adk | 0.145249263 |
| Pcdh12 | 0.145195304 |
| Sco1 | 0.144765834 |
| Mgarp | 0.144187385 |
| Helz | 0.144084123 |
| Hrsp12 | 0.143829546 |
| Srpr | 0.142619907 |

| | |
|---|---|
| Zxdc | 0.141750306 |
| Larp6 | 0.141702072 |
| Ppp1r36 | 0.141539428 |
| Pcnxl4 | 0.140833112 |
| Tfe3 | 0.139962479 |
| Mmab | 0.139678922 |
| Rgs19 | 0.13959827 |
| Ubxn2a | 0.139401369 |
| Fgf12 | 0.138910644 |
| Rtkn2 | 0.138867163 |
| Coq5 | 0.138827671 |
| Mettl8 | 0.138097042 |
| Ilk | 0.136512182 |
| Eps8 | 0.136489463 |
| Clpb | 0.136306253 |
| Brd1 | 0.136101169 |
| Dhx33 | 0.135637077 |
| Pex13 | 0.135488397 |
| Smurf1 | 0.134900124 |
| Col13a1 | 0.134847493 |
| Gmds | 0.134737846 |
| Necab3 | 0.134380359 |
| Urgcp | 0.133978771 |
| Ttc9b | 0.133756847 |
| Golga7b | 0.133736073 |
| Usp10 | 0.133731026 |
| Rpl5 | 0.133558328 |
| Med20 | 0.133543388 |
| Triqk | 0.133303215 |
| Nedd4 | 0.132888687 |
| Chd7 | 0.132690406 |
| Sestd1 | 0.1318403 |
| Apbb1 | 0.131817868 |
| Rgs14 | 0.131643996 |
| Sytl4 | 0.131603673 |
| Gamt | 0.131425022 |
| Neu1 | 0.13129724 |
| Nap1l1 | 0.131046169 |
| Ube4a | 0.130941827 |
| Srl | 0.129259314 |
| Ms4a2 | 0.129105324 |
| Bid | 0.128418243 |
| Lsm1 | 0.128150547 |
| Lag3 | 0.128044281 |
| Fbxo28 | 0.12791656 |
| Hoxa1 | 0.127584149 |
| Btf3l4 | 0.127247803 |

| | |
|---|---|
| Slc25a26 | 0.127247078 |
| Pgrmc1 | 0.12547854 |
| Asrgl1 | 0.124631858 |
| Kctd9 | 0.123680943 |
| Dnajc27 | 0.122185768 |
| Max | 0.122163474 |
| Abhd8 | 0.121752343 |
| Tra2b | 0.12141447 |
| Ncs1 | 0.120879783 |
| Top2a | 0.120538286 |
| Trim2 | 0.120450523 |
| Abhd12 | 0.120348021 |
| Cklf | 0.119343262 |
| Gatad2a | 0.118953854 |
| Tmem171 | 0.118738349 |
| Slc25a4 | 0.118244403 |
| Rassf7 | 0.117461319 |
| Slc25a33 | 0.117165309 |
| Lrrc59 | 0.116477661 |
| Map2k1 | 0.115203785 |
| Eaf1 | 0.115173917 |
| Cblb | 0.114584023 |
| Rapgef2 | 0.114427973 |
| Cwc15 | 0.114250518 |
| Hcfc2 | 0.114133575 |
| Insig2 | 0.113676417 |
| Grap | 0.113512652 |
| Tmem144 | 0.113464312 |
| Fam193a | 0.113015613 |
| Fdps | 0.112776928 |
| Dll1 | 0.111646148 |
| Tpm1 | 0.110776058 |
| Anapc5 | 0.110441612 |
| Kif13a | 0.109809625 |
| Mre11a | 0.10965967 |
| Timm44 | 0.109653056 |
| Psmb6 | 0.108819573 |
| Fam212b | 0.108759204 |
| Phf8 | 0.108371657 |
| Pdpr | 0.108151531 |
| Bace2 | 0.106782534 |
| Cog3 | 0.106760488 |
| Cdc42ep5 | 0.105856948 |
| Nr2c2 | 0.1052109 |
| Hivep1 | 0.105182322 |
| Smim19 | 0.105157865 |
| Tmem260 | 0.104730276 |

| | |
|---|---|
| Rpl10 | 0.104596122 |
| Nrf1 | 0.10334006 |
| Pttg1ip | 0.103189041 |
| Ndufaf2 | 0.103114952 |
| Lrrc17 | 0.103031375 |
| Fam161b | 0.102987061 |
| Rgs5 | 0.102958763 |
| Ppp1ca | 0.102300449 |
| Rps6kc1 | 0.101638754 |
| Lars | 0.10151495 |
| Tbc1d16 | 0.101403138 |
| Cep72 | 0.101158236 |
| Anxa11 | 0.101136922 |
| Slc25a24 | 0.100608079 |
| Dok2 | 0.099457913 |
| Dock7 | 0.098636629 |
| Rnf40 | 0.097908853 |
| Samd12 | 0.097724325 |
| Fcgrt | 0.097535527 |
| Zc3h7b | 0.095711749 |
| Vipr1 | 0.095435917 |
| Ndufa1 | 0.09507097 |
| Nhlrc4 | 0.094790021 |
| Slc39a6 | 0.094165137 |
| Rbm42 | 0.094125069 |
| Usp9x | 0.093919305 |
| Polr3k | 0.093740279 |
| Ttf2 | 0.092720849 |
| Mocs3 | 0.092594505 |
| Arhgap22 | 0.092585897 |
| Pex19 | 0.091883959 |
| Asb16 | 0.091493554 |
| Sec22b | 0.091211506 |
| Rhpn1 | 0.091038193 |
| Slc10a3 | 0.090621663 |
| Calcoco1 | 0.089769973 |
| Gps2 | 0.08932443 |
| Ndfip2 | 0.088973832 |
| St6galnac3 | 0.088026591 |
| Rpp30 | 0.087381692 |
| Tmem138 | 0.086287327 |
| Snx24 | 0.085859455 |
| Lyrm1 | 0.085359673 |
| Myo7b | 0.08523723 |
| Por | 0.08480913 |
| Comtd1 | 0.084422795 |
| Mfng | 0.083905784 |

| | |
|---|---|
| Tead1 | 0.083604095 |
| Tmem164 | 0.083476361 |
| Pelp1 | 0.082864608 |
| Arl4a | 0.082589525 |
| Srsf11 | 0.082534234 |
| Mtmr4 | 0.082488625 |
| Cav2 | 0.081558867 |
| Mmaa | 0.081387106 |
| U2af1l4 | 0.081137935 |
| Arhgap31 | 0.081050704 |
| Tcn2 | 0.080211769 |
| Ripk1 | 0.079895274 |
| Lef1 | 0.079624199 |
| Trim16 | 0.078332479 |
| Gpr173 | 0.078225989 |
| Cbfb | 0.07815738 |
| Pdgfd | 0.076586405 |
| Tma7 | 0.076560828 |
| Mxra8 | 0.07639928 |
| Lpar5 | 0.076321884 |
| Tor1aip2 | 0.076104376 |
| Ankfy1 | 0.075342589 |
| Cfp | 0.075308197 |
| Cdadc1 | 0.074659398 |
| Nf1 | 0.074649715 |
| Txnl4b | 0.074559174 |
| Ppp1r13l | 0.074105091 |
| Nrbp2 | 0.073940103 |
| Plce1 | 0.073766136 |
| Rsbn1 | 0.072632972 |
| Pdxp | 0.072169392 |
| Hist1h4h | 0.071893506 |
| Zyx | 0.07187309 |
| Rpgr | 0.071273074 |
| Abcg1 | 0.070781244 |
| Ahi1 | 0.070465963 |
| C4a | 0.069850867 |
| Cdc42 | 0.069764008 |
| Mylk4 | 0.069739866 |
| Dock10 | 0.069506872 |
| Bend3 | 0.069000556 |
| Trappc11 | 0.068518772 |
| Foxm1 | 0.067965931 |
| Dcbld2 | 0.066544101 |
| Als2cl | 0.066444128 |
| Gemin5 | 0.066432512 |
| Chkb | 0.065839337 |

| | |
|---|---|
| Lsm14b | 0.065741365 |
| Fat3 | 0.065051185 |
| Elmo1 | 0.064938233 |
| Snupn | 0.064359794 |
| Smco3 | 0.064152962 |
| Ndp | 0.063848214 |
| Capg | 0.063521844 |
| Ubr5 | 0.063302339 |
| Wnt2b | 0.063231462 |
| Tiam2 | 0.062992769 |
| Dhx29 | 0.062796294 |
| Smpd2 | 0.0625116 |
| Rpl36al | 0.06249362 |
| Gpbp1 | 0.062411839 |
| Nr6a1 | 0.061811026 |
| Hypk | 0.061572815 |
| Cse1l | 0.061021402 |
| Eif2ak4 | 0.060987381 |
| Phf1 | 0.06087351 |
| Aurkb | 0.060092448 |
| Hoxa5 | 0.059227266 |
| Txlna | 0.058615785 |
| Hps5 | 0.057971133 |
| Mff | 0.057324992 |
| Fsbp | 0.057162372 |
| Kif1c | 0.056447208 |
| Pof1b | 0.055959413 |
| Lig3 | 0.055838703 |
| Kctd15 | 0.055268203 |
| Mocs1 | 0.055089556 |
| Plin2 | 0.054469981 |
| Osgin2 | 0.054060006 |
| Trmt1l | 0.054047656 |
| Rlf | 0.054005026 |
| Ccny | 0.053155458 |
| Nme5 | 0.052437093 |
| Ift52 | 0.052323413 |
| Plxnb2 | 0.051860741 |
| Dctn3 | 0.051136509 |
| Myof | 0.050406317 |
| Magi3 | 0.050393886 |
| Atad2b | 0.04967921 |
| Vegfb | 0.049440228 |
| Npc1 | 0.048207913 |
| Atxn7l3b | 0.048181009 |
| Fam98b | 0.047991723 |
| Vwa5a | 0.047003395 |

| | |
|---|---|
| Gramd4 | 0.046917062 |
| Egln3 | 0.046711987 |
| Adar | 0.045970214 |
| Apba2 | 0.045674307 |
| Rtn4 | 0.04535325 |
| Knop1 | 0.044791536 |
| Parl | 0.043315839 |
| Tmem14c | 0.042180322 |
| Sesn1 | 0.042166504 |
| Mtrr | 0.041970853 |
| Zc2hc1c | 0.041775223 |
| Cdk19 | 0.041693929 |
| Ica1 | 0.041280104 |
| Mutyh | 0.040835178 |
| Ebi3 | 0.040511837 |
| Rpl29 | 0.040214666 |
| Lsm8 | 0.039989389 |
| Trim38 | 0.039935726 |
| Hmox2 | 0.039349048 |
| Irs2 | 0.039186715 |
| Dhrs4 | 0.038786019 |
| Wscd1 | 0.038756764 |
| Mib1 | 0.038569581 |
| Nfkb2 | 0.038410804 |
| Hinfp | 0.038380359 |
| Abhd3 | 0.037296059 |
| Tbc1d15 | 0.036767564 |
| Shprh | 0.03670761 |
| Pim1 | 0.036595491 |
| Zmat1 | 0.035472764 |
| Ddx24 | 0.035416643 |
| Cfi | 0.034961707 |
| Gfm2 | 0.034713508 |
| Fignl2 | 0.034345123 |
| Pth1r | 0.034159279 |
| Pak3 | 0.033301551 |
| Cd9 | 0.03328158 |
| Cacng8 | 0.032930532 |
| Zfp14 | 0.032703293 |
| Esco2 | 0.032603332 |
| Brat1 | 0.031740119 |
| E2f6 | 0.030855476 |
| Lrrc8c | 0.030789547 |
| Sbf1 | 0.03047282 |
| Tceal3 | 0.030223729 |
| Pm20d2 | 0.030050801 |
| Eif4h | 0.029834567 |

| | |
|---|---|
| Csf2rb | 0.02967578 |
| Mr1 | 0.029359425 |
| Ggcx | 0.029000306 |
| Hes1 | 0.02894867 |
| Magt1 | 0.028845121 |
| Macrod1 | 0.02883167 |
| Ppp4r1 | 0.028182105 |
| Ifih1 | 0.028133479 |
| Rmnd5b | 0.027969283 |
| Irf2bpl | 0.027858623 |
| Zbtb25 | 0.02748691 |
| Ctsl | 0.027103102 |
| Spata6 | 0.026696888 |
| Psmg2 | 0.026135641 |
| Prr14l | 0.025873293 |
| Mettl13 | 0.025741761 |
| Zfyve9 | 0.025177562 |
| Aldh5a1 | 0.024090209 |
| Cmc2 | 0.023842113 |
| Btaf1 | 0.023768811 |
| Mex3c | 0.023347086 |
| Pmp22 | 0.023258342 |
| Arfip1 | 0.023203719 |
| Nanos1 | 0.023007187 |
| Ddt | 0.021482 |
| Lima1 | 0.021378129 |
| Zc3h14 | 0.021101576 |
| Uqcc1 | 0.020916209 |
| Rgs10 | 0.020248886 |
| Ptk2b | 0.020198496 |
| Dynlt3 | 0.020117162 |
| Ifi27 | 0.020040203 |
| Ghdc | 0.019663081 |
| Lrr1 | 0.019300229 |
| Cyb5r2 | 0.019279381 |
| Neu3 | 0.018685147 |
| Daglb | 0.018672811 |
| Wdr81 | 0.018602874 |
| Rela | 0.018184792 |
| Tcea2 | 0.017917535 |
| Pus10 | 0.016479898 |
| Micu3 | 0.016382567 |
| Coro7 | 0.016029383 |
| Rbm7 | 0.015587629 |
| Rbm33 | 0.015568952 |
| Ehd4 | 0.014956394 |
| Fam196b | 0.014955673 |

| | |
|---|---|
| Cdc5l | 0.014893371 |
| Peg10 | 0.014846015 |
| Gdf11 | 0.014175386 |
| Ubl5 | 0.014137675 |
| Mrpl11 | 0.013936685 |
| Cdc42bpg | 0.013822601 |
| Pnrc1 | 0.012773892 |
| Thada | 0.012759125 |
| Tpd52l1 | 0.012582022 |
| Itpa | 0.012502024 |
| U2af1 | 0.012210619 |
| Syt9 | 0.012169052 |
| Cyb5r4 | 0.012007678 |
| Catsper1 | 0.011320642 |
| Etnk2 | 0.010947757 |
| Nr2c1 | 0.009876704 |
| Sarm1 | 0.009612376 |
| Rfx5 | 0.008202838 |
| Rasgef1b | 0.00808794 |
| Fbxl12 | 0.007573209 |
| C1qtnf5 | 0.006991434 |
| Map2k7 | 0.006378912 |
| Gmfg | 0.006195013 |
| Pum2 | 0.005979947 |
| Atp5o | 0.005940951 |
| Timm23 | 0.005727502 |
| Prrx1 | 0.004972911 |
| Dnlz | 0.004610892 |
| Gm2a | 0.004403098 |
| Nfatc2ip | 0.0040745 |
| Cyp1a1 | 0.003909724 |
| Ddx39b | 0.003414234 |
| Lrrc8d | 0.002911316 |
| Klhl20 | 0.002523023 |
| Dalrd3 | 0.002159448 |
| Atp8a2 | 0.00186124 |
| Brip1 | 0.001203529 |
| Rlim | 0.001203245 |
| Srsf3 | 0.001188106 |
| Desi1 | 0.001124537 |
| Uxs1 | 0.000666414 |
| Farp1 | 0.000390794 |
| Arih2 | 0.000215125 |
| Hist1h1c | -0.00125385 |
| Clstn1 | -0.001284057 |
| Ptpra | -0.001409786 |
| Slc35e4 | -0.001535755 |

| | |
|---|---|
| Map6d1 | -0.0016437 |
| Bclaf1 | -0.001861415 |
| Cisd2 | -0.001883502 |
| Sap30 | -0.002822431 |
| Lfng | -0.003375472 |
| Lyrm2 | -0.003387583 |
| Kpnb1 | -0.003943315 |
| Mtpn | -0.004007974 |
| Dnm2 | -0.00401758 |
| Sms | -0.004125214 |
| Slc47a1 | -0.004207727 |
| Morc4 | -0.004539066 |
| Iffo1 | -0.004668503 |
| Il1rap | -0.004725626 |
| Morc3 | -0.004955121 |
| Dnajc7 | -0.005490895 |
| Osbpl5 | -0.005572203 |
| Dclk1 | -0.005810612 |
| Ccar2 | -0.005842028 |
| Tcp11l1 | -0.006196754 |
| Eif5b | -0.006423503 |
| St13 | -0.00772061 |
| Creb3l2 | -0.008081843 |
| Csrp2 | -0.008209316 |
| Cdca7 | -0.008539225 |
| Copz2 | -0.008567693 |
| Mapk1ip1l | -0.00914917 |
| Seh1l | -0.009293492 |
| Smarce1 | -0.009563459 |
| Arf3 | -0.009855625 |
| Nme3 | -0.010851483 |
| Camk4 | -0.010951408 |
| Synpo | -0.011101091 |
| Mrpl27 | -0.011332171 |
| Rpl31 | -0.012291278 |
| Cnot1 | -0.012305744 |
| Efhc1 | -0.012351615 |
| Mex3b | -0.013440587 |
| Otud5 | -0.013757166 |
| Lemd3 | -0.014019741 |
| Col9a3 | -0.014248337 |
| Creld1 | -0.014576212 |
| Slc26a1 | -0.014737503 |
| Ybey | -0.01480671 |
| Rsrc2 | -0.014824446 |
| Adam19 | -0.014974762 |
| Ldha | -0.015242191 |

| | |
|---|---|
| Lonp1 | -0.015739882 |
| Nek3 | -0.01616733 |
| Stat6 | -0.016649443 |
| Vim | -0.016674259 |
| Bst1 | -0.016707507 |
| Acot1 | -0.017226324 |
| Lace1 | -0.017259427 |
| Wdr13 | -0.017621954 |
| Ubxn6 | -0.017876576 |
| Rnf8 | -0.017890656 |
| Sesn2 | -0.018359413 |
| Rnf217 | -0.020047008 |
| Fxr2 | -0.020381163 |
| Pnpla2 | -0.020542791 |
| Pla2g16 | -0.020657474 |
| Gna14 | -0.021315779 |
| Ctns | -0.021598197 |
| Cacna1a | -0.021934573 |
| Slc27a3 | -0.023083689 |
| Man1a2 | -0.023195048 |
| Rnf168 | -0.023531436 |
| Trim62 | -0.02455688 |
| Slc37a4 | -0.024627998 |
| Srek1ip1 | -0.024794021 |
| Spata2 | -0.025209584 |
| Plekha4 | -0.025419027 |
| Gpd2 | -0.025496561 |
| Ctu1 | -0.026156483 |
| Nat8l | -0.026623735 |
| Glyctk | -0.026804668 |
| Gins2 | -0.02712434 |
| Wwp2 | -0.02715533 |
| Clhc1 | -0.027540658 |
| Xylb | -0.027738108 |
| Rnf32 | -0.028675108 |
| Saal1 | -0.02885299 |
| Gpsm3 | -0.028863197 |
| Enpp4 | -0.029048087 |
| Exosc7 | -0.029061959 |
| Pigk | -0.029347658 |
| Adap2 | -0.031014168 |
| Rnase1 | -0.031014214 |
| Ndc1 | -0.031918048 |
| Fancc | -0.032405416 |
| Cenpk | -0.032863611 |
| Fam120b | -0.034197393 |
| Sec14l1 | -0.034382061 |

| | |
|---|---|
| Cdkn2aipnl | -0.035012348 |
| Kdm4b | -0.035204672 |
| Cs | -0.036239446 |
| Lyrm5 | -0.036525876 |
| Cdr2 | -0.036584519 |
| Slit3 | -0.03659877 |
| Gpx7 | -0.037307275 |
| Nemf | -0.037314457 |
| Chmp3 | -0.038039612 |
| Ablim3 | -0.038046552 |
| Stk32b | -0.038512956 |
| Ppib | -0.038550468 |
| Sytl3 | -0.038646094 |
| Arhgef3 | -0.03876722 |
| Smarca1 | -0.0390201 |
| Creb1 | -0.039330207 |
| Slc16a7 | -0.039353034 |
| Mitf | -0.040818634 |
| Ctso | -0.042262339 |
| Golga1 | -0.042534793 |
| Jag1 | -0.042790226 |
| Armcx1 | -0.043130942 |
| Alkbh1 | -0.04333397 |
| Ugp2 | -0.043450261 |
| Tomm6 | -0.043576304 |
| Lyrm7 | -0.043930537 |
| Pofut1 | -0.044479473 |
| Tspan7 | -0.044777566 |
| Gal | -0.045111372 |
| Uchl5 | -0.045215867 |
| Lnpep | -0.04541103 |
| Impg2 | -0.045801358 |
| Mmp10 | -0.045982446 |
| Vrk1 | -0.047001903 |
| Cdkn2b | -0.047306363 |
| Actrt3 | -0.04784686 |
| Rbck1 | -0.048741501 |
| Lrrc49 | -0.049330527 |
| Hadha | -0.049469686 |
| Acvr2a | -0.05021515 |
| Csnk1a1 | -0.0504535 |
| Ddx17 | -0.050717273 |
| Plekhm3 | -0.050998588 |
| Fnip1 | -0.051644078 |
| Cd83 | -0.052225139 |
| Fbxo39 | -0.052868612 |
| Fbxl17 | -0.053184716 |

| | |
|---|---|
| Upf2 | -0.053230945 |
| P4ha3 | -0.053496745 |
| Plcg1 | -0.053587597 |
| Ppil2 | -0.053850891 |
| Ndfip1 | -0.054314615 |
| Fam20c | -0.054322824 |
| Radil | -0.05442595 |
| Ppp2r5c | -0.055346712 |
| Gstk1 | -0.055858345 |
| Slc12a5 | -0.055908097 |
| Cct5 | -0.056008502 |
| Bcl9l | -0.056179664 |
| Myl4 | -0.056205132 |
| Tmem91 | -0.056227958 |
| Gnb2 | -0.056751269 |
| Lypla1 | -0.057069591 |
| Grpel1 | -0.057286882 |
| Mef2c | -0.058015259 |
| Aff1 | -0.05832275 |
| Tm2d2 | -0.058668061 |
| Snhg11 | -0.058876919 |
| Ackr3 | -0.060062165 |
| Med25 | -0.060192285 |
| Hnrnpd | -0.06032595 |
| Mcts1 | -0.060743482 |
| Ccdc152 | -0.06099955 |
| Gsdmc | -0.061195653 |
| Tstd2 | -0.061251984 |
| Efna3 | -0.061334299 |
| Fam78a | -0.063114004 |
| Nars | -0.06329094 |
| Gtpbp3 | -0.063537849 |
| Ppp3cb | -0.064071527 |
| Klhl29 | -0.064284534 |
| Srd5a1 | -0.064335854 |
| Mtus1 | -0.06439308 |
| Braf | -0.064496087 |
| App | -0.064922948 |
| Zmynd19 | -0.064935355 |
| Ulk3 | -0.065186927 |
| Spata20 | -0.06558002 |
| Bbs10 | -0.066054443 |
| Rheb | -0.066135823 |
| Exosc3 | -0.066941204 |
| Vip | -0.068328661 |
| Fbxw4 | -0.069027166 |
| Bmpr2 | -0.070455687 |

| | |
|---|---|
| Ncapd2 | -0.070639315 |
| Sycp2 | -0.070754849 |
| Rnpc3 | -0.070944447 |
| Aldh1a1 | -0.072599258 |
| Dopey1 | -0.073740081 |
| Dpm3 | -0.073867471 |
| Rps28 | -0.073883881 |
| R3hdm1 | -0.074143202 |
| Sort1 | -0.074213287 |
| Efcab6 | -0.074241 |
| Nubp2 | -0.074334194 |
| Brwd3 | -0.074363067 |
| Sfxn4 | -0.074489107 |
| Nudt11 | -0.074792707 |
| Naa50 | -0.074849325 |
| Spg11 | -0.074917368 |
| Lmf1 | -0.07496924 |
| Jup | -0.076685127 |
| Slc5a3 | -0.077190343 |
| Reep3 | -0.077434454 |
| Polg | -0.077543258 |
| Nup133 | -0.077703916 |
| Dock11 | -0.078208063 |
| Gli2 | -0.078812795 |
| Hace1 | -0.079888157 |
| Aplf | -0.080354575 |
| Nab2 | -0.080923351 |
| Stom | -0.081960893 |
| Ikzf2 | -0.082239148 |
| Srprb | -0.082305083 |
| Cox7c | -0.083256936 |
| Gpatch11 | -0.083422588 |
| Echdc1 | -0.083429268 |
| Plxnb3 | -0.084122172 |
| Rreb1 | -0.085080903 |
| Pde4b | -0.085432128 |
| Zc3h7a | -0.08683525 |
| Mul1 | -0.087457399 |
| Abcb9 | -0.087529807 |
| Atxn2l | -0.088477142 |
| Ccnb1ip1 | -0.089334967 |
| Tirap | -0.089402092 |
| Tmem220 | -0.09038894 |
| Syt3 | -0.091214547 |
| Cpt2 | -0.091220918 |
| Arhgef18 | -0.091425426 |
| Elp6 | -0.091671118 |

| | |
|---|---|
| Tgfbrap1 | -0.091876765 |
| Orc1 | -0.09187978 |
| Vcl | -0.092611143 |
| Lta4h | -0.093438439 |
| Tle2 | -0.093755385 |
| Gckr | -0.094370394 |
| Rfx2 | -0.096102261 |
| Adcy9 | -0.096340124 |
| Nip7 | -0.096367998 |
| Prkcq | -0.09722621 |
| Amt | -0.098189355 |
| Sh3gl1 | -0.098315616 |
| Klf5 | -0.098583293 |
| Pkn1 | -0.098843061 |
| Cachd1 | -0.101456159 |
| Atg9b | -0.101556414 |
| Mfsd3 | -0.101921475 |
| Mthfd2 | -0.102489613 |
| Adipor2 | -0.102938355 |
| Atp1a1 | -0.10368152 |
| Bysl | -0.103912518 |
| Map4k2 | -0.103979896 |
| Itpripl1 | -0.104148469 |
| Sgk1 | -0.10488829 |
| Pdhx | -0.105115479 |
| Cep95 | -0.105171399 |
| Sos2 | -0.105191762 |
| Ncaph | -0.105354208 |
| Mlh1 | -0.105613813 |
| Sephs1 | -0.105791048 |
| Mfsd2a | -0.105869001 |
| Hsd17b2 | -0.105918337 |
| Cpt1c | -0.10617262 |
| Bbx | -0.106504082 |
| Maneal | -0.106682657 |
| Ghitm | -0.108267757 |
| Arpc4 | -0.108778801 |
| Cirh1a | -0.109518827 |
| Sh3bp5l | -0.109712466 |
| Pttg1 | -0.109837062 |
| Psmd9 | -0.110508428 |
| Slc35e2 | -0.110680729 |
| Ccdc28b | -0.111103045 |
| Tspan9 | -0.11161137 |
| Rnf6 | -0.111807394 |
| Traip | -0.11185022 |
| Tead3 | -0.111960474 |

| | |
|---|---|
| Hn1 | -0.112850656 |
| Ccnt1 | -0.113447909 |
| Hip1r | -0.113577442 |
| Pgm2l1 | -0.113769047 |
| Cntn6 | -0.114370622 |
| Nudt14 | -0.115195422 |
| Dnttip1 | -0.115238095 |
| Fam89b | -0.115321087 |
| Foxo1 | -0.116366073 |
| Masp2 | -0.116772877 |
| Ccdc136 | -0.117237242 |
| Ino80e | -0.117534258 |
| Hist2h2be | -0.117714162 |
| Slc25a44 | -0.118154651 |
| Ubp1 | -0.118494477 |
| Rbbp5 | -0.118887496 |
| Acrbp | -0.119641828 |
| Gareml | -0.119861526 |
| Rhot1 | -0.120061429 |
| Lysmd3 | -0.120249364 |
| Slc29a3 | -0.120533123 |
| Mpdu1 | -0.120547031 |
| Il33 | -0.120987106 |
| Fnbp1l | -0.12163332 |
| Ppm1j | -0.121740226 |
| Ash2l | -0.122175985 |
| Nudt6 | -0.122507823 |
| Park2 | -0.12263589 |
| Hspg2 | -0.122799987 |
| Chst1 | -0.123127283 |
| Nck2 | -0.124269775 |
| Plcxd1 | -0.124709118 |
| Nkiras2 | -0.124988808 |
| Poll | -0.125486745 |
| Gulp1 | -0.125929706 |
| Gbgt1 | -0.126371134 |
| Pde8a | -0.127022645 |
| Ipo7 | -0.127428241 |
| Cox6a1 | -0.127704431 |
| Pyroxd1 | -0.129328298 |
| Rcan1 | -0.129513065 |
| Btbd1 | -0.130036234 |
| Iqub | -0.130351567 |
| Cyp39a1 | -0.130425295 |
| Brox | -0.130740207 |
| Rmi2 | -0.131000507 |
| Lzts2 | -0.13102614 |

| | |
|---|---|
| Polr1e | -0.131355922 |
| Rpe | -0.131667963 |
| Bnip1 | -0.132180481 |
| Sptlc2 | -0.132343595 |
| Iscu | -0.133064611 |
| Akna | -0.133679408 |
| Secisbp2l | -0.134197183 |
| Ldlrad2 | -0.134518525 |
| Rmi1 | -0.134598247 |
| Fabp4 | -0.135017252 |
| Sik2 | -0.135147676 |
| Grb2 | -0.135334325 |
| Adrbk1 | -0.135785115 |
| Hmgn1 | -0.13634164 |
| Hspa14 | -0.136614779 |
| Adck1 | -0.136833884 |
| Mars | -0.137304331 |
| Ift74 | -0.138447326 |
| Atp6v1b2 | -0.13855554 |
| Tmem38b | -0.140671124 |
| Mpc2 | -0.140686325 |
| Stx12 | -0.141488658 |
| Smpd4 | -0.141598797 |
| Tcof1 | -0.141685505 |
| Trim45 | -0.141720777 |
| Dcakd | -0.141748851 |
| Sdhc | -0.142309487 |
| Ttyh2 | -0.142710898 |
| Hmgxb4 | -0.143058521 |
| Frmd5 | -0.143249936 |
| Mfsd11 | -0.144305362 |
| Slc41a2 | -0.144489063 |
| Zfyve16 | -0.14520642 |
| Taf11 | -0.14538492 |
| Cog6 | -0.146229464 |
| Bace1 | -0.146604861 |
| Zfpm2 | -0.146728117 |
| Nenf | -0.147799393 |
| Ndufs2 | -0.148922708 |
| Otud3 | -0.149793226 |
| Aebp2 | -0.150797573 |
| Top3b | -0.152231367 |
| Uqcrq | -0.152274291 |
| Tomm7 | -0.152325941 |
| Tsn | -0.153742291 |
| Zkscan5 | -0.153755763 |
| Ap3s1 | -0.153762397 |

| | |
|---|---|
| Plau | -0.153776089 |
| Fam69b | -0.153943405 |
| Pgs1 | -0.154365167 |
| Poli | -0.155245367 |
| Dip2c | -0.15548081 |
| Map3k6 | -0.155613665 |
| Krt18 | -0.155625903 |
| Ethe1 | -0.156255995 |
| Alg1 | -0.156334632 |
| Vdac1 | -0.156810465 |
| Cox10 | -0.156909809 |
| Arpc2 | -0.157028217 |
| Rfc2 | -0.157591455 |
| Def8 | -0.157847869 |
| Birc2 | -0.157917777 |
| Xrn2 | -0.158260565 |
| Hyou1 | -0.158922419 |
| Hr | -0.159271023 |
| Gcnt2 | -0.159905821 |
| Dnajc5 | -0.16087345 |
| Rngtt | -0.161161731 |
| Glt8d2 | -0.161440386 |
| Cnnm2 | -0.161584076 |
| Surf4 | -0.162066305 |
| Tkt | -0.162866485 |
| Zbtb46 | -0.163017693 |
| Mc1r | -0.163965692 |
| Tmod3 | -0.164163309 |
| Irf1 | -0.164270226 |
| Ndufa8 | -0.164766774 |
| Nid2 | -0.16506825 |
| Prelid2 | -0.165299447 |
| Syngr2 | -0.165469484 |
| Higd1a | -0.166531353 |
| Tmco1 | -0.166665971 |
| Mras | -0.166843425 |
| Fbxo34 | -0.167337459 |
| Npc2 | -0.167551836 |
| Mcmdc2 | -0.167600582 |
| Prkag2 | -0.16770175 |
| Tgs1 | -0.168158456 |
| Kifap3 | -0.168251892 |
| Cars2 | -0.1682653 |
| Ahcyl1 | -0.168309839 |
| Psmf1 | -0.169812707 |
| Endog | -0.170314835 |
| Asb8 | -0.170577524 |

| | |
|---|---|
| Eif2ak1 | -0.170640606 |
| Flii | -0.1716565 |
| Pafah2 | -0.171708395 |
| Ikbkb | -0.17256136 |
| Mdfic | -0.172623793 |
| Slc16a14 | -0.172705343 |
| Ncstn | -0.173024954 |
| Mrpl14 | -0.173270158 |
| Cdc14b | -0.173300573 |
| Rps8 | -0.17411081 |
| Rnf214 | -0.174554621 |
| Spen | -0.174716258 |
| Wls | -0.175307093 |
| Gpkow | -0.17598425 |
| Pycr2 | -0.176528304 |
| Proser2 | -0.17685676 |
| Ppa1 | -0.177363724 |
| Tab2 | -0.177837672 |
| Frmd8 | -0.179233101 |
| C1qtnf1 | -0.180139823 |
| Hars | -0.180642482 |
| Cxadr | -0.181075025 |
| Ccdc68 | -0.181401464 |
| Ctdsp2 | -0.18149941 |
| Prr5l | -0.181597572 |
| Sh3rf1 | -0.181633086 |
| Yy2 | -0.181738696 |
| Spata33 | -0.182186399 |
| Slc37a1 | -0.182720046 |
| Aup1 | -0.182730565 |
| Ggnbp2 | -0.182745558 |
| Clns1a | -0.182763309 |
| Gata2 | -0.183434466 |
| Lmbr1l | -0.18380506 |
| Nlgn1 | -0.18473912 |
| Oas2 | -0.18475735 |
| S100a2 | -0.185035067 |
| Chic1 | -0.18551086 |
| Raver1 | -0.186285523 |
| Trim52 | -0.187124451 |
| Kcnab1 | -0.187610067 |
| Gp6 | -0.187891088 |
| Ccdc88c | -0.189762013 |
| Arel1 | -0.189829173 |
| Cdv3 | -0.190398551 |
| Ept1 | -0.190752728 |
| Tcaim | -0.191223405 |

| | |
|---|---|
| Nmrk1 | -0.191267415 |
| Mbnl3 | -0.19170764 |
| Il21r | -0.193104383 |
| Thnsl1 | -0.19472643 |
| Atxn10 | -0.194792427 |
| Ndufab1 | -0.194889372 |
| Aldh1a2 | -0.195271319 |
| Slc18b1 | -0.196035846 |
| Atpaf1 | -0.196120688 |
| Myl9 | -0.196325809 |
| Ddx58 | -0.196350416 |
| Zbtb2 | -0.19708738 |
| Mrpl30 | -0.19711357 |
| Parp14 | -0.197488139 |
| Acsf3 | -0.198747525 |
| Qtrtd1 | -0.198977062 |
| Trrap | -0.199562758 |
| Rbm15 | -0.20090171 |
| Gal3st4 | -0.201016051 |
| Socs1 | -0.201113982 |
| Bcl2l11 | -0.201255884 |
| Ift20 | -0.20242645 |
| Mipep | -0.202601062 |
| Setmar | -0.202613332 |
| Rangrf | -0.202685854 |
| Zranb1 | -0.202695759 |
| Tesk2 | -0.203204852 |
| Naa10 | -0.203727842 |
| Map1lc3b | -0.203956792 |
| Spryd3 | -0.20406582 |
| Hoxb9 | -0.20420648 |
| Glud1 | -0.204386423 |
| Nudt4 | -0.204941928 |
| Lmf2 | -0.204987224 |
| Polr3b | -0.205526652 |
| Garnl3 | -0.205759967 |
| Ednrb | -0.206070416 |
| Dock3 | -0.207525575 |
| Tle4 | -0.207767086 |
| Vcpip1 | -0.207806407 |
| Lrriq1 | -0.208600802 |
| Thbs3 | -0.209363567 |
| Arhgap26 | -0.209664578 |
| Tmed3 | -0.210503701 |
| Vash1 | -0.210712785 |
| Agfg1 | -0.211069857 |
| Ptpn13 | -0.211119505 |

| | |
|---|---|
| Reps2 | -0.211547433 |
| Tmem14a | -0.211582682 |
| Fads1 | -0.211596032 |
| Mina | -0.214119501 |
| Aes | -0.214328162 |
| Fbxo44 | -0.214892171 |
| Snrpd2 | -0.215259651 |
| Osgepl1 | -0.215327247 |
| Pcm1 | -0.21553791 |
| Dbndd2 | -0.215697532 |
| Ints1 | -0.216577165 |
| Atp5f1 | -0.217002673 |
| Hip1 | -0.217152234 |
| Nbr1 | -0.217887588 |
| Rnf13 | -0.218117562 |
| Vps16 | -0.218486525 |
| Cct4 | -0.219822832 |
| Pgk1 | -0.220288144 |
| Ypel1 | -0.220290577 |
| Derl3 | -0.220300249 |
| Atl2 | -0.220454475 |
| Setbp1 | -0.221158439 |
| Rpl6 | -0.221633887 |
| Fbxo42 | -0.222353965 |
| Tasp1 | -0.222522495 |
| Prmt1 | -0.222939402 |
| Haus8 | -0.223548042 |
| Rassf2 | -0.224293166 |
| Cenpp | -0.224740478 |
| Rpa3 | -0.22499412 |
| Rps25 | -0.225388967 |
| Pls3 | -0.225835312 |
| Reck | -0.228635201 |
| Lepr | -0.2295645 |
| Trappc8 | -0.229872591 |
| Hdac2 | -0.230330494 |
| Supv3l1 | -0.23075683 |
| Fam50a | -0.230930028 |
| Rps26 | -0.231654726 |
| Clint1 | -0.232089657 |
| Eed | -0.232345611 |
| Mob3c | -0.232675527 |
| Gemin7 | -0.233749931 |
| Upp1 | -0.233788234 |
| Kif1b | -0.234608194 |
| Prrc2c | -0.235478231 |
| Mrpl28 | -0.235935085 |

| | |
|---|---|
| B3galnt2 | -0.236299453 |
| Manba | -0.236328454 |
| Lpar2 | -0.23642896 |
| Lnx1 | -0.237046225 |
| Hid1 | -0.237581986 |
| Ston1 | -0.237959966 |
| Polr2h | -0.23808643 |
| Map2k2 | -0.238738512 |
| Kpna4 | -0.238751764 |
| Casd1 | -0.238936706 |
| Tmem168 | -0.239090532 |
| Ccdc178 | -0.240222002 |
| Ppfibp1 | -0.240225008 |
| Gak | -0.24113627 |
| Fam46a | -0.242100994 |
| Map3k8 | -0.242343879 |
| Zbtb34 | -0.243831347 |
| Fam136a | -0.244574856 |
| Ptchd2 | -0.245106054 |
| Fam187a | -0.245406077 |
| Coro1a | -0.245579793 |
| Metap2 | -0.246630336 |
| Zfand6 | -0.246638161 |
| Gramd1c | -0.247735706 |
| Arhgap11a | -0.247784473 |
| Ogfrl1 | -0.248868635 |
| Nudt18 | -0.248970815 |
| Srek1 | -0.250092928 |
| Ebag9 | -0.250472261 |
| Ccdc9 | -0.250600909 |
| Cops8 | -0.251554521 |
| Cntn1 | -0.251983248 |
| Gas1 | -0.252181907 |
| Nlrp2 | -0.253076611 |
| Pyurf | -0.255070033 |
| Skiv2l | -0.255567744 |
| Igfbp3 | -0.255699085 |
| Slc39a7 | -0.256322328 |
| Fam118a | -0.256400513 |
| Kif18a | -0.256987538 |
| Has3 | -0.257570281 |
| Ddx54 | -0.258495249 |
| Syngr1 | -0.259156398 |
| Lsr | -0.262407233 |
| Papolg | -0.262453618 |
| Arsg | -0.263591695 |
| Esam | -0.263617202 |

| | |
|---|---|
| Tarbp2 | -0.264155646 |
| B3gat2 | -0.264316701 |
| Idi1 | -0.264587721 |
| Ormdl3 | -0.264865566 |
| Ctf1 | -0.265900637 |
| Depdc1b | -0.266266988 |
| Dnajc13 | -0.266484464 |
| Mrpl17 | -0.267845819 |
| Paip1 | -0.268576592 |
| Fut1 | -0.268611089 |
| Oat | -0.26861737 |
| Ncoa5 | -0.268729853 |
| Pacrgl | -0.268909999 |
| Stim1 | -0.269498999 |
| Rhog | -0.270060231 |
| Slc25a25 | -0.270544982 |
| Baz1b | -0.270610715 |
| Prdx5 | -0.271368728 |
| Mettl1 | -0.271579892 |
| Mapkapk5 | -0.271707574 |
| Rab40c | -0.272101399 |
| Sybu | -0.273018016 |
| B3galt4 | -0.273190327 |
| P4ha2 | -0.273274449 |
| Ube2g1 | -0.273769415 |
| Cpt1a | -0.274299968 |
| Themis2 | -0.274362135 |
| Tubb2a | -0.274731116 |
| Csk | -0.274781366 |
| Apitd1 | -0.275587022 |
| Numbl | -0.275676836 |
| Lztr1 | -0.275679834 |
| Ubr3 | -0.275702833 |
| Fhl3 | -0.275705718 |
| Rgs4 | -0.276384784 |
| Lhx6 | -0.276480986 |
| Rnmt | -0.277706904 |
| Mrpl21 | -0.278205459 |
| Mllt10 | -0.278941353 |
| Serpini1 | -0.27902095 |
| Leng9 | -0.279547061 |
| Kdm1a | -0.279860647 |
| Tmem150c | -0.279900672 |
| Pcgf1 | -0.279971894 |
| Rac1 | -0.280819408 |
| Lactb2 | -0.281704982 |
| Rccd1 | -0.282307997 |

| | |
|---|---|
| Homer1 | -0.283175147 |
| Ppp2r1a | -0.283363878 |
| Ccdc159 | -0.283673165 |
| Gpcpd1 | -0.283729489 |
| Ltbp3 | -0.284205434 |
| Twf2 | -0.285427618 |
| Mta2 | -0.28576289 |
| Rala | -0.285847612 |
| Ezh1 | -0.285956429 |
| Rps11 | -0.286095991 |
| Gfm1 | -0.28625893 |
| Impad1 | -0.287359386 |
| Capn2 | -0.28797877 |
| Bax | -0.288125006 |
| Edc4 | -0.288634498 |
| Rab18 | -0.288656619 |
| Ceacam1 | -0.28872473 |
| Arid4a | -0.288763468 |
| Maged2 | -0.289010673 |
| Vwa8 | -0.29021048 |
| Vps37a | -0.290534852 |
| Smad4 | -0.292209051 |
| Rpl30 | -0.292666239 |
| Ddx19a | -0.29387051 |
| Slc43a2 | -0.294403896 |
| Tmem5 | -0.294579581 |
| Rasa4 | -0.294807479 |
| Ddit4l | -0.294965154 |
| Dcps | -0.295056082 |
| Bend5 | -0.295362743 |
| Fam179b | -0.295674971 |
| Fdft1 | -0.295700257 |
| Rgs7bp | -0.295872699 |
| Stt3a | -0.296861769 |
| Zdhhc1 | -0.296955507 |
| Gp1bb | -0.297245384 |
| Hsbp1l1 | -0.297287683 |
| Gins1 | -0.298464264 |
| Taf7 | -0.300095776 |
| Lrsam1 | -0.300624352 |
| Snx14 | -0.301164674 |
| Gfod2 | -0.30238589 |
| Tubd1 | -0.302640554 |
| Ccdc167 | -0.303291379 |
| Fam192a | -0.303556074 |
| Gosr1 | -0.305949643 |
| Kpna6 | -0.306395876 |

| | |
|---|---|
| Timmdc1 | -0.306791043 |
| Cir1 | -0.30691967 |
| Tpi1 | -0.308468892 |
| Tmem135 | -0.308521597 |
| Lrrc4 | -0.308884127 |
| Slc25a28 | -0.30895746 |
| Wdr35 | -0.309238938 |
| Etaa1 | -0.310155299 |
| Acad8 | -0.310219681 |
| Gid4 | -0.310405567 |
| Lipe | -0.311643705 |
| Cox15 | -0.311718883 |
| Tmem189 | -0.312045853 |
| Mphosph6 | -0.312057923 |
| Cyth3 | -0.312181368 |
| Nsun6 | -0.312368607 |
| Nat6 | -0.312418737 |
| Slc30a7 | -0.312586707 |
| Oaz2 | -0.312968434 |
| Slc16a6 | -0.312999437 |
| Cers2 | -0.314338068 |
| Ptpn2 | -0.314435689 |
| Phb2 | -0.314544287 |
| Uvrag | -0.315194555 |
| Cpsf6 | -0.315675631 |
| U2surp | -0.315906137 |
| Gmnn | -0.316343044 |
| Setd6 | -0.316644426 |
| Uggt2 | -0.316945083 |
| Pak2 | -0.31728857 |
| Arrdc2 | -0.317389783 |
| Ssh1 | -0.31738983 |
| Heatr5a | -0.317420108 |
| Oas3 | -0.317516396 |
| Rbm8a | -0.317793953 |
| Arf4 | -0.317848255 |
| Abat | -0.31896775 |
| Myo9b | -0.319249827 |
| Sipa1l1 | -0.319625935 |
| Idnk | -0.320183759 |
| Cdc42bpa | -0.321088815 |
| Col6a1 | -0.322835441 |
| Spg7 | -0.323804415 |
| Utp14a | -0.324492969 |
| Pxmp2 | -0.324793296 |
| Acp1 | -0.324925297 |
| Dennd6a | -0.325651449 |

| | |
|---|---|
| Kmt2a | -0.325757873 |
| Topors | -0.326352134 |
| Sdc3 | -0.326707537 |
| Slc38a1 | -0.329070859 |
| Ift43 | -0.330237322 |
| Gdf3 | -0.33030077 |
| Tbrg1 | -0.330837416 |
| Lonrf3 | -0.33105048 |
| Pcnxl2 | -0.331801497 |
| Aplnr | -0.332505234 |
| Errfi1 | -0.333836421 |
| Zbed6 | -0.333853198 |
| Per1 | -0.333918796 |
| 42622 | -0.335229506 |
| Lama3 | -0.335559274 |
| Sh3rf3 | -0.335771239 |
| Dtymk | -0.336752563 |
| Rmdn1 | -0.336917358 |
| Cdc42se2 | -0.338052225 |
| Atg4c | -0.339378538 |
| Inhbb | -0.340253804 |
| Dirc2 | -0.340575771 |
| Arih1 | -0.340981744 |
| Polq | -0.341105439 |
| Col1a1 | -0.341516259 |
| Rspo3 | -0.342699595 |
| Rock1 | -0.342926582 |
| Tnfrsf10b | -0.343778318 |
| Atad5 | -0.34385183 |
| Vkorc1 | -0.343881334 |
| Tubb2b | -0.344714512 |
| Mccc1 | -0.3452092 |
| Prmt7 | -0.346627731 |
| Fam13a | -0.347943972 |
| Troap | -0.348639272 |
| St8sia2 | -0.349200852 |
| Apool | -0.349955883 |
| Podnl1 | -0.350644816 |
| Trim28 | -0.351540815 |
| Hmgcl | -0.351587006 |
| Mvp | -0.351684149 |
| Carm1 | -0.352187903 |
| Tmem59 | -0.353213403 |
| Vps4b | -0.354209325 |
| Myo6 | -0.355486938 |
| Ccser1 | -0.355490486 |
| Fli1 | -0.356545031 |

| | |
|---|---|
| Pcdh17 | -0.356651795 |
| Rtn4ip1 | -0.356690786 |
| Ddx41 | -0.356698616 |
| Ppargc1b | -0.357643956 |
| Syngr3 | -0.358084015 |
| Pip4k2a | -0.358203154 |
| Xxylt1 | -0.358337227 |
| Dbndd1 | -0.359410441 |
| Slc25a23 | -0.359720052 |
| Ralbp1 | -0.359848105 |
| Rltpr | -0.359983761 |
| Tmem37 | -0.360305463 |
| Mgme1 | -0.360554687 |
| Baalc | -0.360908002 |
| Tcerg1 | -0.361339319 |
| Iars | -0.361909329 |
| Sepp1 | -0.362399191 |
| Zmat2 | -0.36265622 |
| Pts | -0.363097714 |
| Creb3l1 | -0.364203485 |
| Vdr | -0.365231662 |
| Ankrd46 | -0.365475749 |
| Ccdc71l | -0.366235714 |
| Hoxb5 | -0.366764203 |
| Synj1 | -0.367025222 |
| Nol11 | -0.367627514 |
| Oxnad1 | -0.36770034 |
| Phf20l1 | -0.368592585 |
| Itga6 | -0.368788275 |
| Spock3 | -0.369952472 |
| Glo1 | -0.371378124 |
| Rbm43 | -0.371988619 |
| Gtf3a | -0.372097346 |
| Lamc2 | -0.37233753 |
| Zeb2 | -0.372782756 |
| Midn | -0.373113808 |
| Fosl1 | -0.373444269 |
| Nppc | -0.37390604 |
| Vit | -0.374171947 |
| Hebp2 | -0.374851253 |
| Ccndbp1 | -0.375626288 |
| Tk2 | -0.375718993 |
| Tmem238 | -0.375893359 |
| Tnip2 | -0.375982627 |
| Smarca5 | -0.376857329 |
| Marveld2 | -0.377049176 |
| Tram2 | -0.37784712 |

| | |
|---|---|
| Smc4 | -0.378214099 |
| Adamts18 | -0.378219353 |
| Dgcr6 | -0.378364806 |
| Gnptg | -0.379047596 |
| Fnbp1 | -0.379528763 |
| Xpr1 | -0.379553019 |
| Pdcd1lg2 | -0.379901251 |
| Mepce | -0.38010194 |
| Matn3 | -0.380567011 |
| Olfml3 | -0.380870569 |
| Exosc2 | -0.381403298 |
| Nudt5 | -0.382263348 |
| Stag3 | -0.38256288 |
| Tcf7l2 | -0.382789424 |
| Atg16l1 | -0.383369072 |
| Adora3 | -0.384306013 |
| Ankrd23 | -0.384379557 |
| Suclg2 | -0.384549887 |
| Myrf | -0.384794165 |
| Ndufs1 | -0.384908247 |
| Gpr143 | -0.386173156 |
| Hmmr | -0.387588663 |
| Clasp2 | -0.38778844 |
| Nynrin | -0.388565234 |
| Clcc1 | -0.389290766 |
| Cyc1 | -0.389319408 |
| Gnptab | -0.391314595 |
| Whsc1l1 | -0.39210564 |
| Sbds | -0.392119611 |
| Pbk | -0.392250379 |
| Clec11a | -0.393013873 |
| Slc11a2 | -0.393646095 |
| Bub1 | -0.393806752 |
| Exosc4 | -0.393923804 |
| Zyg11a | -0.394128018 |
| Wdr90 | -0.394474866 |
| Cenpv | -0.394748653 |
| Gdi2 | -0.395502526 |
| Hibadh | -0.396365549 |
| Lpgat1 | -0.397680775 |
| Zbtb22 | -0.397791902 |
| Commd4 | -0.397979604 |
| Snrpd3 | -0.398591179 |
| Itga11 | -0.39909564 |
| Pla2g4a | -0.400490703 |
| Arl16 | -0.400749683 |
| Ddx52 | -0.401237295 |

| | |
|---|---|
| Nisch | -0.401453525 |
| Fastkd1 | -0.4017252 |
| Bak1 | -0.401756072 |
| Ppp1r26 | -0.402508341 |
| Toe1 | -0.403579441 |
| Eif3m | -0.404095795 |
| Actn1 | -0.404393807 |
| Rabepk | -0.404543479 |
| Yipf1 | -0.405362942 |
| Syncrip | -0.406516628 |
| Cd2ap | -0.407535726 |
| Hivep2 | -0.407660758 |
| Rpf2 | -0.409122142 |
| Dnase1 | -0.409845299 |
| Scfd1 | -0.409941113 |
| Dyrk3 | -0.41094328 |
| Rpl10a | -0.411761013 |
| Cic | -0.412074873 |
| Stab2 | -0.412333011 |
| Slc35f5 | -0.412970953 |
| Map3k4 | -0.413523806 |
| Msl3 | -0.414004543 |
| Tex10 | -0.414230878 |
| Ppp3ca | -0.415509244 |
| Setd3 | -0.415785719 |
| Jade2 | -0.415811599 |
| Bbc3 | -0.416044498 |
| Hmcn1 | -0.416272433 |
| Tmem243 | -0.416862315 |
| Tctn2 | -0.41711298 |
| Rab39b | -0.417786388 |
| Batf3 | -0.417810912 |
| Myef2 | -0.418193438 |
| Arf5 | -0.419029308 |
| Ppp1r21 | -0.41929363 |
| Lclat1 | -0.419301732 |
| Ankef1 | -0.419761512 |
| Ifitm1 | -0.420917328 |
| Sugp1 | -0.421287936 |
| Taf1 | -0.421313917 |
| Lsm7 | -0.421453673 |
| Cend1 | -0.421897785 |
| Cnot6 | -0.422977272 |
| Arl15 | -0.423126186 |
| Shoc2 | -0.423196372 |
| Acot9 | -0.423757895 |
| Tmem52 | -0.423846897 |

| | |
|---|---|
| Smad7 | -0.42396491 |
| Eral1 | -0.424573321 |
| Commd3 | -0.424643647 |
| Cpeb2 | -0.424836578 |
| Znhit2 | -0.425160062 |
| Ccdc15 | -0.425172229 |
| Pcbp2 | -0.425961632 |
| Gdi1 | -0.426687428 |
| Ncdn | -0.426802935 |
| Cog2 | -0.428300699 |
| Hcn2 | -0.428339656 |
| Gpx3 | -0.428499661 |
| Cers5 | -0.428908039 |
| Myl6b | -0.429169313 |
| Vamp2 | -0.429803463 |
| Izumo4 | -0.430016367 |
| Nfxl1 | -0.431412889 |
| Nfkbil1 | -0.431430876 |
| Bcl3 | -0.431461519 |
| Polr3f | -0.431532376 |
| Ppdpf | -0.431744253 |
| Mlst8 | -0.432361411 |
| Tmem63b | -0.433312362 |
| Osbpl9 | -0.433472082 |
| Clstn3 | -0.433679596 |
| St5 | -0.434095323 |
| Cand1 | -0.434100913 |
| Acox3 | -0.434266836 |
| Prrx2 | -0.435586603 |
| Tmem241 | -0.436006664 |
| Abcf3 | -0.43683784 |
| Pgm5 | -0.437210953 |
| Dcun1d1 | -0.437500134 |
| Timm22 | -0.4379043 |
| Fam129a | -0.437992154 |
| Socs4 | -0.439126407 |
| Pex12 | -0.439236627 |
| Rac2 | -0.439572631 |
| Cops6 | -0.439716515 |
| Mdh2 | -0.43989079 |
| Foxj3 | -0.441004046 |
| Iqck | -0.441129227 |
| Stradb | -0.441180105 |
| Zdhhc16 | -0.44178712 |
| Adsl | -0.4419573 |
| Pnp | -0.442197237 |
| Pmch | -0.442540206 |

| | |
|---|---|
| Gpr157 | -0.442893212 |
| Inpp5k | -0.443326312 |
| Dmrta1 | -0.445109525 |
| Slc39a14 | -0.445284577 |
| Ccdc17 | -0.445409036 |
| Prob1 | -0.446777131 |
| Ltbp1 | -0.446959309 |
| Esf1 | -0.447001229 |
| Ppcdc | -0.447301457 |
| Ggps1 | -0.447809921 |
| Zkscan2 | -0.449706482 |
| Rif1 | -0.452319307 |
| Pcca | -0.452919305 |
| Cdk4 | -0.453510032 |
| Fam188b | -0.454791946 |
| Chp1 | -0.455410511 |
| Gins4 | -0.455552756 |
| Txndc9 | -0.455771514 |
| Hrh1 | -0.456118599 |
| Ltc4s | -0.456190364 |
| Prkar2b | -0.456554287 |
| Stk3 | -0.456563809 |
| Fgfr1op2 | -0.456844734 |
| Tcf15 | -0.457675812 |
| Ackr4 | -0.458290005 |
| Anapc2 | -0.458717605 |
| Lrrc1 | -0.459448147 |
| Bap1 | -0.4597839 |
| Coa4 | -0.459957803 |
| Polk | -0.460006795 |
| Cox11 | -0.460141784 |
| Pdp2 | -0.462456427 |
| Pcsk7 | -0.462659952 |
| Denr | -0.463526057 |
| Mcm7 | -0.463936288 |
| Tanc1 | -0.464742737 |
| Mmrn2 | -0.465217784 |
| Zxdb | -0.465570131 |
| Ube2ql1 | -0.46645637 |
| Ppp3r1 | -0.467286493 |
| Siva1 | -0.467458219 |
| Abca6 | -0.468509209 |
| Cdh13 | -0.468995025 |
| Mllt1 | -0.469895731 |
| Zc3h11a | -0.47034858 |
| Rab5b | -0.470362588 |
| Cdt1 | -0.470646072 |

| | |
|---|---|
| Ftsj1 | -0.47065201 |
| Vasp | -0.47107481 |
| Use1 | -0.471309838 |
| Gimap4 | -0.471498502 |
| Prpf8 | -0.471696858 |
| Exosc1 | -0.471727598 |
| Dph2 | -0.471917345 |
| Reps1 | -0.472078516 |
| Sephs2 | -0.472098352 |
| Gspt2 | -0.472550845 |
| Ywhah | -0.472845443 |
| Fam160b1 | -0.47323091 |
| Ubxn4 | -0.474011378 |
| Gnl3 | -0.474318888 |
| Rpl22l1 | -0.474353367 |
| Eda | -0.474533915 |
| Frmd6 | -0.474885815 |
| Dpysl3 | -0.477274871 |
| Kdm2a | -0.478752048 |
| Stat2 | -0.479248366 |
| Kazn | -0.480291155 |
| Echdc2 | -0.480404716 |
| Atf2 | -0.481328412 |
| Terf2ip | -0.481417077 |
| Cdipt | -0.481644293 |
| Csf1 | -0.482365306 |
| Slc30a6 | -0.482461509 |
| Rbm38 | -0.484831231 |
| Wfs1 | -0.48613823 |
| Csnk1d | -0.487773775 |
| Hook3 | -0.488405316 |
| Bcr | -0.488629296 |
| Herc1 | -0.488677153 |
| Fto | -0.489013058 |
| Scmh1 | -0.489417143 |
| Gpn2 | -0.490564642 |
| Ndufaf5 | -0.490817708 |
| Hmga1 | -0.491438912 |
| Swi5 | -0.492039331 |
| Sorbs1 | -0.493907715 |
| Scml2 | -0.494019249 |
| Fam69a | -0.494131275 |
| Ubald1 | -0.494151444 |
| Rap1gap2 | -0.49510361 |
| 42621 | -0.496072474 |
| Pxdc1 | -0.496120593 |
| Soat1 | -0.49652714 |

| | |
|---|---|
| Cct8 | -0.497355222 |
| Lix1l | -0.497829735 |
| Mrps11 | -0.498127622 |
| Sncaip | -0.498881224 |
| Cib1 | -0.500430762 |
| Rps6ka5 | -0.501631745 |
| Chaf1b | -0.502802017 |
| Bub1b | -0.503682282 |
| Lrrc56 | -0.50459396 |
| Xpo7 | -0.504681076 |
| Thap2 | -0.50652559 |
| Atf3 | -0.508407458 |
| Tet1 | -0.508625582 |
| Bche | -0.508729719 |
| Rpl41 | -0.509107954 |
| Cryab | -0.510121281 |
| Creb3 | -0.511111114 |
| Usp53 | -0.511309484 |
| Bad | -0.511669149 |
| Mospd1 | -0.512708479 |
| Fam227b | -0.514791441 |
| Fam149a | -0.516921727 |
| Pam | -0.516926421 |
| Sec61a1 | -0.51957133 |
| Fez1 | -0.519604882 |
| Api5 | -0.521502424 |
| Fam161a | -0.522279651 |
| Xylt1 | -0.52261054 |
| Cdc25a | -0.523439736 |
| Arhgef26 | -0.524698002 |
| Acp5 | -0.524976581 |
| Wrn | -0.525264607 |
| Acacb | -0.526237317 |
| Rps6kl1 | -0.528191322 |
| Zkscan3 | -0.529139555 |
| Rmdn2 | -0.529900071 |
| Raver2 | -0.530082189 |
| Arid4b | -0.530546762 |
| Xrcc3 | -0.531617975 |
| Csde1 | -0.531629518 |
| Actr2 | -0.532373126 |
| Ebf3 | -0.532811802 |
| Ring1 | -0.53298519 |
| Usp49 | -0.5340124 |
| Tfam | -0.534061656 |
| Rgs17 | -0.534213021 |
| Ufl1 | -0.534490886 |

| | |
|---|---|
| Pcdh1 | -0.534795153 |
| Cct3 | -0.535199105 |
| Rab21 | -0.535941577 |
| Zmynd11 | -0.538104073 |
| Galns | -0.538203724 |
| Hint3 | -0.538483028 |
| Rabl6 | -0.540160319 |
| Bmp4 | -0.54055703 |
| Fam98c | -0.541286727 |
| Wnt3 | -0.541510234 |
| Cox7a2 | -0.541686063 |
| Rmnd5a | -0.542158521 |
| Clec16a | -0.543043947 |
| Fam219a | -0.543376737 |
| Kdm6a | -0.543899523 |
| Btbd9 | -0.543951865 |
| Rgs11 | -0.544317184 |
| Vat1 | -0.544664443 |
| Azin1 | -0.544858535 |
| Vcpkmt | -0.545558783 |
| Tmpo | -0.54580931 |
| Kcnk13 | -0.546690777 |
| Prps1 | -0.547266365 |
| Stim2 | -0.547415092 |
| Rmnd1 | -0.547456122 |
| Prdm15 | -0.54842467 |
| Wdr19 | -0.550172142 |
| Lamp3 | -0.552322908 |
| Ermard | -0.552659932 |
| Blzf1 | -0.552828825 |
| Fnbp4 | -0.553207623 |
| Setdb2 | -0.555080478 |
| Lat | -0.555565322 |
| Golga2 | -0.556416831 |
| Fam198b | -0.557349671 |
| Atp5j2 | -0.557602085 |
| Ttll3 | -0.558773957 |
| Arvcf | -0.559303291 |
| Pin4 | -0.559618547 |
| Rnf7 | -0.559716745 |
| Heatr3 | -0.560591815 |
| Lsm3 | -0.561821096 |
| Thbd | -0.561884408 |
| Etv2 | -0.562400051 |
| Znfx1 | -0.562593096 |
| Tbcc | -0.563269225 |
| Hsd17b4 | -0.563502594 |

| | |
|---|---|
| Efnb3 | -0.563641761 |
| Cebpg | -0.563667328 |
| Homer2 | -0.563993343 |
| Stoml1 | -0.565598255 |
| Rep15 | -0.566483514 |
| Rhob | -0.566851743 |
| Edem1 | -0.566884159 |
| Mfge8 | -0.56750859 |
| Phip | -0.568394164 |
| Tbck | -0.568407659 |
| Mum1 | -0.568972811 |
| Dffa | -0.570203619 |
| Akap6 | -0.570978999 |
| C2cd4c | -0.571995629 |
| Snapin | -0.57225185 |
| Donson | -0.572390351 |
| Lrfn5 | -0.575226975 |
| Adtrp | -0.576360594 |
| Sppl3 | -0.576448737 |
| Il17rb | -0.576760133 |
| Chka | -0.577454739 |
| Melk | -0.578558636 |
| Fam96b | -0.580535323 |
| Pou2f2 | -0.581974703 |
| Pbx2 | -0.582180859 |
| Taf7l | -0.584766583 |
| Hn1l | -0.584873953 |
| Parp1 | -0.586137674 |
| Poldip3 | -0.586262773 |
| Nt5c2 | -0.586384832 |
| Serpinh1 | -0.586438206 |
| Sucla2 | -0.587492072 |
| Ptch1 | -0.588164836 |
| Rrm2b | -0.588376888 |
| Thy1 | -0.588422979 |
| Rnf208 | -0.589107574 |
| Ddx19b | -0.589864506 |
| Fndc3a | -0.590178738 |
| Luzp1 | -0.590675703 |
| Tmem56 | -0.591001236 |
| Mrpl20 | -0.591289227 |
| Herpud2 | -0.591296189 |
| Prim1 | -0.591394486 |
| Cdc25b | -0.591754804 |
| Sympk | -0.591809897 |
| Tex22 | -0.592579934 |
| Asap2 | -0.592663916 |

| | |
|---|---|
| Ndc80 | -0.593693221 |
| Thumpd1 | -0.593948181 |
| Pip4k2c | -0.593954021 |
| Spidr | -0.595145465 |
| Cdh12 | -0.596028031 |
| Ssc5d | -0.596510278 |
| Cep44 | -0.59719574 |
| Ift27 | -0.597500206 |
| Hdac6 | -0.59799064 |
| Thtpa | -0.600281604 |
| S1pr4 | -0.60125356 |
| Trpm7 | -0.60196049 |
| Ltn1 | -0.605974032 |
| Hspbap1 | -0.606338922 |
| Apol6 | -0.608018532 |
| Ppm1d | -0.60876844 |
| Bloc1s2 | -0.609089563 |
| Ncapd3 | -0.60920401 |
| Ckb | -0.609465389 |
| Il1rapl1 | -0.609528524 |
| Snrpe | -0.610181428 |
| Zfp37 | -0.610779155 |
| Josd2 | -0.611820181 |
| Pfn2 | -0.611898294 |
| Dyrk1a | -0.612570179 |
| Pdcl3 | -0.612591375 |
| Hdac9 | -0.614744993 |
| Tspyl5 | -0.614946632 |
| Krr1 | -0.615059447 |
| Ifnar1 | -0.616741118 |
| Patz1 | -0.617015317 |
| Pik3r1 | -0.617308782 |
| Rp9 | -0.617629276 |
| Rasef | -0.617690933 |
| Zscan20 | -0.617927543 |
| Fancm | -0.618073553 |
| Grpel2 | -0.619997814 |
| Cyb561 | -0.620544639 |
| Zbtb26 | -0.621066445 |
| Snapc5 | -0.621440522 |
| Zdhhc8 | -0.621923149 |
| Mctp1 | -0.623106725 |
| Ppp1r3b | -0.62406723 |
| Epg5 | -0.624180327 |
| Lpin3 | -0.624578531 |
| Hoxa2 | -0.624658646 |
| Atp7b | -0.625591542 |

| | |
|---|---|
| Slc39a9 | -0.626207788 |
| Rbpms2 | -0.627391778 |
| Atp6v1a | -0.628160966 |
| Hes7 | -0.629740802 |
| Hoxa7 | -0.629741639 |
| Cyth1 | -0.631074781 |
| Ddx43 | -0.631509122 |
| Cd40 | -0.631845492 |
| Cldn12 | -0.632280252 |
| Alyref | -0.632773831 |
| Prr14 | -0.632926993 |
| Slc22a4 | -0.633428242 |
| Bicd1 | -0.634851952 |
| Vcp | -0.635241984 |
| Lipg | -0.635878025 |
| Tm9sf4 | -0.636091209 |
| Mospd2 | -0.636444136 |
| Rps6 | -0.636547573 |
| Ctnnal1 | -0.636924035 |
| Prss3 | -0.638956078 |
| Aadat | -0.639457397 |
| Ldlrad3 | -0.641061779 |
| Syt16 | -0.642311845 |
| Tsen34 | -0.642615089 |
| Ndufs4 | -0.643471954 |
| Rab32 | -0.643971144 |
| Macf1 | -0.644905816 |
| Ube2k | -0.645871389 |
| Nt5dc3 | -0.646478218 |
| L3hypdh | -0.64708687 |
| Ppp1r10 | -0.648209397 |
| Gtf3c3 | -0.650014833 |
| Ubxn2b | -0.650383083 |
| Nup205 | -0.650429535 |
| Mkrn1 | -0.65088457 |
| Mrps36 | -0.65200406 |
| Nudt3 | -0.652178703 |
| Wdr82 | -0.652497513 |
| Cox7a1 | -0.654442307 |
| Sgsm2 | -0.655268054 |
| Ube2l3 | -0.655733636 |
| Cdc37 | -0.656109369 |
| Khsrp | -0.656315838 |
| Pelo | -0.656384949 |
| Kdm4c | -0.657924966 |
| Zmym3 | -0.658870231 |
| Klhl9 | -0.65941189 |

| | |
|---|---|
| Zfand2a | -0.659880565 |
| Ppfibp2 | -0.660327737 |
| Vps53 | -0.660360315 |
| Hint1 | -0.66038053 |
| Pigo | -0.661487177 |
| Ucp2 | -0.661749661 |
| Rbms1 | -0.663501987 |
| Stx11 | -0.664398923 |
| Cpped1 | -0.664920843 |
| Tfb1m | -0.665504215 |
| Hibch | -0.666076848 |
| Ccdc125 | -0.666183381 |
| Aldh9a1 | -0.667542543 |
| Stard4 | -0.668102355 |
| Tbc1d22a | -0.66826234 |
| Trnt1 | -0.668741751 |
| Nefh | -0.669670034 |
| Dach1 | -0.669786235 |
| Tspyl4 | -0.670611891 |
| Phactr2 | -0.671152207 |
| Rora | -0.671447548 |
| Ube2t | -0.672870693 |
| Pear1 | -0.673344981 |
| Pole4 | -0.674965997 |
| Ccdc107 | -0.675145968 |
| Ada | -0.67524348 |
| Hmha1 | -0.675739585 |
| Dcaf12l1 | -0.675894828 |
| Prkca | -0.675950775 |
| Prdx1 | -0.676063271 |
| Tdrkh | -0.6762854 |
| Shb | -0.678696862 |
| Bloc1s3 | -0.679040754 |
| Igbp1 | -0.679188111 |
| Celsr3 | -0.680070388 |
| Nutf2 | -0.683671485 |
| Chid1 | -0.683971852 |
| Pkm | -0.684581407 |
| Ctnna1 | -0.684785854 |
| Cbx1 | -0.684815436 |
| Rpl12 | -0.68603873 |
| Imp3 | -0.686359724 |
| Fkbp3 | -0.686768399 |
| Lif | -0.687193606 |
| Luc7l | -0.687653669 |
| Kcnk2 | -0.688102123 |
| Capn10 | -0.688220003 |

| | |
|---|---|
| Ostc | -0.689289828 |
| Ticrr | -0.68960024 |
| Ndufv3 | -0.690205351 |
| Vav2 | -0.691371267 |
| Zscan29 | -0.69260531 |
| Mapre1 | -0.693058345 |
| Cttnbp2nl | -0.693062666 |
| Gltscr2 | -0.693300786 |
| Entpd4 | -0.69340341 |
| Nifk | -0.695516772 |
| Ppp1r15a | -0.695694496 |
| Dnajb4 | -0.696329604 |
| Josd1 | -0.697029125 |
| Slc39a11 | -0.698541815 |
| Ddit4 | -0.699311829 |
| Katnal2 | -0.699481474 |
| Ripk3 | -0.699507917 |
| Hykk | -0.699735931 |
| Yae1d1 | -0.700680024 |
| Cpne2 | -0.701267137 |
| Chchd4 | -0.701650733 |
| Cetn2 | -0.702087563 |
| Rrn3 | -0.70231204 |
| Timm8b | -0.70299362 |
| Pomk | -0.704107717 |
| Pla2g6 | -0.704375727 |
| Polr2g | -0.704907594 |
| Pard6g | -0.705322145 |
| Lipt2 | -0.70539357 |
| Bag1 | -0.706069648 |
| Med30 | -0.706860802 |
| Eif2b5 | -0.707317728 |
| Mief2 | -0.707750193 |
| Hoxb2 | -0.710601918 |
| Cd63 | -0.711471932 |
| Cpsf4 | -0.711682863 |
| Med8 | -0.711961211 |
| Thbs2 | -0.712233505 |
| Pik3r4 | -0.713646188 |
| Fam110b | -0.713988988 |
| Zmym4 | -0.714513498 |
| Sfmbt2 | -0.71515024 |
| Kif20a | -0.715775211 |
| Fahd1 | -0.717616989 |
| Aco2 | -0.719047133 |
| Eci1 | -0.722450234 |
| Rtfdc1 | -0.722656584 |

| | |
|---|---|
| Fam101a | -0.723622747 |
| Eif5a2 | -0.724288152 |
| Sp4 | -0.724504863 |
| Vars | -0.724660396 |
| Suco | -0.725615075 |
| Arpc1b | -0.728030338 |
| Flywch1 | -0.728481793 |
| Raph1 | -0.728816015 |
| Spsb2 | -0.730867218 |
| Csnk2a1 | -0.731527284 |
| Lsm4 | -0.731593308 |
| Fam45a | -0.731985401 |
| Sema5a | -0.732290876 |
| Cdyl | -0.732711545 |
| Tmem87a | -0.733955727 |
| Fat4 | -0.73502824 |
| Ccrl2 | -0.73506537 |
| Slc37a2 | -0.738601207 |
| Osbp2 | -0.739361254 |
| Samd9l | -0.7401904 |
| Papola | -0.74045251 |
| Tex2 | -0.741508433 |
| Nploc4 | -0.74397864 |
| Nav3 | -0.744354124 |
| Styx | -0.744811214 |
| Cnr1 | -0.745414431 |
| Asb14 | -0.745956409 |
| Inpp5f | -0.746141013 |
| Derl2 | -0.74701792 |
| Asb13 | -0.747055694 |
| Zcrb1 | -0.747459315 |
| Kif22 | -0.748823171 |
| Atg4d | -0.749338986 |
| Cog8 | -0.74955918 |
| Ppcs | -0.749829412 |
| Kdelr3 | -0.750300675 |
| Tgfa | -0.751289304 |
| Gls | -0.754511212 |
| Actr3b | -0.757052943 |
| Zrsr2 | -0.757062245 |
| Setd1b | -0.757637439 |
| Lrch3 | -0.759025092 |
| Rai14 | -0.759760727 |
| Sirt4 | -0.7619969 |
| Xrcc4 | -0.762552527 |
| Sh3glb2 | -0.763815408 |
| Crybb2 | -0.764231346 |

| | |
|---|---|
| Hsdl1 | -0.764669644 |
| Gpnmb | -0.765068748 |
| Tomm70a | -0.7662717 |
| Atxn2 | -0.766780639 |
| Dnmt3b | -0.767755028 |
| Ctdsp1 | -0.768480583 |
| Il1a | -0.769303141 |
| Fap | -0.769520824 |
| Kcnt2 | -0.769843766 |
| Ankle2 | -0.77217203 |
| Htatip2 | -0.77333652 |
| Pole3 | -0.77448915 |
| Galnt4 | -0.775519436 |
| Map3k3 | -0.7755338 |
| Crbn | -0.776171303 |
| Stmn1 | -0.776418767 |
| Papd7 | -0.776934695 |
| Mturn | -0.777191762 |
| Ppid | -0.777200167 |
| Vti1a | -0.777822558 |
| Pter | -0.77845328 |
| Dctn2 | -0.778556817 |
| Taok1 | -0.778854272 |
| Mapk8ip2 | -0.779123694 |
| Oxa1l | -0.779139735 |
| Dnph1 | -0.780118227 |
| Stil | -0.780191472 |
| Supt7l | -0.780544903 |
| Abhd14b | -0.781101316 |
| Rxra | -0.782471911 |
| Il13ra2 | -0.784560505 |
| Cel | -0.784687291 |
| Zcchc10 | -0.785355252 |
| Tlr3 | -0.785582741 |
| Zdhhc21 | -0.785878984 |
| Prpf6 | -0.78603759 |
| Tmem205 | -0.786398072 |
| Pcdh19 | -0.786801937 |
| Serpinb8 | -0.787766441 |
| Akap7 | -0.788787869 |
| Tmem69 | -0.789133806 |
| Wipi2 | -0.789168275 |
| Plekhg5 | -0.789521168 |
| Aldh4a1 | -0.79009684 |
| Dad1 | -0.790404844 |
| Clgn | -0.791339405 |
| Nhsl1 | -0.79180361 |

| | |
|---|---|
| Drg1 | -0.792676172 |
| Pex11g | -0.794252671 |
| Epha1 | -0.794998834 |
| Gde1 | -0.795137706 |
| Eno1 | -0.795397419 |
| Wdr43 | -0.797242033 |
| Kank1 | -0.797254504 |
| Rrp8 | -0.801000355 |
| Srsf1 | -0.802490609 |
| Dek | -0.804302416 |
| Dolk | -0.804486641 |
| Hjurp | -0.805030155 |
| Ahdc1 | -0.805977753 |
| Picalm | -0.806018943 |
| Gtpbp2 | -0.806344475 |
| Hmgcs1 | -0.806728343 |
| Crhbp | -0.80759013 |
| Fam83g | -0.807820551 |
| S1pr2 | -0.808488575 |
| Sema3a | -0.809355414 |
| Obfc1 | -0.809507146 |
| Tbc1d9b | -0.811315407 |
| Dvl2 | -0.812531503 |
| Ak6 | -0.813665416 |
| Pcgf6 | -0.814656495 |
| Traf3ip2 | -0.817732122 |
| Adamts12 | -0.818092532 |
| Cisd1 | -0.818534764 |
| Dennd2c | -0.819777131 |
| Ankrd16 | -0.820532219 |
| Atp2c1 | -0.821197054 |
| Cradd | -0.822964101 |
| Helq | -0.823656933 |
| Odf2l | -0.824052819 |
| Hnrnpl | -0.824382672 |
| Lrrc29 | -0.826126361 |
| Socs3 | -0.828203492 |
| Cacng7 | -0.828628488 |
| Kcnrg | -0.829795241 |
| Ccdc174 | -0.830263015 |
| Alcam | -0.830535979 |
| Tuft1 | -0.830857975 |
| Clip4 | -0.830916906 |
| Zbtb43 | -0.833550531 |
| Pank2 | -0.833615429 |
| Thap4 | -0.833890206 |
| Smarcd2 | -0.834039657 |

| | |
|---|---|
| Fam186b | -0.835260319 |
| Nucb1 | -0.835332719 |
| Glcci1 | -0.835947189 |
| Tmem261 | -0.836004266 |
| Agpat4 | -0.836037963 |
| Akap3 | -0.836452172 |
| Serf2 | -0.837950176 |
| Emilin2 | -0.838318757 |
| Syde1 | -0.838532276 |
| Mtcp1 | -0.841653916 |
| Trip6 | -0.841916632 |
| Rassf9 | -0.845010665 |
| Ccdc58 | -0.846589536 |
| Phkb | -0.847730343 |
| Nom1 | -0.849196379 |
| Dok5 | -0.85128376 |
| Plk3 | -0.852493119 |
| Sltm | -0.853786144 |
| Pard3b | -0.853975105 |
| Inpp5b | -0.854319486 |
| Ddx21 | -0.855126998 |
| Dhx8 | -0.856565194 |
| Usp16 | -0.857825548 |
| Aqr | -0.859807216 |
| Kcng1 | -0.862061018 |
| Eri1 | -0.862981003 |
| Slc37a3 | -0.864235668 |
| Asl | -0.867354202 |
| Gab3 | -0.867563578 |
| Ptprk | -0.868071585 |
| Pdcd4 | -0.868380907 |
| Zzz3 | -0.868812355 |
| Slc1a4 | -0.869321714 |
| Lrp2bp | -0.869326618 |
| Actl10 | -0.869908568 |
| Gorasp2 | -0.87256256 |
| Ly6k | -0.872631727 |
| Rab8a | -0.872818326 |
| Ica1l | -0.873295336 |
| Tmem126b | -0.875293655 |
| Trappc3 | -0.875590792 |
| Ctps2 | -0.8771004 |
| Prss12 | -0.878322428 |
| Arntl2 | -0.878724878 |
| Lmnb1 | -0.878856361 |
| Ldlrad4 | -0.880154422 |
| Wdr76 | -0.880248682 |

| | |
|---|---|
| Eef1d | -0.881362909 |
| Ltb4r2 | -0.88165967 |
| Dhfr | -0.885534875 |
| Ttc5 | -0.886274352 |
| Herc3 | -0.886635146 |
| Dctn4 | -0.889725722 |
| Tmem64 | -0.890624783 |
| Smg1 | -0.891337428 |
| Vps26a | -0.891672677 |
| Bri3 | -0.893057606 |
| Mecp2 | -0.893157813 |
| Slc35f6 | -0.894813248 |
| Ctcf | -0.897934872 |
| Kcnc3 | -0.899530389 |
| Osbpl8 | -0.899844186 |
| Fam122c | -0.900050515 |
| Pacsin3 | -0.900698249 |
| Sar1b | -0.900826719 |
| Slc12a6 | -0.901631494 |
| Rfk | -0.905207139 |
| Ppm1g | -0.905535312 |
| Lpcat2 | -0.908656465 |
| Pef1 | -0.910906224 |
| Cmpk1 | -0.911315205 |
| Mtx2 | -0.911351854 |
| Edem3 | -0.911743507 |
| Gart | -0.911958845 |
| Chchd1 | -0.913374683 |
| Ing3 | -0.913546859 |
| Pmel | -0.91471704 |
| Zc3h12c | -0.915329413 |
| Fem1c | -0.917234908 |
| Hpse | -0.917383696 |
| Brf1 | -0.917871695 |
| Ddx46 | -0.919935772 |
| Actb | -0.920016063 |
| Bud31 | -0.921421645 |
| Ctu2 | -0.922015553 |
| Prkci | -0.922232735 |
| Krt81 | -0.923199918 |
| Mark4 | -0.924851136 |
| Hsd17b14 | -0.926379138 |
| Phf7 | -0.928065468 |
| Eif4g2 | -0.929171279 |
| Stxbp4 | -0.929338274 |
| Cd14 | -0.932002608 |
| Trappc3l | -0.934937336 |

| | |
|---|---|
| Rras2 | -0.935048999 |
| Usp5 | -0.93612662 |
| Sgk3 | -0.936965135 |
| Ube2j1 | -0.937832747 |
| Rps6ka2 | -0.940666472 |
| Ska2 | -0.941010699 |
| Nampt | -0.943203285 |
| Cog5 | -0.945227915 |
| Dhrsx | -0.946425383 |
| Acbd3 | -0.947744024 |
| Nvl | -0.947993755 |
| Calm1 | -0.948360327 |
| Edil3 | -0.949609242 |
| Tdp1 | -0.950888923 |
| Mdn1 | -0.951187542 |
| Tmem115 | -0.95236752 |
| Fam49a | -0.952646759 |
| Rnf31 | -0.952735148 |
| Ckap5 | -0.953941786 |
| Nkiras1 | -0.954313836 |
| Lrrc40 | -0.955451456 |
| Mtfp1 | -0.956167183 |
| Ankrd52 | -0.956932013 |
| Camsap2 | -0.957382019 |
| Tor4a | -0.958880817 |
| Tex261 | -0.960032537 |
| Eif4a3 | -0.960323583 |
| Tead2 | -0.960442118 |
| Trpm4 | -0.96380485 |
| Tsnaxip1 | -0.964123717 |
| Accs | -0.965143302 |
| Ccdc171 | -0.967194542 |
| Sema4f | -0.97025955 |
| Tomm5 | -0.970985749 |
| Micalcl | -0.971196475 |
| Nrde2 | -0.9731797 |
| Lrrc66 | -0.973183754 |
| Ccnd3 | -0.973663681 |
| Dnajc18 | -0.974472652 |
| Bccip | -0.974940145 |
| Ccr10 | -0.975915066 |
| F2rl2 | -0.97704893 |
| Ficd | -0.977061199 |
| Rpgrip1l | -0.977124423 |
| Nptxr | -0.977866199 |
| Srfbp1 | -0.977885171 |
| Acads | -0.978710066 |

| | |
|---|---|
| Tmeff2 | -0.978951306 |
| Gripap1 | -0.981380787 |
| Stard13 | -0.984424224 |
| Zfc3h1 | -0.986456888 |
| Taf6l | -0.987536851 |
| Hspa4l | -0.991474304 |
| Slc19a1 | -0.995365706 |
| Sprn | -0.996162208 |
| Nsun2 | -0.996385989 |
| Arhgef10l | -0.999075805 |
| Dkc1 | -0.99945481 |
| Trim33 | -1.00018362 |
| Kcnmb4 | -1.000375754 |
| Fam102b | -1.000744081 |
| Mrps12 | -1.001249189 |
| Fign | -1.003033174 |
| Snx3 | -1.003438086 |
| Cox8a | -1.003630523 |
| Rad9a | -1.003860994 |
| Xab2 | -1.005839445 |
| Cript | -1.014028862 |
| Tnfsf15 | -1.015154385 |
| Stc1 | -1.016034442 |
| Larp1b | -1.016090624 |
| Uqcrh | -1.019631948 |
| Gpr146 | -1.021522094 |
| Zdhhc22 | -1.022560764 |
| Ddx6 | -1.023579254 |
| Utp6 | -1.024792216 |
| Sec22a | -1.025217943 |
| Dbnl | -1.025563659 |
| Ipo9 | -1.027964427 |
| Tfb2m | -1.028301575 |
| Tiprl | -1.032128025 |
| Etnk1 | -1.03320992 |
| Bin1 | -1.033721696 |
| Csnk1g1 | -1.034431837 |
| Slc4a8 | -1.035514395 |
| Cry1 | -1.036952901 |
| Eif2s2 | -1.037813412 |
| Tmem126a | -1.038766325 |
| Tmem71 | -1.038792932 |
| Hnrnph3 | -1.039391563 |
| Dopey2 | -1.042728829 |
| Cdca5 | -1.044106933 |
| Mark2 | -1.04415579 |
| Rbm14 | -1.044625281 |

| | |
|---|---|
| Nrp2 | -1.044720994 |
| Usp24 | -1.045217595 |
| Mapt | -1.046862407 |
| Agtpbp1 | -1.05056791 |
| Ccdc22 | -1.050579362 |
| Tmem45a | -1.052922742 |
| Tradd | -1.052946323 |
| Dpp4 | -1.054670292 |
| Mb21d1 | -1.056533224 |
| S100a13 | -1.058438998 |
| Eif3e | -1.060514304 |
| Exoc4 | -1.061350872 |
| Fhl1 | -1.063403416 |
| Tmem208 | -1.065040334 |
| Fst | -1.065064849 |
| Map4k5 | -1.065791663 |
| Strbp | -1.071405656 |
| Yif1a | -1.071484045 |
| Nat10 | -1.071679702 |
| Cdc20 | -1.072259461 |
| U2af2 | -1.074125327 |
| Txnip | -1.074312913 |
| Med31 | -1.07686383 |
| Wdr54 | -1.077081602 |
| Prr22 | -1.078083072 |
| Usp14 | -1.07864079 |
| Tmem65 | -1.079529976 |
| Wwc1 | -1.080561008 |
| Tacc3 | -1.080679134 |
| Tctn3 | -1.083243998 |
| Nudt17 | -1.084013336 |
| Ptgis | -1.087203439 |
| Diablo | -1.088265382 |
| Fryl | -1.08936844 |
| Hoxd1 | -1.089788564 |
| Cadps2 | -1.090399947 |
| Slc25a42 | -1.091957645 |
| Fam26e | -1.092828475 |
| Trip10 | -1.093316829 |
| Srp72 | -1.093547492 |
| G0s2 | -1.097489783 |
| Cplx1 | -1.098901866 |
| Skap2 | -1.099622191 |
| Uso1 | -1.100268134 |
| Sp100 | -1.100663707 |
| Rexo2 | -1.100726752 |
| Adam9 | -1.101628039 |

| | |
|---|---|
| Jag2 | -1.104035559 |
| Hsf2bp | -1.105783271 |
| Spag4 | -1.106974885 |
| Exoc2 | -1.10700636 |
| Dgke | -1.108157772 |
| Sec11c | -1.110398472 |
| Prkrip1 | -1.110510076 |
| Lrig1 | -1.110657876 |
| Cttnbp2 | -1.112631559 |
| Cdnf | -1.112735217 |
| Sfpq | -1.113960308 |
| Arrdc3 | -1.116236236 |
| Fer1l4 | -1.117382748 |
| Snrpn | -1.119430887 |
| Brca2 | -1.121934833 |
| Nabp1 | -1.122869284 |
| Otud1 | -1.122970776 |
| Nudt21 | -1.127640769 |
| Nsun3 | -1.128654767 |
| Rara | -1.12918933 |
| St3gal3 | -1.132435918 |
| Eif3h | -1.13439738 |
| Vps25 | -1.13475661 |
| Vps13a | -1.135261295 |
| Ccnc | -1.136030544 |
| Zdhhc2 | -1.137397777 |
| Casp8 | -1.137743789 |
| Asph | -1.139530686 |
| Edf1 | -1.140307017 |
| Mme | -1.142199703 |
| Rc3h2 | -1.142204056 |
| Slc35a4 | -1.144793946 |
| Trim27 | -1.145279043 |
| Arhgap18 | -1.146283189 |
| Vti1b | -1.146853045 |
| Fbxl6 | -1.146860705 |
| Pik3cb | -1.148850408 |
| Cep192 | -1.149998911 |
| Slc39a13 | -1.151257285 |
| Capn15 | -1.151621802 |
| Spatc1l | -1.152090264 |
| Lpxn | -1.152529315 |
| Prkaa2 | -1.15516924 |
| Capn7 | -1.161664405 |
| Bex1 | -1.163196235 |
| Orc4 | -1.164687384 |
| Mef2d | -1.164735016 |

| | |
|---|---|
| Sapcd2 | -1.166443968 |
| Ap5m1 | -1.168890675 |
| Zfp36l2 | -1.169099371 |
| Akt1s1 | -1.169217258 |
| Fmod | -1.169754001 |
| Leng1 | -1.169823128 |
| Abcf1 | -1.170058414 |
| Adat2 | -1.170641298 |
| Hmbs | -1.173319226 |
| Zbtb8a | -1.173403144 |
| Rpl32 | -1.174527121 |
| Pygo1 | -1.175653759 |
| Rnf114 | -1.17823075 |
| Gtf2i | -1.180281479 |
| Pus7l | -1.180866158 |
| Etv6 | -1.18185936 |
| Hs1bp3 | -1.181917792 |
| Ythdc2 | -1.182793489 |
| Bloc1s4 | -1.184849193 |
| Plcd4 | -1.185035215 |
| Rassf5 | -1.185270113 |
| Slx4 | -1.18685683 |
| Trub1 | -1.186954052 |
| Psma2 | -1.188152466 |
| Kcnk3 | -1.198044951 |
| Dhrs1 | -1.199475183 |
| Sox5 | -1.199972435 |
| Itpkb | -1.200849464 |
| Aldh3a1 | -1.201651346 |
| Tpst1 | -1.201763465 |
| Aifm2 | -1.201897711 |
| Rnf215 | -1.206182448 |
| Ccdc102a | -1.206913688 |
| Tnfrsf1b | -1.20886292 |
| Wars | -1.210250691 |
| Foxk2 | -1.210613417 |
| C7 | -1.211203637 |
| Fyb | -1.211994715 |
| Evl | -1.216290411 |
| Gpr3 | -1.218286241 |
| Ppp1r35 | -1.221534819 |
| Hesx1 | -1.222967097 |
| Hdac1 | -1.22536241 |
| Syne3 | -1.227110386 |
| Rab3il1 | -1.227110992 |
| Nup43 | -1.229812891 |
| Fbxw7 | -1.230288911 |

| | |
|---|---|
| Prkar2a | -1.230389748 |
| Sgcb | -1.23087416 |
| Rabl3 | -1.231135899 |
| Dpy19l1 | -1.232232195 |
| Pogz | -1.23603116 |
| Prdx2 | -1.244574915 |
| Sae1 | -1.245218367 |
| Rsrc1 | -1.245898059 |
| Shq1 | -1.24695879 |
| Plekha3 | -1.249559506 |
| Ubl4a | -1.25058147 |
| Flcn | -1.251352645 |
| Sele | -1.253833606 |
| Ccdc170 | -1.25408943 |
| Gjc1 | -1.255666203 |
| Rplp1 | -1.256071238 |
| Top2b | -1.25625963 |
| Tmem131 | -1.256522708 |
| Fbxo11 | -1.257049635 |
| Tgfbi | -1.258924663 |
| Mpp7 | -1.260119933 |
| C1galt1 | -1.262902284 |
| Zcchc17 | -1.262911733 |
| Son | -1.263950746 |
| Pms1 | -1.264000359 |
| Swt1 | -1.265937849 |
| Tsen54 | -1.266359279 |
| Kif15 | -1.269341997 |
| Paip2b | -1.26997933 |
| Snap47 | -1.273271296 |
| Rab8b | -1.277053092 |
| Terf2 | -1.282450666 |
| Slc4a11 | -1.283932673 |
| Minpp1 | -1.285021116 |
| Lynx1 | -1.285312242 |
| Rassf1 | -1.28920348 |
| Gkap1 | -1.290750555 |
| Cep350 | -1.292364553 |
| Copg2 | -1.29252372 |
| Sap130 | -1.29276002 |
| Bckdhb | -1.294567407 |
| Tbc1d5 | -1.299869917 |
| Ubap2l | -1.30212018 |
| Fam217b | -1.302935178 |
| Taf8 | -1.304789446 |
| Stag2 | -1.305194245 |
| Tmem256 | -1.305498534 |

| | |
|---|---|
| Med13 | -1.306276708 |
| Snrpd1 | -1.307304678 |
| Hspa4 | -1.308782921 |
| Prkcdbp | -1.309556585 |
| Tmem179b | -1.311060205 |
| Nat9 | -1.311153046 |
| Pim3 | -1.31292049 |
| Ddr1 | -1.313530242 |
| Ilkap | -1.313668215 |
| Prelid1 | -1.31390129 |
| Hspb8 | -1.317007694 |
| Lipa | -1.319512181 |
| Rwdd2a | -1.321058042 |
| Tusc2 | -1.321407452 |
| Arl5b | -1.322810733 |
| Ap4e1 | -1.324072189 |
| Relt | -1.324357571 |
| Dennd2a | -1.3260201 |
| Ccl2 | -1.327693762 |
| Atad2 | -1.328854661 |
| Snx2 | -1.329988613 |
| 42624 | -1.332082281 |
| Uqcrc2 | -1.333642239 |
| Mapk7 | -1.339167024 |
| Kin | -1.339403092 |
| Ylpm1 | -1.339900126 |
| Cnot11 | -1.340109888 |
| Brcc3 | -1.342915533 |
| Iws1 | -1.34597127 |
| Whsc1 | -1.348135746 |
| Atp6v0c | -1.349889009 |
| Idh1 | -1.351131367 |
| Dnajc24 | -1.352017944 |
| Akirin1 | -1.354836027 |
| Maf | -1.35497078 |
| Ppp2r4 | -1.356835345 |
| Cybrd1 | -1.360631673 |
| Mios | -1.361154339 |
| Dffb | -1.361929614 |
| Elovl1 | -1.363796905 |
| Lrrc57 | -1.365335291 |
| Mpv17l | -1.366616011 |
| Ccser2 | -1.367076741 |
| Rrp36 | -1.369926431 |
| Amotl2 | -1.370647434 |
| Gpbp1l1 | -1.37124289 |
| Uck1 | -1.371597651 |

| | |
|---|---|
| Pax9 | -1.374486021 |
| Pign | -1.375857581 |
| Tprkb | -1.376948168 |
| Fam114a1 | -1.377042813 |
| Nxt1 | -1.377902166 |
| Col25a1 | -1.38140132 |
| Mfsd8 | -1.382598451 |
| Nostrin | -1.383581236 |
| Napb | -1.384869248 |
| Gabrb3 | -1.386860958 |
| Hsp90ab1 | -1.387559694 |
| Rad51 | -1.388748969 |
| Cwf19l2 | -1.388857046 |
| Metrnl | -1.389225198 |
| Ptgr1 | -1.390200834 |
| Nalcn | -1.390203193 |
| Rab4b | -1.391740657 |
| R3hdm2 | -1.391993359 |
| Focad | -1.392606837 |
| Plod3 | -1.398716457 |
| Rab12 | -1.398904611 |
| Ndst3 | -1.401403437 |
| Hnrnpu | -1.401424068 |
| G3bp2 | -1.402174488 |
| Rnase4 | -1.403889643 |
| Exoc7 | -1.40573528 |
| Sh3d21 | -1.408037807 |
| Pcdhga1 | -1.409157367 |
| Shcbp1 | -1.409931738 |
| Hmgxb3 | -1.41362749 |
| Gorab | -1.413963848 |
| Agps | -1.415435134 |
| Rab14 | -1.416438903 |
| Pigz | -1.417822639 |
| Wdr89 | -1.424286407 |
| Rnf20 | -1.428008887 |
| Efcab11 | -1.432335892 |
| Ssfa2 | -1.433847439 |
| Crebbp | -1.436299578 |
| Kif27 | -1.436451646 |
| Map3k1 | -1.443794706 |
| Adamts9 | -1.446646908 |
| Nr1d2 | -1.448402349 |
| Lsm6 | -1.448717113 |
| Pdzd2 | -1.453148448 |
| Sspo | -1.45967217 |
| Nmral1 | -1.465333002 |

| | |
|---|---|
| Plk4 | -1.470111986 |
| Pds5b | -1.474805821 |
| Ddx42 | -1.478528496 |
| Rfc3 | -1.478647462 |
| Aars2 | -1.478885784 |
| Lrrc58 | -1.480046111 |
| Rps6ka6 | -1.482441303 |
| Cubn | -1.482525043 |
| Ppil3 | -1.483510828 |
| Dcun1d3 | -1.483720417 |
| Plcb1 | -1.484288308 |
| Tmem25 | -1.486919653 |
| Gpr75 | -1.493987306 |
| Sf3b1 | -1.496302162 |
| Cela3b | -1.498837311 |
| F8 | -1.500851086 |
| Tfec | -1.502666552 |
| Smg7 | -1.502848876 |
| Ndufa9 | -1.504148139 |
| Ccdc138 | -1.505974375 |
| Fam133b | -1.50756708 |
| Tet2 | -1.509157814 |
| Ttc25 | -1.514222577 |
| Cacna2d4 | -1.514562948 |
| Commd5 | -1.517002488 |
| Prrg4 | -1.521835413 |
| Tsc22d1 | -1.522940018 |
| Bptf | -1.524841004 |
| Ebp | -1.526337983 |
| Cpxm1 | -1.529056163 |
| Mbd1 | -1.529909928 |
| Pde11a | -1.533121701 |
| Sgta | -1.538779015 |
| Gnaz | -1.541529513 |
| Chmp6 | -1.546637908 |
| Map3k10 | -1.551139623 |
| Mn1 | -1.553686076 |
| Ccnk | -1.556854171 |
| Gnai1 | -1.562133478 |
| Haus4 | -1.562234412 |
| Ubxn11 | -1.56316139 |
| F2r | -1.563568041 |
| Cnih1 | -1.565268311 |
| Fyco1 | -1.567144743 |
| Tyw3 | -1.567325348 |
| Bbip1 | -1.567449327 |
| H1f0 | -1.567833982 |

| | |
|---|---|
| Erc2 | -1.569597939 |
| Tmed8 | -1.570070092 |
| Rps4x | -1.570244029 |
| Ripk2 | -1.570782508 |
| 42615 | -1.57228795 |
| Bcl2l12 | -1.575171212 |
| Slc9b1 | -1.575839427 |
| Hspb1 | -1.576289126 |
| Dnajc15 | -1.578680108 |
| Trim26 | -1.579418779 |
| Pgam5 | -1.585896566 |
| Rbx1 | -1.589265317 |
| Tnfaip8 | -1.589840648 |
| Fam122a | -1.590042005 |
| Nudt22 | -1.591434052 |
| Ppp1r3e | -1.592220234 |
| Shf | -1.597312393 |
| Slc35b3 | -1.602961458 |
| Parvb | -1.606544484 |
| Fgd6 | -1.608111872 |
| Lsamp | -1.611016293 |
| Ndufs7 | -1.611400403 |
| Ssbp3 | -1.611621695 |
| Phka1 | -1.613323201 |
| Rabgap1 | -1.615188976 |
| Klhl23 | -1.618393435 |
| Vps52 | -1.623526731 |
| Cdc73 | -1.624050878 |
| Rapgefl1 | -1.629905014 |
| Egr3 | -1.632138775 |
| Jmjd7 | -1.638823992 |
| Tbx3 | -1.643121237 |
| Thrap3 | -1.643176975 |
| Htra1 | -1.643268542 |
| Kif23 | -1.643741904 |
| Proca1 | -1.646909742 |
| Map2k5 | -1.648854605 |
| 42617 | -1.649189477 |
| Tmem106b | -1.650222316 |
| Nmnat1 | -1.654610701 |
| Gpn1 | -1.65910637 |
| Mtfr2 | -1.661004853 |
| Phax | -1.664178505 |
| Heatr5b | -1.665941129 |
| Scfd2 | -1.668799668 |
| Tusc3 | -1.675246863 |
| Adamts1 | -1.680794475 |

| | |
|---|---|
| Tcea3 | -1.682898821 |
| Tusc1 | -1.686171712 |
| Ror1 | -1.690089893 |
| Sf3b2 | -1.692001375 |
| Tgif2 | -1.694609762 |
| Ap1s3 | -1.694628698 |
| Thoc2 | -1.700659952 |
| Dennd4c | -1.702913513 |
| Dpy30 | -1.706200271 |
| Tmem248 | -1.713857104 |
| Btbd10 | -1.717295182 |
| Dhx38 | -1.724338697 |
| Rsph9 | -1.725392631 |
| Ifi35 | -1.726383125 |
| Ncor2 | -1.727352964 |
| Rnf121 | -1.728436112 |
| Hopx | -1.733584397 |
| Pitpnm2 | -1.736701071 |
| Tbc1d8 | -1.737420367 |
| Hsd17b12 | -1.743019354 |
| Efcab7 | -1.743170109 |
| Ddi2 | -1.746776347 |
| Yy1 | -1.747267181 |
| Cst3 | -1.748022563 |
| Taf12 | -1.759851178 |
| Ttll5 | -1.765162569 |
| Tmem240 | -1.765178876 |
| Kalrn | -1.765364366 |
| Kif5b | -1.782236642 |
| Megf8 | -1.784817778 |
| Lrch4 | -1.784867394 |
| Polr2j | -1.785173126 |
| Ubxn7 | -1.788139022 |
| Ptprf | -1.790495676 |
| Gosr2 | -1.793177328 |
| Tmc6 | -1.799954729 |
| Mdh1 | -1.801304521 |
| Man2b2 | -1.801679338 |
| Nek5 | -1.805627422 |
| Dbp | -1.809118379 |
| Pgd | -1.812121324 |
| Dusp16 | -1.813113691 |
| Cyb5d1 | -1.814285898 |
| Rtel1 | -1.816338933 |
| Ormdl2 | -1.821291671 |
| Tjp2 | -1.823214884 |
| Mxd4 | -1.824092018 |

| | |
|---|---|
| Rybp | -1.827155149 |
| Triap1 | -1.832269117 |
| Elavl2 | -1.834058183 |
| Prune | -1.837021225 |
| Erp29 | -1.847815069 |
| Nup85 | -1.848235305 |
| Sfswap | -1.854877271 |
| Rad54b | -1.865036323 |
| Gbe1 | -1.870331267 |
| Slc1a1 | -1.87247453 |
| Snx4 | -1.880996288 |
| Pld2 | -1.889099204 |
| Purg | -1.889829765 |
| Ldlr | -1.893909951 |
| Nup107 | -1.895435786 |
| Eprs | -1.897364008 |
| Gabbr2 | -1.90288698 |
| Lgi2 | -1.907776828 |
| Mapkapk2 | -1.912550569 |
| Cd200 | -1.917514229 |
| Uqcr10 | -1.923092252 |
| Enah | -1.924311314 |
| Smim8 | -1.926051589 |
| Fxn | -1.927166522 |
| Slc44a3 | -1.945309686 |
| Ccdc88a | -1.945414956 |
| Mrpl2 | -1.946545027 |
| Ntan1 | -1.947091776 |
| Plcl1 | -1.957631518 |
| Haghl | -1.970529269 |
| Naif1 | -1.99767264 |
| Ccdc18 | -2.000558283 |
| Ppp6r3 | -2.000640739 |
| Sox12 | -2.003839385 |
| Six4 | -2.007917016 |
| Dock5 | -2.013086386 |
| Pmm2 | -2.020062534 |
| Sec63 | -2.031645207 |
| Dync2li1 | -2.042159007 |
| Neurl1b | -2.049325561 |
| Ttc4 | -2.053128677 |
| Sod1 | -2.053665912 |
| Abcd3 | -2.063643353 |
| Tle1 | -2.064902691 |
| Galk2 | -2.070451215 |
| Cpsf2 | -2.081124633 |
| Cps1 | -2.082903998 |

| | |
|---|---|
| Gtf3c5 | -2.092022193 |
| Osbp | -2.094079405 |
| Plk2 | -2.106029881 |
| Fam96a | -2.135922752 |
| Atp5s | -2.137714128 |
| Epha6 | -2.140253855 |
| Tab3 | -2.141692112 |
| Fam213b | -2.148433916 |
| Msn | -2.157212544 |
| Rasgrf2 | -2.162340912 |
| Noc4l | -2.168168479 |
| Polr3g | -2.172019311 |
| Tbx19 | -2.191303557 |
| Mrps24 | -2.193524105 |
| Tm9sf3 | -2.207856285 |
| Secisbp2 | -2.21208104 |
| Slc36a4 | -2.2239627 |
| Coq6 | -2.226563247 |
| Dstn | -2.239616274 |
| Cep57 | -2.247715271 |
| Stam | -2.252739599 |
| Stam2 | -2.270280867 |
| Emilin1 | -2.272217226 |
| Mau2 | -2.277932268 |
| Ptpmt1 | -2.296100611 |
| Clec14a | -2.304017172 |
| Kbtbd3 | -2.306072556 |
| Phc3 | -2.312104005 |
| Aldoa | -2.3126745 |
| Tmem41b | -2.353505555 |
| Ube2v1 | -2.354546299 |
| Ndufa10 | -2.360183586 |
| Eif4g1 | -2.365583845 |
| Tmem9 | -2.407881179 |
| Sf3b4 | -2.422798842 |
| Sbf2 | -2.436402401 |
| Pip5kl1 | -2.448846627 |
| Lgmn | -2.452011289 |
| Cmtm4 | -2.454860847 |
| Ints4 | -2.46508229 |
| Smad5 | -2.486757767 |
| Naa25 | -2.561008981 |
| Eif4ebp3 | -2.561902567 |
| Eef1a2 | -2.578923937 |
| Dcaf11 | -2.590122401 |
| Pkp2 | -2.658451785 |
| Plat | -2.702218124 |

| | |
|---|---|
| Odc1 | -2.722058958 |
| Map10 | -2.728080617 |
| Cdca7l | -2.753104148 |
| Chst14 | -2.769808893 |
| Ksr2 | -2.771837464 |
| Stk38l | -2.791063276 |
| Zc3h4 | -2.7993311 |
| Tmx1 | -2.874000489 |
| Sirt5 | -2.90382763 |
| Mecr | -2.954860293 |
| Kdelc2 | -2.994976492 |
| Nxf1 | -3.119431512 |
| Elmod2 | -3.287353475 |
| Nol9 | -3.334717111 |
| Csrp2bp | -3.34637265 |
| Flnc | -3.355824705 |
| Zbtb40 | -3.362266523 |
| Plin3 | -3.537840332 |
| Slco4a1 | -3.546600587 |
| Kctd10 | -3.669746041 |
| Asnsd1 | -3.680454261 |

**Table S1b. Cell-surface exposed proteins**

| Gene | Cumulative z (top 3) |
|---|---|
| Pcdhga9 | 6.689505294 |
| Itga3 | 6.21238783 |
| Rtn4r | 5.904293171 |
| Ermap | 5.899203615 |
| Itga9 | 4.95604584 |
| Nlgn2 | 4.751228498 |
| Icam1 | 4.742356641 |
| Icam2 | 4.735612438 |
| Fgfr4 | 4.565617434 |
| B2m | 4.47316319 |
| Selp | 4.236192666 |
| Gp1ba | 3.909901227 |
| Eng | 3.674595657 |
| Il1rl1 | 3.600699219 |
| Heg1 | 3.597256533 |
| Tgfbr3 | 3.516658859 |
| B4galt1 | 3.296562189 |
| Enox1 | 3.160935539 |
| Robo4 | 3.022857665 |
| Ly75 | 3.015433449 |

**Table S2. List of sgRNAs and siRNAs.**

| Gene sg/si | Details |
|---|---|
| Mm Pcdhga9 sg pair | GGGAGCCAGTTCCACACCAG; GTATCTGGTGACCAAGGTGG |
| Mm Pcdhga9 si | Horizon Discovery, L-046725-01-0005 |
| Hs Pcdhga9 si | Horizon Discovery, L-013270-02-0005 |
| Hs Pcdhg si#1 | Custom si, Horizon Discovery, CCGCCCAACACGGACTGGCGTTT |
| Hs Pcdhg si#2 | Custom si, Horizon Discovery, CCCAACAACCAGTTTGACACAGA |
| Hs Klf2 si | Horizon Discovery, L-006928-00-0005 |
| Hs Klf4 si | Horizon Discovery, L-005089-00-0005 |
| Hs RBPJ si | Horizon Discovery, L-007772-00-0005 |

**Table S3. List of primary antibodies.**

| Antibody | Company, Catalog No. | Dilution for IF | Dilution for immunoblot |
|---|---|---|---|
| GFP | ThermoFisher Scientific, A-11122 | 1:1000 | 1:4000 |
| VCAM1 | Abcam, ab134047 | - | 1:4000 |
| Tubulin | Millipore Sigma, 05-829 | - | - |
| Pcdhg | ThermoFisher Scientific, MA5-27615 | 1:200 | 1:2000 |
| Klf4 | Abcam, ab215036 | 1:500 | 1:4000 |
| GAPDH | Cell Signaling, 5174 | - | 1:8000 |
| CD68 | ThermoFisher Scientific, 14-0681-82 | 1:200 | - |
| FLAG | Millipore Sigma, F1804 | 1:400 | 1:2000 |
| FLAG | Cell Signaling, 14793 | 1:400 | 1:2000 |
| RBPJ | Cell Signaling, 5313 | - | 1:2000 |
| NICD (Val 1744) | Cell Signaling, 4147 | - | 1:2000 |
| mAb A9 (0.1 mg/ml) | *This study* | 1:100 | 1:500 |
| mAb B1 (0.1 mg/ml) | *This study* | 1:100 | 1:500 |
| mAb B4 (0.1 mg/ml) | *This study* | 1:100 | 1:500 |
| Rat IgG (1.0 mg/ml) | Vector Labs, I-4000-1 | - | - |
| SMA | Millipore Sigma, A2547 | 1:500 | - |
| Erg | Abcam, ab92513 | 1:800 | - |

**Table S4. List of RT-PCR primers.**

| Gene | Forward primer | Reverse Primer |
|---|---|---|
| Mm Pcdhga9 | GCCTAAGGAGTTAGCGAGCA | TGGCACGGAATAGCGGATTT |
| Mm GAPDH | GGGTCCCAGCTTAGGTTCATC | TACGGCCAAATCCGTTCACA |
| Mm Klf2 | AAGAGCTCGCACCTAAAGGC | CTTTCGGTAGTGGCGGGTAA |
| Hs Klf2 | CGGCAAGACCTACACCAAGA | TGGTAGGGCTTCTCACCTGT |
| Hs Klf4 | GTGGAGAAAGATGGGAGCAG | TGACTTTGGGGTTCAGGTG |
| Hs GAPDH | TGCACCACCAACTGCTTAGC | GGCATGGACTGTGGTCATGAG |
